# Chemoenzymatic Synthesis of Glycosphingolipids having an HNK-1 Epitope for Erythrocyte Cell Surface Remodeling

**DOI:** 10.1101/2025.03.10.642389

**Authors:** Mehman I. Bunyatov, Geert-Jan Boons

**Affiliations:** Chemical Biology and Drug Discovery, Utrecht Institute for Pharmaceutical Sciences, and Bijvoet Center for Biomolecular Research, Utrecht University, 3584 CG Utrecht, The Netherlands; Complex Carbohydrate Research Center, University of Georgia, Athens, Georgia 30602, United States; Department of Chemistry, University of Georgia, Athens, Georgia 30602, United States

## Abstract

Several neuropathies, such as Guillain-Barre syndrome and myelin-associated glycoprotein neuropathy (MAG), are caused by antibodies targeting glycosphingolipids. Several studies have indicated that MAG arises from pathogenic IgM autoantibodies targeting sulfoglucuronyl (HNK-1) containing glycosphingolipids. The exact mechanism by which IgM-neuropathy occurs has not been fully elucidated. Furthermore, no appropriate diagnostic tools are available for MAG using sulfoglucuronyl containing glycosphingolipids. To address these limitations, we describe here a synthetic strategy that makes it possible to prepare sulfoglucuronyl paraglobosides using a *neo*chemo-enzymatic approach. It is based on the enzymatic assembly of *N*-acetyl-lactosamine (LacNAc) backbones as thioglycosides that were subjected to protecting group manipulations to give glycosyl acceptors for the chemical installation of a sulfated glucuronic acid moiety. A late-stage conversion of the thioglycosides into anomeric fluorides made it possible to enzymatically introduce sphingosine. The resulting compounds were acetylated to provide 3-sulfo-glucuronyl- and glucuronyl-containing glycosphingolipids, respectively. The glycosphingolipids were employed to remodel the surface of erythrocytes to examine complement mediated toxicity by an anti-HNK-1 antibody. It was found that erythrocytes remodeled with exogenous administered HNK-1 containing glycosphingolipid undergo complement dependent lysis when incubated with an IgM anti-CD57 IgM antibody, whereas a compound lacking a sulfate was not able to induce this effect. The approach could be extended to the gangliosides GM1a and GD1a, which have been implicated in Guillain-Barre syndrome. The results highlight that cell surface remodeling will be attractive for diagnosis, disease monitoring and immunological research of diseases associated with pathogenic antibodies targeting glycosphingolipids.

**TOC graphic:** 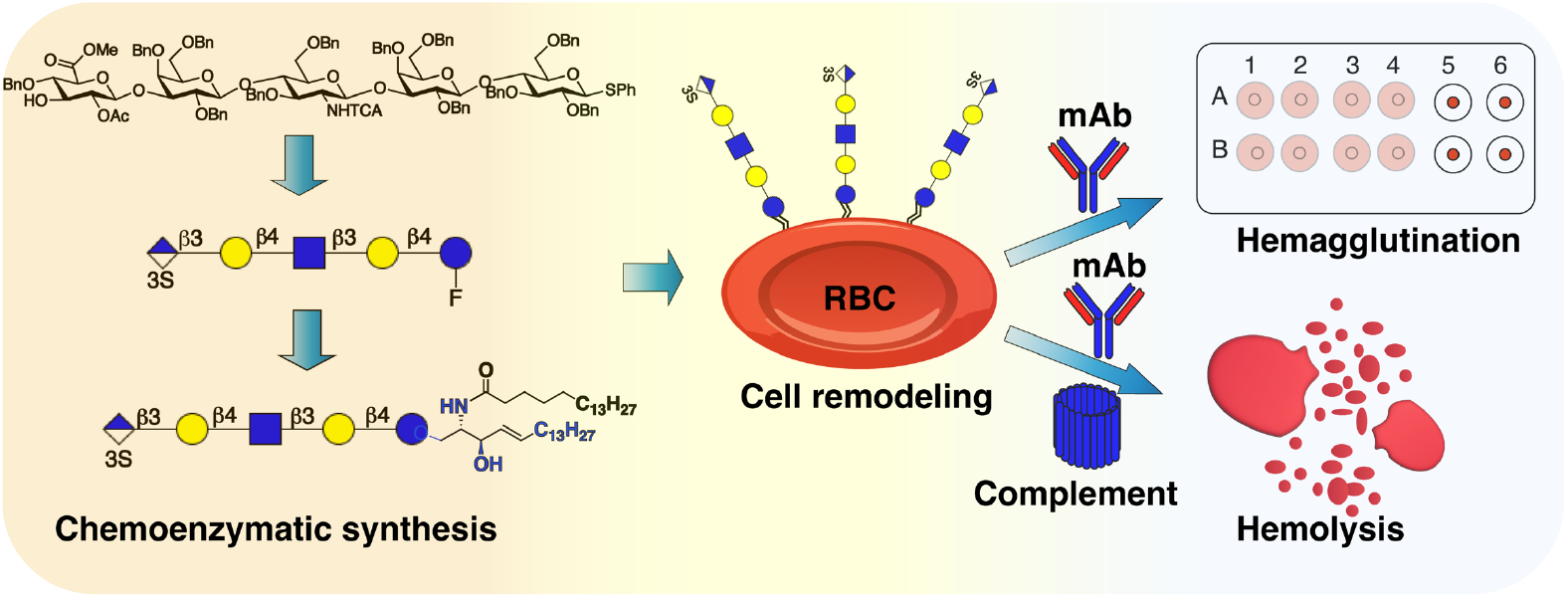

## INTRODUCTION

Human peripheral nervous tissues are rich in acidic glycosphingolipids which are mainly sialic acid containing gangliosides, paraglobosides and their sulfated derivatives.^1^ The spatial organization on the plasma membrane and unique physicochemical properties of glycosphingolipids allow these biomolecules to act as mediators of a wide range of biological processes such as cell-cell communication, immune regulation, signal transduction including regulation of protein tyrosine kinase activity and cellular interactions.^2^ Glycosphingolipids can also be targeted by pathogenic antibodies,^3^ and for example antibodies recognizing gangliosides, which are abundantly expressed in the central nervous system, can lead to neuropathies such as Guillain-Barre syndrome.^4^ The peripheral nervous system is rich in glycosphingolipids of the neolacto series that can be decorated with epitopes such as Lewis^x^, CD75 and human natural killer-1 (HNK-1), (Figure 1A). These structures have also been implicated in autoimmune disorders and one such autoimmune neuropathy involves the interactions of IgM antibodies with HNK-1 bearing paraglobosides (compounds **1a**,**b**, Figure 1B).^5^ The resulting IgM monoclonal gammopathy, which is also known as myelin-associated glycoprotein (MAG) neuropathy, leads to slow but progressive destruction of the myelin layer of nerve cells manifesting in sensory ataxia with impaired gait, tremor, distal muscle weakness and neuropathic pain eventually leading to severe disabilities. The HNK-1 epitope is a trisaccharide composed of a 3-O-sulfated glucuronic acid linked to N-acetyl-lactosamine (HSO_3_-3GlcAß1-3Galß1-4GlcNAc-). Earlier studies have showed that disease progression arises from direct targeting of pathogenic IgM autoantibodies to HNK-1 containing glycoconjugates. However, the exact mechanism by which IgM-neuropathy occurs has not been fully elucidated. Experiments conducted using feline animal models have indicated that in addition to antibodies, complement factor proteins are involved in disease manifestation. Fresh patient serum supplemented with external complement induced extensive demyelination of cat peripheral nerves. Passive immunizations using sera without complement factors or frozen serum with complement components induced minor or no demyelination.^6,7^ Moreover, nerve biopsies obtained from patients indicated the presence of IgM and complement C3d depositions on myelinated fibers.^8,9^ Even though these experimental data implicate complement activation during IgM targeted demyelination, there is no direct evidence at the molecular level for harnessing complement proteins by anti-HNK-1 autoantibodies. A platform is needed that can determine the involvement of specific glycans, antibody as well as complement in disease manifestation. Such a platform may also find use in the diagnosis of IgM monoclonal gammopathies, which for MAG has been fraught with many difficulties.^10^ In this respect, currently employed Enzyme-Linked Immunosorbent Assay (ELISA) are based on the use of a HNK-1 containing *N*-glycans of myelin associated glycoprotein, which do properly resemble HNK-1 presented on glycosphingolipids. ELISA is performed at 4 °C that does not necessarily reflect the binding activity *in vivo*. The severity of the neuropathy also depends likely to depend on other effector mechanisms that are not measured in these assays.

**Figure 1.**
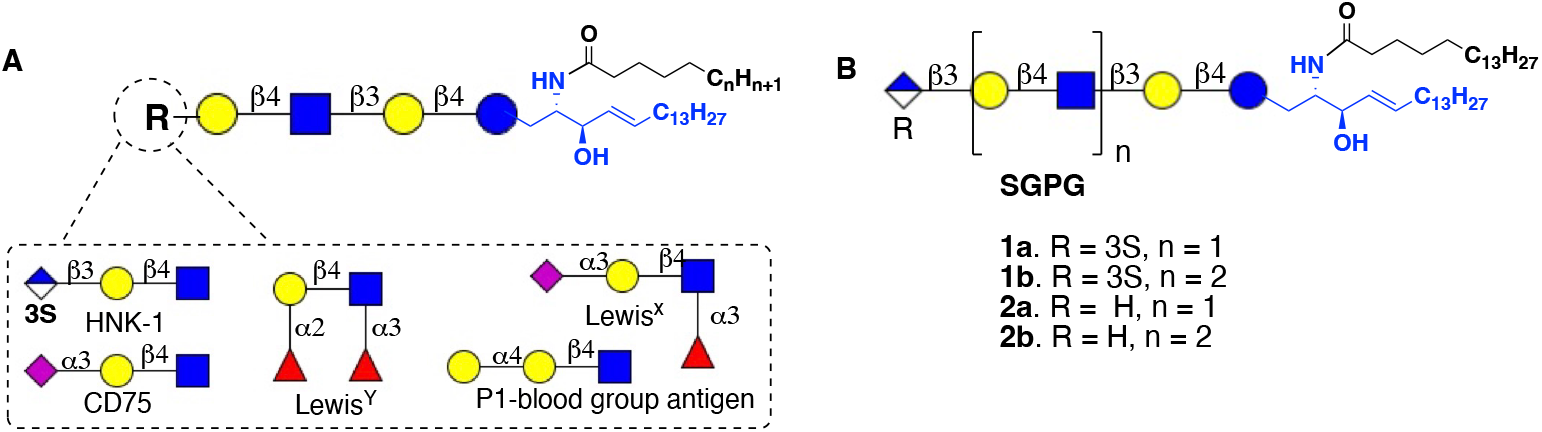
**A.** Core *neo-*lacto glycosphingolipids series with diverse functional epitopes B. Target library of synthetic sulfo glucuronyl paragloboside glycolipids.

Previously, we described binding preferences of serum IgM autoantibodies of MAG patients for sulfoglucuronyl and sulfoglucuronyl lactosaminyl paraglobosides using synthetic glycans immobilized on microarray slides.^11^ The synthetic glycans were devoid of a ceramide moiety and therefore could not be employed in functional studies. To address this limitation, we describe here, for the first time, a synthetic strategy for the preparation of sulfoglucuronyl paraglobosides. It is based on a neochemo-enzymatic approach in which *N*-acetyl-lactosamine (LacNAc) backbones are enzymatic assembled that are then subjected to protecting group manipulations to give a glycosyl acceptor for chemical installation of a sulfated glucuronic acid moiety. A key feature of the synthetic approach was a late-stage conversion of a thioglycoside into an anomeric fluoride which made it possible to enzymatically introduce sphingosine. The resulting compounds were acetylated to provide the targeted 3-sulfo-glucuronyl and glucuronyl containing glycosphingolipids **1a** and **2a** (Figure 1B), respectively. The latter compounds were employed to remodel the cell surface of erythrocytes to examine complement mediated hemolysis by anti-HNK-1 antibodies. It was found that cell surface remodeling of erythrocytes with exogenous administered HNK-1 containing glycosphingolipid **1a** resulted in complement dependent lysis when incubated with an IgM anti-CD57 IgM antibody, whereas compound **2a** lacking a sulfate was not able to induce this effect. The cell surface remodeling and complement mediated lysis could be extended to the gangliosides GM1a and GD1a in the presence of appropriate antibodies. The results support that auto-antibodies cause cell damage in a complement mediated manner. Furthermore, it highlights that the cell surface remodeling approach will be attractive for diagnosis, disease monitoring and immunological research of diseases associated with pathogenic antibodies targeting glycosphingolipids.

## RESULTS AND DISCUSSION

### Chemical Synthesis of HNK-1 Containing Anomeric Fluoride

The amphiphilic properties of glycosphingolipids make this class of compounds not only biologically important but also synthetically challenging.^12^ In particular, the presence of a hydrophobic ceramide moiety complicates chemical or chemoenzymatic extensions of lactosyl ceramide to give more complex glycosphingolipids. To address this challenge, late-stage installation of sphingosine using a mutant endoglycoceramidase enzyme (EGC) was introduced.^13^ It utilizes a-lactosyl fluoride that enzymatically is extended to a more complex glycan for subsequent condensation with sphingosine using a mutant EGC. This approach can, however, not easily be adapted to the preparation of glycosphingolipids that are equipped with epitopes that cannot be assembled by enzymatic procedures alone such as HNK-1 containing compounds. In this respect, the chemical lability of anomeric glycosyl fluorides limits many chemical manipulations, and for example is not compatible with commonly used glycosylation conditions such as the use of Lewis acids for activation of glycosyl donors.^14,15^ Late-stage installation of fluoride is also difficult because it requires acidic conditions that may not be compatible with sensitive functionalities such a sialosides and sulfates.

Previously, it was shown that Barluenga’s reagent (IPy_2_BF_4_) in a combination with hydrofluoric acid in pyridine can convert thioglycosides having a free hydroxyl, into the corresponding glycosyl fluorides with high a-anomeric selectivity.^16^ Thus, we anticipated that treatment of thioglycoside 4 with IPy_2_BF_4_ and HF×Py would give the anomeric fluoride 5 (Figure. 2). The latter compound has a free C-3 hydroxyl and sulfation with SO_3_×Py followed by deprotection was expected to yield 6. This compound can then serve as glycosyl donor for enzymatic installation of sphingosine using a mutant endoglycoceramidase^17^ to yield lyso-sulfoglucuronyl paragloboside analogues. Acylating of the amine of the sphingosine moiety with stearoyl succinimide would then provide target compound **1a**. A similar sequence of reactions without the sulfation step was expected to provide **2a**. Thiophenyl lactoside **3** was chosen as the starting glycan for assembly of key intermediate **4**.

**Figure 2.**
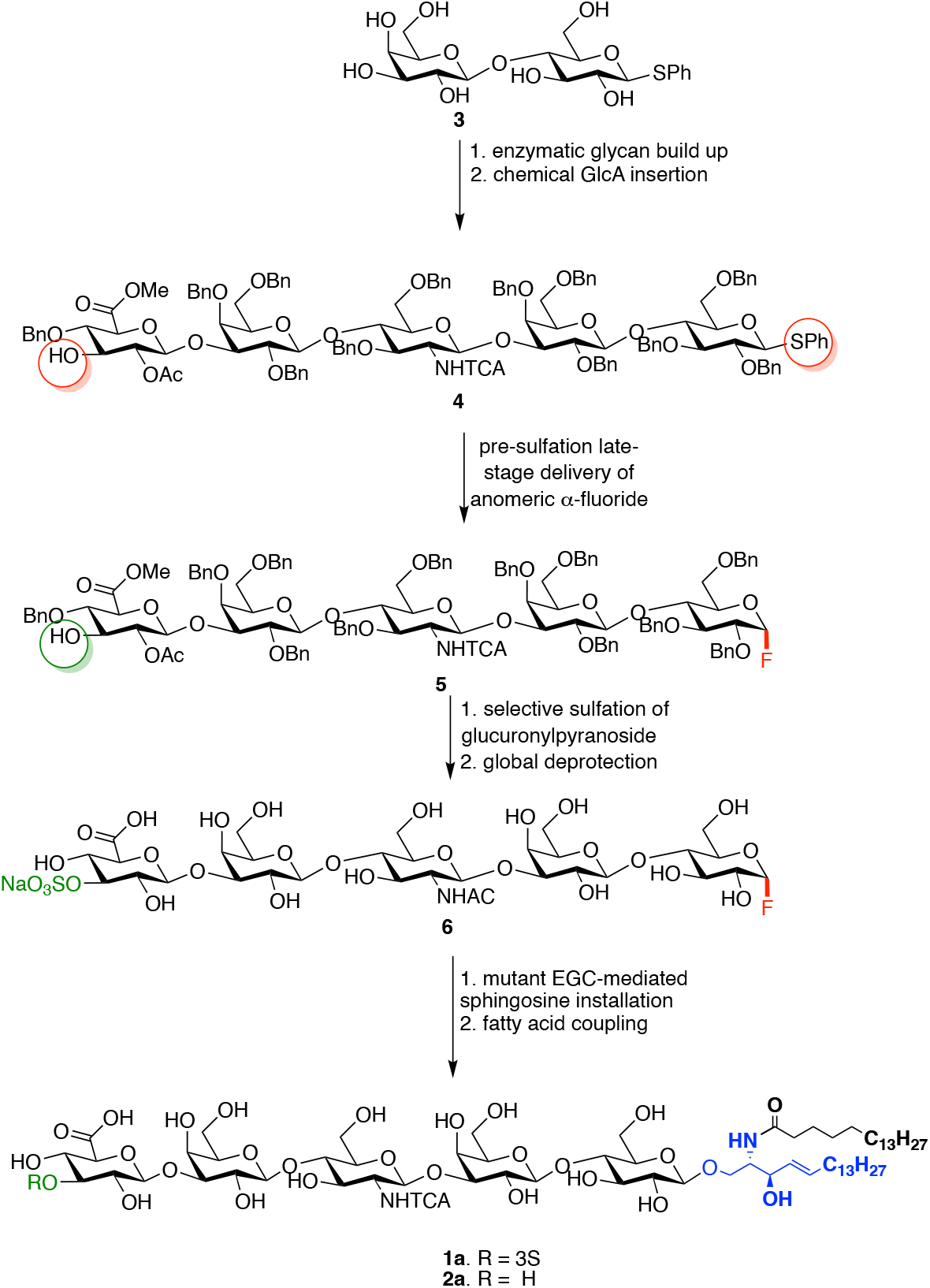
Key synthetic steps for the *neo*chemoenzymatic synthesis of glucuronyl paraglobosides **1a** and **2a**.

Phenyl-1-ß-thiolactoside **3** was prepared from per-*O*-acetylated-ß-lactoside, which was converted into lacto-*N*-neotetraose **7** by the sequential addition of *N*-trifluoroacetylglucosamine (GlcNHTFA) and galactose (Gal), using UDP-GlcNHTFA and UDP-Gal as nucleotide-sugar donors and the microbial glycosyltransferases HpB3GnT^18^ and NmLgtB,^19^ respectively. The resulting tetrasaccharide **7** was subjected to aqueous NaOH to remove the TFA moieties, and the amines of resulting compound **8** were converted into azides by an azido-transfer reaction using imidazolonium sulfonyl azide^20^ to give **9** (Figure 3).

**Figure 3.**
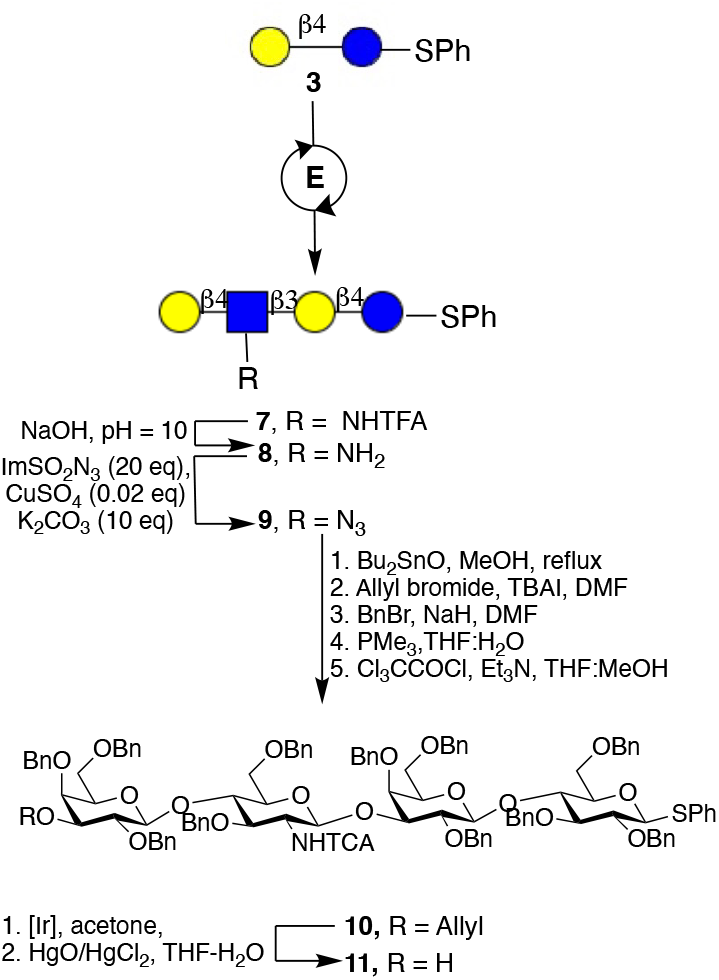
Chemoenzymatic assembly of properly protected thioglycoside **11**. E = UDP-GlcNHTFA and HpB3GnT and then UDP-Gal and NmLgtB.

Previously, we demonstrated that the C-3 hydroxyl of a terminal galactoside of an oligo-LacNAc derivative can selectively be protected as an allyl ether by formation of a dibutyl stannelene complex followed by reaction with allyl bromide.^11^ The application of these reaction conditions resulted in the selective allylation of the C-3 hydroxyl of the terminal galactoside of **9**. The remaining alcohols were protected as benzyl ethers by treatment with benzyl bromide and NaH in DMF, which was followed by PMe_3_ mediated azide reduction for subsequent acylation with trichloroacetyl chloride to give fully protected **10**. The latter compound was converted into glycosyl acceptor **11** by a two-step procedure involving isomerization of the allyl into a vinyl ether using hydrogen activated (1,5-cyclooctadiene) bis(methyldiphenylphosphine) iridium(I) hexafluorophosphate in THF for 18 h,^21^ which was hydrolyzed using a mixture of HgO/HgCl_2_. The removal of allyl ether was monitored by TLC and confirmed by MALDI-TOF MS and ^1^H-NMR spectroscopy which showed loss of a characteristic peak at 5.5 ppm. The use of PdCl_2_ in methanol to remove the allyl ether, which was successfully employed for *O*-glycosides, failed for **11** which is likely due to catalyst poisoning by the thioglycoside.

Next, oligosaccharide acceptor **11** was glycosylated with glucuronic acid donor **12** in the presence of TMSOTf (1.5 eq of per GlcA donor) in DCM at -50 ^0^C to give, after a reaction time of 2 h, glucoronate **13** in high yield (Figure 4). The proton at C-3 of the terminal galactoside showed a downfield chemical shift in the ^1^H-NMR spectrum and gave a through space correlation (NOEsy) with the anomeric H-1 of glucuronic acid indicating the formation of the desired glycosidic linkage. Compound **13** was subjected to KOH (1 M) in a mixture of dioxane and H_2_O to hydrolyze the methyl ester. The resulting solution was concentration under reduced pressure and the resulting residue was dissolved in methanol to remove the benzoate esters. Next, the crude product was dissolved in acetic anhydride and heated to 85 ^°^C to convert the glucuronic acid moiety into the corresponding 3,6-lactone. This was followed by cooling the reaction mixture to room temperature and addition of pyridine, which resulted in acetylation of the remaining hydroxyl to yield compound **14**. The lactone of the latter compound was hydrolyzed under mild basic conditions using a methanolic solution of NaOAc to give compound **4** that has a free 3-OH at glucuronic acid. The identify of this compound was confirmed by MALDI-TOF MS and homo- and hetero nuclear NMR, which showed the presence of the 3-OH of GlcA and a shift of H-3_GlcA_ to a more up field region (see Figure S2). With compound **4** in hand, attention was focused on the selective installation of an anomeric fluoride with a-anomeric configuration leaving the hydroxyl at C-3 of GlcA free for subsequent sulfation. Thus, the thiophenyl glucoside of **4** was activated with Barluenga’s reagent (2.0 equiv.)^16^ at -40 ^°^C and the resulting intermediate subjected to fluoride displacement by treatment with an excess (20 equiv.) of HF/Py complex in dichloromethane. The progress of fluoride installation was monitored by TLC (R_*f*_ *=* 0.6 →R_*f*_ = 0.5, Tol:Acetone = 7:1) and formation of **5** was confirmed by MALDI-TOF MS. ^1^H NMR spectroscopy of the resulting compounds showed the absence of characteristic H-1_Glc_ of thioglycosides, and instead a new signal for H-1_Glc_ was observed as a doublet of doublets at 5.4–5.3 ppm with coupling constants of *J*_1_ = 2.7 Hz, *J*_2_ = 53.3 Hz indicating the formation of the expected *α*-anomeric fluoride (Figure S3). ^19^F NMR also confirmed the formation of the a-anomeric fluoride with a characteristic peak at -149.7 ppm (dd, *J*_1_ = 25.6 Hz, *J*_2_ = 53.1 Hz).^16^ H-2_GlcA_ and H-3_GlcA_ did not exhibit a change in chemical shift in the ^1^H NMR spectrum, indicating that neither fluorination of 3-OH_GlcA_ nor acetyl migration from C2 to C3 of glucuronic acid had occurred. The hydroxyl of compound **5** was sulfated using an excess of SO_3_×Py complex in pyridine. The reaction was quenched by the addition of methanol and the sulfate ester was subjected to ion exchange using Dowex-Na^+^ ion exchange resin. The resulting compound was hydrogenated over Pd(OH)_2_/C (20% wet, Degussa type) to remove the benzyl ethers and reduce the NHTCA moieties to acetamides. Gratifyingly, the anomeric fluoride had stayed intact during the hydrogenation, and no dehalogenation was detected by LC-MS. Finally, the methyl ester and acetyl esters at C-6 and C-2 of glucuronic acid, respectively, were hydrolyzed under mild basic conditions (pH = 7.8) to give compound **6**. Similarly, compound **5** was subjected to global deprotection to yield unsulfated glycosyl fluoride **15**.

**Figure 4.**
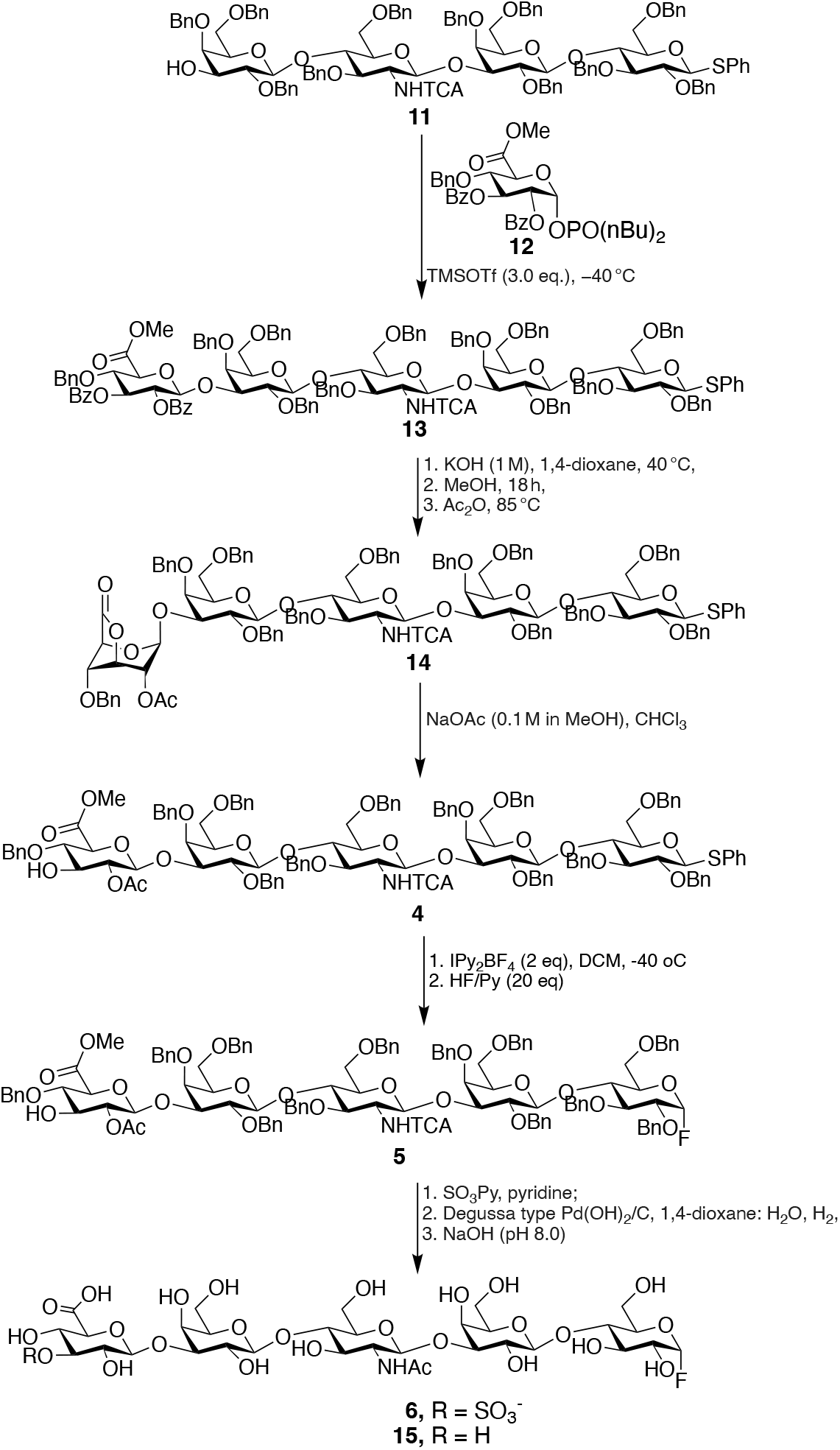
Synthesis of glycosyl fluorides containing an HNK-1 or glucuronyl moiety.

### Enzymatic *En Bloc* Transfer of Anomeric Fluorides to Sphingosine

Enzymatic transfer of glycosyl fluorides to sphingosine is a versatile approach for the preparation of glycosphingolipids (GSLs) and has been applied to the synthesis of several gangliosides (GM3, GM1), globosides (Gb3), and lactosides (LnNT, Pk-antigen) using a mutant endoglycoceramidase obtained from *Rhodococcus* strain M-777.^13,22^ Here, we examined the feasibility of transfer of chemoenzymatically synthesized sulfogluronyl paragloboside fluorides to sphingosine using a double mutated endoglycoceramidase to yield lyso-sulfoglucuronyl paraglobosides that can be converted to glycosphingolipids by acylation of the amine of the sphingosine moiety. Thus, erythro-sphingosine was prepared from commercially available phytosphingosine in an overall yield of 15% over six steps.^23^ Previously, different versions of endoglycoceramidase enzymes have been engineered to broaden its substrate promiscuity^22,24^ and in particular the double mutant EGC E351S/D314Y showed increased efficiency over the single mutant parent EGC E351S in analytical scale synthesis of glycosphingolipids. Furthermore, dimethoxyethane (DME) was shown to be a superior detergent to solubilize sphingosine. Thus, glycans **6** and **12** were condensed with sphingosine (1.5 equiv.) using endoglycoceramidase (EGC II D351S/E314Y) in NaOAc buffer (pH = 5.0, 50 mM) in the presence of 1% (v/v) DME to yield *lyso* forms of glycosphingolipids **14** and **15**, respectively. The progress of the reactions was monitored by MALDI-TOF MS until complete consumption of the glycosyl fluorides was observed. The products were purified over a reverse C18-cartridge to remove traces of hydrolyzed glycosyl donor and unreacted sphingosine. LC-MS and ^1^H NMR spectroscopy confirmed the structural integrity of the compounds (Figure 5). Next, the amine of the sphingosine moiety of **14** and **15** was acylated to provide the targeted glycosphingolipids. The composition of fatty acid of naturally occurring glycosphingolipids is heterogenous, ranging from stearic acid (C18) to lignoceric acid (C24) with variable degrees of unsaturation.^25^ The most abundant form is, however, stearic acid. Thus, stearic acid was activated as an *N*-succinimide ester by treatment with *N*-hydroxy succinimide (NHS) in the presence of 1-ethyl-3-(3-dimethylaminopropyl)carbodiimide in DCM. The resulting stearyl-NHS precipitated in DCM and used immediately in subsequent acylation reaction. Next, compounds **14** and **15** were reacted with stearyl-HNS in a mixture of THF and MeOH (3:1) in the presence of Et_3_N to obtain, after purification by C8 reverse phase chromatography, the sulfated and unsulfated glucuronyl paraglobosides **1a** and **2a** (Figure 5).

**Figure 5.**
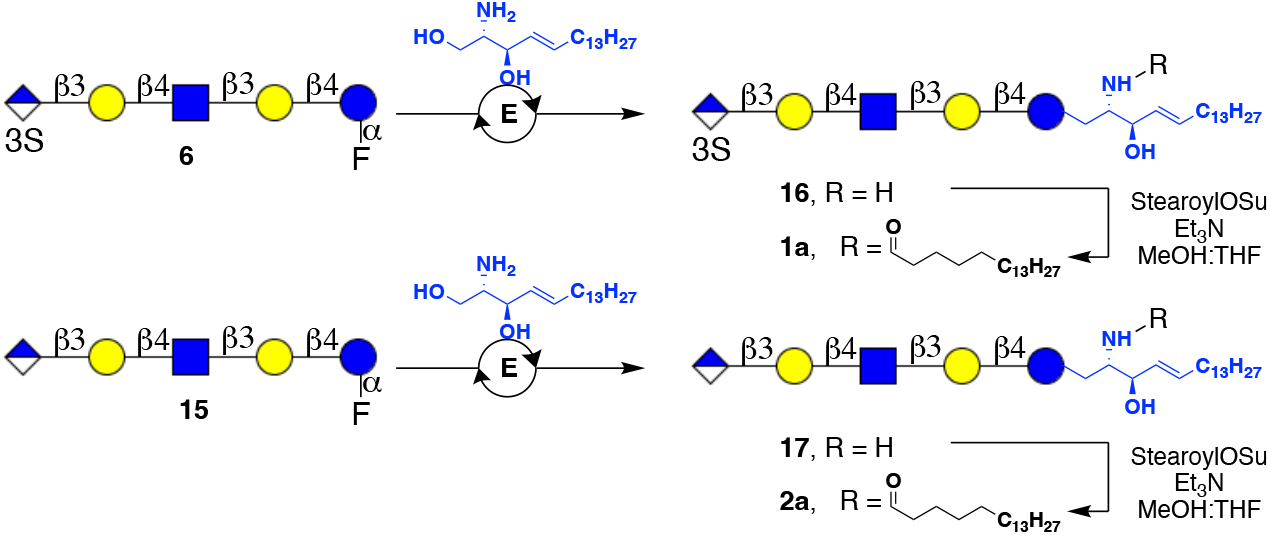
*En block* enzymatic transfer of *α*-glycosyl fluorides to sphingosine and subsequent fatty acid acylation to give sulfated (**1a**) and non-sulfated (**2a**) paraglobosides. Conditions: E: endoglycoceramidase (EGC II D351S/E314Y), NaOAc buffer (pH = 4.5 – 5.5), 37 ^°^C.

We extended the *neo*chemo-enzymatic approach to the preparation of glucuronyl lactosamine paraglobosides **1b** and **2b** with and without a sulfate moiety, respectively (see Schemes S4 and S5 for details). Compound **7** was elongated by an additional LacNAc moiety, and the resulting compound was then subjected to the established protecting group manipulation to facilitate the chemical installation of a glucuronic acid moiety. Anomeric fluorination and sulfation gave a heptasaccharide that was transformed into a lyso-glucuronyl lactosamine paraglobosides. This compound was subsequently acylated with stearic acid, resulting in sulfated glucuronyl lactosamine paraglobosides **1b**. In a similar way, compound **2b** lacking a sulfate was prepared demonstrating the power of the methodology.

### Glycoengineering of Erythrocytes by Glycosphingolipids 1a and 2a to Examine Agglutination and Complement-Mediated Hemolysis

Red blood cells (RBCs) are prone to complement-mediated cell lysis releasing hemoglobin that can be measured by spectrophotometry. This unique feature of erythrocytes makes them well suited to examine complement and antibody dependent cytotoxicities.^26,27^ Thus, we were compelled to investigate the effector function of an IgM anti-HNK1 monoclonal (anti-CD57) antibody on erythrocytes that exogenously were exposed to glycosphingolipids **1a** and **2a** for incorporation of these lipids in their cell membranes. To assess the incorporation efficiency of glycosphingolipids **1a** and **2a** into cell membranes (hence glyco-remodeled cells, RBCg), turkey erythrocytes were first tested for hemagglutination by an anti-CD57 IgM antibody. Thus, fresh turkey RBCs (25% suspension) were incubated with glycosphingolipids **1a** and **2a** (1 mg/mL) at 37 ^°^C for 1 h and then washed twice with PBS and then incubated with anti-CD57 antibody using serial dilutions from 100 *μ*M to 0.1 nM for 30 min at room temperature. Erythrocytes decorated with sulfated glucuronyl paragloboside **1a** could be agglutinated with anti-CD57 IgM antibody at a concentration as low as 10 nM, however, no agglutination was observed for erythrocytes that were remodelled with non-sulfated bearing glycosphingolipid **2a**. Moreover, prolonged incubation of erythrocytes with the same concentration of glycosphingolipids at 37 ^°^C did not markedly improve the hemagglutination efficiency. In contrast, RBCs that were incubated with glycosphingolipid **1a** at a lower temperature (22 ^°^C) or reduced concentration (0.1 mg/mL) required a higher antibody concentration for agglutination, indicating the density of the incorporated glycolipids influences the binding of the IgM antibody (see Figure S6 for details).

Several studies have indicated that exogenously administered gangliosides can bind to the cell-surface proteins and only partially be incorporated into membranes.^28^ To account for this possible effect, RBCs remodeled with compound **1a** were treated with 0.1% freshly prepared fetal calf serum to remove not properly incorporated compounds. In addition, the erythrocytes were treated with 1% trypsin solution to remove the protein bound **1a**, thus exposing only glycolipids that are properly incorporated into cell-membranes. RBCs treated with serum did not show any change in agglutinating, while trypsin treatment exhibited agglutination at a slightly lower antibody concentration, presumably due to reduced shielding of neighboring proteins exerted on the incorporated gangliosides. Thus, these results indicate that most of the gangliosides were properly incorporated into the cell membrane. It is worth noting that trypsinization followed by the glycosphingolipid **1a** incorporation required a higher anti-CD57 antibody concentration for agglutination. This observation indicates that the exogenously administered glycosphingolipids first encounter cell-surface proteins, which facilitate proper incorporation into the cell membrane.

Next, we investigated effector functions of the anti-CD57 IgM antibody using the glycosphingolipid remodeled erythrocytes. Thus, RBCs were remodeled with compounds **1a** and **2a** and then exposed to the anti-CD57 IgM antibody followed by treatment with guinea pig serum complement to induce complement dependent hemolysis (Figure 6A). Unmodified erythrocytes showed almost no hemolysis at a wide range of antibody concentrations indicating that these cells do not express appropriate receptors. Modification of erythrocytes with non-sulfated glucuronyl paragloboside **1b** gave similar results. On the other hand, RBCs exposed to HNK-1 containing glyosphingolipid **1a** demonstrated hemolysis in the presence of serum complement. The degree of hemolysis was dependent on the anti-CD57 IgM antibody concentration, starting at sub-nanomolar and reaching a maximum at micromolar concentration. The same erythrocytes did not exhibit substantial hemolysis in the absence of serum complement, indicating that the lysis is complement dependent.

**Figure 6.**
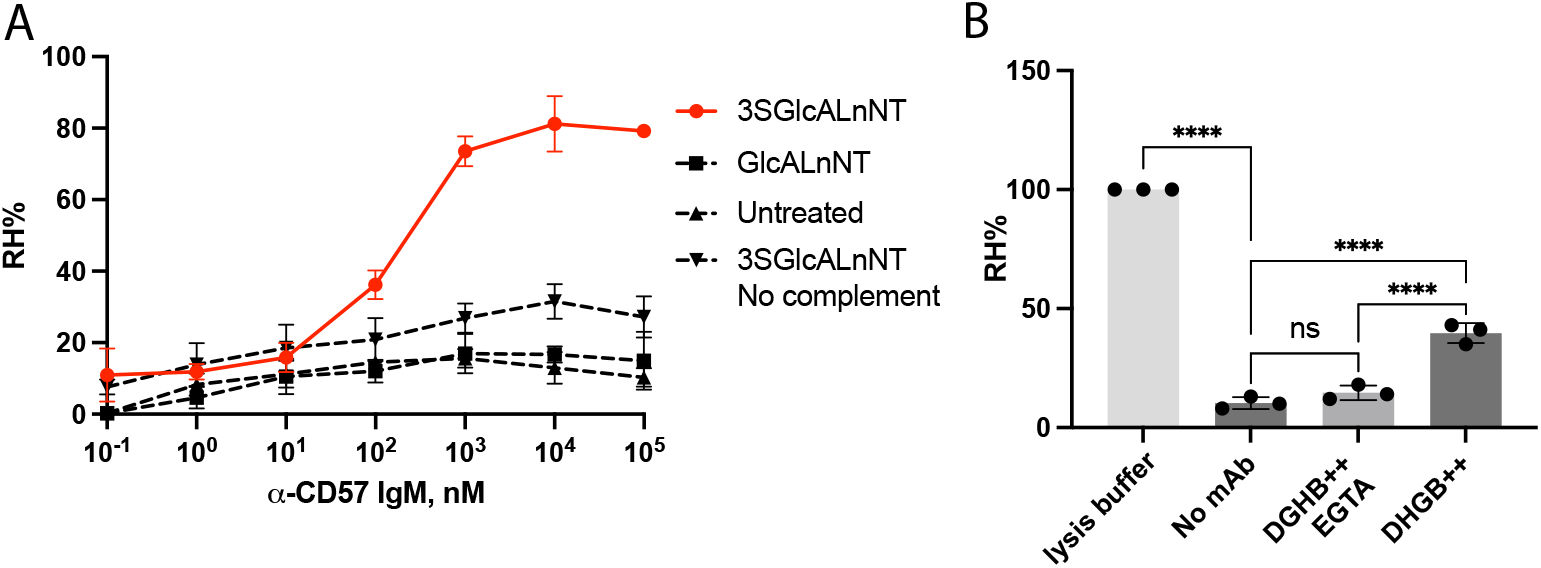
Complement-dependent hemolysis of glucuronyl paragloboside remodeled erythrocytes by anti-HNK-1 IgM antibody. (A) Sulfoglucuronyl paragloboside remodeled erythrocytes undergo hemolysis in the presence of anti-CD57 (anti-HNK-1) antibody and guinea pig serum complement (EC_50_ = 163.7 nM). (B) Complement activation is inhibited in the absence of Ca^2+^, indicating anti-CD57 IgM antibodies (100 nM) involves the activation of complement via classical pathway. Statistical significance was calculated using ordinary one-way ANOVA. *P* values are determined as <0.0001 for ****, ns means not significant.

To assess whether the antibody mediated cell lysis involves complement activation *via* the classical or alternative pathway, we further tested two different conditions for hemolysis (Figure 6B). After glycolipid incorporation to give tRBC^g^, the cells were stored in DGH (Dextrose-Gelatin-Hepes) buffer supplemented with Mg^2+^ and Ca^2+^ ions and used for the subsequent lysis experiments. In addition, the tRBC^g^ in storage media was supplemented with ethylene glycol-*bis*(β-aminoethyl)-*N,N,N’,N’*-tetraacetic acid (EGTA) which chelates Ca^2+^ ions while keeping the Mg^2+^ ions free. Next, the cells were treated with the anti-CD57 IgM antibody (100 nM) and guinea pig serum complement. tRBC^g^ containing both calcium and magnesium ions. Inhibition of hemolysis was observed for erythrocytes sensitized with EGTA, indicating that the complement activation follows the classical pathway which is abolished in the absence of calcium ions.

### Glycoengineering of Erythrocytes with the Gangliosides GM1a and GD1a for Assessing Complement-Mediated Hemolysis by Anti-Ganglioside Antibodies

Next, we extended the cell surface remodeling approach to GM1a and GD1a gangliosides and demonstrated selective agglutination and hemolysis of the resulting erythrocytes by anti-GM1a and anti-GD1a IgG antibody, respectively. These gangliosides have been implicated in Guillain-Barré Syndrome, which is immune disorder that is associated with *Campylobacter jejuni* infections.^29^ It results in inducing antibodies to lipo-oligosaccharide (LOS) of this gram-negative bacterium that cross-react with gangliosides at peripheral nerves causing polyneuropathy^4^. Glycolipidomic studies of human erythrocytes have indicated the presence of small amounts of GM1 and no detectable amounts of GD1a gangliosides. However, the presence of GM1 and GD1a gangliosides on the surface of fowl erythrocytes has not been established yet. The cell surface of turkey erythrocytes was remodeled with exogenously administered GM1a or GD1a ganglioside followed by a hemagglutination assay in the presence of anti-GM1a and anti-GD1a IgG antibodies. Since IgG antibodies are less effective in agglutinating than IgM, a secondary anti-IgG antibody was added to facilitate lattice formation. Ganglioside remodeled erythrocytes were agglutinated in the presence of appropriate anti-ganglioside antibodies and secondary anti-IgG monoclonal antibody (see Figure S7), however no hemagglutination was observed for erythrocytes lacking exogenously administered gangliosides.

Next, attention was turned to hemolysis of erythrocytes remodeled with GM1 and GD1a gangliosides in the presence of anti-ganglioside IgG monoclonal antibodies and guinea-pig serum complement. Native erythrocytes did not show any hemolysis with the antibodies in the presence of guinea pig serum complement. GM1a decorated erythrocytes showed significant hemolysis with a wide range concentration of anti-GM1a IgG antibody in the presence of serum complement (Figure 7A). Similarly, erythrocytes remodeled with GD1a exhibited cell lysis in the presence of the anti-GD1a antibody and serum complement (Figure 7B). The lysis was very selective and GM1a remodeled erythrocytes did not undergo substantial cell lysis in the presence of anti-GD1a antibodies and complement, and the anti-GM1a antibody did not induce lysis of GD1a coated red-blood cells. Thus, these results demonstrate the absence of cross-reactivity of the antibodies. Moreover, none of the ganglioside remodeled erythrocytes were lysed in the absence of the serum complement with appropriate antibodies supporting a complement dependent mechanism.

**Figure 7.**
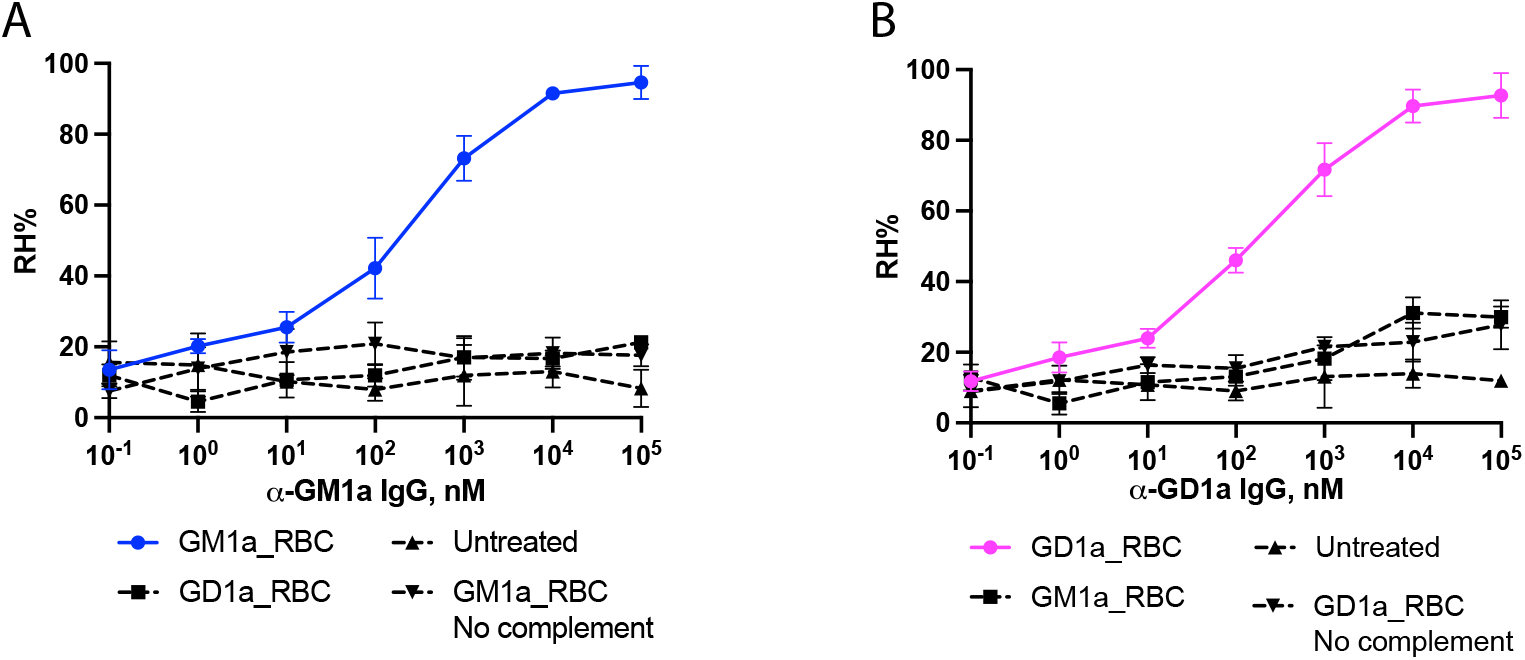
Complement-dependent hemolysis of GM1a and GD1a remodeled erythrocytes by anti-ganglioside antibodies. (A) GM1a remodeled erythrocytes undergo hemolysis in the presence of anti-GM1a IgG antibody and guinea pig serum complement (EC_50_ = 275 nM). (B) GD1a remodeled erythrocytes undergo hemolysis in the presence of anti-GD1a IgG antibody and guinea pig serum complement (EC_50_ = 201 nM).

## CONCLUSIONS

Despite advances in chemical and enzymatic synthesis of glycosphingolipids,^30,31^ the preparation of this class of compounds having complex glycan moieties has remained challenging. Here, we introduce a chemoenzymatic approach for the synthesis of immunologically important glycosphingolipids equipped with HNK-1 epitope and their unsulfated counterparts. Key to the approach was the enzymatic assembly of lacto-*neo*tetraose and lacto-*neo*hexaose as thioglycosides which were subjected to protecting group manipulations and chemical glycosylation to install a glucuronic acid moiety having a C-3 hydroxyl. The thioglycosides could be converted into *α*-anomeric fluorides by treatment with Barluenga’s reagent in the presence of HF-pyridine complex which was followed by sulfation of the C-3 hydroxyl to install a protected HNK-1 epitope. The strategy avoids incompatibilities of fluorination with the presence of sensitive functionalities such as a sulfate. After global deprotection, the fluorides were employed as substrate for enzymatic transfer to sphingosine catalyzed by a mutant endoglycoceramidase to give lyso-forms of glycosphingolipids (**16** and **17**). The latter compounds were further modified by stearic acid to yield the targeted glycosphingolipids (**1**-**2**). The compounds were employed to examine the mechanisms of cytotoxicity of an anti-HNK-1 antibody. Serum antibodies of anti-MAG neuropathy patients interact with HNK-1 epitopes that have been found on *N*-linked glycoproteins such as myelin associated glycoprotein (MAG) and myelin proteins (P0, P22) and NCAM and as part of glycosphingolipids. The latter compounds, which are expressed by peripheral nervous system, are thought to play a main role in the pathogenesis of anti-MAG neuropathy.^11,32^ The interaction of pathogenic autoantibodies with sulfoglucuronyl paraglobosides is expected to activate the complement system leading to the degradation of myelin sheath of nerve cells but there is no direct evidence for such a mechanism. Here, we took advantage of the amphiphilic properties of the synthetic glycosphingolipids to incorporate these compounds into the cell membrane of red blood cells. These cells do not express HNK-1 carrying glycoconjugates, and hence the incorporation of the synthetic glycolipids provides an opportunity to examine functional properties HNK-1 containing glycoconjugates. We found that exogenously administered sulfoglucuronyl paragloboside glycosphingolipid can easily be incorporated into the cell membrane of RBCs, which can be agglutinated by an anti-HNK-1 monoclonal antibody. In addition to hemagglutination, we also showed that the red blood cells undergo hemolysis in the presence of serum complement, which is dependent on the concentration of the IgM antibody and the amount of exogenously administered glycosphingolipid. Furthermore, depletion of Ca^2+^ ions abrogated the hemolysis, indicating that the lysis is mediated by the classical pathway of complement activation.

The erythrocyte remodeling approach could be extended to gangliosides and offers the prospect of a diagnostic tool for immune disorders involving auto- or cross-reacting antibodies such as MAG and GBS. It will not only provide binding but also functional properties such as complement mediated hemolysis. Detection of anti-antibodies is usually performed by ELISA^33^ or Western blotting.^32^ In the case of MAG, myelin associated glycoprotein is employed for antibody detection, which can lead to errors because it does not present HNK-1 as part of a glycosphingolipid. In the case of GBS, isolated gangliosides are employed to detect antibodies which is also challenging because ELISA requires coating, blocking and several washing steps which can lead to errors.^34^ A number of these problems can be circumvented by employing fully synthetic glycosphingolipids for cell surface remodeling of erythrocytes. It is worth noting that such erythrocytes have a rather longer shelf-life and a 25% suspension incubation with a known concentration of glycospingolipids can be stored in Alsever’s solution for up to 4 weeks without a notable degree of autohemolysis. Furthermore, the hemagglutination and hemolysis assays performed with stored RBCs were reproducible, indicating that very little or no shedding of glycolipids occur upon storage.

HNK-1 containing glycoconjugates have been implicated nervous system development, plasticity and dendritic spine morphogenesis.^35,36^ It acts as neural-recognition molecules employing laminin, P- and L-selectins and galectins as receptors for cell adhesion.^37,38^ Furthermore, the expression of HNK-1 is substantially reduced in brains of Alzheimer disease patients and may influence β-amyloid protein. Cell surface remodeling of cells with compounds such as **1** may provide opportunities to examine, at a molecular level, biological properties mediated by HNK-1.

It is to be expected that the employment chemoenzymatic strategy can provide entry into other classes of glycosphingolipids having sensitive glycan architectures and functional groups. For example, tumor-associated glycolipids such as VIM-2 and their sulfated and sialylated derivatives should be assessable by a similar chemoenzymatic strategy. The fucosylation pattern of such compounds can be controlled through the introduction of unnatural glycan modifications, whereas the use of chemical protecting group manipulations can accommodate the installation of functionalities such as sulfates.

## ASSOCIATED CONTENT

### Supporting Information

The Supporting Information is available free of charge at https://pubs.cs.org/doi/ Methods, analytical data, additional figures, and copies of NMR spectra (PDF).

## AUTHOR INFORMATION

**Corresponding Author**

**Geert-Jan Boons -** Chemical Biology and Drug Discovery, Utrecht Institute for Pharmaceutical Sciences, Utrecht University, Utrecht 3584 CG, The Netherlands; Complex Carbohydrate Research Center, University of Georgia, Athens, Georgia 30602, United States; Bijvoet Center for Biomolecular Research, Utrecht University, Utrecht 3584, The Netherlands; Chemistry Department, University of Georgia, Athens, Georgia 30602, United
States;

**Author**

**Mehman I. Bunyatov** - Chemical Biology and Drug Discovery, Utrecht Institute for Pharmaceutical Sciences, Utrecht University, Utrecht 3584 CG, The Netherlands; Complex Carbohydrate Research Center, University of Georgia, Athens, Georgia 30602, United States;

## ACKNOWLEDGMENTS

We thank Dr. G.P. Bosman (Utrecht University) for supplying glycosyltransferases and glycosynthase, and C.K. Page and J.D. Shepard (University of Georgia) for assisting with hemaglutination studies.

