## Supplemental Information for "Chemoenzymatic Synthesis of Glycosphingolipids having an HNK-1 Epitope for Erythrocyte Cell Surface Remodeling"

### Table of contents

1. Materials and methods
   Chemicals ………………………………………………………………….……..……….... S3
   Enzyme expression …………………………………………………….…......……......… S3
2. NMR nomenclature and properties of key compounds
   Glycan chain numbering (Figure S1) …………………………….………….........……... S4
   NMR characteristics of key intermediates (Figures S2 and S3) ……….………..............…... S5
3. Chemoenzymatic synthesis of oligosaccharide acceptors with various length
   General protocol for the installation of unit using UDP-GlcNHTFA and
      UDP-Gal in combination with HpB3GnT and LgtB, respectively …………….........… S6
4. Chemical modification of enzymatically assembled oligosaccharides
   General protocol for TFA removal ……………………………………….……….…..… S8
   General procedure for azido-transfer reaction ……………………….……...……...… S8
   General protocol for tin mediated allylation ………………………...…...…...…...... S9
   General protocol for benzylation ………………………………………......……..…..….. S9
   General procedure for conversion of azide into NHTCA ………………...……...…....……..... S9
   General procedure for allyl ether removal …………………………...……..……...………... S10
5. Synthesis of target glycans with diverse terminal epitopes
   General procedure for glycosylation of lactosyl acceptors with glucuronic acid donor …..…. S15
   General procedure for saponification and intralactone formation reactions ……..……........… S16
   General protocol for methanolysis of lactones ……………..…….…………...……....… S17
   General protocol for O-sulfation ……………………...…..……………….……...……... S17
   General procedure for anomeric a-fluoride installation ……...…...……………….……...... S17
   General procedure for global deprotection …………………....………………..……...……... S18
6. Synthesis of lyso- and full-length glycosphingolipids
   General procedure for the synthesis of *lyso*-glycosphingolipids via enzymatic *en block*
      transfer of glycosyl fluorides …...…........….........…......…......…........….................. S26
   General protocol for acylation of glycosyl sphingosines ………….…………….………..... S26
7. Synthesis of glycosphingolipids having hexasaccharide backbones
   Figure S4 ……………………………………………….………..…………….……..... S32
   Figure S5 ……….……………………………………….………….……………….……..... S33
8. Glycoremodeling and hemagglutination of red-blood cells and cell-based assays
   General protocol for glycolipid remodeling of red blood cell ……………………………....... S50
   General protocol for antibody-agglutination of SGPG (1a) and GPG (2a)
      modified erythrocytes ………...………...……..……….…...…….……….…………… S51
   Figure S6 ……………………………………………………….…………...……….......... S52
   Protocol for antibody-agglutination of GM1a and GD1a remodeled erythrocytes …...…........ S52
   Figure S7 ………………………………………….……………………...……...…... S53
   Protocol for antibody-mediated complement dependent cytotoxicity …. ….……….………... S53
9. References ……………………………………………………………………….......... S54
10. Copies of NMR spectra …….………………….……………………………..…........ S54

### 1. Materials and methods

#### Chemicals

Unless otherwise specified, all chemicals were sourced from Sigma-Aldrich. Monosaccharide for building block synthesis were purchased from Carbosynth Limited (UK). UDP-Galactose was purchased from Roche (material number: 07703562103), UDP-GlcNHTFA[1] was prepared according to the published protocols. Acetonitrile, dichloromethane, toluene, tetrahydrofuran and N,N-dimethylformamide used for synthesis were anhydrous grade and were supplied by an MB SPS 5 solvent purification system. Other organic solvents used for reactions were obtained from Biosolve Chemie. Technical grade organic solvents for work-up procedures were supplied from VWR Chemicals. Deuterated solvents for NMR experiments were purchased from Cambridge Isotope Laboratories.

$^{1}$H and $^{13}$C NMR spectra were recorded on Bruker 600 UltraShield. Chemical shifts are reported in parts per million (ppm) relative to $CDCl_3$ as the internal standard. NMR data are presented as follows: Chemical shift, multiplicity (s = singlet, d = doublet, t = triplet, dd = doublet of doublets, m = multiplet, b = broad); coupling constants are reported in Hertz (Hz). All NMR signals were assigned on the basis of $^{1}$H NMR, COSY, HSQC, HMBC, TOCSY and NOESY experiments. Mass spectra were recorded on either an Applied Biosystems SCIEX MALDI-TOF/TOF 5800 mass spectrometer or a Shimadzu Biotech Axima-CFR MALDI-TOF. Column chromatography was performed on silica gel G60 (Silicycle, 60-200 µm, 60 Å). TLC analysis was performed using precoated silica gel 60 F-254 plates (Merck) with detection by UV light (254 nm) where applicable, and by charring with 10% sulfuric acid in ethanol or a *p*-anisaldehyde staining solutionn ethanol (1% v/v). C8 and C18 cartridges were purchased from Thermo Fisher (Hypersep™ Catalog Number 60108) for reverse phase chromatography. Unless specified otherwise, all moisture sensitive reactions were carried out under argon atmosphere and in the presence of activated molecular sieves. Unless otherwise stated, all reactions were carried out at room temperature (RT) in glassware with magnetic stirring. Solutions in organic solvents were dried with $Na_2SO_4$ and concentrated at 40 °C/2 kPa. Molecular sieves were flame-dried in vacuo immediately prior to use.

#### Enzyme expression

Glycosyltransferases HpB3GnT and NmLgtB were expressed in *E. coli* BL21 cell culture according to published protocols.[2] Glycosynthase mutant endoglycoceramidase (EGC II D351S/E314Y) was prepared according to published protocol.[3] All enzymes were expressed as soluble proteins containing a hexahistidine tag and purified by $Ni^{2+}$-NTA affinity column chromatography.

### 2. NMR nomenclature and analysis of target glycans

#### Glycan chain numbering

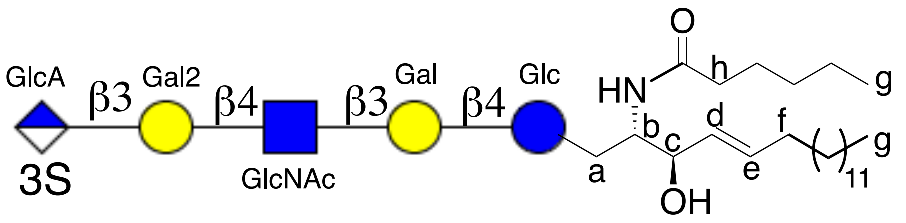


**Figure S1.** General approach to the labeling of monosaccharides of target glycolipids and ceramide moiety in protons for NMR assignment.

The numbering of the glycan chain followed a direction from the reducing end to the non-reducing terminus. Chemical shift assignments for each proton of the protected glycans were performed starting from the most downfield proton and continuing sequentially to the most upfield proton. For deprotected glycans, the assignments were performed separately for carbohydrate and non-carbohydrate protons. Proton assignments within each monosaccharide were performed individually, counting from H-1 to H-6 for all residues, except for GlcA, which followed the sequence H-1 to H-5. Aglycone moieties were labeled depending on the nature of residue. For thiophenyl group only the chemical shift for aromatic protons were estimated. Sphingosine and ceramide are labeled according to the provided figure: **a** – C1 protons of sphingosine, **b** – C2-NH protons, **c** – C3 protons, **d** – HC= alkenyl proton of sphingosine, **e** – C=CH- alkenyl proton of sphingosine, **f** – C=C-$CH_2$- allylic protons of sphingosine, **h** – α-$CH_2$ of stearic acid, and **g** – terminal $CH_3$ of sphingosine and fatty acid, protons with the most upfield chemical shifts. Chemical shifts of aliphatic -$CH_2$- of ceramide were not individually assigned.

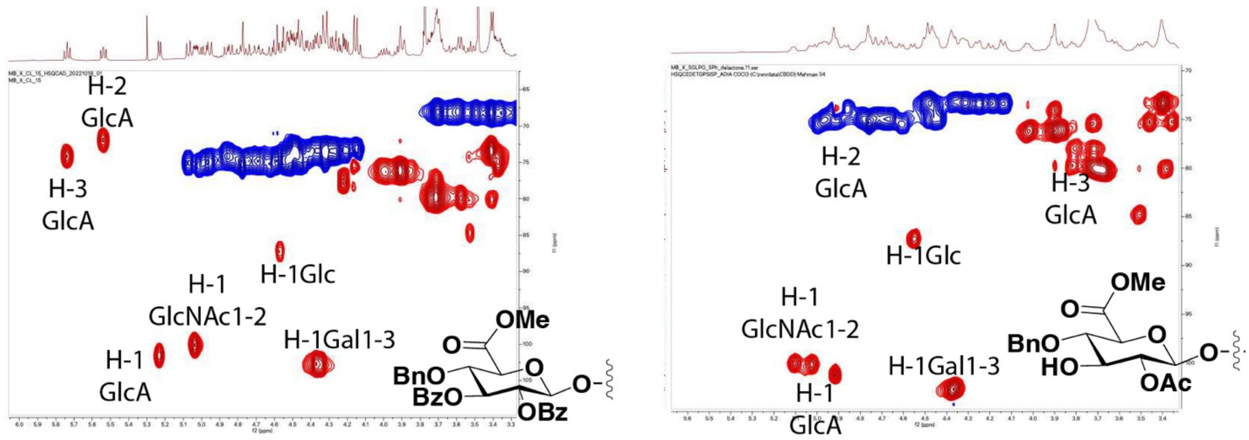


**Figure S2**. Chemical shift change of H-3 proton of glucuronic acid upon deprotection.

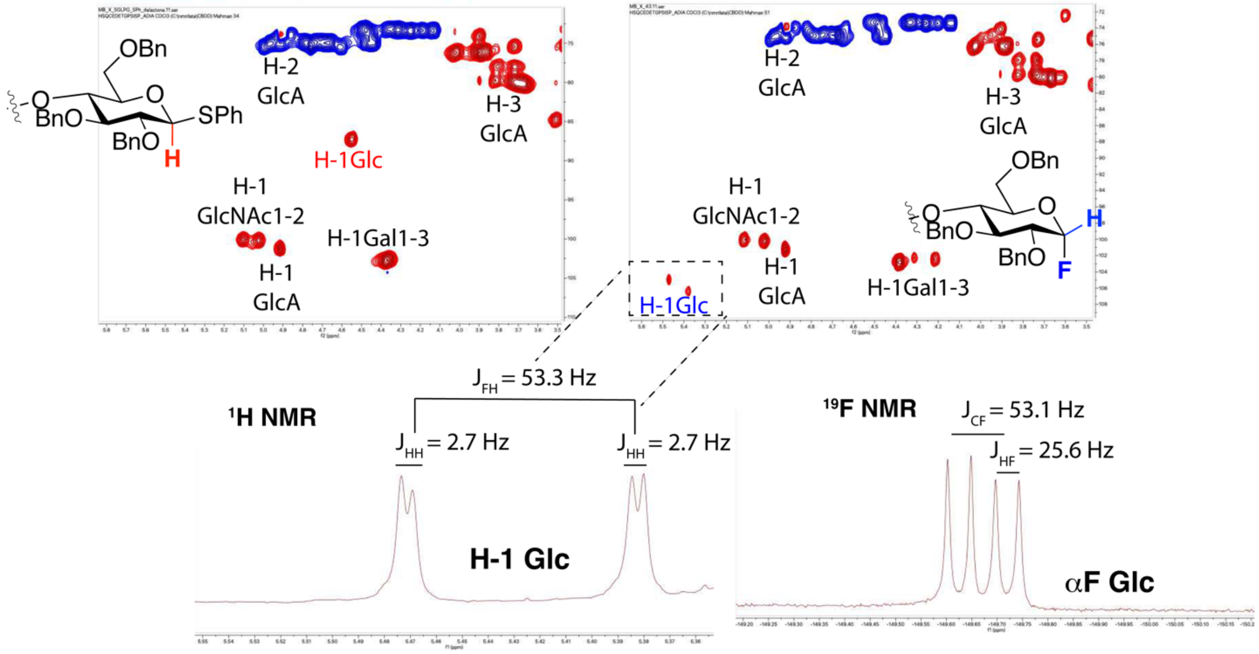


**Figure S3**. NMR characteristics of thioglycoside and a-glycosyl fluoride. Anomeric fluoride installation is characterized by the disappearance of diagnostic chemical shift of H-$1_{Glc}$ of thioglycoside and formation of a new signal as doublet of doublets with a characteristic chemical shift and J-coupling constants corresponding to the formation of anomeric a-fluorides. $^{19}$F NMR spectroscopy of the product matches with the reported chemical shifts and spin-spin splitting coupling constants of anomeric α-fluorides.

### 3. Chemoenzymatic synthesis of oligosaccharide acceptors with various length

**General protocol for the installation of unit using UDP-GlcNHTFA and UDP-Gal in combination with HpB3GnT and LgtB, respectively**

UDP-GlcNHTFA (1.5 equiv) and acceptor **(3, 7)** were dissolved in Tris buffer (pH = 7.8, 100 mM) containing 10 mM $MgCl_2$ and 1 mM DTT to give a final concentration of 10 mM of acceptor. To this solution, Hpβ3GlcNAcT (1%, w/w) and CIAP (10 mU) were added, and the reaction mixture was incubated at 30 ℃ overnight in an incubator with shaking. MALDI-TOF MS was used to monitor the progress of the reaction and in case of incomplete conversion an extra portion of UDP-GlcNHTFA and Hpβ3GlcNAcT were added. Upon completion of the enzymatic transformation, the reaction mixture was centrifuged, and supernatant was decanted using a 1 mL syringe and passed through Sep-Pak C18-reverse phase cartridge by first washing with 5% aqueous acetonitrile to remove inorganic salts and nucleotide bases, followed by elution with 40% aqueous acetonitrile to obtain the desired intermediate compounds. Fractions containing product were pooled and freeze-dried. The resulting residue was redissolved in water to which Tris buffer (100 mM, pH = 7.5), UDP-Gal (1.5 equiv), $MgCl_2$ (10 mM), CIAP (0.1 U) and NmLgtB enzyme were added. The reaction mixture was incubated at 25 °C. After several hours, MALDI-TOF MS analysis indicated the completion of galactose transfer. After centrifugation, the mixture was passed through a Sep-Pak C18 cartridge and washed with low concentration of acetonitrile and eluted with 30% ACN. Fractions containing product were combined and lyophilized. The crude product was applied to a Biogel-P-2 size exclusion column and eluted with 10 mM aqueous $NH_4HCO_3$. Product containing fractions were analyzed by MALDI-TOF MS, concentrated, lyophilized and used in the subsequent steps. Titled compounds were analyzed by NMR and MS.

#### ***β*-D-galactopyranosyl-(1→4)-1-phenylthio-*β*-D-glucopyranoside (3)**

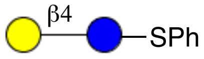


Compound **3** was synthesized in a multi-gram scale as described previously.[4] MALDI-TOFMS *m/z* $C_{32}H_{45}NO_{13}$ $(M+Na)^+$ calculated 457.45, found 457.25.

$^1$H NMR (600 MHz, $D_2O$) δ: (non carbohydrate): 7.48 – 7.44 (m, 2H, SPh), 7.22 – 7.14 (m, 3H,

SPh).

$^{13}$C NMR (151 MHz, $D_2O$): $\delta$ (non carbohydrate): 131.8, 128.5, 127.0 (C Ar).

| **3** | H1 | H2 | H3 | H4 | H5 | H6 |
|---|---|---|---|---|---|---|
| Glc | 4.51<br>d, *J* = 9.9 Hz | 3.18 | 3.47 | 3.49 | 3.35 | 3.80<br>dd, $J_1$ = 2.8 Hz, $J_2$ = 12.8 Hz<br>3.79<br>dd, $J_1$ = 4.0 Hz, $J_2$ =13.0 Hz |
| Gal | 4.26<br>d, *J* = 7.8 Hz | 3.45 | 3.48 | 3.71,<br>d, *J* = 3.3 Hz | 3.38,<br>d, J = 3.44 Hz | 3.60<br>3.66 |

| **3** | C1 | C2 | C3 | C4 | C5 | C6 |
|---|---|---|---|---|---|---|
| Glc | 87.9 | 72.2 | 78.8 | 75.8 | 79.4 | 60.6 |
| Gal | 103.9 | 76.6 | 75.5 | 68.7 | 73.5 | 61.0 |

**$\beta$-D-galactopyranosyl)-(1→4)-(2-trifluoroacetamido-2-deoxy-$\beta$-D-glucopyranosyl)-(1→3)-($\beta$-D-galactopyranosyl)-(1→4)-1-phenylthio-$\beta$-D-glucopyranoside (7)**

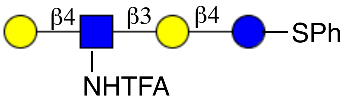


Compound **7** (1.8 g, 2.05 mmol) was obtained from compound **3** (1.2 g, 2.6 mmol) using the general protocol for LacNHTFA installation using UDP-GlcNHTFA (1.7 g, 2.5 mmol) and UDP-Gal (1.5 g, 2.5 mmol). MALDI-TOF MS *m/z* $C_{32}H_{46}F_3NO_{20}S$ $(M+Na)^+$ 876.22 found 876.43.

$^1$H NMR (600 MHz, $D_2O$) δ: (non-carbohydrate): 7.49 – 7.43 (m, 2H, SPh), 7.21 – 7.14 (m, 3H, SPh).

$^{13}$C NMR (151 MHz, $D_2O$): $\delta$ (non-carbohydrate): 132.3, 128.1, 127.0 (C Ar).

| **7** | H1 | H2 | H3 | H4 | H5 | H6 |
|---|---|---|---|---|---|---|
| Glc | 4.52 | 3.21 | 3.51 | 3.54 | 3.45 | 3.86 – 3.80 |
| Gal | 4.35<br>d, *J* = 7.3 Hz | 3.47 | 3.50 | 4.11<br>d, *J* = 2.8 Hz | 3.27 | 3.70 – 3.66 |
| GlcNHTFA | 4.74 | 3.81 | 3.54 | 3.66 | 3.60 | 3.80 |
| Gal(2) | 4.36 | 3.44 | 3.46 | 3.94 | na | 3.70 – 3.66 |

| 7 | C1 | C2 | C3 | C4 | C5 | C6 |
|---|---|---|---|---|---|---|
| Glc | 87.6 | 72.8 | 74.3 | 78.8 | 75.1 | 58.9 |
| Gal | 103.1 | 69.6 | 80.1 | 68.3 | 72.4 | 61.8 |
| GlcNAc | 101.6 | 52.1 | 74.6 | 77.8 | 75.2 | 60.1 |
| Gal(2) | 103.4 | 75.1 | 72.2 | 68.5 | 72.3 | 61.8 |

### 4. Chemical modification of enzymatically assembled tetra- and hexasaccharide

This process involves a variety of chemical transformations to modify tetra- and hexasaccharide, including deacylation, azido-transfer, allylation, benzylation, and further modifications.

**General protocol for TFA removal.** Compounds **7** and **S3** were dissolved in water and the pH of the resulting solutions were adjusted to 10 by dropwise addition of 1 M NaOH solution. The reaction mixtures were incubated at 37 ℃ and progress of reactions were monitored by MALDI-TOF MS. Upon completion of the deacylation, the pH of the reaction was brought to neutral by dropwise addition of 1 M HCl solution and then the samples were freeze-dried. Crude products were applied to a Biogel-P-2 size exclusion column and eluted with 10 mM aqueous $NH_4HCO_3$. Carbohydrate containing fractions were identified by MALDI-TOF MS and fractions containing products were pooled and lyophilized.

**General procedure for azido-transfer reaction.** To a solution of compound **8** and **S4** in water, $K_2CO_3$ (10 equiv), imidazole-1-sulfonyl azide hydrogen chloride (10 equiv) (prepared following the protocol,[5] and cat. amount of $CuSO_4{\cdot}5H_2O$ were added and the reaction mixture was incubated at 37 ℃ for 2 h. The conversion was monitored by MALDI-TOF MS and in case starting material was remaining an additional portion of diazo-transfer reagent was added followed by adjustment of pH to 8. The reaction mixture was further incubated until full conversion was observed by mass spectrometry and TLC analyses. Insoluble solid particles were removed by centrifugation and the supernatant was lyophilized. The resulting material was dissolved in water and applied onto benchtop C18 beads, which were washed with 10% acetonitrile in water followed by elution with 50% aqueous acetonitrile. Fractions containing products were collected and freeze-dried using $H_2O$:$^t$BuOH (3:1, v/v) as solvent mixture.

**General protocol for tin-mediated allylation.** Compounds **9** and **S5** and $Bu_2SnO$ (1.5 equiv) were suspended in anhydrous methanol and heated under reflux at 80 °C for 2 h, after which the suspension turned clear. The reaction mixture was stirred for a further 20 min under refluxing conditions after which, it was allowed cool to RT. Methanol was evaporated under reduced pressure, and the resulting residue was coevaporated from toluene to remove the residual methanol and then dried *in-vacuo* for 2 h. The residue was taken up in anhydrous DMF (50 mL) and allyl bromide (3 equiv) and TBAI (0.4 equiv) were added, and the reaction mixture was stirred at RT for 18 h. The progress of the reaction was monitored by TLC. In case of slow conversion 1 more equivalent of allyl bromide and 0.2 equivalent of TBAI were added. Upon completion, the reaction mixture was concentrated under reduced pressure, the residue dissolved up in a minimum amount of DCM:MeOH (1/1, v/v), dry loaded on a silica gel column, and chromatographed using a gradient of EtOAc:MeOH (100% → 60%). Fractions containing product were collected, concentrated under reduced pressure and the product characterized by MS and NMR.

**General protocol for benzylation.** Intermediates were dissolved in DMF and the resulting solution cooled in an ice-bath. NaH (1.5 equiv per each hydroxyl) and BnBr (2 equiv per each hydroxyl) were added and the reaction mixture was stirred for 30 min at 0 °C and then 2 h at RT. The progress of the reactions was monitored by TLC and when completed, the reaction was quenched by the addition of ice-cold methanol. The solvents were evaporated under reduced pressure and the residue was dissolved in DCM, subsequently washed with 1M aqueous HCl, sat. $NaHCO_3$ solution and brine. The organic layer was separated, dried ($Na_2SO_4$) and concentrated under reduced pressure. The residue was applied to a column silica gel and eluted with a gradient of petroleum ether: EtOAc = 20:1 → 6:1. Fractions containing product were concentrated under reduced pressure and used in the next step without further characterization.

**General procedure for conversion of azide into NHTCA.** Intermediate compounds obtained after benzylation of **21b-c** were dissolved in 90% aq. THF and to this solution 1M solution of $PMe_3$ in THF was added untill a final 5-fold excess of azido glycan (caution: no base was used for the reduction to avoid the *N*-benzylation with traces of remaining benzyl bromide from the previous reaction). The reaction mixtures stirred for 2 h at RT, after which TLC indicated full consumption of the starting material. The solvents were evaporated under reduced pressure and the residue was coevaporated twice from toluene to remove trace amounts of moisture and trimethyl phosphine. The residue was further dried in high *vacuo* for 2 h. The solid residue was dissolved in DCM, triethylamine (2 equiv to starting material) was added followed by the addition of trichloroacetyl chloride (1.5 equiv per each $NH_2$). TLC monitoring indicated the formation of trichloroacetamide product within 15 min. The reaction mixture was further stirred for 30 min, quenched with sat. solution of aqueous $NaHCO_3$, and washed with 1M HCl, brine. Organic phases were collected, dried ($Na_2SO_4$) and concentrated *in vacuo*. The residue was applied to a silica gel column chromatography and eluted with Tol:EtOAc=9:1, v:v.

**General procedure for allyl ether removal.** To solutions of compounds **10** and **S6** in dry THF a solution of catalytic amount of $[Ir(cod)(PPh_2Me)_2]PF_6$ catalyst (0.2 equiv)* in THF was added and the solution stirred overnight at RT. After which, it was concentrated under reduced pressure and obtained crude residue dissolved in acetone:$H_2O$ (9:1, v/v) mixture. To the obtained solution $HgCl_2$ (2 equiv.) and HgO (0.4 equiv.) were added, the reaction mixture stirred at RT for 1 h. The progress of deallylation was monitored by TLC and when no more starting material was observed, the reactions were diluted with DCM and washed with 10% aqueous solution of KI** and brine. Organic layers were separated, dried over anhydrous $Na_2SO_4$ and concentrated by rotary evaporation under reduced pressure. The remaining crude product was applied to silicagel column chromatography and eluted with Tol:EtOAc=5:1, v:v eluent to afford the required compound. Fractions containing product were collected, concentrated and characterized the by MALDI-TOF MS and NMR spectroscopy.

*$[Ir(cod)(PPh_2Me)_2]PF_6$ catalyst needs to be activated by bubbling $H_2$ through the solution of catalyst in THF for 2 h.

** KI solution was pretreated with 2% $Na_2S_2O_4$ solution to remove the trace amounts of $I_2$, which can act as a promoter for the thioglycoside activation.

**β-D-galactopyranosyl)-(1→4)-(2-amino-2-deoxy-β-D-glucopyranosyl)-(1→3)-(β-D-galactopyranosyl)-(1→4)-1-phenylthio-β-D-glucopyranoside (8)**

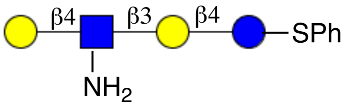


Compound **7** (1.0 g, 1.11 mmol) was subjected to deprotection according to general procedure to obtain 820 mg of compound **8** (96%). MALDI-TOF MS *m/z* $C_{30}H_{47}NO_{19}S$ $(M+Na)^+$ calculated 780.74, found 781.02.

$^1H$ NMR (600 MHz, $D_2O$) δ: (non-carbohydrate): 7.50 – 7.14 (m, 5H, SPh).
$^{13}C$ NMR (151 MHz, $D_2O$): *δ* (non-carbohydrate): 127.0, 129.6, 131.3 (C Ar).

Carbohydrate region:

| **8** | H1 | H2 | H3 | H4 | H5 | H6 |
|---|---|---|---|---|---|---|
| Glc | 4.71 | 3.28<br>t, J = 8.9 Hz | 3.57 | 3.54 | na | 3.70 – 3.61 |
| Gal | 4.38<br>d, *J* = 7.8 Hz | 3.59 | 3.77 | 3.80,<br>d, *J* = 2.8 Hz | na | 3.70 – 3.61 |
| GlcNAc | 4.87<br>d, *J* = 8.4 Hz | 3.02<br>d, J = 9.6 Hz | 3.72 | 3.57 | na | 3.70 – 3.61 |
| Gal(2) | 4.34<br>d, *J* = 7.9 Hz | 3.47 | 3.56 | 4.07<br>d, *J* = 2.2 Hz | na | 3.70 – 3.61 |

| **8** | C1 | C2 | C3 | C4 | C5 | C6 |
|---|---|---|---|---|---|---|
| Glc | 87.2 | 71.3 | 75.0 | 74.4 | na | 59.0 – 61.0 |
| Gal | 102.2 | 70.0 | 82.5 | 68.2 | na | 59.0 – 61.0 |
| GlcNAc | 100.5 | 55.8 | 70.8 | 72.7 | na | 59.0 – 61.0 |
| Gal(2) | 103.1 | 70.7 | 78.7 | 68.5 | na | 59.0 – 61.0 |

**β-D-galactopyranosyl-(1→4)-(2-azido-2-deoxy-β-D-glucopyranosyl)-(1→3)-(β-D-galactopyranosyl)-(1→4)-1-phenylthio-β-D-glucopyranoside (9)**

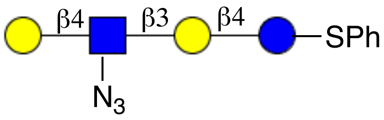


Compound **8** (800 mg, 1.02 mmol) was converted into **9** (720 mg) in 87% yield according to the general protocol for azido transfer. MALDI-TOF MS *m/z* $C_{30}H_{45}N_3O_{19}S$ $(M+Na)^+$

calculated 806.74, found 806.61.

$^{1}H$ NMR (600 MHz, $D_2O$) δ: (non-carbohydrate): 7.51 – 7.11 (m, 5H, SPh).
$^{13}C$ NMR (151 MHz, $D_2O$): $\delta$ (non-carbohydrate): 126.0, 129.1, 131.2 (C Ar).

| **9** | H1 | H2 | H3 | H4 | H5 | H6 |
|---|---|---|---|---|---|---|
| Glc | 4.69 | 3.26 | 3.51 | 3.60 | 3.24 | 3.81<br>3.72 |
| Gal | 4.40<br>d, $J$ = 7.8 Hz | 3.37 | 3.56 | 4.10<br>d, $J$ = 2.6 Hz | na | 3.79 – 3.56 |
| GlcNAc | 4.72 | 3.26 | 3.70 | 3.65 | 3.30 | 3.88 |
| Gal(2) | 4.36<br>d, $J$ = 8.2 Hz | 3.40 | 3.54 | 3.92<br>d, $J$ = 2.3 Hz | na | 3.79– 3.55 |

| **9** | C1 | C2 | C3 | C4 | C5 | C6 |
|---|---|---|---|---|---|---|
| Glc | 87.5 | 72.5 | 74.0 | 75.4 | 70.4 | 59.7 |
| Gal | 102.1 | 71.6 | 82.6 | 67.4 | na | 61.8 |
| GlcNAc | 103.0 | 57.6 | 75.7 | 75.8 | 72.0 | 59.6 |
| Gal(2) | 101.9 | 72.1 | 80.6 | 67.8 | na | 61.8 |

**3-O-allyl-*β*-D-galactopyranosyl)-(1→4)-(2-azido-2-deoxy-*β*-D-glucopyranosyl)-(1→3)-(*β*-D-galactopyranosyl)-(1→4)-1-phenylthio-*β*-D-glucopyranoside (S1)**

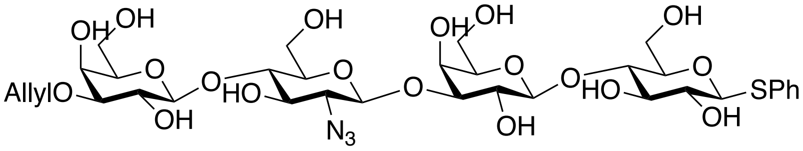


Compound **S1** (520 mg, 74%) was synthesized according to the general allylation procedure starting from **9** (660 mg, 0.82 mmol), $Bu_2SnO$ (310 mg, 1.25 mmol), allyl bromide (280 $\mu$l, 3.28 mmol) and TBAI (121 mg, 0.328 mmol). MALDI-TOF MS *m/z* $C_{33}H_{49}N_3O_{19}S$ $(M+2Na-N_2)^+$ calculated 846.81, found 847.34.

$^{1}H$ NMR (600 MHz, $D_2O$) δ 7.47–7.08 (m, 5H, Ar-H), 5.86 (m, 1H, $CH_2$=*C<u>H</u>*-$CH_2$-), 5.20 (d, J = 17.1 Hz, 1H, C*<u>H</u>*H=CH-$CH_2$-), 5.20 (d, J = 11.0 Hz, 1H, CH*<u>H</u>*=CH-$CH_2$-), 4.62 (H-1, GlcNAc), 4.54 (d, J = 8.8 Hz, 1H, H-1 Glc), 4.34 – 4.28 (m, 3H, H-1 Gal-2, H-1 Gal-1,), 4.20 – 3.99 (m, 4H, $CH_2$=CH-C*<u>H</u>*$_2$, H-4 Gal1-2), 3.88 – 3.84 (m, 1H, H-6a Glc), 3.76 (dd, $J_1$ = 8.5 Hz, $J_2$ = 4.4 Hz, 1H, H-6b Glc), 3.72 – 3.56 (m, 6H, H-6 GlcNAc, H-6 Gal1-2, H-2 Gal1, H-3 Gal1), 3.51 – 3.34 (m,

5H, H-2 Gal2, H-3 Gal2, H-3 GlcNAc, H-4 GlcNAc, H-3 Glc), 3.28 (t, *J* = 8.7 Hz, 1H, H-2 GlcNAc) 3.18 (m, 1H, H-5 Glc).

$^{13}C$ NMR (151 MHz, $D_2O$): *δ* 57.3 (C6 GlcNAc), 60.1 (C6 Gal1-2, C6 Glc), 66.0 (C4 Gal1-2), 67.1 (C2 GlcNAc), 70.1 ($CH_2$=CH-$\underline{C}H_2$-), 73.4 (C2 Glc), 76.1 (C4 Glc), 76.2 (C4 GlcNAc), 78.4 (C3 Gal1), 79.5 (C3 Gal2), 79.8 (C2 Glc), 80.2 (C3 Glc), 82.2 (C3 GlcNAc), 87.9 (C1 Glc), 101.4 (C1 GlcNAc), 102.5 (C1 Gal1), 102.7 (C1 Gal-2), 120.1 ($\underline{CH_2}$=CH-$CH_2$-), 126.2 – 127.8 (C Ar), 135.4 ($CH_2$=$\underline{CH}$-$CH_2$-).

**3-O-allyl-2,4-6-tri-O-benzyl-β-D-galactopyranosyl)-(1→4)-(2-azido-3,6-di-O-benzyl-2-deoxy-β-D-glucopyranosyl)-(1→3)-2,4-6-tri-O-benzyl-β-D-galactopyranosyl)-(1→4)-2,3,6-tri-O-benzyl-1-phenylthio-β-D-glucopyranoside (10)**

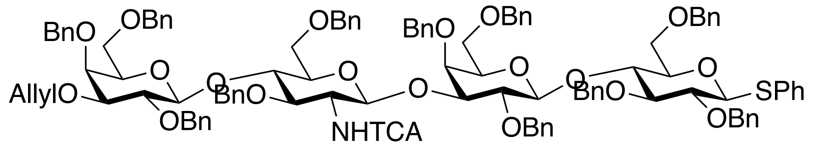


Compound **S1** (500 mg, 0.59 mmol) was benzylated using the general procedure in the presence of NaH (520 mg, 13.0 mmol) and benzyl bromide (1.5 mL), and then subjected to Staudinger azide reduction using a $PMe_3$ (3.6 mL) solution. The obtained amine intermediate was protected as a trichloroacetamide using 130 μL trichloroacetyl chloride following the general procedure for amine protection to furnish **10** (620 mg, 53% over 3 steps). MALDI-TOF MS *m/z* $C_{112}H_{116}Cl_3NO_{20}S$ $(M+Na)^+$ calculated 1957.55, found 1958.23.

$^1H$ NMR (600 MHz, Chloroform-d) δ 7.52–7.03 (m, 45H, Ar-H), 6.61 (m, 1H, -N*H*TCA), 5.93 (m, 1H, $CH_2$=$\underline{CH}$-$CH_2$-), 5.33 (dd, $J_1$ = 1.8 Hz, $J_2$ = 17.2 Hz, 1H, CH$\underline{H}$=CH-$CH_2$-), 5.18 (dd, $J_1$ = 1.7 Hz, $J_2$ = 10.4 Hz, 1H, C$\underline{H}$H=CH-$CH_2$-), 5.08 (d, *J* = 7.3 Hz, 1H, H-1 GlcNAc), 5.01 (d, *J* = 10.9 Hz, 1H, -OC$\underline{H_2}$Ph), 4.99 (d, *J* = 11.8 Hz, 1H, -OC$\underline{H_2}$Ph), 4.94 (d, *J* = 9.9 Hz, 1H, -OC$\underline{H_2}$Ph), 4.83 – 4.73 (m, 4H, -OC$\underline{H}$HPh, -OC$\underline{H_2}$Ph), 4.68 (d, *J* = 12.7 Hz, 1H, -OC$\underline{H_2}$Ph), 4.62 (d, J = 10.9 Hz, 1H, -OC$\underline{H_2}$Ph), 4.56 (d, J = 9.9 Hz, 1H, H-1 Glc), 4.53 – 4.46 (m, 4H, -OC$\underline{H_2}$Ph), 4.42 (d, *J* = 7.3 Hz, 1H, H-1 Gal2), 4.38 – 4.35 (m, 2H, -OC$\underline{H_2}$Ph, H-1 Gal1), 4.32 (d, J = 11.6 Hz, 1H, -OC$\underline{H}$HPh), 4.28 (d, J = 11.9 Hz, 1H, -OC$\underline{H}$HPh), 4.20 (d, J = 11.9 Hz, 1H, -OC$\underline{H}$HPh), 4.19 (d, J

= 12.2 Hz, 1H, -OC*H*HPh), 4.18 – 4.15 (m, 2H, $CH_2$=CH-C*$H_2$*-), 4.04 (m, 1H, H-4 GlcNAc), 3.93 (d, *J* = 2.3 Hz, 1H, H-4 Gal2), 3.92 (t, $J_1$ = 9.8 Hz, 1H, H-4 Glc), 3.86 (d, *J* = 3.3 Hz, 1H, H-4 Gal1), 3.83 (dd, $J_1$ = 4.4 Hz, $J_2$ = 7.7 Hz, 1H, H-6a Glc), 3.79 – 3.74 (m, 3H, H-2 GlcNAc, H-3 Gal1, H-3 GlcNAc), 3.74 – 3.68 (m, 5H, H-3 Gal2, H-2 Gal2, H-2 Gal1, H-6a GlcNAc, H-6b Glc), 3.67 (d, *J* = 10.1 Hz, 1H, H-6b Glc), 3.63 (dd, $J_1$ = 1.8 Hz, $J_2$ = 10.4 Hz, 1H, H-6b GlcNAc), 3.55 – 3.47 (m, 3H, H-3 Glc, H-6a Gal1, H-5 GlcNAc), 3.44 – 3.40 (m, 2H, H-6a Gal2, H-2 Glc), 3.39 – 3.33 (m, 4H, H-6b Gal1, H-5 Gal1, H-5 Gal2), 3.26 (d, $J_1$ = 3.5 Hz, $J_2$ = 6.9 Hz, 1H, H-6b Gal2), 3.20 (d, $J_1$ = 3.9 Hz, $J_2$ = 9.5 Hz, 1H, H-5 Glc).

$^{13}$C NMR (151 MHz, $CDCl_3$): $\delta$ 57.8 (C2 GlcNAc), 68.1 (C6 Gal2), 68.2 (C6 Gal1), 68.3 (C6 Glc, C6 GlcNAc), 71.6 (-*C*$H_2$ allyl), 73.1 (-O*C*$H_2$Ph), 73.2 (C5 Gal2, C5 Gal1), 73.6 (C4 Gal2), 73.4 (-O*C*$H_2$Ph), 73.5 (-O*C*$H_2$Ph), 73.9 (-O*C*$H_2$Ph), 74.6 (-O*C*$H_2$Ph), 74.9 (-O*C*$H_2$Ph), 75.0 (-O*C*$H_2$Ph), 75.5 (-O*C*$H_2$Ph), 75.6 (-O*C*$H_2$Ph), 76.6 (C5 GlcNAc), 76.3 (C4 Gal1), 76.6 (C4 GlcNAc), 78.2 (C3 Gal1), 79.3 (C5 Glc), 79.8 (C2 Glc), 79.9 (C3 GlcNAc), 80.0 (C3 Gal1), 80.1 (C3 Gal2), 82.2 (C75.4 (C5 GlcNAc), 76.3 (C4 Gal1), 76.3 (C4 Glc), 76.6 (C4 GlcNAc), 78.0 (C3 GlcNAc), 79.83 (C3 Gal), 79.9 (C2 Gal), 80.3 (C2 Gal), 81.7 (C3 Gal2), 82.2 (C2 Glc), 87.3 (C1 Glc), 100.3 (C1 GlcNAc), 102.6 (C1 Gal), 103.0 (C1 Gal2), 116.5 (*C*$H_2$=CH-$CH_2$-), 127.2, 127.4, 127.5, 127.5, 127.6, 127.7, 127.7, 127.8, 127.8, 127.9, 128.0, 128.1, 128.2, 128.2, 128.3, 128.9, 132.0 (C Ar), 134.6 ($CH_2$=*C*H-$CH_2$-).

**2,4-6-tri-O-benzyl-β-D-galactopyranosyl)-(1→4)-(2-azido-3,6-di-O-benzyl-2-deoxy-β-D-glucopyranosyl)-(1→3)- 2,4-6-tri-O-benzyl-β-D-galactopyranosyl)-(1→4)-2,3,6-tri-O-benzyl-1-phenylthio-β-D-glucopyranoside (11)**

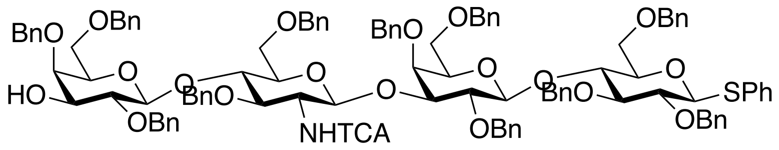


360 mg of compound **11** was obtained from the reaction of 600 mg (0.306 mmol) of **10** with [Ir(cod)$(PPh_2Me)_2$]$PF_6$ catalyst (50 mg, 0.062 mmol) and $HgCl_2$ (165 mg, 0.61 mmol) and HgO (27.0 mg, 0.122 mmol) according to general procedure for allyl removal.

MALDI-TOF MS *m/z* $C_{123}H_{131}Cl_3N_2O_{23}$ $(M+Na)^+$ calculated 1917.48, found 1917.72.

$^1$H NMR (600 MHz, Chloroform-d) δ 7.51–7.04 (m, 45H, Ar-H), 6.60 (d, J = 7.5 Hz, 1H, -N*H*TCA), 5.08 (d, *J* = 7.3 Hz, 1H, H-1 GlcNAc), 5.07 (d, *J* = 11.1 Hz, 1H, -OC$\underline{H_2}$Ph), 5.05 (d, *J* = 7.8 Hz, 1H, H-1 GlcNAc), 5.01 (d, *J* = 12.8 Hz, 1H, -OC$\underline{H_2}$Ph), 4.96 (d, *J* = 9.5 Hz, 1H, -OC$\underline{H_2}$Ph), 4.86 – 4.68 (m, 5H, -OC$\underline{H}$HPh, -OC$\underline{H_2}$Ph), 4.64 (d, J = 11.2 Hz, 1H, -OC$\underline{H_2}$Ph), 4.54 (d, J = 8.3 Hz, 1H, H-1 Glc), 4.52 – 4.45 (m, 4H, -OC$\underline{H_2}$Ph), 4.43 (d, *J* = 7.8 Hz, 1H, H-1 Gal2), 4.38 – 4.32 (m, 3H, -OC$\underline{H_2}$Ph, H-1 Gal1), 4.30 (d, J = 12.3 Hz, 1H, -OC$\underline{H}$HPh), 4.21 (m, 1H, -OC$\underline{H}$HPh), 4.17 (d, J = 12.0 Hz, 1H, -OC$\underline{H}$HPh), 4.16 – 4.14 (m, 2H, $CH_2$=CH-C$\underline{H_2}$-), 4.04 (m, 1H, H-4 GlcNAc), 3.90 (d, *J* = 2.6 Hz, 1H, H-4 Gal2), 3.86 (t, $J_1$ = 9.4 Hz, 1H, H-4 Glc), 3.84 (d, *J* = 2.3 Hz, 1H, H-4 Gal1), 3.82 (dd, $J_1$ = 4.1 Hz, $J_2$ = 7.8 Hz, 1H, H-6a Glc), 3.76 – 3.71 (m, 2H, H-3 Gal1, H-3 GlcNAc), 3.70 – 3.66 (m, 6H, H-2 GlcNAc, H-3 Gal2, H-2 Gal2, H-2 Gal1, H-6a GlcNAc, H-6b Glc), 3.62 (dd, $J_1$ = 1.8 Hz, $J_2$ = 10.4 Hz, 1H, H-6b GlcNAc), 3.54 – 3.47 (m, 3H, H-3 Glc, H-6a Gal1, H-5 GlcNAc), 3.46 – 3.40 (m, 3H, H-6a Gal2, H-6b Gal1, H-2 Glc), 3.37 – 3.34 (m, 2H, H-5 Gal1, H-5 Gal2), 3.28 (m, 1H, H-6b Gal2), 3.18 (d, $J_1$ = 3.9 Hz, $J_2$ = 9.5 Hz, 1H, H-5 Glc).

$^{13}$C NMR (151 MHz, $CDCl_3$): *δ* 56.2 (C2 GlcNAc), 68.0 (C6 Gal2), 68.1 (C6 Gal1), 68.3 (C6 Glc, C6 GlcNAc), 72.0 (-O$\underline{C}H_2$Ph), 74.1 (C5 Gal2, C5 Gal1), 74.6 (C4 Gal2), 75.5 (-O$\underline{C}H_2$Ph), 75.9 (-O$\underline{C}H_2$Ph), 76.0 (-O$\underline{C}H_2$Ph), 76.5 (-O$\underline{C}H_2$Ph), 76.8 (C5 GlcNAc), 76.8 (C4 Gal2), 76.9 (C4 GlcNAc), 77.1 (C4 Gal1), 79.3 (C5 Glc), 79.8 (C2 Glc), 79.9 (C3 GlcNAc), 80.0 (C3 Gal1), 80.1 (C3 Gal2), 82.2 (C5 GlcNAc), 80.3 (C2 Gal), 82.2 (C2 Glc), 86.4 (C1 Glc), 100.5 (C1 GlcNAc), 102.4 (C1 Gal), 102.8 (C1 Gal2), 127.6, 127.7, 127.7, 127.9, 128.0, 128.1, 128.2, 128.4, 128.6, 128.9, 129.0, 129.1, 129.3, 129.4, 129.5, 130.1, 133.0 (C Ar).

### 5. Synthesis of target glycans with diverse terminal epitopes

**General procedure for glycosylation of lactosyl acceptors with glucuronic acid donor.** A mixture of acceptor and donor (1 equiv and 2 equiv, respectively) were twice coevaporated from anhydrous toluene and then dried in high *vacuo* for 2 h. It was taken up in DCM to a final concentration of donor and acceptor of 40 mM and 20 mM, respectively. To this solution 4Å molecular sieves was added and the mixture stirred for 2 h at RT. The solution was cooled to -40

℃, and then a solution of TMSOTf in DCM (3 equiv in total) was added dropwise and the reaction mixture left stirring allowing the temperature gradually to rise to -20 ℃. The progress of the glycosylation was monitored by TLC and if necessary an additional portions of donor was added to drive the reaction to completion. When TLC analysis showed almost no acceptor remaining, triethylamine was added to quench the reaction. The mixture was filtered through a pad of celite and washed with saturated aqueous $NaHCO_3$. The aqueous phase was decanted, and the DCM layer was dried ($Na_2SO_4$) and concentrated to dryness. The residue was dissolved in a minimum amount of toluene/Acetone (1/1, v/v) and applied to SX-1 size exclusion chromatography to separate the product from unreacted starting materials. The products was characterized by MALDI-TOF MS and NMR.

**General procedure for saponification and intralactone formation reactions.** To a solution of compounds **19a-c** in dioxane was added 1 M aqueous KOH until the pH reached 11 and the mixture was stirred vigorously at 40 ℃. The progress of the reaction was monitored by TLC until disappearance of starting material. The reaction mixture was concentrated under reduced pressure and the residue redissolved in methanol. The solution was stirred for 18 h until all benzoyl ester were removed. The pH of the solution was adjusted to neutral with 1 M HCl and then concentrated under reduced pressure. The residue was taken up in chloroform and washed with sat. aqueous $NaHCO_3$. The procedure was repeated until the chloroform layer was clear. The aqueous phase was decanted, and the organic layer was dried ($Na_2SO_4$), concentrated under reduced pressure and the residue dried *in vacuo* before subjected to the next reaction.

The crude glucuronyl carboxylates were dissolved in $Ac_2O$ and heated to 85 ℃ for 2 h, after which TLC analysis indicated the formation of a 3,6-lactone. The reaction mixture was cooled to RT, after which pyridine, acetic anhydride, and catalytic amount of DMAP were added. The reaction mixture stirred for 18 h and the progress of the reaction was monitored by MALDI-TOF MS. The mixture was evaporated to dryness under reduced pressure, twice coevaporated form toluene to ensure the complete removal of acetic anhydride and acetic acid. The crude product was dissolved in a minimum amount of toluene and applied on silica-gel column chromatography eluting with a gradient of EtOAc in toluene (5% → 20%). Fractions containing product were collected, concentrated under reduced pressure and used in the next step after full characterized by NMR and MS.

**General protocol for methanolysis of lactones.** To 100 mM solution of anhydrous NaOAc in anhydrous methanol, 3Å MS were added and the mixture was stirred for 2 h under an inert atmosphere. Compounds **20a-c** were dissolved in anhydrous chloroform (1 mL) and the methanolic solution of sodium acetate (6 mL) was added and the reaction mixture was stirred for 2 h or until the starting material was fully consumed. After completion of the reaction, the solution was buffered by the addition of 50 mL glacial acetic acid and the solvent was evaporated under a stream of $N_2$. The crude product was taken up in toluene and transferred to Eppendorf tube and solid particles removed by centrifugation. The supernatant was applied to a silica-gel column and elutied with 20% EtOAc in toluene to afford the product.

**General protocol for *O*-sulfation.** Compounds **21a-c** were dissolved in pyridine to which $SO_3$Py (20 equiv) was added. The reaction mixture stirred until TLC indicated complete consumption of the alcohol, after which methanol was added to quench unreacted pyridinium sulfur trioxide. The mixture was concentrated under reduced pressure at RT. The residue was dissolved in methanol and then passed through Dowex $Na^+$ ion exchange resin to convert the pyridinium form of sulfates to more stable sodium salts.

**General procedure for anomeric α-fluoride installation.** Compounds x-x were dissolved in dry toluene and evaporated to dryness under rotary evaporator. The procedure was repeated 3-times and further dried under high-vacuum. The obtained residue was dissolved in ultra-dry dichloromethane, to which flame-dried molecular sieves (4 Å) were added. The reaction mixture stirred at RT for 2 h, after which 2 equivalents of Barluenga's reagent (Bis(pyridine)iodonium(I) tetrafluoroborate) was added. The reaction mixture was rapidly cooled to -40 $^{o}$C and HF/Py (40 equiv. by HF) was added, and the reaction mixture was left stirring at that temperature for 30 min. TLC (Tol:acetone, 5:1) indicated the consumption of the starting material and formation of a new spot corresponding to the glycosyl fluoride (hydrolyzed product does not run on TLC in the same solvent system). After the completion of the reaction, it was diluted with dichloromethane and poured into saturated $NaHCO_3$ solution. Organic phase was separated, washed with 10% $Na_2S_2O_4$ solution, dried over anhydrous $Na_2SO_4$ and concentrated. Crude residue was applied to SX-1 size exclusion column chromatography and eluted with tol:acetone (1:1) to obtain glycosyl fluorides.

**General procedure for global deprotection.** Compounds **21a-c** and **22a-c** were dissolved in 1:1 mixture of dioxane and water to which Degussa type $Pd(OH)_2/C$ catalyst was added along with 50 mL of acetic acid. The reaction mixture was stirred overnight under $H_2$ atmosphere. In case of incomplete hydrogenation, additional catalyst was added, and the reaction mixture was stirred for a further 18 h under an atmosphere of $H_2$. The formation of desired products was confirmed with ESI-TOF MS in negative mode. The catalyst was filtered off over a pad of compressed celite, the filter washed with water and the supernatant lyophilized. The resulting white fluffy material were dissolved in water and the pH was adjusted to 8. The solution was incubated at 37 °C to hydrolyze acetyl and methyl esters. The reaction mixture was freeze-dried, and the residue applied to P2 biogel size exclusion column chromatography using aqueous 10 mM $NH_4HCO_3$ as the eluent. Fractions containing product were were collected and lyophilized.

**Methyl 2,3-di-O-benzoyl-4-O-benzyl-*β*-D-glucopyranosyluronate)-(1→3)-2,4-6-tri-O-benzyl-*β*-D-galactopyranosyl)-(1→4)-(2-azido-3,6-di-O-benzyl-2-deoxy-*β*-D-glucopyranosyl)-(1→3)-2,4-6-tri-O-benzyl-*β*-D-galactopyranosyl)-(1→4)-2,3,6-tri-O-benzyl-1-phenylthio-*β*-D-glucopyranoside (13)**

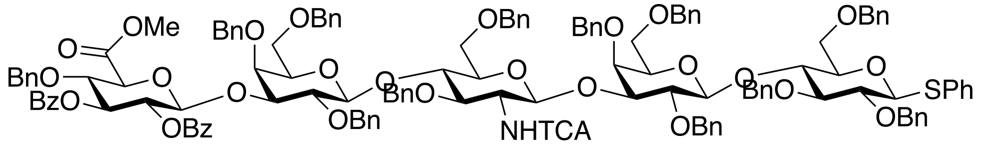


Compound **11** (320 mg, 0.17 mmol) was reacted with donor **12** (250 mg, 0.34 mmol) followed the general protocol for glycosylations to give **13** (260 mg, 65%). MALDI-TOF MS *m/z* $C_{137}H_{136}Cl_3NO_{28}S$ $(M+Na)^+$ calculated 2405.97, found 2406.24.

$^1$H NMR (600 MHz, $CDCl_3$) δ 7.89 (dd, $J_1$ = 1.3 Hz, $J_2$ = 7.9 Hz, 2H, Bz), 7.89 (dd, $J_1$ = 1.4 Hz, $J_2$ = 8.3 Hz, 2H, Bz), 7.52 – 7.48 (m, 2H, Bz), 7.43–7.04 (m, 100H, Ar-H), 6.61 (d, $J_1$ = 4.9 Hz, $J_2$ = 7.8 Hz 1H, -N*H*TCA), 5.06 (d, *J* = 11.1 Hz, 1H, -OCH*H*Ph), 5.01 – 4.93 (m, 2H, $-OCH_2Ph$), 4.86 (d, *J* = 10.8 Hz, 1H, -OCH*H*Ph), 4.83 (d, *J* = 10.1 Hz, 1H, -OCH*H*Ph), 4.80 – 4.72 (m, 3H, -$OCH_2Ph$), 4.71 – 4.65 (m, 2H, $-OCH_2Ph$), 4.62 – 4.56 (m, 2H, $-OCH_2Ph$), 4.56 – 4.33 (m, 2H, -$OCH_2Ph$), 4.33 – 4.23 (m, 7H, $-OCH_2Ph$), 4.20 (d, *J* = 11.4 Hz, 1H, -OCH*H*Ph), 4.20 (d, *J* = 10.1 Hz, 2H, -OC$\underline{H_2}$Ph) 3.76 (-COO*Me*).

$^{13}$C NMR (151 MHz, $CDCl_3$): $\delta$ 52.8 (-COO*Me*), 73.1 (-O*C*$H_2$Ph), 73.2 (-O*C*$H_2$Ph), 73.4 (-O*C*$H_2$Ph), 73.5 (-O*C*$H_2$Ph), 73.8 (-O*C*$H_2$Ph), 74.0 (-O*C*$H_2$Ph), 74.5 (-O*C*$H_2$Ph), 74.8 (-O*C*$H_2$Ph), 75.0 (-O*C*$H_2$Ph), 75.1 (-O*C*$H_2$Ph), 75.2 (-O*C*$H_2$Ph), 75.4 (-O*C*$H_2$Ph), 75.6 (-O*C*$H_2$Ph), 125.3, 127.1, 127.2, 127.3, 127.4, 127.5, 127.6, 127.7, 127.8, 127.9, 127.9, 128.0, 128.1, 128.2, 128.3, 128.4, 128.5, 128.9, 129.1, 129.7, 131.9, 133.1, 133.3, 133.7, 136.9, 137.8, 138.0, 138.1, 138.2, 138.3, 138.4, 138.7, 138.8, 139.1, 139.3 (C Ar), 161.6 (-NHTCA), 165.1 (-OO*C* Bz), 165.5 (-OO*C* Bz), 168.4 (-C6 GlcA).

| **13** | H1 | H2 | H3 | H4 | H5 | H6 |
|---|---|---|---|---|---|---|
| Glc | 4.55 | 3.39 | 3.51<br>t, $J$ = 8.4 Hz | 3.91<br>t, $J$ = 9.0 Hz | 3.20<br>dd, $J_1$ = 4.4 Hz, $J_2$ = 10.8 Hz | 3.72 – 3.21 |
| Gal | 4.37 | 3.69 | 3.71 | 3.90<br>d, $J$ = 2.0 Hz | 3.36 | 3.72 – 3.21 |
| GlcNAc | 5.02<br>d, $J$ = 7.8 Hz | 3.72 | 3.73 | 3.99 | 3.39 | 3.72 – 3.21 |
| Gal2 | 4.33 | 3.56 | 3.68 | 3.88<br>d, $J$ = 2.1 Hz | 3.40 | 3.72 – 3.21 |
| GlcA | 5.21<br>d, $J$ = 7.9 Hz | 5.53<br>t, $J$ = 8.0 Hz | 5.72<br>t, J =9.4 Hz | 4.23 | 4.15 | - |

| **13** | C1 | C2 | C3 | C4 | C5 | C6 |
|---|---|---|---|---|---|---|
| Glc | 87.4 | 80.2 | 84.8 | 75.9 | 79.2 | 68.2 |
| Gal | 102.5 | 80.0 | 80.9 | 76.0 | 75.1 | 68.2 |
| GlcNAc | 100.1 | 57.6 | 79.9 | 76.0 | 80.3 | 68.2 |
| Gal2 | 102.8 | 79.9 | 79.5 | 76.0 | 73.2 | 68.2 |
| GlcA | 101.5 | 71.9 | 74.3 | 77.7 | 74.7 | 168.0 |

**2-O-acetyl-4-O-benzyl-*β*-D-glucopyranosylurono-6,3-lactone-(1→3)-2,4-6-tri-O-benzyl-*β*-D-galactopyranosyl)-(1→4)-(2-trichloroacetamido-3,6-di-O-benzyl-2-deoxy-*β*-D-glucopyranosyl)-(1→3)-2,4-6-tri-O-benzyl-*β*-D-galactopyranosyl)-(1→4)-2,3,6-tri-O-benzyl-1-phenylthio-*β*-D-glucopyranoside (14)**

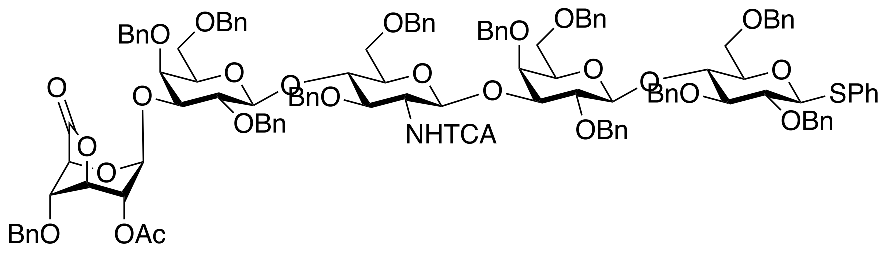


Compound **13** (240 mg, 0.1 mmol) was subjected to lactonization using the general procedure to yield **14** 160 mg (72%). MALDI-TOF MS *m/z* $C_{125}H_{130}Cl_3NO_{26}S$ $(M+Na)^+$ calculated 2223.76, found 2224.49.

$^1$H NMR (600 MHz, $CDCl_3$) δ 7.38–7.14 (m, 70H, Ar-H), 6.54 (d, *J* = 7.1 Hz, 1H, -N*H*TCA), 5.50 (d, *J* = 12.1 Hz, 1H, -OC*H*HPh), 5.48 (m, 2H, H-1 GlcA, H-2 GlcA), 4.97 (m, 1H, H-1 GlcNAc), 4.96 – 4.90 (m, 3H, -$OCH_2Ph$, -OC*H*HPh), 4.86 – 4.74 (m, 4H, H-3 GlcA, -$OCH_2Ph$, -OC*H*HPh), 4.72 – 4.56 (m, 10H, -$OCH_2Ph$, H-1 Glc), 4.54 – 4.40 (m, 7H, -$OCH_2Ph$, H-1 Gal2), 4.36 – 4.10 (m, 9H, -$OCH_2Ph$, H-1 Gal1), 4.08 – 4.00 (m, 2H, H-5 GlcA, H-4 GlcA, H-4 GlcNAc), 3.93 – 3.80 (m, 6H, H-4 Gal1-2, H-4 Glc, H-3 Gal1, H-2 Gal2), 3.77– 3.70 (m, 4H, H-6 Glc, H-2 GlcNAc, H-3 GlcNAc), 3.68 – 3.64 (m, 2H, H-2 Gal1, H-3 Gal1), 3.55 – 3.52 (m, 2H, H-6 Glc), 3.45 (dd $J_1$ = 7.7 Hz, $J_2$ = 2.7 Hz, 1H, H-$6_a$ Gal1), 3.46 – 3.40 (m, 3H, H-6b Glc, H-5 Gal2, H-3 Glc), 3.39 – 3.29 (m, 5H, H-6b Gal, H-2 Glc, H-5 GlcNAc, H-5 Gal), 3.30 (dd, $J_1$ = 6.6 Hz, $J_2$ = 3.3 Hz, 1H, H-$6_b$ Gal), 3.18 (m, 1H, H-5 Glc).

$^{13}$C NMR (151 MHz, $CDCl_3$): *δ* 57.8 (C2 GlcNAc), 68.0 (C6 GlcNAc), 68.2 (C6 Gal1-2, C6 Glc), 67.1 (C5 GlcA), 69.5 (C3 GlcA), 72.8 (-O*C*$H_2$Ph), 73.0 (C2 GlcA), 73.4 (C4 GlcA), 73.1 (-O*C*$H_2$Ph), 73.2 (C5 Gal), 73.3, 73.4 (-O*C*$H_2$Ph), 73.8 (C5 Gal2), 74.6, 74.9 (-O*C*$H_2$Ph), 75.0 (C5 GlcNAc), 75.3 (C5 Glc), 75.9 (C4 Glc), 75.9 (C4 Gal2), 76.3 (C4 Gal1), 76.4 (C4 GlcNAc), 76.5 (C3 Gal1), 77.0 (C3 Glc), 79.4 (C3 Gal2), 81.2 (C2 Gal2), 81.4 (C2 Glc), 83.0 (C3 Glc), 87.6 (C1 Glc), 100.0 (C1 GlcNAc), 100.3 (C1 GlcA), 103.0 (C1 Gal), 103.4 (C1 Gal2), 126.1, 126.5, 126.7, 127.0, 127.2, 127.3, 127.4, 127.5, 127.6, 127.7, 127.8, 127.9, 128.0, 128.1, 128.2, 128.3, 128.4, 128.5, 128.6, 129.3, 137.0. 138.1, 138.3, 138.6, 138.8, 139.0, 139.3 (C Ar), 162.4 (-NHTCA), 169.2 (-$C6_{GlcA}$), 170.5 (-OAc).

**2-O-acetyl-4-O-benzyl-*β*-D-glucopyranosyluronate-(1→3)-2,4-6-tri-O-benzyl-*β*-D-galactopyranosyl)-(1→4)-(2-azido-3,6-di-O-benzyl-2-deoxy-*β*-D-glucopyranosyl)-(1→3)-2,4-**

**6-tri-O-benzyl-$\beta$-D-galactopyranosyl)-(1→4)-2,3,6-tri-O-benzyl-$\beta$-D-glucopyranoside (4)**

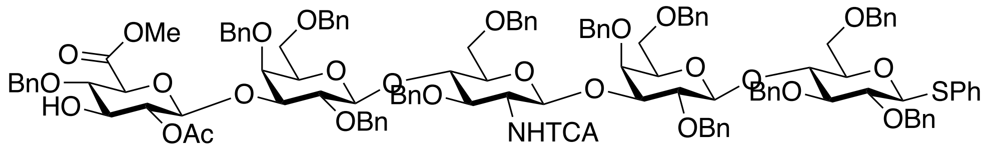


Compound **4** (90 mg, 0.04 mmol, 74%) was obtained from **14** (120 mg, 0.54 mmol) according to the general procedure.

MALDI-TOF MS *m/z* $C_{139}H_{149}Cl_3N_2O_{30}$ $(M+Na)^+$ calculated 2255.84, found 2256.70.

$^1$H NMR (600 MHz, $CDCl_3$) δ 7.38–7.04 (m, 75H, Ar-H), 6.6 (d, *J* = 8.1 Hz, 1H, -*NH*TCA), 5.14 (d, *J* = 12.7 Hz, 2H, $CH_2$ Cbz), 5.1 (d, *J* = 7.5 Hz, 1H, H-1 GlcNAc), 4.9 (d, *J* = 9.2 Hz, 1H, -OC*H*HPh), 4.92 - 4.86 (m, 5H, -OC*H*HPh, -$OCH_2$Ph, H-1 GlcA, H-2 GlcA), 4.8 (m, 4H, -$OCH_2$Ph), 4.70 – 4.60 (m, 5H, -$OCH_2$Ph, -OC*H*HPh), 4.51 – 4.41 (m, 7H, -OC*H*HPh, -$OCH_2$Ph, -$NCH_2$Ph), 4.38 (d, *J* = 7.1 Hz, 1H, H-1Gal2), 4.35 (d, J = 7.2 Hz, 1H, H-1 Gal1), 4.33 – 4.29 (m, 3H, -OC*H*HPh, -$OCH_2$Ph), 4.27 – 4.22 (m, 2H, H-1 Glc, OC*H*HPh), 4.19 (d, *J* = 11.7 Hz, 1H, OCH*H*Ph), 4.13 (d, J = 11.8 Hz, 1H, OCH*H*Ph), 4.03 (t, *J* = 8.8 Hz, 1H, H-4 GlcNAc), 3.92 – 3.83 (m, 4H, H-4 Glc, H-5 GlcA, H-4 Gal1-2), 3.87 – 3.83 (m, 1H, -OC*H*H linker), 3.82 (t, *J* = 8.7 Hz, 1H, H-4 GlcA), 3.78 – 3.60 (m, 13H, H-3 GlcNAc, H-6a GlcNAc, H-2 GlcNAc, H-3 GlcA, H-3 Gal1-2, H-2 Gal1-2, H-6 Gal1-2, -COO*Me*), 3.56 (d, *J* = 10.1 Hz, 1H, H-6b Gal), 3.50 (m, 1H, H-6a Glc), 3.46 – 3.42 (m, H-3 Glc, H-5 GlcNAc), 3.42 – 3.30 (m, 6H, -OC*H*H linker, H-6b Gal, H-5 Gal1-2, H-2 Glc), 3.28 – 3.10 (m, 4H, H-6b Glc, H-5 Glc, -$CH_2$ linker), 1.90 (s, 3H, -OAc), 1.64 – 1.42 (m, 4H, -$CH_2$ linker), 1.34 – 1.19 (m, 2H, -$CH_2$ linker).

$^{13}$C NMR (151 MHz, $CDCl_3$): $\delta$ 21.0 (-COO*C*$H_3$), 23.1 (-*C*$H_2$ linker), 27.8 (-*C*$H_2$ linker), 29.5 (-*C*$H_2$ linker), 46.7 (-*C*$H_2$ linker), 50.4 (-N*C*$H_2$Ph), 52.4 (-OMe GlcA), 57.8 (C2 GlcNAc), 67.1 (-*C*$H_2$ Cbz), 67.9 (C6 GlcNAc), 68.2 (C6 Glc, C6 Gal1-2), 69.6 (-*C*$H_2$ linker), 73.1 (-O*C*$H_2$Ph), 73.2 (C5 Gal1,2), 73.3 (-O*C*$H_2$Ph), 73.4 (-O*C*$H_2$Ph), 73.8 (C2 GlcA), 74.1 (C5 GlcA), 74.9 (-O*C*$H_2$Ph), 75.0 (-O*C*$H_2$Ph), 75.0 (C5 Glc), 75.1 (-O*C*$H_2$Ph), 75.3 (C5 GlcNAc), 75.4 (C3 GlcA), 75.5 (-O*C*$H_2$Ph), 76.2 (C4 Gal1-2), 76.3 (C4 Glc), 76.5 (C4 GlcNAc), 78.1 (C3 GlcNAc), 79.7 (C4 GlcA), 79.9 (C3 Gal1-2), 80.0 (C3 Gal1-2), 80.3 (C2 Gal1-2), 80.3 (C2 Gal1-2), 81.7 (C2 Glc), 82.7 (C3 Glc), 101.1 (C1 GlcNAc), 101.3 (C1 GlcA), 102.5 (C1 Gal1), 102.7 (C1 Gal2), 103.6 (C1 Glc), 127.4, 127.5, 127.6, 127.7, 127.8, 127.9, 128.0, 128.1, 128.2, 128.3, 128.4, 128.55, 128.6 137.6, 137.9, 138.0, 138.2, 138.3, 138.7, 138.9 (C Ar), 139.0 (-COO Cbz), 161.5 (-NHTCA), 168.5

(-*C*OOMe GlcA), 170.5 (-*C*OOCH$_3$ OAc).

**Methyl 2-O-acetyl-4-O-benzyl-*β*-D-glucopyranosyluronate)-(1→3)-(2,4-6-tri-O-benzyl-*β*-D-galactopyranosyl)-(1→4)-(2-trichloracetamido-3,6-di-O-benzyl-2-deoxy-*β*-D-glucopyranosyl)-(1→3)-2,4-6-tri-O-benzyl-*β*-D-galactopyranosyl)-(1→4)-2,3,6-tri-O-benzyl--α-D-glucopyranosyl fluoride (5)**

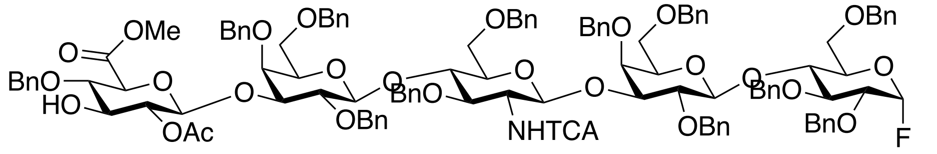


Compound **5** (76 mg, 0.035 mmol, 94%) was obtained from **4** (85 mg, 0.037 mmol) according to the general procedure for anomeric fluoride installation, using Barluenga's reagent (28 mg, 0.074 mmol) and HF/Py (145 µL, 1.48 mmol).
MALDI-TOF MS *m/z* $C_{120}H_{129}Cl_3FNO_{27}$ $(M+Na)^+$ calculated 2165.67, found 2166.74.

$^1$H NMR (600 MHz, CDCl$_3$) δ 7.49 (d, J = 7.4 Hz, 1H, Ar-H), 7.39–7.08 (m, 70H, Ar-H), 6.66 (d J = 8.3 Hz, 1H, -*NH*TCA), 5.03 – 4.88 (m, 5H, -OCH$_2$Ph), 4.83 – 4.60 (m, 10H, -O*C*H$_2$Ph, -OC*H*HPh), 4.54 – 4.44 (m, 5H, -OCH$_2$Ph), 4.39 – 4.12 (m, 8H, -OCH$_2$Ph), 4.14 (d, J = 11.7 Hz, 1H, -OC*H*HPh), 3.70 (s, 3H, -COO*Me*).

$^{13}$C NMR (151 MHz, CDCl$_3$): *δ* 52.7 (-OMe GlcA), 73.2, 73.3, 73.4, 73.5, 73.6, 73.9, 74.3, 75.0, 75.1, 75.6 (-O*C*H$_2$Ph), 127.2, 127.3, 127.4, 127.5, 127.6, 127.9, 128.0, 128.1, 128.2, 128.3, 128.4, 128.5, 128.6, 129.0, 137.6, 137.8, 137.9, 138.2, 138.3, 138.6, 138.9, 139.0, 139.2, 139.9 (C Ar).
$^{19}$F NMR (564 MHz, CDCl$_3$): *δ* -149.66 (dd, $J_1$ = 25.6 Hz, $J_2$ = 53.1 Hz)

| **5** | H1 | H2 | H3 | H4 | H5 | H6 |
|---|---|---|---|---|---|---|
| Glc | 5.42 – 5.37<br>dd, $J_{HH}$ = 2.7 Hz<br>$J_{FH}$ = 53.3 Hz | 3.41 | 3.46 | 3.95<br>t, J = 9.1 Hz | 3.25 | 3.76 |
| Gal | 4.37<br>d, J = 7.1 Hz | 3.70 | 3.74 | 3.91<br>d, J = 3.0 Hz | 3.38 | 3.48<br>3.40 |

| | | | | | | |
|---|---|---|---|---|---|---|
| GlcNAc | 5.08<br>d, J = 8.4 Hz | 3.72 | 3.76 | 4.02<br>t, J = 8.4 Hz | 3.46 | 3.64<br>3.50 |
| Gal(2) | 4.23<br>d, J = 7.8 Hz | 3.71 | 3.80 | 3.86<br>d, *J* = 2.6 Hz | 3.38 | 3.40 |
| GlcA | 4.91 | 4.90 | 3.72 | 3.82 | 3.90 | - |

| **5** | C1 | C2 | C3 | C4 | C5 | C6 |
|---|---|---|---|---|---|---|
| Glc | 104.4<br>–<br>106.4 | 78.8 | 81.0 | 75.0 | 75.1 | 68.0 |
| Gal | 103.0 | 79.6 | 79.7 | 76.4 | 73.3 | 68.2 |
| GlcNAc | 99.9 | 57.8 | 79.6 | 76.3 | 75.5 | 68.1 |
| Gal(2) | 102.6 | 79.8 | 79.9 | 76.1 | 73.3 | 68.2 |
| GlcA | 101.5 | 73.9 | 75.7 | 77.7 | 76.1 | |

**Methyl 2-O-acetyl-3-O-sulfate-*β*-D-glucopyranosyluronate-(1→3)-*β*-D-galactopyranosyl)-(1→4)-(2-acetamido-2-deoxy-*β*-D-glucopyranosyl)-(1→3)-*β*-D-galactopyranosyl)-(1→4)-α-D-glucopyranosyl fluoride (S2)**

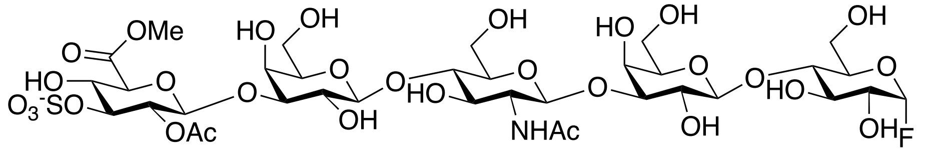


Compound **S2** (13 mg, 0.013 mmol, 82%) was obtained from **5** (35 mg, 0.016 mmol) according using the general protocols for *O*-sulfation and hydrogenation.

MALDI-TOF MS *m/z* $C_{36}H_{59}FNO_{30}S$ $(M\text{-}H)^-$ calculated 1036.90, found 1036.83.

$^1H$ NMR (600 MHz, $CDCl_3$) δ 3.75 (s, 3H, -COO<u>*Me*</u>), 2.03 (s, 3H, -NHCO<u>$CH_3$</u>), 1.95 (s, 3H, -OCO<u>$CH_3$</u>).

$^{13}C$ NMR (151 MHz, $CDCl_3$): *δ* 20.1 (-NHCO<u>$CH_3$</u>), 22.3 (-OCO<u>$CH_3$</u>), 53.2 (-OMe GlcA), 170.3 (-NH<u>*C*</u>O$CH_3$), 174.7 (-O<u>*C*</u>O$CH_3$).

| **S2** | H1 | H2 | H3 | H4 | H5 | H6 |
|---|---|---|---|---|---|---|
| Glc | 5.56<br>dd, $J_{HH}$ = 2.6 Hz<br>$J_{FH}$ = 53.5 Hz | 3.62 | 3.63 | 3.77 | 3.48 | 3.83<br>3.76 |

| | | | | | | |
|---|---|---|---|---|---|---|
| Gal | 4.35<br>d, J = 7.1 Hz | 3.47 | 3.61 | 3.91<br>d, J = 3.0 Hz | 3.47 | 3.67 |
| GlcNAc | 4.62<br>d, J = 8.3 Hz | 3.70 | 3.64 | 3.86 | 3.61 | 3.60 |
| Gal(2) | 4.37<br>d, J = 7.8 Hz | 3.47 | 3.61 | 3.86<br>d, *J* = 2.6 Hz | 3.47 | 3.67 |
| GlcA | 4.87 | 4.87 | 4.42 | 3.73 | 4.12<br>d, J = 9.6 Hz | - |

| **S2** | C1 | C2 | C3 | C4 | C5 | C6 |
|---|---|---|---|---|---|---|
| Glc | 107.0 | 74.4 | 82.0 | 70.3 | 74.8 | 59.7 |
| Gal | 102.7 | 74.8 | 79.7 | 74.8 | 69.9 | 60.9 |
| GlcNAc | 102.9 | 52.3 | 83.3 | 72.8 | 75.0 | 59.6 |
| Gal(2) | 102.7 | 74.8 | 79.9 | 74.8 | 69.9 | 60.9 |
| GlcA | 101.9 | 71.6 | 80.6 | 76.2 | 74.4 | 170.6 |

**3-O-sulfate-*β*-D-glucopyranosyluronate-(1→3)-*β*-D-galactopyranosyl-(1→4)-(2-acetamido-2-deoxy-*β*-D-glucopyranosyl-(1→3)-*β*-D-galactopyranosyl-(1→4)-*β*-D-glucopyranoside (6)**

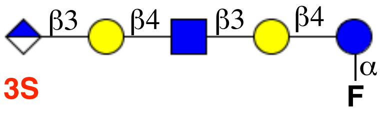


Compound **6** (8 mg, 82%) was obtained from **S2** (10 mg, 0.01 mmol) using the general protocol for saponification. ESI TOF-MS *m/z* $C_{33}H_{54}FNO_{29}S$ $(M - H)^-$ calculated 980.84, found 981.47.

$^1$H NMR (600 MHz, $CDCl_3$) δ 2.03 (s, 3H, -NHCO*<u>CH</u>$_3$*).

$^{13}$C NMR (151 MHz, $CDCl_3$): *δ* 20.1 (-NHCO*<u>CH</u>$_3$*), 170.3 (-NH<u>C</u>OCH$_3$).

| **6** | H1 | H2 | H3 | H4 | H5 | H6 |
|---|---|---|---|---|---|---|
| Glc | 5.61<br>dd, $J_{HH}$ = 2.9 Hz<br>$J_{FH}$ = 53.4 Hz | 3.49 | 3.64 | 3.79 | 3.49 | 3.84<br>3.76 |
| Gal | 4.37<br>d, J = 7.5 Hz | 3.63 | na | 4.1<br>d, J = 3.2 Hz | 3.47 | 3.53 |
| GlcNAc | 4.61<br>d, J = 8.5 Hz | 3.71 | 3.74 | 3.69 | 3.60 | 3.67 |

| | | | | | | |
|---|---|---|---|---|---|---|
| Gal(2) | 4.45<br>d, J = 7.5 Hz | 3.63 | na | 4.07<br>d, *J* = 2.9 Hz | 3.60 | 3.56 |
| GlcA | 4.67 | 3.62 | 4.25<br>t, J = 9.1 Hz | 3.65 | 3.87 | - |

| **6** | C1 | C2 | C3 | C4 | C5 | C6 |
|---|---|---|---|---|---|---|
| Glc | 106.9 | 72.4 | 82.0 | 71.3 | 72.8 | 59.7 |
| Gal | 102.7 | 75.5 | na | 68.2 | 70.4 | 61.0 |
| GlcNAc | 102.9 | 55.1 | 82.4 | 72.8 | 72.6 | 61.0 |
| Gal(2) | 102.7 | 75.5 | na | 68.2 | 70.4 | 57.0 |
| GlcA | 103.4 | 75.0 | 82.0 | 72.2 | 72.4 | 171.4 |

**3-O-sulfate-*β*-D-glucopyranosyluronate-(1→3)-*β*-D-galactopyranosyl)-(1→4)-(2-acetamido-2-deoxy-*β*-D-glucopyranosyl)-(1→3)-*β*-D-galactopyranosyl)-(1→4)-α-D-glucopyranosyl fluoride (15)**

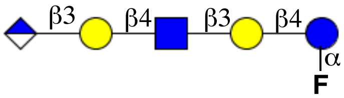


Compound **15** (6 mg, 0.007 mmol, 51%) was obtained from **5** (30 mg, 0.014) using the general protocols for *O*-sulfation, hydrogenation and saponification. ESI TOF-MS *m/z* $C_{33}H_{56}N_2FO_{26}$ $(M - H)^-$ calculated 900.78, found 900.65.

$^1$H NMR (600 MHz, $CDCl_3$) δ 2.00 (s, 3H, -NHCO*CH₃*).

$^{13}$C NMR (151 MHz, $CDCl_3$): *δ* 21.0 (-NHCO*CH₃*), 172.1 (-NH*C*OCH₃).

| **15** | **H1** | **H2** | **H3** | **H4** | **H5** | **H6** |
|---|---|---|---|---|---|---|
| Glc | 5.47<br>dd, $J_{HH}$ =<br>2.98Hz<br>$J_{FH}$ = 52.4<br>Hz | 3.28 | 3.55 | 3.65 | na | 3.78 – 3.66 |
| Gal | 4.35<br>d, *J* = 7.6 Hz | 3.50 | 3.62 | 4.08<br>d, *J* = 3.1 Hz | na | 3.66 – 3.60 |
| GlcNAc | 4.71<br>d, *J* = 8.3 Hz | 3.70 | 3.55 | 3.84 | na | 3.88 |
| Gal2 | 4.47<br>d, *J* = 8.1 Hz | 3.64 | 3.72 | 4.10<br>d, *J* = 3.0 Hz | na | 3.66 – 3.60 |
| GlcA | 4.70<br>d, *J* = 7.8 Hz | 3.36 | 3.48 | 3.42 | na | - |

| **15** | C1 | C2 | C3 | C4 | C5 | C6 |
|---|---|---|---|---|---|---|
| Glc | 104.4 | 72.1 | 83.0 | 70.8 | 70.3 | 60.1 |
| Gal | 101.2 | 74.5 | na | 66.2 | na | 62.0 |
| GlcNAc | 102.4 | 56.2 | 81.5 | 71.4 | 72.3 | 61.1 |
| Gal(2) | 103.3 | 75.1 | na | 68.6 | na | 62.0 |
| GlcA | 103.4 | 75.2 | 81.0 | 71.1 | 70.4 | 170.1 |

### 6. Synthesis of lyso- and full-length glycosphingolipids

**General procedure for the synthesis of *lyso*-glycosphingolipids via enzymatic en bloc transfer of glycosyl fluorides.** To a solution of α-glycosyl fluorides (10 mM) in NaOAc buffer (pH = 5.0, 50 mM), erythron-sphingosine (15 mM) was added. The reaction mixture was vortexed to suspend all solid residues, after which the suspension was supplemented with dimethoxyethane to final concentration of 1% (v/v). Endoglycoceramidase (EGC II D351S/E314Y) enzyme was added (1% w/w per substrate) and the reaction mixture incubated at 37 ºC overnight. The progress of transformations was monitored by LC-MS ESI and the reaction was quenched upon the consumption of the fluoride donors. Crude reaction mixture was applied on pre-conditioned benchtop C18 reverse-phase chromatography cartridges. Trace of unreacted or hydrolyzed glycosyl fluorides were removed with 5% acetonitrile in $H_2O$, after which the products were eluted with 50% acetonitrile in $H_2O$. Product containing fractions were identified by TLC spotting and confirmed with ESI-TOF mass spectrometry, which were later pooled and lyophilized to afford white fluffy glycosyl sphingosines.

**General protocol for acylation of glycosyl sphingosines.** Lyso-glycosphingolipids were suspended in methanol, to which triethylamine (10 equiv.) was added. Later, a solution of stearoyl N-hydroxy succinimide in THF (5 equiv. per $NH_2$) was added dropwise under rigorous stirring. The reaction mixture was left stirring at RT overnight. The progress of fatty acid acylation was monitored by TLC and confirmed by MALDI-TOF mass spectroscopy. After completion of the reaction, the reaction mixture was concentrated and unreacted stearic acid, and hydroxy succinimide were precipitated out of water. Solid particles were removed by centrifugation and the supernatant applied on C18 reverse-phase chromatography. Glycosyl ceramides were eluted with 60% (v/v) acetonitrile in $H_2O$.

**(*2S,3R,4E*)-2-Amino-3-hydroxyoctadec-4-en-1-yl-3-O-sulfo-*β*-*D*-glucopyranosyl-(1→3)-*β*-D-galactopyranosyl)-(1→4)-2-acetamido-2-deoxy-*β*-D-glucopyranosyl-(1→3)-*β*-D-galactopyranosyl-(1→4)-*β*-D-glucopyranoside (16)**

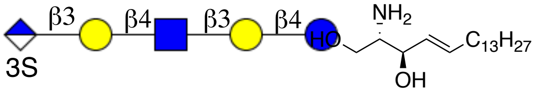


Compound **6** (5 mg, 0.0037 mmol) and *erythro*-sphingosine (2.3 mg, 0.0076 mmol) are condensed using general protocol for sphingosine installation to afford 3.9 mg (62%) of lyso-sulfoglucuronyl paragloboside (**16**).
ESI TOF-MS *m/z* $C_{51}H_{91}N_2O_{31}S$ $(M - H)^-$ calculated 1243.2591, found 1243.2594.

$^1$H NMR (600 MHz, $D_2O$) δ: (non-carbohydrate): 5.75 (m, 1H, =CH (sphingosine)), 5.38 (dt, $J_1$ = 6.3 Hz, $J_2$ = 15.6 Hz, 1H, HC=C (sphingosine)), 4.18 (m, 1H, -C*H*NH$_2$(sphingosine)), 3.68 (m, 2H, -OC*H*$_2$- C1-sphingosine), 3.56 (m, 2H, =CHC*H*$_2$-), 1.99 (m, 2H, -C*H*$_2$), 1.88 (s, 3H, -NHAc), 1.32 (m, 4H, -CH$_2$), 1.25 – 1.14 (m, 44H, -CH$_2$), 0.79 (t, J = 6.2 Hz, 3H, CH$_3$).

$^{13}$C NMR (151 MHz, $D_2O$) δ: (non-carbohydrate): 135.7 (=*C*H (sphingosine)), 127.5 (H*C*=C (sphingosine)), 70.2 (-*C*HNH$_2$ (sphingosine)), 58.1 (-O*C*H$_2$- C1-sphingosine), 58.1 (=CHC*H*$_2$-), 32.1 (-*C*H$_2$), 32.0 (-CH$_2$), 30.5 – 28.3 (-CH$_2$), 21.7 (-NHAc), 12.8 (CH$_3$).

| **16** | H1 | H2 | H3 | H4 | H5 | H6 |
|---|---|---|---|---|---|---|
| Glc | 4.33 | 3.47 | 3.67 | 3.76 | 3.44 | 3.88<br>3.77 |
| Gal | 4.35<br>d, *J* = 7.5 Hz | 3.62 | Na | 3.95<br>d, *J* = 3.2 Hz | 3.41 | 3.50 |
| GlcNAc | 4.67<br>d, *J* = 7.6 Hz | 3.70 | 3.75 | 3.75 | 3.55 | 3.78 |
| Gal(2) | 4.46<br>d, *J* = 7.8 Hz | 3.53 | na | 4.01 | 3.42 | 3.56 |
| GlcA | 4.67 | 3.62 | 4.20 | 3.69 | 3.86 | - |

| **16** | C1 | C2 | C3 | C4 | C5 | C6 |
|---|---|---|---|---|---|---|
| Glc | 101.3 | 71.4 | 81.9 | 71.5 | 71.8 | 60.0 |
| Gal | 102.5 | 73.5 | Na | 67.7 | 71.6 | 61.3 |
| GlcNAc | 102.4 | 55.0 | 82.5 | 72.1 | 72.3 | 61.3 |
| Gal(2) | 101.7 | 75.0 | Na | 68.5 | 70.1 | 57.5 |
| GlcA | 103.5 | 74.0 | 83.4 | 72.0 | 72.6 | 171.4 |

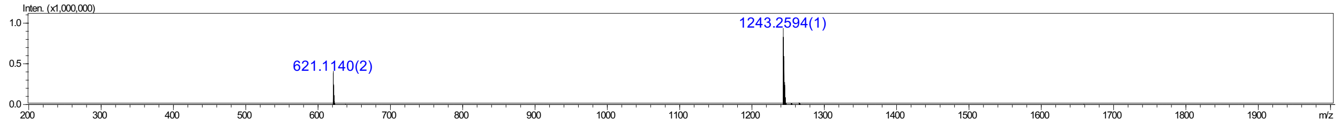


**(*2S,3R,4E*)-2-Amino-3-hydroxyoctadec-4-en-1-yl-3-β-*D*-glucopyranosyl-(1→3)-*β*-D-galactopyranosyl)-(1→4)-2-acetamido-2-deoxy-*β*-D-glucopyranosyl-(1→3)-*β*-D-galactopyranosyl-(1→4)-*β*-D-glucopyranoside (17)**

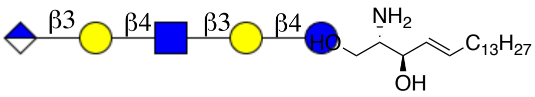


Compound **15** (3 mg, 0.0033 mmol) and *erythro*-sphingosine (1.5 mg, 0.005 mmol) are condensed using general protocol for sphingosine installation to afford 1.6 mg (41%) of lyso-glucuronyl paragloboside (**17**).

ESI TOF-MS *m/z* $C_{51}H_{92}N_2O_{28}$ $(M - H)^-$ calculated 1163.3831, found 1163.3540.

$^1$H NMR (600 MHz, $D_2O$) δ: (non-carbohydrate): 5.76 (m, 1H, =CH (sphingosine)), 5.38 (dt, $J_1$ = 6.4 Hz, $J_2$ = 15.3 Hz, 1H, HC=C (sphingosine)), 4.19 (m, 1H, -C*H*NH$_2$(sphingosine)), 3.68 (m, 2H, -OC*H*$_2$- C1-sphingosine), 3.54 (m, 2H, =CHC*H*$_2$-), 1.98 (m, 2H, -C*H*$_2$), 1.88 (s, 3H, -NHAc), 1.25 – 1.14 (m, 44H, -CH$_2$), 0.81 (t, J = 6.2 Hz, 3H, CH$_3$).

$^{13}$C NMR (151 MHz, $D_2O$) δ: (non-carbohydrate): 135.6 (=*C*H (sphingosine)), 127.4 (H*C*=C (sphingosine)), 70.6 (-*C*HNH$_2$ (sphingosine)), 57.4 (-O*C*H$_2$- C1-sphingosine), 58.2 (=CHC*H*$_2$-), 32.2 (-*C*H$_2$), 32.1 (-CH$_2$), 32.5 – 26.3 (-CH$_2$), 20.1 (-NHAc), 13.0 (CH$_3$).

| 17 | H1 | H2 | H3 | H4 | H5 | H6 |
|---|---|---|---|---|---|---|
| Glc | 4.30 | 3.20 | 3.51 | 3.66 | na | 3.76 – 3.64 |
| Gal | 4.32<br>d, $J$ = 7.6 Hz | 3.51 | 3.61 | 4.09<br>d, $J$ = 2.9 Hz | na | 3.62 – 3.61 |
| GlcNAc | 4.70<br>d, $J$ = 8.3 Hz | 3.71 | 3.54 | 3.84 | na | 3.86 |
| Gal2 | 4.45<br>d, $J$ = 8.1 Hz | 3.62 | 3.74 | 4.10<br>d, $J$ = 3.0 Hz | na | 3.60 – 3.58 |
| GlcA | 4.68 | 3.30 | 3.52 | 3.43 | na | - |

| 17 | C1 | C2 | C3 | C4 | C5 | C6 |
|---|---|---|---|---|---|---|
| Glc | 101.2 | 72.0 | 82.6 | 71.9 | 71.0 | 58.1 |
| Gal | 101.3 | 74.3 | Na | 66.6 | Na | 62.4 |
| GlcNAc | 102.1 | 56.1 | 81.4 | 71.5 | 72.2 | 60.4 |
| Gal(2) | 103.1 | 74.6 | Na | 68.2 | Na | 62.6 |
| GlcA | 102.1 | 74.2 | 81.1 | 72.4 | 70.4 | 170.6 |

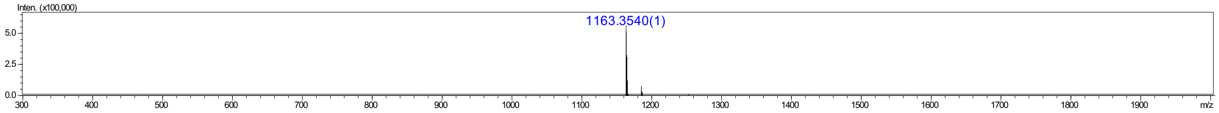


**(*2S,3R,4E*)-2-hexadecanamido-3-hydroxyoctadec-4-en-1-yl-*β*-*D*-glucopyranosyl-(1→3)-*β*-D-galactopyranosyl)-(1→4)-2-acetamido-2-deoxy-*β*-D-glucopyranosyl-(1→3)-*β*-D-galactopyranosyl-(1→4)-*β*-D-glucopyranoside (2a)**

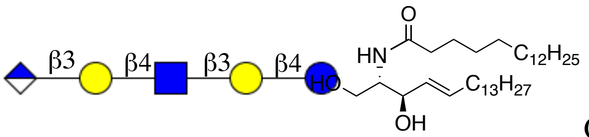


Compound **17** (1 mg, 0.0008 mmol) was acylated using general protocol for stearic acid installation to afford compound **2a** (0.8 mg, 65%). ESI TOF-MS *m/z* $C_{69}H_{124}N_2O_{29}$ $(M - H)^-$ calculated 1445.74, found 1444.99.

$^1$H NMR (600 MHz, $D_2O$) δ: (non-carbohydrate): 5.58 (m, 1H, =CH (sphingosine)), 5.35 (dd, $J_1$ = 7.9 Hz, $J_2$ = 15.1 Hz, 1H, HC=C (sphingosine)), 4.09 (d, $J_1$ = 4.1 Hz, $J_2$ = 10.5 Hz, 1H, -OCH*H*-

C1-sphingosine), 3.97 (t, J = 8.3 Hz, 1H, -C*H*NH$_2$(sphingosine)), 3.47 (m, 1H, -OC*H*H- C1-sphingosine), 2.07 (t, J = 7.6 Hz, 2H, α-CH$_2$-stearic acid), 1.93 (m, 2H, -C*H*$_2$), 1.89 (s, 3H, -NHAc), 1.49 (m, 4H, -CH$_2$), 1.24 – 1.15 (m, 73H, -CH$_2$), 0.80 (t, J = 6.2 Hz, 3H, CH$_3$).

$^{13}$C NMR (151 MHz, D$_2$O) δ: (non-carbohydrate): 133.9 (=*C*H (sphingosine)), 130.0 (H*C*=C (sphingosine)), 72.0 (-*C*HNH$_2$ (sphingosine)), 68.0 (-O*C*H$_2$- C1-sphingosine), 58.1 (=CHC*H*$_2$-), 35.2 (α-*C*H$_2$-stearic acid -), 32.5 (-CH$_2$), 32.8 (-CH$_2$), 30.6 – 28.1 (-CH$_2$), 26.1 (-CH$_2$), 22.4 (-CH$_2$), 21.5 (-NHAc), 13.1 (CH$_3$).

| **2a** | H1 | H2 | H3 | H4 | H5 | H6 |
|---|---|---|---|---|---|---|
| Glc | 4.20 | 3.16 | 3.67 | 3.76 | na | 3.81 |
| Gal | 4.24 | 3.54 | Na | 3.91<br>d, *J* = 2.9 Hz | na | 3.65 |
| GlcNAc | 4.55<br>d, *J* = 7.8 Hz | 3.70 | 3.72 | 3.74 | 3.57 | 3.70 |
| Gal(2) | 4.36<br>d, *J* = 8.1 Hz | 3.50 | na | 3.95<br>d, *J* = 2.9 Hz | na | 3.62 |
| GlcA | 4.54 | 3.20 | 3.22 | 3.69 | 3.74 | - |

| **2a** | C1 | C2 | C3 | C4 | C5 | C6 |
|---|---|---|---|---|---|---|
| Glc | 101.1 | 71.4 | 80.1 | 71.5 | Na | 60.1 |
| Gal | 102.4 | 73.5 | Na | 68.5 | na | 61.0 |
| GlcNAc | 101.8 | 56.2 | 82.1 | 72.1 | 70.1 | 60.2 |
| Gal(2) | 102.9 | 75.1 | Na | 68.8 | na | 61.2 |
| GlcA | 101.6 | 79.0 | 79.6 | 72.0 | na | 168.2 |

**(*2S,3R,4E*)-2-hexadecanamido-3-hydroxyoctadec-4-en-1-yl-3-O-sulfo-*β*-*D*-glucopyranosyl-(1→3)-*β*-D-galactopyranosyl)-(1→4)-2-acetamido-2-deoxy-*β*-D-glucopyranosyl-(1→3)-*β*-D-galactopyranosyl-(1→4)-*β*-D-glucopyranoside (1a)**

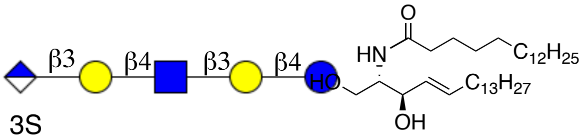


Compound **16** (2 mg, 0.0016 mmol) was acylated using general protocol for stearic acid installation to afford compound **1a** (1.9 mg, 79%). ESI TOF-MS *m/z* $C_{69}H_{123}N_2O_{32}S$ $(M - H)^-$ calculated 1524.79, found 1524.22.

$^1$H NMR (600 MHz, $D_2O$) δ: (non-carbohydrate): 5.60 (m, 1H, =CH (sphingosine)), 5.37 (dd, $J_1$ = 7.6 Hz, $J_2$ = 15.1 Hz, 1H, HC=C (sphingosine)), 4.10 (d, $J_1$ = 4.6 Hz, $J_2$ = 10.4 Hz, 1H, -OCH*H*-C1-sphingosine), 3.98 (t, J = 8.1 Hz, 1H, -C*H*$NH_2$(sphingosine)), 3.51 (m, 1H, -OC*H*H- C1-sphingosine), 2.06 (t, J = 7.6 Hz, 2H, α-$CH_2$-stearic acid), 1.93 (m, 2H, -C*H*$_2$), 1.89 (s, 6H, -NHAc), 1.47 (m, 4H, -$CH_2$), 1.26 – 1.16 (m, 67H, -$CH_2$), 0.81 (t, J = 6.1 Hz, 3H, $CH_3$).

$^{13}$C NMR (151 MHz, $D_2O$) δ: (non-carbohydrate): 133.9 (=*C*H (sphingosine)), 130.1 (H*C*=C (sphingosine)), 74.0 (-*C*H$NH_2$ (sphingosine)), 68.5 (-O*C*$H_2$- C1-sphingosine), 58.6 (=CHC*H*$_2$-), 35.4 (α-*C*$H_2$-stearic acid -), 33.5 (-$CH_2$), 30.1 – 28.3 (-$CH_2$), 26.2 (-$CH_2$), 22.6 (-$CH_2$), 21.5 (-NHAc), 21.3 (-NHAc), 13.0 ($CH_3$).

| **1a** | H1 | H2 | H3 | H4 | H5 | H6 |
|---|---|---|---|---|---|---|
| Glc | 4.19 | 3.19 | 3.67 | 3.76 | 3.44 | 3.80 |
| Gal | 4.25<br>d, *J* = 8.7 Hz | 3.62 | Na | 3.95<br>d, *J* = 2.8 Hz | 3.41 | 3.67 |
| GlcNAc | 4.55<br>d, *J* = 7.6 Hz | 3.68 | 3.75 | 3.75 | 3.55 | 3.76 |
| Gal(2) | 4.35<br>d, *J* = 8.8 Hz | 3.53 | na | 4.01 | 3.42 | 3.60 |
| GlcA | 4.57<br>d, J = 7.8 Hz | 3.46 | 4.20 | 3.69<br>d, *J* = 3.3 Hz | 3.86 | - |

| 1a | C1 | C2 | C3 | C4 | C5 | C6 |
|---|---|---|---|---|---|---|
| Glc | 103.3 | 71.4 | 81.9 | 71.5 | 71.8 | 60.4 |
| Gal | 103.6 | 73.5 | Na | 68.5 | 71.6 | 61.1 |
| GlcNAc | 102.8 | 55.0 | 82.5 | 72.1 | 72.3 | 60.5 |
| Gal(2) | 103.1 | 75.0 | Na | 68.8 | 70.1 | 61.0 |
| GlcA | 103.7 | 79.0 | 83.3 | 72.0 | 72.6 | 171.4 |

### 7. Synthesis of glycosphingolipids having hexasaccharide backbones

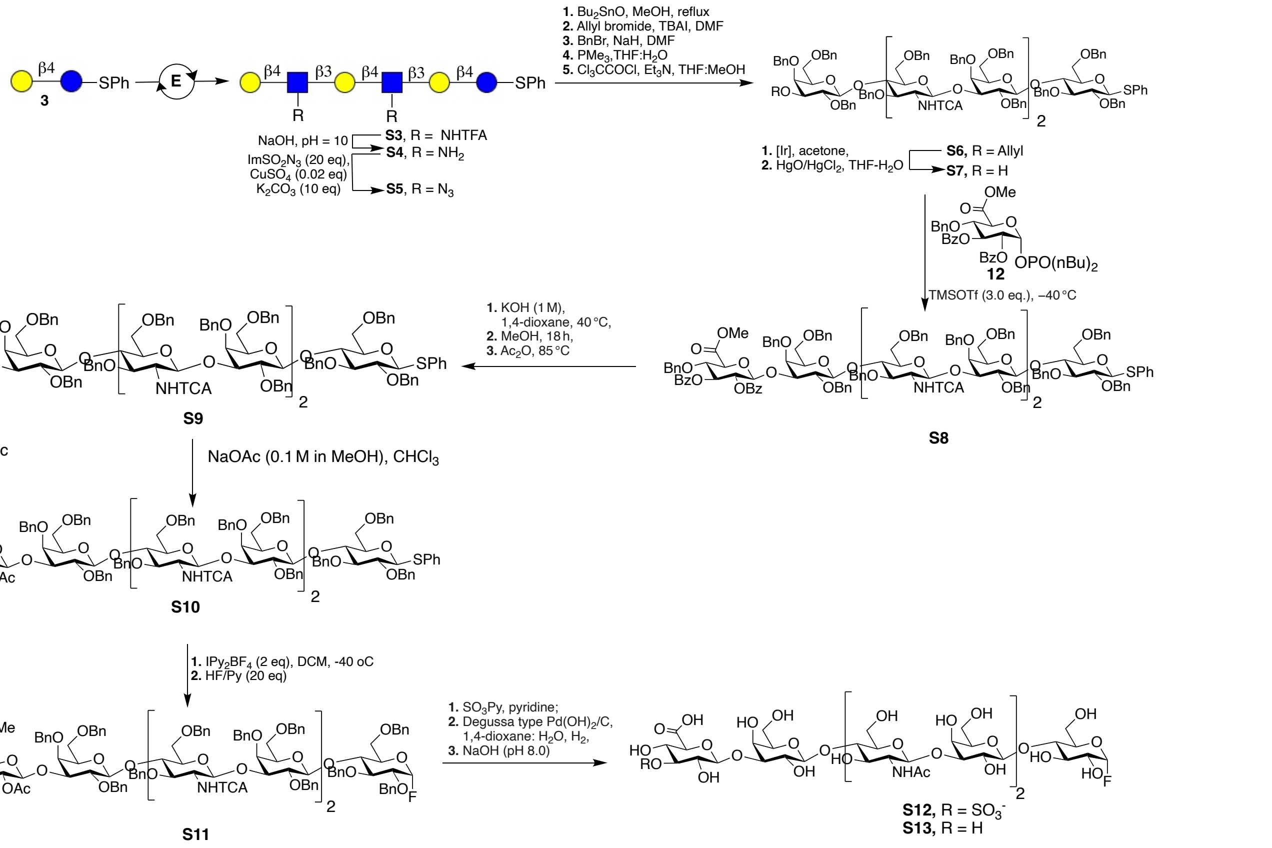


**Figure S4.** Chemoenzymatic synthesis of glycosyl a-fluorides of sulfated and unsulfated glucuronyl paraglobosides.

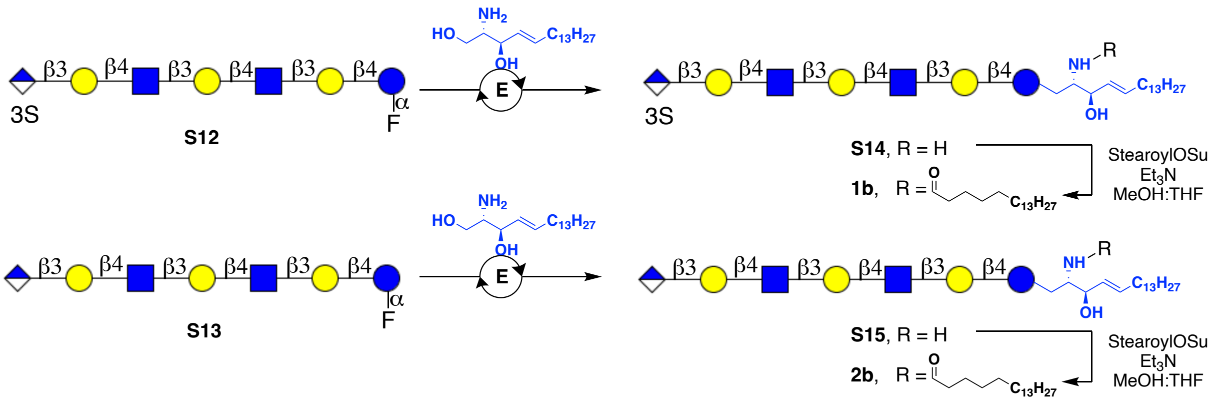


**Figure S5.** Enzymatic *en bloc* installation of sphingosine with subsequent stearic acid acylation.

**β-D-galactopyranosyl-(1→4)-2-trifluoroacetamido-2-deoxy-β-D-glucopyranosyl-(1→3)-β-D-galactopyranosyl-(1→4)-2-trifluoroacetamido-2-deoxy-β-D-glucopyranosyl-(1→3)-β-D-galactopyranosyl-(1→4)-1-phenylthio-β-D-glucopyranoside (S3)**

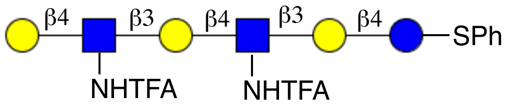


Compound **7** (500 mg, 0.57 mmol) was reacted with UDP-GlcNHTFA (570 mg, 0.86 mmol) and UDP-Gal (508 mg, 0.90 mmol) according to the general protocol for LacNHTFA installation to yield compound **15c** (556 mg, 78%).

MALDI-TOF MS *m/z* $C_{46}H_{66}F_6N_2O_{30}S$ $(M+Na)^+$ calculated 1296.08, found 1295.89.

$^1H$ NMR (600 MHz, $D_2O$) δ: (non-carbohydrate): 7.38 – 7.04 (m, 5H, SPh).
$^{13}C$ NMR (151 MHz, $D_2O$): $\delta$ (non-carbohydrate): 132.5, 129.3, 128.1 (C Ar).

| **S3** | H1 | H2 | H3 | H4 | H5 | H6 |
|---|---|---|---|---|---|---|
| Glc | 4.52 | 3.22 | 3.54 | 3.58 | 3.18 | 3.88 – 3.80 |
| Gal | 4.29<br>d, *J* = 7.1 Hz | 3.45 | 3.58 | 4.04<br>d, *J* = 2.4 Hz | 3.25 | 3.74 – 3.70 |
| GlcNAc | 4.72 | 3.71 | 3.56 | 3.68 | 3.37 | 3.78 – 3.76 |
| Gal(2) | 4.36<br>d, *J* = 7.7 Hz | 3.41 | 3.62 | 4.11<br>d, *J* = 2.8 Hz | 3.25 | 3.74 – 3.70 |

| | | | | | | |
|---|---|---|---|---|---|---|
| GlcNAc2 | 4.70 | 3.71 | 3.55 | 3.65 | 3.28 | 3.80 – 3.76 |
| Gal (3) | 4.30 | 3.38 | 3.61 | 3.95 | 3.34 | 3.62 – 3.54 |

| **S3** | H1 | H2 | H3 | H4 | H5 | H6 |
|---|---|---|---|---|---|---|
| Glc | 87.3 | 73.1 | 74.1 | 78.2 | 74.1 | 59.5 |
| Gal | 103.1 | 70.1 | 81.1 | 69.1 | 72.1 | 61.1 |
| GlcNAc | 102.8 | 55.8 | 74.9 | 77.6 | 74.6 | 60.1 |
| Gal(2) | 103.6 | 75.0 | 73.1 | 70.1 | 71.7 | 61.4 |
| GlcNAc2 | 102.5 | 56.2 | 74.2 | 78.3 | 75.6 | 60.1 |
| Gal (3) | 102.1 | 71.4 | 72.6 | 69.4 | 75.4 | 61.0 |

**β-D-galactopyranosyl-(1→4)-2-amino-2-deoxy-β-D-glucopyranosyl-(1→3)-β-D-galactopyranosyl-(1→4)-2-amino-2-deoxy-β-D-glucopyranosyl-(1→3)-β-D-galactopyranosyl-(1→4)-1-phenylthio-β-D-glucopyranoside (S4)**

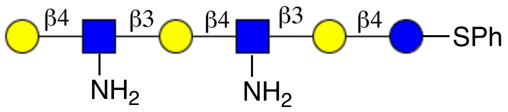


Compound **S3** (550 mg, 0.43 mmol) was subjected to TFA removal according to the general procedure to yield 420 mg (87%) of **S4**.

MALDI-TOF MS *m/z* $C_{42}H_{68}N_2O_{28}S$ $(M+Na)^+$ calculated 1104.04, found 1101.08.

$^1$H NMR (600 MHz, $D_2O$) δ: (non-carbohydrate): 7.51 – 7.11 (m, 5H, SPh).

$^{13}$C NMR (151 MHz, $D_2O$): *δ* (non-carbohydrate): 126.0, 129.1, 131.2 (C Ar).

Carbohydrate region:

| | H1 | H2 | H3 | H4 | H5 | H6 |
|---|---|---|---|---|---|---|
| Glc | 4.67 | 3.21 | 3.51 | 3.53 | 3.16 | 3.72 – 3.68 |
| Gal | 4.35<br>d, J = 8.2 Hz | 3.57 | 3.65 | 3.95 | 3.32 | 3.62 |
| GlcNAc | 4.84<br>d, J = 7.6 Hz | 2.91 | 3.67 | 3.71 | na | 3.68 |
| Gal(2) | 4.33 | 3.43 | 3.49 | 4.05 | 3.34 | 3.62 |

| | | | | | | |
|---|---|---|---|---|---|---|
| | d, J = 8.0 Hz | | | d, J = 2.6 Hz | | |
| GlcNAc2 | 4.85<br>d, $J$ = 8.6 Hz | 3.0 | 3.71 | 3.66 | na | 3.68 |
| Gal(3) | 4.36<br>d, $J$ = 7.9 Hz | 3.56 | 3.55 | 3.96<br>d, $J$ = 2.3 Hz | 3.37 | 3.62 |

| | C1 | C2 | C3 | C4 | C5 | C6 |
|---|---|---|---|---|---|---|
| Glc | 88.7 | 73.2 | 74.4 | 76.0 | 75.2 | na |
| Gal | 103.0 | 72.4 | 78.0 | 69.9 | 75.9 | na |
| GlcNAc | 102.1 | 57.3 | 76.3 | 76.5 | 75.5 | na |
| Gal(2) | 103.1 | 71.2 | 82.3 | 69.9 | 75.8 | na |
| GlcNAc (2) | 102.2 | 57.4 | 74.8 | 77.3 | 76.0 | na |
| Gal(3) | 103.8 | 71.4 | 81.9 | 69.8 | 76.2 | na |

**β-D-galactopyranosyl-(1→4)-2-azido-2-deoxy-β-D-glucopyranosyl-(1→3)-β-D-galactopyranosyl-(1→4)-2-azido-2-deoxy-β-D-glucopyranosyl-(1→3)-β-D-galactopyranosyl-(1→4)-1-phenylthio-β-D-glucopyranoside (S5)**

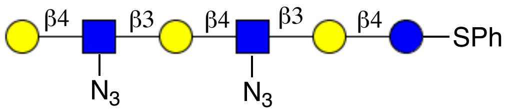


Compound **S5** (356 mg, 0.31 mmol) was obtained from starting material, **S4** (400 mg, 0.36 mmol) following the general protocol for azido transfer. MALDI-TOF MS *m/z* $C_{42}H_{64}N_6O_{28}S$ $(M+Na)^+$ calculated 1156.04, found 1155.87.

$^1$H NMR (600 MHz, $D_2O$) δ: (non-carbohydrate): 7.47 – 7.09 (m, 5H, SPh).

$^{13}$C NMR (151 MHz, $D_2O$): $\delta$ (non-carbohydrate): 127.0, 128.9, 130.9 (C Ar).

| **S5** | H1 | H2 | H3 | H4 | H5 | H6 |
|---|---|---|---|---|---|---|
| Glc | 4.69 | 3.22 | 3.54 | 3.66 | 3.18 | 3.78 – 3.72 |
| Gal | 4.37<br>d, J = 7.7 Hz | 3.55 | 3.55 | 4.00 | na | 3.60 |
| GlcNAc | 4.86<br>d, $J$ = 8.1 Hz | 3.30 | 3.67 | 3.72 | 3.25 | 3.66 |
| Gal(2) | 4.34<br>d, $J$ = 8.0 Hz | 3.49 | na | 4.10<br>d, $J$ = 2.8 Hz | na | 3.62 |

| | | | | | | |
|---|---|---|---|---|---|---|
| GlcNAc2 | 4.85<br>d, $J$ = 7.8 Hz | 3.34 | 3.75 | 3.70 | na | 3.66 |
| Gal(3) | 4.30 | 3.46 | 3.55 | 3.95 | na | 3.62 |

| **S5** | C1 | C2 | C3 | C4 | C5 | C6 |
|---|---|---|---|---|---|---|
| Glc | 89.3 | 72.8 | 75.8 | 76.8 | 72.7 | 61 |
| Gal | 102.4 | 73.1 | 83.8 | 68.5 | na | 62.4 |
| GlcNAc | 103.5 | 67.4 | 74.8 | 76.3 | na | 58.4 |
| Gal(2) | 103.2 | 71.9 | na | 68.8 | na | 62.8 |
| GlcNAc(2) | 103.8 | 68.0 | 74.5 | 75.8 | na | 58.4 |
| Gal(3) | 103.6 | 70.6 | 81.1 | 68.0 | na | 62.8 |

**3-O-allyl-2,4-6-tri-O-benzyl-*β*-D-galactopyranosyl)-(1→4)-(2-azido-3,6-di-O-benzyl-2-deoxy-*β*-D-glucopyranosyl)-(1→3)- (2,4-6-tri-O-benzyl-*β*-D-galactopyranosyl)-(1→4)-(2-azido-3,6-di-O-benzyl-2-deoxy-*β*-D-glucopyranosyl)-(1→3)-2,4-6-tri-O-benzyl-*β*-D-galactopyranosyl)-(1→4)-2,3,6-tri-O-benzyl-1-phenylthio-*β*-D-glucopyranoside (S6)**

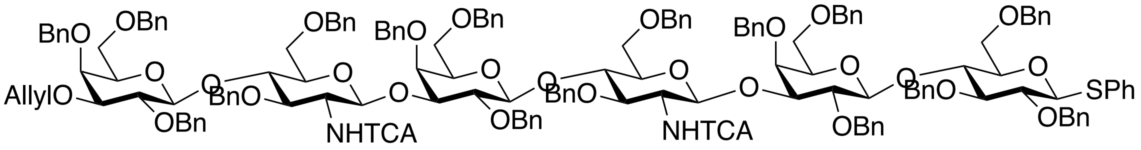


Compound **S6** (350 mg, 46% over 4 steps) was obtained from compound **S5** (300 mg, 0.26 mmol) using the general protocols for tin-mediated allylation, benzylation and reductive acylation using appropriate amounts of each reagent for corresponding reactions.

MALDI-TOF MS *m/z* $C_{161}H_{166}Cl_6N_2O_{30}S$ $(M+Na)^+$ calculated 2876.83, found 2876.29.

$^1$H NMR (600 MHz, Chloroform-d, non-carbohydrate) δ 7.55–7.03 (m, 82H, Ar-H), 6.65 (d, J = 7.8 Hz, 1H, -N*H*TCA), 6.60 (d, J = 8.2 Hz, 1H, -N*H*TCA), 5.94 (m, 1H, $CH_2$=*C<u>H</u>*-$CH_2$-), 5.33 (dd, $J_1$ = 1.8 Hz, $J_2$ = 17.8 Hz, 1H, CH*<u>H</u>*=CH-$CH_2$-), 5.18 (dd, $J_1$ = 1.2 Hz, $J_2$ = 10.8 Hz, 1H, C*<u>H</u>*H=CH-$CH_2$-), 5.0 (d, $J$ = 10.8 Hz, 1H, -OC*<u>H</u>*$_2$Ph), 4.97 (d, $J$ = 11.4 Hz, 2H, -OC*<u>H</u>*$_2$Ph), 4.91 (d, $J$ = 11.4 Hz, 2H, -OC*<u>H</u>*$_2$Ph), 4.87 (d, $J$ = 10.9 Hz, 2H, -OC*<u>H</u>*$_2$Ph), 4.83 – 4.71 (m, 7H, -

OC$\underline{H_2}$Ph), 4.68 (d, $J$ = 12.3 Hz, 1H, -OC$\underline{H_2}$Ph), 4.61 (d, $J$ = 10.3 Hz, 2H, -OC$\underline{H_2}$Ph), 4.53 – 4.43 (m, 8H, -OC$\underline{H_2}$Ph), 4.40 – 4.23 (m, $J$ = 11.4 Hz, 8H, -OC$\underline{H_2}$Ph), 4.22 – 4.12 (m 6H, -OC$\underline{H}$HPh, $CH_2$=CH-C$\underline{H_2}$-).

[13]C NMR (151 MHz, $CDCl_3$ non-carbohydrate): $\delta$ 71.6 (-$\underline{C}H_2$ allyl), 73.1 (-O$\underline{C}H_2$Ph), 73.2 (-O$\underline{C}H_2$Ph), 73.4 (-O$\underline{C}H_2$Ph), 73.5 (-O$\underline{C}H_2$Ph), 73.9 (-O$\underline{C}H_2$Ph), 74.6 (-O$\underline{C}H_2$Ph), 74.8 (-O$\underline{C}H_2$Ph), 74.9 (-O$\underline{C}H_2$Ph), 75.0 (-O$\underline{C}H_2$Ph), 75.4 (-O$\underline{C}H_2$Ph), 75.5 (-O$\underline{C}H_2$Ph), 118.8 ($\underline{C}H_2$=CH-$CH_2$-), 127.2 – 128.8 (C Ar), 131.9 (C Ar), 135.0 ($CH_2$=$\underline{C}$H-$CH_2$-).

Carbohydrate region:

| **S6** | H1 | H2 | H3 | H4 | H5 | H6 |
|---|---|---|---|---|---|---|
| Glc | 4.61<br>d, J = 10.16 Hz | 3.37 | 3.51 | 3.90 | 3.20<br>dd, $J_1$ = 3.8 Hz, $J_2$ = 10.5 Hz | 3.70<br>3.61 |
| Gal | 4.38 | 3.78 – 3.67 | 3.78 – 3.67 | 3.85<br>d, J = 2.8 Hz | 3.28 – 3.25 | 3.45<br>3.34 |
| GlcNAc | 5.10<br>d, J = 7.4 Hz | 3.79 | 3.79 | 4.05 – 4.00 | 3.53 | 3.82<br>3.77 |
| Gal(2) | 4.42<br>d, J = 8.0 Hz | 3.78 – 3.67 | 3.78 – 3.67 | 3.90 – 3.91 | 3.28 – 3.25 | 3.49<br>3.41 |
| GlcNAc2 | 5.03<br>d, J = 7.4 Hz | 3.76 | 3.73 | 4.05 – 4.00 | 3.36 | 3.82<br>3.77 |
| Gal(3) | 4.37 | 3.78 – 3.67 | 3.78 – 3.67 | 3.90 – 3.91 | 3.28 – 3.25 | 3.27 |

| **S6** | C1 | C2 | C3 | C4 | C5 | C6 |
|---|---|---|---|---|---|---|
| Glc | 87.2 | 82.1 | 75.4 | 76.1 | 79.2 | 68.4 |
| Gal | 102.9 | 80.1 | 79.8 | 73.4 | 82.3 | 68.2 |
| GlcNAc | 100.5 | 57.6 | 78.1 | 76.4 | 75.4 | 68.2 |
| Gal(2) | 103.1 | 80.1 | 79.8 | 76.0 | 82.3 | 68.4 |
| GlcNAc2 | 100.5 | 57.6 | 78.1 | 76.4 | 75.3 | 68.2 |
| Gal(3) | 102.6 | 80.1 | 79.8 | 76.3 | 82.3 | 68.0 |

**2,4-6-tri-O-benzyl-*β*-D-galactopyranosyl)-(1→4)-(2-azido-3,6-di-O-benzyl-2-deoxy-*β*-D-glucopyranosyl)-(1→3)- (2,4-6-tri-O-benzyl-*β*-D-galactopyranosyl)-(1→4)-(2-azido-3,6-di-O-benzyl-2-deoxy-*β*-D-glucopyranosyl)-(1→3)-2,4-6-tri-O-benzyl-*β*-D-galactopyranosyl)-(1→4)-2,3,6-tri-O-benzyl-1-phenylthio-*β*-D-glucopyranoside (S7)**

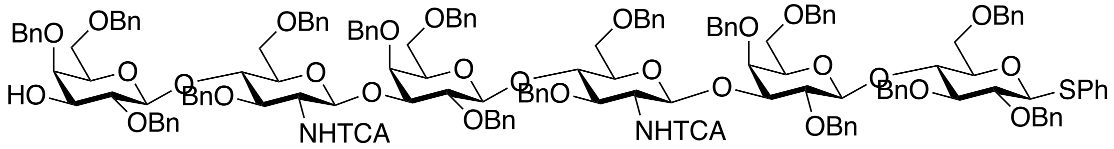


220 mg (71%) of compound **S7** was obtained from the reaction of 300 mg (0.11 mmol) of **S6** with [Ir(cod)(PPh$_2$Me)$_2$]PF$_6$ catalyst (10 mg, 0.012 mmol) and HgCl$_2$ (67 mg, 0.24 mmol) and HgO (10 mg, 0.044 mmol) according to general procedure for allyl removal.

MALDI-TOF MS *m/z* $C_{158}H_{162}Cl_6N_2O_{30}S$ $(M+Na)^+$ calculated 2836.77, found 2838.47.

$^1$H NMR (600 MHz, Chloroform-d) δ 7.52–7.04 (m, 96H, Ar-H), 6.67 (d, *J* = 8.3 Hz, 1H, -N*H*TCA GlcNAc2), 6.60 (d, *J* = 8.0 Hz, 1H, -N*H*TCA GlcNAc1), 5.00 (d, J = 10.8 Hz, 1H, -OCH$_2$Ph), 4.98 (d, J = 11.4 Hz, 1H, -OCH$_2$Ph), 4.97 (d, J = 11.5 Hz, 1H, -OCH$_2$Ph), 4.94 (d, J = 10.8 Hz, 1H, -OCH$_2$Ph), 4.87 (d, J = 10.8 Hz, 1H, -OCH$_2$Ph), 4.85 (d, J = 11.4 Hz, 1H, -OCH$_2$Ph), 4.82 (d, *J* = 11.5 Hz, 1H, -OC*H*HPh), 4.77 – 4.65 (m, 8H, -OCH$_2$Ph), 4.62 – 4.59 (m, 2H, -OCH$_2$Ph), 4.54 – 4.46 (m, 6H, -OCH$_2$Ph), 4.43 – 4.31 (m, 6H, -OCH$_2$Ph), 4.26 (d, J = 11.3 Hz, 1H, -OCH$_2$Ph), 4.23 (d, J = 11.8 Hz, 1H, -OCH$_2$Ph), 4.19 (d, J = 12.0 Hz, 1H, -OCH$_2$Ph), 4.14 (d, J = 11.8 Hz, 1H, -OCH$_2$Ph).

$^{13}$C NMR (151 MHz, CDCl$_3$): *δ* 73.1, 73.2, 73.3, 73.5, 73.8, 74.0, 74.8, 74.9, 75.1, 75.2, 75.5, 75.6 (-OCH$_2$Ph), 127.1, 127.2, 127.3, 127.4, 127.5, 127.6, 127.7, 127.8, 127.9, 127.9, 128.0, 128.1, 128.2, 128.3, 128.4, 128.5, 128.6, 128.8, 131.9 (C$_{Ar}$),

| **S7** | H1 | H2 | H3 | H4 | H5 | H6 |
|---|---|---|---|---|---|---|
| Glc | 4.56<br>d, *J* = 9.6 Hz | 3.39 | 3.51 | 3.91 | 3.19<br>dd, $J_1$ = 3.8 Hz,<br>$J_2$ = 10.5 Hz | 3.84<br>3.75 |
| Gal | 4.38 | 3.54 | na | 3.83<br>d, *J* = 2.5 Hz | na | 3.48 |

| | | | | | | |
|---|---|---|---|---|---|---|
| GlcNAc | 5.13<br>d, $J$ = 7.6 Hz | 3.75 | 3.84 | 4.07<br>t, $J$ = 8.0 Hz | 3.53 | 3.75<br>3.69 |
| Gal(2) | 4.42<br>d, $J$ = 8.0 Hz | 3.76 – 3.73 | na | 3.90 | na | 3.41<br>3.27 |
| GlcNAc2 | 5.04<br>d, $J$ = 6.9 Hz | 3.73 | 3.74 | 4.02<br>t, $J$ = 8.4 Hz | 3.36 | 3.75<br>3.69 |
| Gal(3) | 4.37 | 3.76 – 3.73 | na | 3.91 | na | 3.34 |

| **S7** | C1 | C2 | C3 | C4 | C5 | C6 |
|---|---|---|---|---|---|---|
| Glc | 87.2 | 80.3 | 80.5 | 76.4 | 79.4 | 68.4 |
| Gal | 102.9 | 74.8 | na | 75.5 | na | 67.9 |
| GlcNAc | 100.0 | 67.8 | 77.8 | 76.4 | 75.3 | 68.2 |
| Gal(2) | 103.1 | 80.0 | na | 76.1 | na | 68.2 |
| GlcNAc2 | 100.2 | 67.8 | 77.8 | 76.4 | 75.3 | 68.2 |
| Gal(3) | 102.6 | 80.0 | na | 76.1 | na | 67.9 |

**Methyl 2,3-di-O-benzoyl-4-O-benzyl-$\beta$-D-glucopyranosyluronate)-(1→3)-2,4-6-tri-O-benzyl-$\beta$-D-galactopyranosyl)-(1→4)-(2-azido-3,6-di-O-benzyl-2-deoxy-$\beta$-D-glucopyranosyl)-(1→3)- (2,4-6-tri-O-benzyl-$\beta$-D-galactopyranosyl)-(1→4)-(2-azido-3,6-di-O-benzyl-2-deoxy-$\beta$-D-glucopyranosyl)-(1→3)-2,4-6-tri-O-benzyl-$\beta$-D-galactopyranosyl)-(1→4)-2,3,6-tri-O-benzyl-1-phenylthio-$\beta$-D-glucopyranoside (S8)**

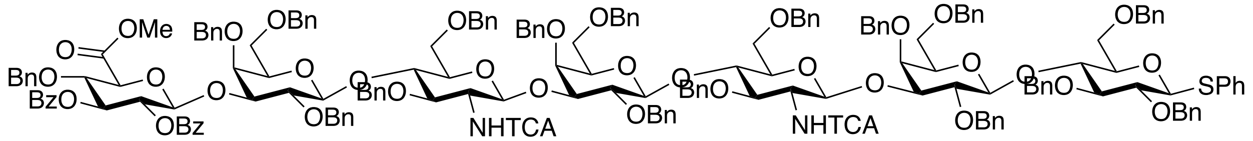


Compound **S7** (200 mg, 0.07 mmol) was reacted with donor **12** (96 mg, 0.14 mmol) according to general glycosylation protocol to give **S8** (170 mg, 74%). MALDI-TOF MS *m/z* $C_{186}H_{186}Cl_6N_2O_{38}$ $(M+Na)^+$ calculated 3325.26, found 3326.20.

$^{1}$H NMR (600 MHz, Chloroform-d) δ 7.91 (d, $J$ = 7.6 Hz, 2H, Bz), 7.78 (d, $J$ = 7.6 Hz, 2H, Bz), 7.51 (m, 2H, Bz), 7.37–7.01 (m, 105H, Ar-H), 6.62 (m, 2H, -N*H*TCA), 5.05 (d, $J$ = 7.8 Hz, 1H, -OCH*H*Ph), 4.98 – 4.93 (m, 3H, -OCH$_2$Ph), 4.89 – 4.79 (m, 3H, -OCH$_2$Ph), 4.71 – 4.14 (m, 34H, -CH$_2$Ph), 3.48 (-COO*Me* GlcA).

$^{13}$C NMR (151 MHz, CDCl$_3$): $\delta$ 50.5 (-COO*Me*), 73.1, 73.2, 73.5, 73.7, 73.8, 73.9, 74.1, 74.3 74.7, 74.8, 74.9, 75.0, 75.1, 75.2, 75.2 (-O*C*H$_2$Ph), 126.3, 126.5, 127.1, 127.2, 127.3, 127.4, 127.5, 127.6, 127.7, 127.8, 127.9, 128.0, 128.1, 128.2, 128.3, 128.4, 128.5, 129.7, 129.8, 129.9, 132.6, 132.9, 133.1, 133.3, 137.9. 138.0, 138.1, 138.2, 138.7, 138.8, 139.0 ($C_{Ar}$), 139.4 (-COO Cbz), 161.0 (-NHTCA), 164.0 (-OO$C_{Bz}$), 164.1 (-OO$C_{Bz}$), 165.5 (-C6$_{GlcA}$).

| **S8** | H1 | H2 | H3 | H4 | H5 | H6 |
|---|---|---|---|---|---|---|
| Glc | 4.56 | 3.41 | 3.51 | 3.91 | 3.20 d, $J_1$ = 11.2 Hz | 3.84 3.75 |
| Gal | 4.35 | 3.56 | 3.73 | 3.83 d, $J$ = 2.5 Hz | na | 3.48 |
| GlcNAc | 5.03 | 3.78 | 3.65 | 3.99 | 3.36 | 3.72 – 3.21 |
| Gal(2) | 4.36 | 3.70 | 3.73 | 3.99 | 3.39 | 3.72 – 3.21 |
| GlcNAc2 | 5.03 | 3.70 | 3.65 | 3.99 | 3.40 | 3.72 – 3.21 |
| Gal(3) | 4.36 | 3.76 – 3.73 | 3.75 | 4.23 | 4.15 | - |
| GlcA | 5.23 d, $J$ = 7.8 Hz | 5.54 dd, $J_1$ = 1.7 $J_2$ = 9.8 Hz, | 5.73 t, $J$ = 9.7 Hz | 4.22 | 4.21 | |

| **S8** | C1 | C2 | C3 | C4 | C5 | C6 |
|---|---|---|---|---|---|---|
| Glc | 87.3 | 80.1 | 84.6 | 76.4 | 79.3 | 68.4 |
| Gal | 102.9 | 74.8 | na | 75.5 | na | 67.9 |
| GlcNAc | 100.0 | 52.8 | 77.8 | 76.4 | 75.3 | 68.2 |
| Gal(2) | 103.1 | 80.0 | na | 76.1 | na | 68.2 |

| | | | | | | |
|---|---|---|---|---|---|---|
| GlcNAc2 | 100.0 | 57.4 | 77.8 | 76.4 | 75.3 | 68.2 |
| Gal(3) | 102.6 | 80.0 | na | 76.1 | na | 67.9 |
| GlcA | 101.3 | 72.1 | 74.4 | 77.2 | 77.7 | 165.5 |

**Methyl 2-O-acetyl-4-O-benzyl-*β*-D-glucopyranosyluronate)-(1→3)-2,4-6-tri-O-benzyl-*β*-D-galactopyranosyl)-(1→4)-(2-azido-3,6-di-O-benzyl-2-deoxy-*β*-D-glucopyranosyl)-(1→3)-(2,4-6-tri-O-benzyl-*β*-D-galactopyranosyl)-(1→4)-(2-azido-3,6-di-O-benzyl-2-deoxy-*β*-D-glucopyranosyl)-(1→3)-2,4-6-tri-O-benzyl-*β*-D-galactopyranosyl)-(1→4)-2,3,6-tri-O-benzyl-1-phenylthio-*β*-D-glucopyranoside (S9)**

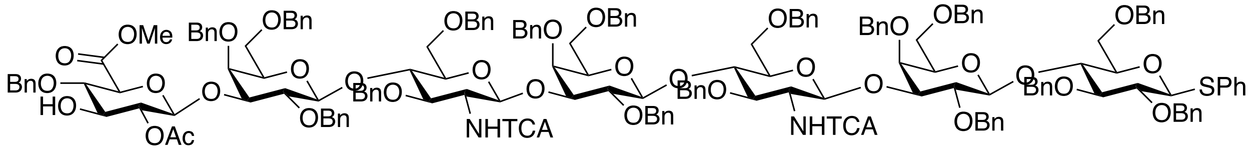


Compound **S9 (**67 mg, 0.021 mmol, 45%) was obtained from **S8** (160 mg, 0.048 mmol) according to general procedure.
MALDI-TOF MS *m/z* $C_{174}H_{180}Cl_6N_2O_{37}S$ $(M+Na)^+$ calculated 3159.09, found 3160.63.

$^1$H NMR (600 MHz, $CDCl_3$) δ 7.55–7.00 (m, 90H, Ar-H), 6.65 (m, 1H, -<u>*NH*</u>$TCA_2$), 6.57 (d, 1H, -<u>*NH*</u>$TCA_1$), 5.01– 4.86 (m, 6H, $-OCH_2Ph$), 4.82 – 4.53 (m, 12H, -O<u>*C*</u>$H_2$Ph, -OC<u>*H*</u>HPh), 4.53 – 4.40 (m, 8H, $-OCH_2Ph$), 4.40 – 4.10 (m, 12H, $-OCH_2Ph$), 3.71(s, 3H, -COO<u>*Me*</u>).

$^{13}$C NMR (151 MHz, $CDCl_3$): *δ* 52.1 (-OMe GlcA), 68.1, 68.2, 73.1, 73.2, 73.3, 73.4, 73.5, 74.2, 74.8, 74.9, 75.0, 75.1, 75.5, 75.6 (-O<u>*C*</u>$H_2$Ph), 127.2, 127.4, 127.5, 127.6, 127.7, 127.8, 127.9, 128.0, 128.1, 128.2, 128.9, 128.5, 128.6, 128.8, 131.9 (C Ar), 165.1 (-<u>*C*</u>OOMe GlcA).

| **S9** | H1 | H2 | H3 | H4 | H5 | H6 |
|---|---|---|---|---|---|---|
| Glc | 4.55 | 3.37 | 3.50 | 3.90 | 3.18<br>d, *J* = 6.6 Hz | 3.71<br>3.66 |
| Gal | 4.36 | 3.70 | 3.73 | 3.90 | 3.39 | 3.46<br>3.40 |

| | | | | | | |
|---|---|---|---|---|---|---|
| GlcNAc | 5.09 | 3.72 | 3.80 | 4.01 | 3.45 | 3.74<br>3.60 |
| Gal(2) | 4.36 | 3.70 | 3.73 | 3.86 | 3.39 | 3.40<br>3.33 |
| GlcNAc2 | 5.03 | 3.72 | 3.79 | 4.01 | 3.35 | 3.74<br>3.60 |
| Gal(3) | 4.36 | 3.76 – 3.73 | 3.73 | 3.90 | 3.39 | 3.30<br>3.25 |
| GlcA | 4.91 | 4.90 | 3.71 | 3.81 | 3.90 | |

| **S9** | C1 | C2 | C3 | C4 | C5 | C6 |
|---|---|---|---|---|---|---|
| Glc | 87.6 | 80.2 | 84.7 | 78.1 | 79.3 | 68.3 |
| Gal | 102.9 | 80.1 | 80.0 | 76.3 | 73.3 | 67.9 |
| GlcNAc | 100.1 | 57.8 | 80.1 | 76.5 | 75.4 | 67.8 |
| Gal(2) | 102.9 | 80.0 | 80.1 | 76.3 | 73.4 | 68.2 |
| GlcNAc2 | 100.3 | 57.8 | 78.2 | 76.5 | 75.2 | 67.8 |
| Gal(3) | 102.9 | 80.0 | 80.1 | 76.3 | 73.4 | 67.9 |
| GlcA | 101.3 | 74.1 | 75.4 | 79.9 | 74.0 | 165.1 |

**Methyl 2-O-acetyl-4-O-benzyl-*β*-D-glucopyranosyluronate)-(1→3)-2,4-6-tri-O-benzyl-*β*-D-galactopyranosyl)-(1→4)-(2-trichloracetamido-3,6-di-O-benzyl-2-deoxy-*β*-D-glucopyranosyl)-(1→3)-(2,4-6-tri-O-benzyl-*β*-D-galactopyranosyl)-(1→4)-(2-trichloracetamido-3,6-di-O-benzyl-2-deoxy-*β*-D-glucopyranosyl)-(1→3)-2,4-6-tri-O-benzyl-*β*-D-galactopyranosyl)-(1→4)-2,3,6-tri-O-benzyl-α-D-glucopyranosyl fluoride (S10)**

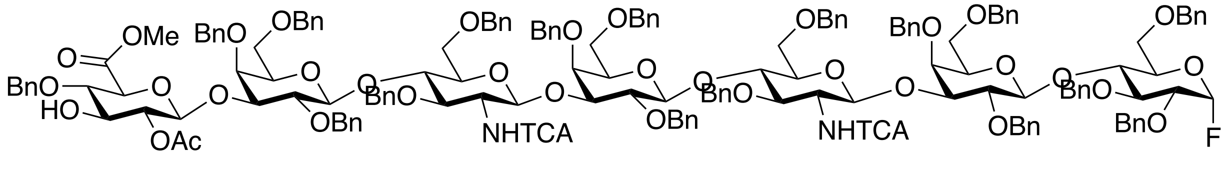


Compound **S10** (33 mg, 0.011 mmol, 57%) was obtained from **S9** (60 mg, 0.019 mmol) according to the general procedure for anomeric fluoride installation, using Barluenga's reagent (14 mg,

0.038 mmol) and HF/Py (74 μL, 0.76 mmol).

MALDI-TOF MS *m/z* $C_{168}H_{175}Cl_6FN_2O_{37}$ $(M+Na)^+$ calculated 3068.91, found 3071.48.

$^1$H NMR (600 MHz, $CDCl_3$) δ 7.40–7.04 (m, 70H, Ar-H), 6.65 (m, 1H, -*NH*TCA$_2$), 6.57 (d, J = 8.0 Hz, 1H, -*NH*TCA$_1$), 4.99 – 4.86 (m, 6H, -OCH$_2$Ph), 4.83 – 4.62 (m, 9H, -O*C*H$_2$Ph, -OC*H*HPh), 4.53 – 4.40 (m, 8H, -OCH$_2$Ph), 4.36 – 4.18 (m, 7H, -OCH$_2$Ph), 4.14 (d, J = 12.40 Hz, 1H, -OC*H*HPh), 3.71(s, 3H, -COO*Me*).

$^{13}$C NMR (151 MHz, $CDCl_3$): *δ* 52.8 (-OMe GlcA), 73.2, 73.3, 73.4, 73.5, 73.6, 73.8, 74.0, 74.1, 74.2, 74.5, 74.8, 74.9, 75.0, 75.1, 75.2, 75.4 (-O*C*H$_2$Ph), 126.2 – 129.8 (C Ar), 162.0 (-NH*TCA*), 167.6 (-*C*OOMe GlcA).

$^{19}$F NMR (564 MHz, $CDCl_3$): *δ* -149.67 (dd, $J_1$ = 25.4 Hz, $J_2$ = 53.5 Hz).

| **S10** | H1 | H2 | H3 | H4 | H5 | H6 |
|---|---|---|---|---|---|---|
| Glc | 5.42 – 5.37<br>dd, $J_{HH}$ = 2.8 Hz<br>$J_{FH}$ = 54.1 Hz | 3.42 | 3.46 | 3.95<br>t, J = 10.9 Hz | 3.18<br>d, *J* = 6.6 Hz | 3.75<br>3.72 |
| Gal | 4.36<br>d, J = 7.3 Hz | 3.65 | 3.61 | 3.86<br>d, J = 2.3 Hz | 3.36 | 3.48<br>3.41 |
| GlcNAc | 5.11<br>d, J = 7.9 Hz | 3.71 | 3.81 | 4.02<br>t, J = 8.5 Hz | 3.45 | 3.62 |
| Gal(2) | 4.36<br>d, J = 7.3 Hz | 3.71 | 3.71 | 3.88<br>d, *J* = 2.8 Hz | 3.40 | 3.40 |
| GlcNAc2 | 5.01 | 3.71 | 3.82 | 4.02<br>t, *J* = 8.5 Hz | 3.35 | 3.62 |
| Gal(3) | 4.36 | 3.70 | 3.74 | 3.90 | 3.39 | 3.26 |
| GlcA | 4.91 | 4.91 | 3.71 | 3.82 | 3.89 | - |

| **S10** | C1 | C2 | C3 | C4 | C5 | C6 |
|---|---|---|---|---|---|---|
| Glc | 105.0 – 106.8 | 80.2 | 84.7 | 75.0 | 79.3 | 68.3 |

| | | | | | | |
|---|---|---|---|---|---|---|
| Gal | 102.9 | 80.1 | 80.2 | 76.2 | 73.0 | 67.9 |
| GlcNAc | 99.9 | 57.8 | 80.1 | 75.9 | 75.4 | 67.8 |
| Gal(2) | 102.9 | 79.8 | 80.4 | 76.1 | 73.3 | 68.2 |
| GlcNAc2 | 100.3 | 57.8 | 78.2 | 75.9 | 75.2 | 67.8 |
| Gal(3) | 102.9 | 80.0 | 80.2 | 74.1 | 73.4 | 67.9 |
| GlcA | 101.3 | 73.8 | 75.4 | 77.7 | 74.1 | |

**3-*O*-sulfate-*β*-D-glucopyranosyluronate-(1→3)-*β*-D-galactopyranosyl-(1→4)-(2-acetamido-2-deoxy-*β*-D-glucopyranosyl-(1→3)-*β*-D-galactopyranosyl-(1→4)-(2-acetamido -2-deoxy-*β*-D-glucopyranosyl-(1→3)-*β*-D-galactopyranosyl-(1→4)-1-phenylthio-*β*-D-glucopyranoside (S11)**

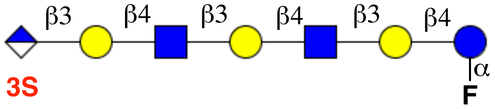


Compound **S11** (2 mg, 30%) was obtained from **S10** (15 mg, 0.005 mmol) using the general protocols for *O*-sulfation, hydrogenation and saponification.

ESI TOF-MS *m/z* $C_{46}H_{74}FN_2O_{39}S$ (M - H)$^-$ calculated 1330.13, found 1331.37.

$^1$H NMR (600 MHz, $CDCl_3$) δ 1.98, 1.95 (s, 3H, -NHCO$\underline{CH_3}$).

$^{13}$C NMR (151 MHz, $CDCl_3$): *δ* 22.8 (-NHCO$\underline{CH_3}$), 168.1 (-NH$\underline{C}$OCH$_3$).

| **S11** | H1 | H2 | H3 | H4 | H5 | H6 |
|---|---|---|---|---|---|---|
| Glc | 5.61<br>dd, $J_{HH}$ = 2.5 Hz<br>$J_{FH}$ = 53.6 Hz | 3.48 | 3.64 | 3.79 | 3.49 | 3.84<br>3.76 |
| Gal | 4.37<br>d, $J$ = 7.5 Hz | 3.60 | 3.65 | 4.07<br>d, $J$ = 3.3 Hz | 3.51 | 3.53 |

| | | | | | | |
|---|---|---|---|---|---|---|
| GlcNAc | 4.62<br>d, $J$ = 8.2 Hz | 3.71 | 3.74 | 3.69 | 3.60 | 3.87<br>3.76 |
| Gal(2) | 4.45<br>d, $J$ = 7.5 Hz | 3.59 | 3.65 | 4.07<br>d, $J$ = 3.3 Hz | 3.51 | 3.66 |
| GlcNAc2 | 4.62<br>d, $J$ = 8.2 Hz | 3.71 | 3.74 | 3.69 | 3.60 | 3.87<br>3.76 |
| Gal(3) | 4.45<br>d, $J$ = 7.5 Hz | 3.63 | 3.74 | 4.10<br>d, $J$ = 2.8 Hz | 3.61 | 3.66 |
| GlcA | 4.67 | 4.62 | 4.24<br>t, $J$ = 9.1 Hz | 3.64 | 3.87 | - |

| **S11** | C1 | C2 | C3 | C4 | C5 | C6 |
|---|---|---|---|---|---|---|
| Glc | 106.9 | 74.7 | 82.0 | 71.3 | 72.8 | 59.7 |
| Gal | 102.7 | 75.5 | na | 68.2 | 70.4 | 61.0 |
| GlcNAc | 102.7 | 55.1 | 82.4 | 72.8 | 72.6 | 61.0 |
| Gal(2) | 102.7 | 75.5 | na | 68.2 | 70.4 | 57.0 |
| GlcNAc | 102.7 | 55.1 | 82.4 | 72.8 | 72.6 | 61.0 |
| Gal(2) | 102.7 | 75.5 | 77.9 | 68.2 | 70.4 | 57.0 |
| GlcA | 103.4 | 75.0 | 82.0 | 72.2 | 72.4 | 171.4 |

***β-D*-glucopyranosyluronate-(1→3)-*β*-D-galactopyranosyl)-(1→4)-(2-acetamido-2-deoxy-*β*-D-glucopyranosyl-(1→3)-*β*-D-galactopyranosyl-(1→4)-1-phenylthio-*β*-D-glucopyranoside (S12)**

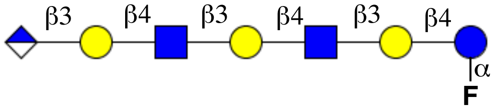


Compound **S12 (**1 mg, 25%) was obtained from **S13** (10 mg, 0.003 mmol) using the general protocols for hydrogenation and saponification. ESI TOF-MS *m/z* $C_{46}H_{74}FN_2O_{36}$ (M - H)$^-$ calculated 1250.08, found 1252.54.

$^1$H NMR (600 MHz, $CDCl_3$) δ 1.97, 1.94 (s, 3H, -NHCO*CH₃*).

$^{13}$C NMR (151 MHz, $CDCl_3$): *δ* 21.9 (-NHCO*CH₃*), 167.4 (-NH*C*OCH₃).

| **S12** | H1 | H2 | H3 | H4 | H5 | H6 |
|---|---|---|---|---|---|---|
| Glc | 5.56<br>dd, $J_{HH}$ = 2.4 Hz<br>$J_{FH}$ = 52.5 Hz | 3.43 | 3.58 | 3.74 | 3.40 | 3.79 |
| Gal | 4.32<br>d, $J$ = 7.1 Hz | 3.55 | 3.60 | 4.02 | 3.45 | 3.53 |
| GlcNAc | 4.56<br>d, $J$ = 8.2 Hz | 3.66 | 3.65 | 3.64 | 3.55 | 3.82 |
| Gal(2) | 4.40<br>d, $J$ = 7.7 Hz | 3.50 | 3.60 | 4.02 | 3.46 | 3.61 |
| GlcNAc2 | 4.57<br>d, $J$ = 7.9 Hz | 3.66 | 3.69 | 3.64 | 3.55 | 3.82 |
| Gal(3) | 4.40 | 3.57 | 3.69 | 4.01 | 3.56 | 3.66 |
| GlcA | 4.62 | 4.56 | 3.65 | 3.55 | 3.82 | - |

| **S11** | C1 | C2 | C3 | C4 | C5 | C6 |
|---|---|---|---|---|---|---|
| Glc | 105.7 | 73.5 | 80.7 | 70.0 | 71.6 | 58.5 |
| Gal | 101.5 | 73.8 | na | 66.6 | na | 59.5 |
| GlcNAc | 101.6 | 53.9 | 81.2 | 71.6 | na | 59.8 |
| Gal(2) | 102.5 | 74.3 | na | 66.9 | 69.2 | 55.9 |
| GlcNAc | 101.8 | 53.9 | 80.8 | 70.9 | 71.0 | 59.4 |
| Gal(2) | 102.1 | 74.5 | 76.8 | 67.0 | na | 55.8 |
| GlcA | 103.4 | 73.4 | 80.5 | 70.9 | 71.2 | 170.2 |

**(*2S,3R,4E*)-2-Amino-3-hydroxyoctadec-4-en-1-yl-3-O-sulfo-*β-D*-glucopyranosyl-(1→3)-*β*-D-galactopyranosyl)-(1→4)-(2-acetamido-2-deoxy-*β*-D-glucopyranosyl-(1→3)-*β*-D-galactopyranosyl-(1→4)-*β*-D-glucopyranoside (S13)**

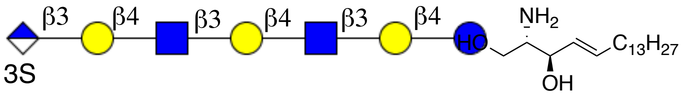


Compound **S12** (1 mg, 0.0008 mmol) and *erythro*-sphingosine (0.4 mg, 0.0012 mmol) are condensed using general protocol for sphingosine installation to afford 0.7 mg (55%) of lyso-sulfoglucuronyl paragloboside (**S13**).
ESI TOF-MS *m/z* $C_{64}H_{110}N_3O_{41}S$ $(M - H)^-$ calculated 1609.69, found 1610.42.

$^1$H NMR (600 MHz, $D_2O$) δ: (non-carbohydrate): 5.76 (m, 1H, =CH (sphingosine)), 5.40 (dt, $J_1$ = 6.4 Hz, $J_2$ = 15.3 Hz, 1H, HC=C (sphingosine)), 4.17 (m, 1H, -C*H*NH$_2$ (sphingosine)), 3.70 (m, 2H, -OC*H*$_2$- C1-sphingosine), 3.54 (m, 2H, =CHC*H*$_2$-), 1.89 (m, 2H, -C*H*$_2$), 1.88 (s, 3H, -NHAc), 1.32 (m, 4H, -CH$_2$), 1.28 – 1.12 (m, 44H, -CH$_2$), 0.83 (t, J = 6.1 Hz, 3H, CH$_3$).

$^{13}$C NMR (151 MHz, $D_2O$) δ: (non-carbohydrate): 136.1 (=*C*H (sphingosine)), 128.4 (H*C*=C (sphingosine)), 70.1 (-*C*HNH$_2$ (sphingosine)), 58.1 (-O*C*H$_2$- C1-sphingosine), 58.1 (=CHC*H*$_2$-), 32.1 (-*C*H$_2$), 32.0 (-CH$_2$), 30.1 – 27.7 (-CH$_2$), 21.7 (-NHAc), 12.3 (CH$_3$).

| **S13** | H1 | H2 | H3 | H4 | H5 | H6 |
|---|---|---|---|---|---|---|
| Glc | 4.32 | 3.47 | 3.67 | 3.76 | na | 3.88<br>3.77 |
| Gal | 4.37 | 3.62 | Na | 3.95<br>d, *J* = 3.2 Hz | na | 3.50 |
| GlcNAc | 4.65 | 3.71 | 3.76 | na | 3.55 | 3.78 |
| Gal(2) | 4.46 | 3.52 | na | 4.01 | na | 3.56 |
| GlcNAc2 | 4.67 | 3.71 | 3.76 | na | 3.55 | 3.78 |
| Gal(3) | 4.39 | 3.52 | na | 4.01 | na | 3.56 |
| GlcA | 4.67 | 3.60 | 4.18 | 3.69 | 3.86 | - |

**(*2S,3R,4E*)-2-hexadecanamido-3-hydroxyoctadec-4-en-1-yl-3--O-sulfo-*β*-D-galactopyranosyl-(1→4)-2-acetamido-2-deoxy-*β*-D-glucopyranosyl-(1→3)-*β*-D-galactopyranosyl-(1→4)-2-acetamido-2-deoxy-*β*-D-glucopyranosyl-(1→3)-*β*-D-galactopyranosyl-(1→4)-*β*-D-glucopyranoside (1b)**

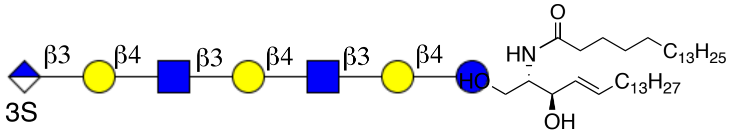


Compound **S13** (0.6 mg, 0.0036 mmol) was acylated using general protocol for stearic acid installation to afford compound **1b** (0.4 mg, 57%). ESI TOF-MS *m/z* $C_{82}H_{142}N_3O_{42}S$ $(M - H)^-$ calculated 1874.08, found 1872.46.

$^1$H NMR (600 MHz, $D_2O$) δ: (non-carbohydrate): 5.58 (m, 1H, =CH (sphingosine)), 5.36 (dd, $J_1$ = 7.9 Hz, $J_2$ = 15.3 Hz, 1H, HC=C (sphingosine)), 4.10 (d, $J_1$ = 4.6 Hz, $J_2$ = 10.4 Hz, 1H, -OCH*H*-C1-sphingosine), 3.98 (t, J = 8.1 Hz, 1H, -C*H*$NH_2$(sphingosine)), 3.51 (m, 1H, -OC*H*H- C1-sphingosine), 2.06 (t, J = 7.6 Hz, 2H, α-$CH_2$-stearic acid), 1.93 (m, 2H, -C$H_2$), 1.89 (s, 6H, -NHAc), 1.47 (m, 4H, -$CH_2$), 1.26 – 1.16 (m, 67H, -$CH_2$), 0.81 (t, J = 6.1 Hz, 3H, $CH_3$).

$^{13}$C NMR (151 MHz, $D_2O$) δ: (non-carbohydrate): 133.9 (=*C*H (sphingosine)), 130.1 (H*C*=C (sphingosine)), 74.0 (-*C*H$NH_2$ (sphingosine)), 68.5 (-O*C*$H_2$- C1-sphingosine), 58.6 (=CHC$H_2$-), 35.4 (α-*C*$H_2$-stearic acid -), 33.5 (-$CH_2$), 30.1 – 28.3 (-$CH_2$), 26.2 (-$CH_2$), 22.6 (-$CH_2$), 21.5 (-NHAc), 21.3 (-NHAc), 13.0 ($CH_3$).

Carbohydrate region:

| **1b** | H1 | H2 | H3 | H4 | H5 | H6 |
|---|---|---|---|---|---|---|
| Glc | 4.20<br>d, *J* = 8.1 Hz | 3.20 | 3.66 | 3.74 | na | na |
| Gal | 4.24<br>d, *J* = 8.3 Hz | 3.60 | 3.64 | 3.95 | na | na |
| GlcNAc | 4.64<br>d, *J* = 8.1 Hz | 3.68 | 3.76 | 3.75 | na | na |
| Gal2 | 4.35 | 3.51 | 3.62 | 4.01 | na | na |
| GlcNAc2 | 4.66 | 3.68 | 3.75 | 3.67 | na | na |
| Gal3 | 4.40 | na | na | 4.06 | na | na |
| GlcA | 4.58<br>d, *J* = 7.6 Hz | 3.48 | 4.20 | 3.68 | 3.86 | |

| **1b** | C1 | C2 | C3 | C4 | C5 | C6 |
|---|---|---|---|---|---|---|
| Glc | 103.2 | 73.0 | 74.1 | 76.4 | na | na |
| Gal | 103.4 | 74.1 | 82.1 | 68.5 | na | na |
| GlcNAc | 102.8 | 55.4 | 78.9 | 76.1 | na | na |
| Gal2 | 103.4 | 76.7 | 82.5 | 68.1 | na | na |
| GlcNAc2 | 102.7 | 55.1 | 78.0 | 76.0 | na | na |
| Gal3 | 103.6 | na | na | 67.8 | na | na |
| GlcA | 103.5 | 79.1 | 82.5 | 71.5 | 72.6 | 170.3 |

**(*2S,3R,4E*)-2-hexadecanamido-3-hydroxyoctadec-4-en-1-yl-*β*-D-galactopyranosyl-(1→4)-2-acetamido-2-deoxy-*β*-D-glucopyranosyl-(1→3)-*β*-D-galactopyranosyl-(1→4)-2-acetamido-2-deoxy-*β*-D-glucopyranosyl-(1→3)-*β*-D-galactopyranosyl-(1→4)-*β*-D-glucopyranoside (2b)**

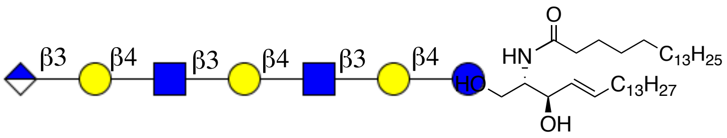


Compound **S14** (0.6 mg, 0.0005 mmol) was acylated using general protocol for stearic acid installation to afford compound **2a** (0.5 mg, 55%). ESI-TOF MS *m/z* $C_{82}H_{143}N_3O_{39}$ $(M-H)^-$ calculated 1794.02, found 1793.53.

$^1H$ NMR (600 MHz, $D_2O$) δ: (non-carbohydrate): 5.56 (m, 1H, =CH (sphingosine)), 5.37 (dd, $J_1$ = 8.6 Hz, $J_2$ = 15.7 Hz, 1H, HC=C (sphingosine)), 4.09 (d, $J_1$ = 4.2 Hz, $J_2$ = 10.4 Hz, 1H, -OCH*H*-C1-sphingosine), 3.95 (t, J = 8.1 Hz, 1H, -C*H*$NH_2$(sphingosine)), 3.45 (m, 1H, -OC*H*H- C1-sphingosine), 2.08 (t, J = 7.6 Hz, 2H, α-$CH_2$-stearic acid), 1.93 (m, 2H, -C*$H_2$*), 1.91 (s, 6H, -NHAc), 1.46 (m, 4H, -$CH_2$), 1.26 – 1.11 (m, 73H, -$CH_2$), 0.81 (t, J = 6.2 Hz, 3H, $CH_3$).

$^{13}C$ NMR (151 MHz, $D_2O$) δ: (non-carbohydrate): 133.1 (=*C*H (sphingosine)), 131.0 (H*C*=C (sphingosine)), 71.0 (-*C*H$NH_2$ (sphingosine)), 68.3 (-O*C*$H_2$- C1-sphingosine), 58.4 (=CHC*$H_2$*-), 35.1 (α-*C*$H_2$-stearic acid -), 33.6 (-$CH_2$), 31.3 – 29.0 (-$CH_2$), 24.3 (-$CH_2$), 21.4 (-$CH_2$), 21.2 (-NHAc), 20.5 (-NHAc), 11.0 ($CH_3$).

| **2a** | H1 | H2 | H3 | H4 | H5 | H6 |
|---|---|---|---|---|---|---|
| Glc | 4.21 | 3.18 | 3.68 | na | na | 3.80 |
| Gal | 4.26 | 3.52 | na | 3.90 | na | 3.64 |

| | | | | | | |
|---|---|---|---|---|---|---|
| GlcNAc | 4.56<br>d, $J$ = 7.7 Hz | 3.71 | na | 3.75 | na | 3.68 |
| Gal(2) | 4.34 | 3.51 | na | 4.01<br>d, $J$ = 2.6 Hz | na | 3.64 |
| GlcNAc2 | 4.55<br>d, $J$ = 7.7 Hz | 3.70 | na | 3.75 | na | 3.68 |
| Gal(3) | 4.36 | 3.50 | 3.66 | 4.10<br>d, $J$ = 2.6 Hz | na | 3.64 |
| GlcA | 4.52 | 3.20 | 3.22 | 3.68 | 3.72 | - |

| **2a** | C1 | C2 | C3 | C4 | C5 | C6 |
|---|---|---|---|---|---|---|
| Glc | 100.5 | 72.4 | 80.1 | na | Na | 60.4 |
| Gal | 101.3 | 73.5 | Na | 68.5 | na | 61.0 |
| GlcNAc | 102.0 | 56.2 | na | na | na | 60.6 |
| Gal(2) | 103.0 | 74.1 | na | na | na | 61.0 |
| GlcNAc | 102.0 | 56.2 | na | na | na | 60.6 |
| Gal(2) | 103.0 | 74.1 | na | na | na | 61.2 |
| GlcA | 101.6 | 79.0 | 79.6 | 72.0 | 74.2 | 170.0 |

### 8. Glycoremodeling and hemagglutination of red-blood cells and cell-based assays

Fresh fowl blood was centrifuged (10 min, 1000 rpm) followed by removal of the supernatant. Cell pellets were washed three times in sterile PBS (pH 7.4, Gibko$^{TM}$) with intermittent centrifugation (1000 rpm, 10 min). Erythrocyte solutions were stored in a 25% solution in Alsever's solution until further use.

#### General protocol for glycolipid remodeling of red blood cells

Equal volumes of red-blood cells (25%) and glycosphingolipids (1 mg/mL) were mixed and incubated at 37 $^{o}$C for 1 h. After incubation the cell suspension was centrifuged (10 min, 1000 rpm) and the supernatant was discarded. Cell pellets were resuspended in PBS and centrifuged again at 1000 rpm for 10 min. The washing procedure was performed three times to ensure the removal of leftover glycolipids and the final cell-suspension in PBS was directly used for hemagglutination and hemolysis assays or buffer exchanged to Alsever's solution for further storage.

**General protocol for antibody-agglutination of SGPG (1a) and GPG (2a) modified erythrocytes**

Glycolipid remodeled red-blood cells were diluted in PBS to 1% final cell-suspension. To 50 µL of RBC suspension in 96-well V-shaped microtiter plate (Nunc™ 249662), 50 µL of monoclonal mouse anti-CD57 IgM antibody (ANT-288, ProspecBio) was added with 10-fold serial dilutions (100 µM to 0.1 nM) and incubated at RT for 30 min. The button formations were observed with naked eye, which were verified by plate tilting. All assays were performed in full biological replicates.

Red-blood cells that were glycoremodeled under different conditions were tested for hemagglutination to assess the efficiency of antigen incorporation:

- **Prolonged (6 h)** incubation of erythrocytes with glycosphingolipids (1 mg/mL)
- Incubation of erythrocytes with glycosphingolipids (1 mg/mL) at lower (**22 ºC**) temperatures
- Incubation of erythrocytes with reduced concentration of glycosphingolipids (**0.1 mg/mL**)
- Incubation of erythrocytes with glycosphingolipids (1 mg/mL), followed by **0.1% FCS** wash
- Incubation of erythrocytes with glycosphingolipids (1 mg/mL), followed by ***trypsin*** treatment
- Erythrocytes **trypsinization**, followed by incubation with glycosphingolipids (1 mg/mL).

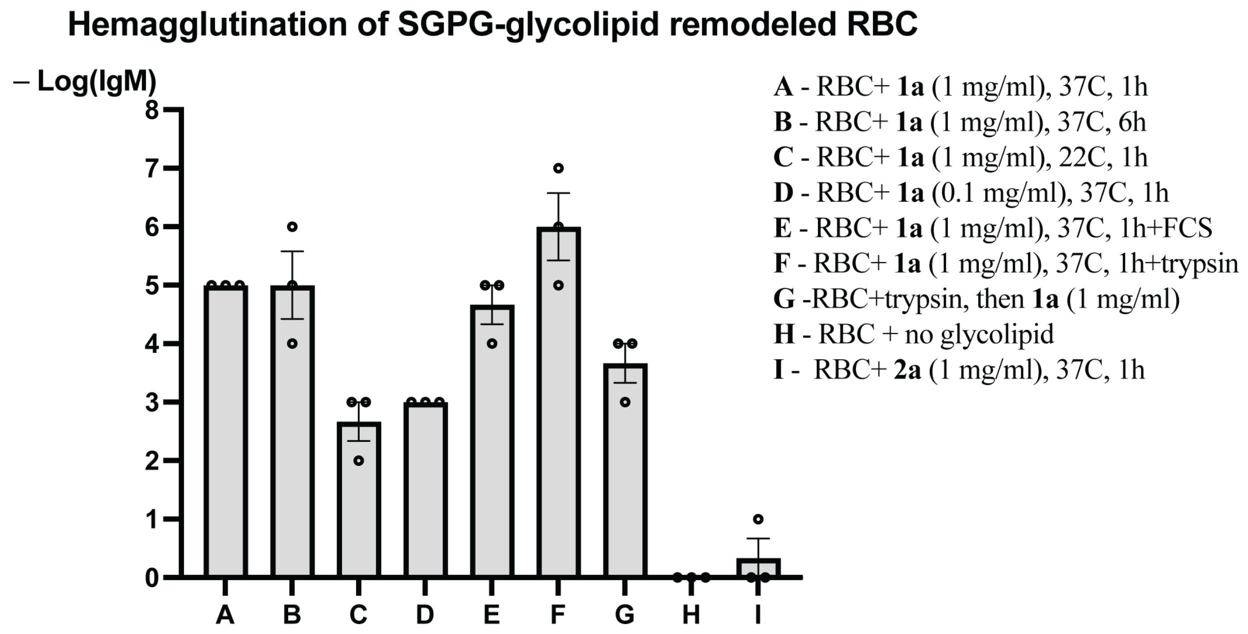


**Figure S6**. Hemagglutination of SGPG-remodeled erythrocytes with monoclonal anti-CD57 IgM antibody.

**Protocol for antibody-agglutination of GM1a and GD1a remodeled erythrocytes**

General protocol for glycolipid remodeling and hemagglutination is used for ganglioside incorporated erythrocytes. Briefly, red-blood cells were remodeled with GM1a (1 mg/mL) and GD1a (1 mg/mL) gangliosides. Monoclonal rabbit anti-GM1a IgG1 antibody (GTX50971, GeneTex) and monoclonal mouse anti-GD1a IgG1 antibody (Sigma Aldrich, MAB5606Z) were used as primary antibodies for each ganglioside. Polyclonal goat anti-rabbit IgG (Sigma Aldrich, B8895) and goat anti-mouse IgG (Sigma Aldrich, B7264) were used as secondary antibodies against anti-GM1a IgG and anti-GD1a IgG respectively.

To 50 µL of RBC suspension in 96-well V-shaped microtiter plate (Nunc™ 249662), 50 µL of anti-GM1a or anti-GD1a antibody was added with 10-fold serial dilutions (100 µM to 0.1 nM) and incubated at RT for 30 min. After which cell-suspensions were pipetted up-and-down, followed by the addition of secondary antibodies and incubated for 30 min. The button formations were observed with naked eye, which were verified by plate tilting. All assays were performed in full biological replicates.

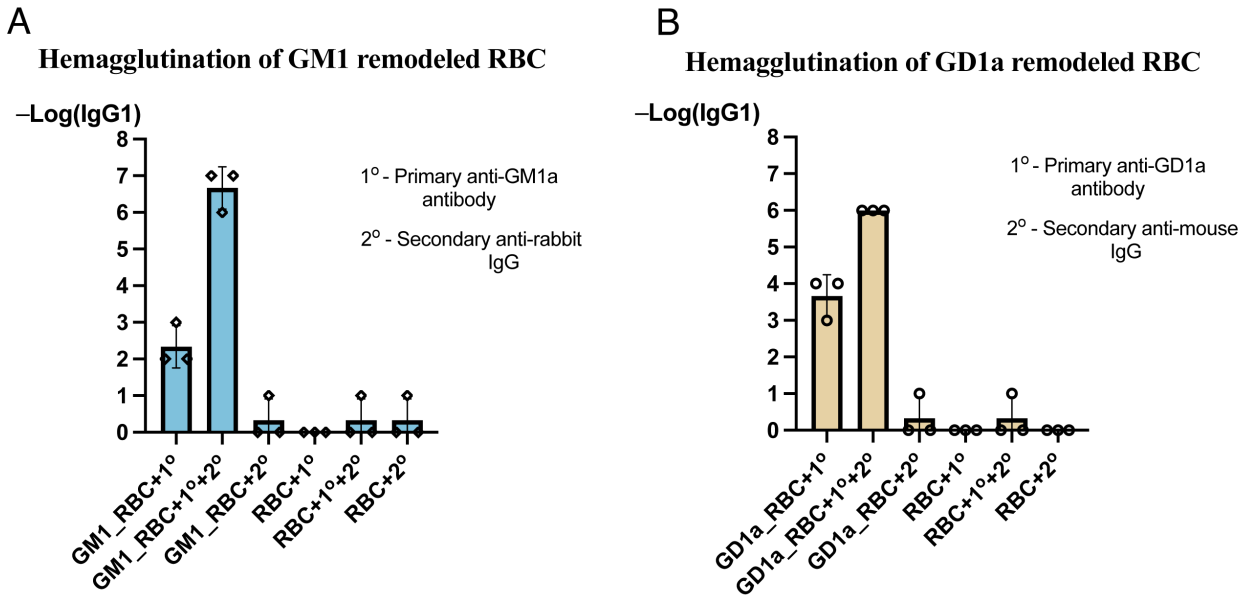


**Figure S7.** Hemagglutination of ganglioside-remodeled erythrocytes: (A) GM1 remodeled red-blood cells show slight agglutination with anti-GM1a IgG antibody, which increases upon the addition of secondary anti-IgG antibody. (B) GD1a remodeled erythrocytes are agglutinated with anti-GD1a IgG antibody and the presence of secondary anti-IgG antibody markedly increase the agglutination of erythrocytes. None of the ganglioside remodeled red-blood cells shows any degree of agglutination in the presence of respective secondary only anti-IgG antibodies.

### Protocol for antibody-mediated complement dependent cytotoxicity

50 µL of 1% suspension of glycan remodeled erythrocytes in Dextrose-Gelatin-Hepes buffer were seeded in 96-well V-shaped microtiter plate (Nunc™ 249662) , to which anti-glycolipid antibodies were added, followed by the addition of 1:4 diluted guinea-pig serum complement as an exogenous source of complement. The cell suspension was incubated at 37 °C for 2 h, after which it was centrifuged for 5 min at 1000 rcf. Supernatants were transferred to 96-well flat bottom microtiter plate (Nunc™ 243656) and absorbance at 415 nm was measured using plate reader. Relative hemolysis was calculated using the formula: %RH = $(A_{\text{sample}} - A_{\text{blank}})/\ (A_{\text{lysis buffer}} - A_{\text{blank}})\text{x}100$, where $A_{\text{sample}}$ – represents the absorbance of released hemoglobin from the glyolipid remodeled erythrocytes, $A_{\text{lysis buffer}}$ – represents the absorbance of released hemoglobin from the native erythrocytes treated with ACK-lysis buffer, $A_{\text{blank buffer}}$ – represents the absorbance of spontaneously released hemoglobin from non-treated erythrocytes.

### 10. Copies of NMR spectra

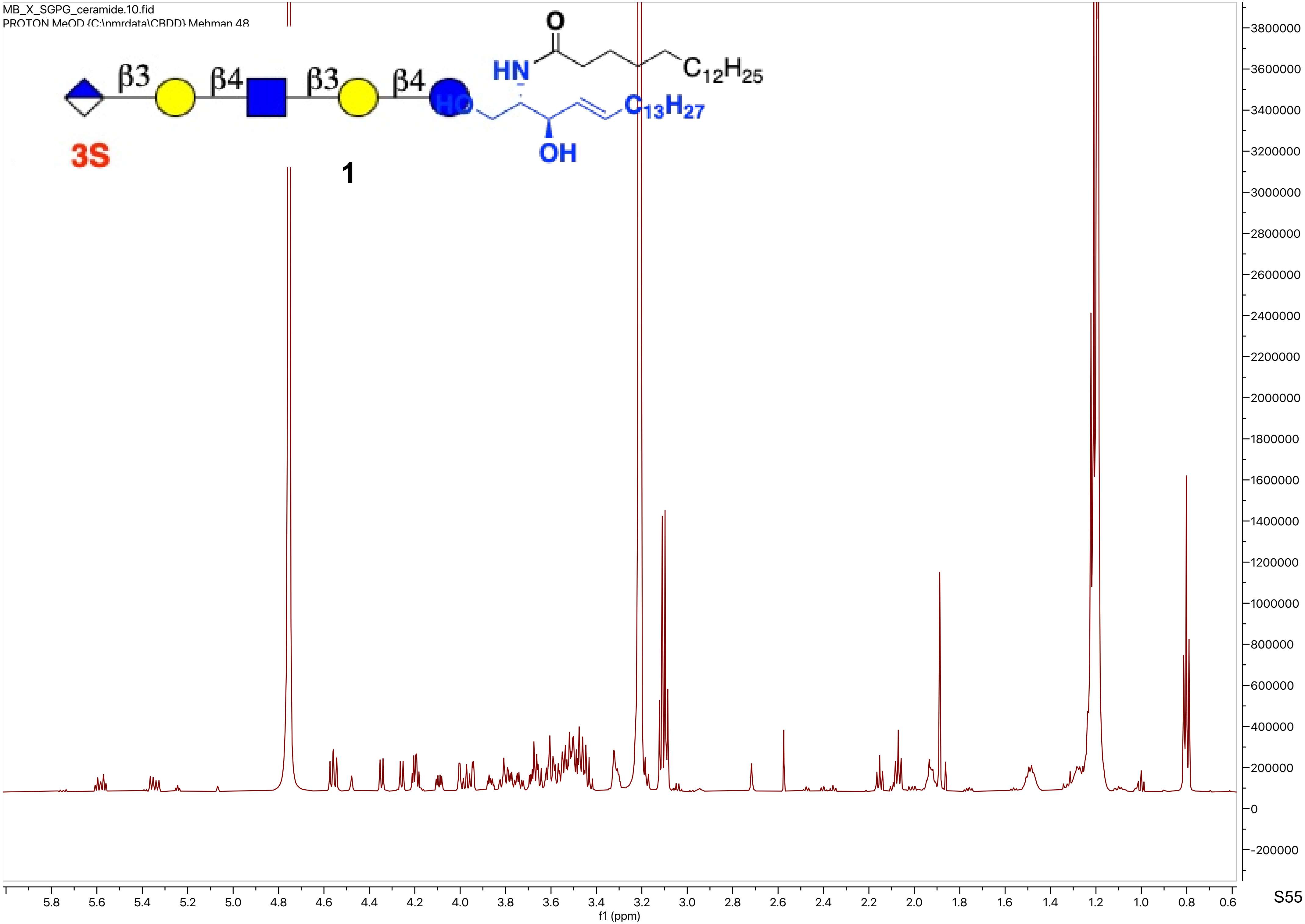
MB_X_SGPG_ceramide.10.fid
PROTON MeOD {C:\nmrdata\CBDD} Mehman 48
β3
β4
β3
β4
3S
1
HN
O
$C_{12}H_{25}$
HO
$C_{13}H_{27}$
OH
f1 (ppm)

MB_X_SGPG_ceramide.10.fid
PROTON MeOD {C:\nmrdata\CBDD} Mehman 48

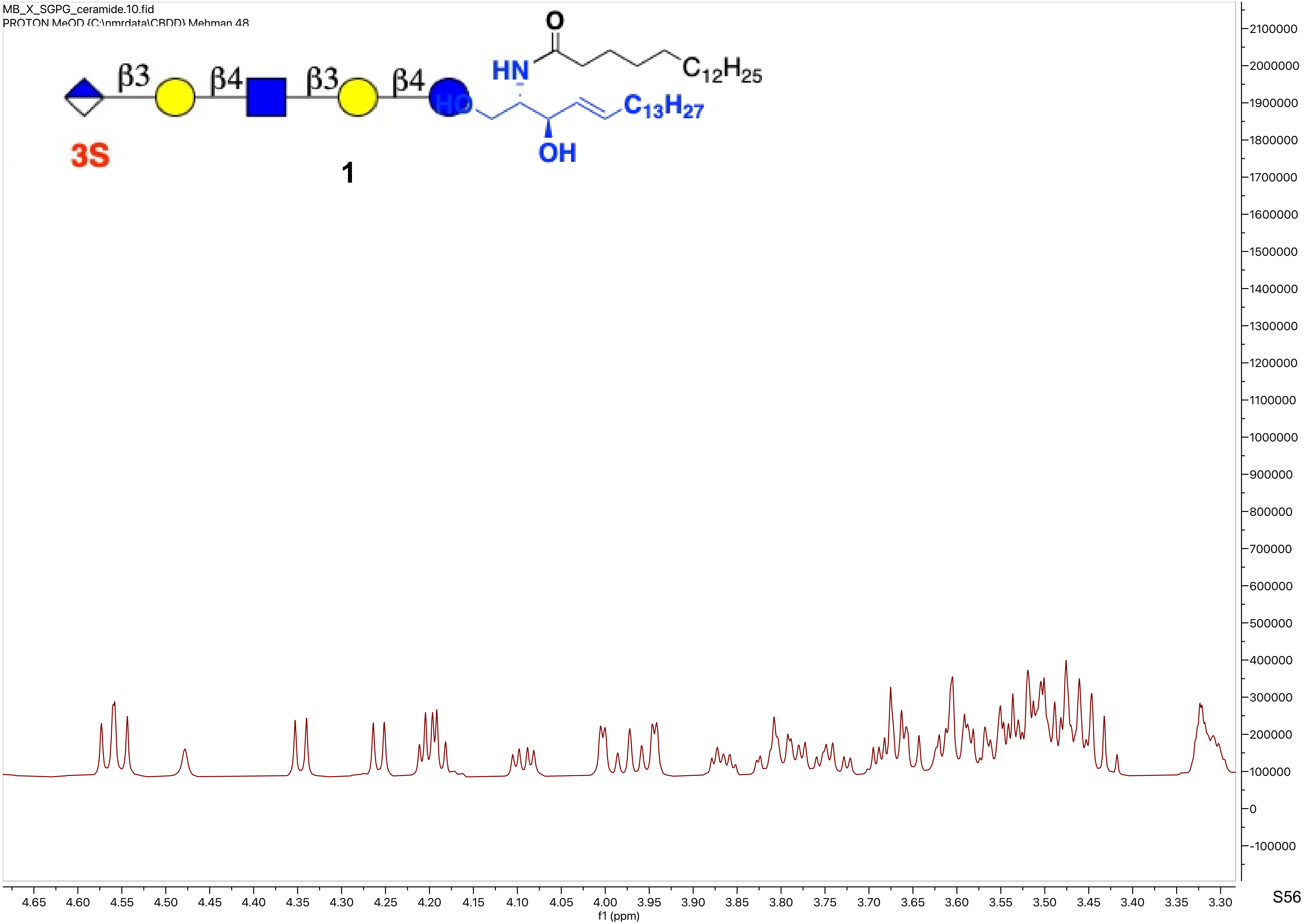

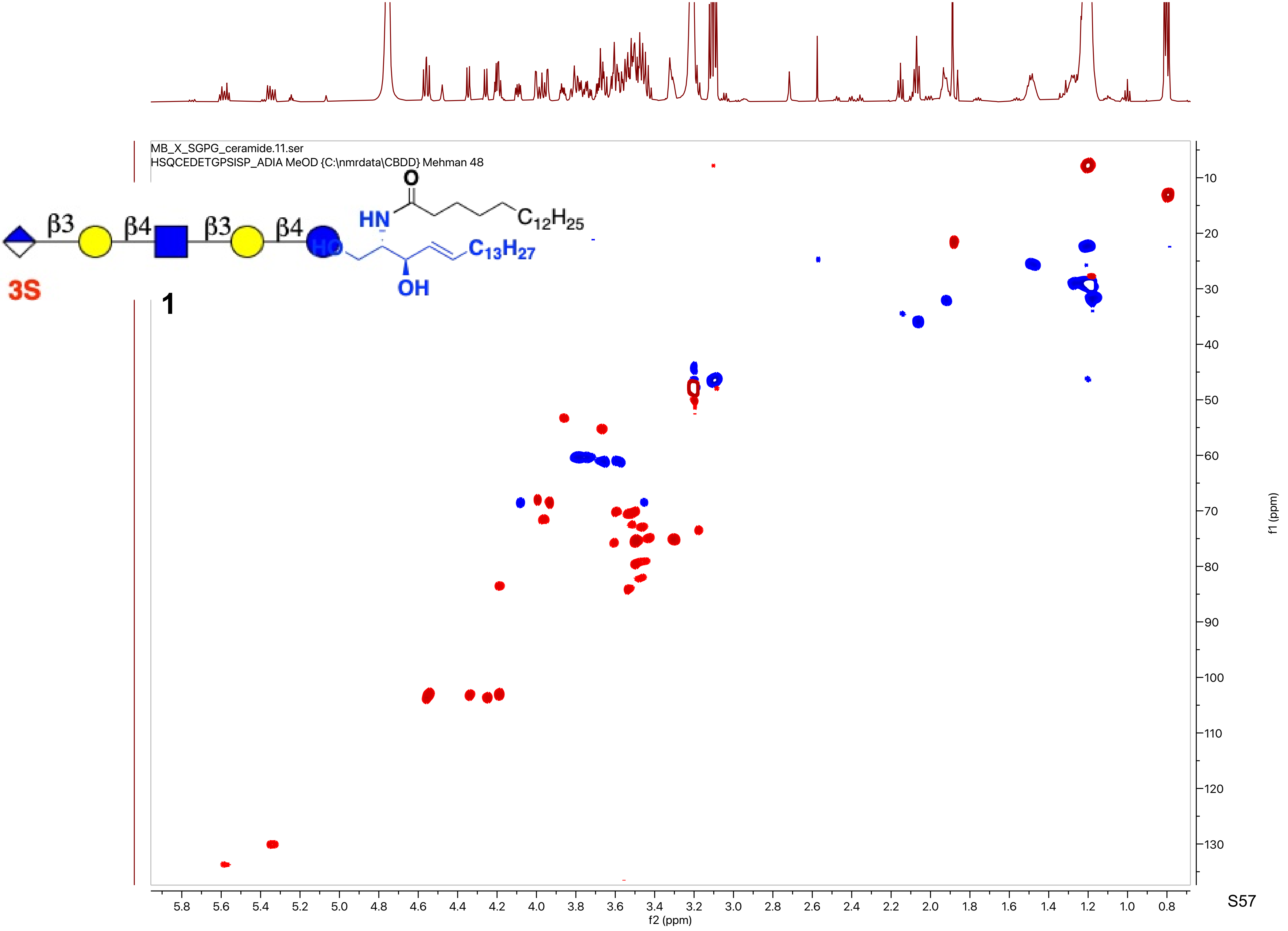
MB_X_SGPG_ceramide.11.ser
HSQCEDETGPSISP_ADIA MeOD {C:\nmrdata\CBDD} Mehman 48
β3
β4
β3
β4
3S
O
HN
$C_{12}H_{25}$
HO
$C_{13}H_{27}$
OH
1
f2 (ppm)
f1 (ppm)

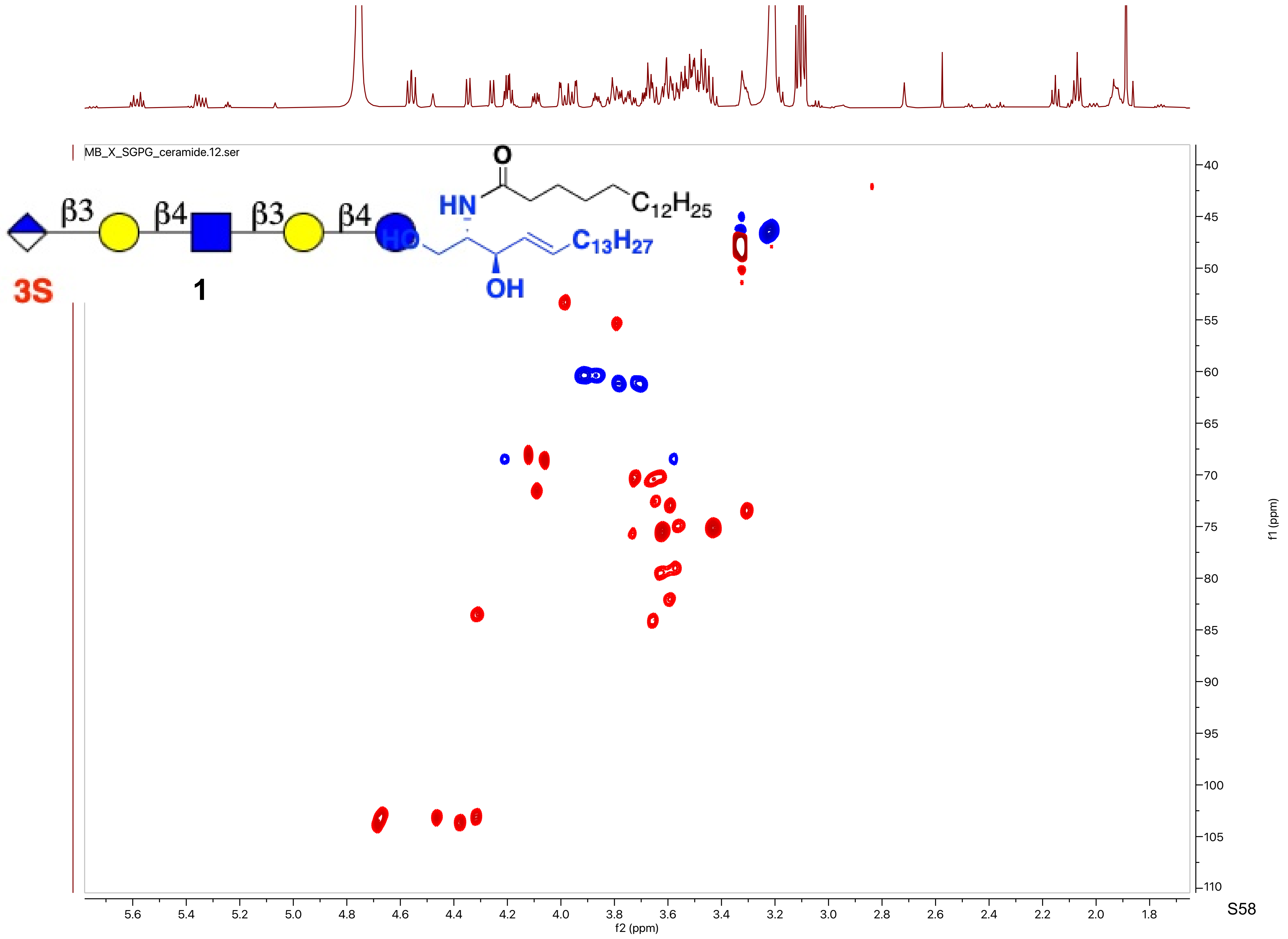

MB_X_SGPG_ceramide.12.ser
β3
β4
β3
β4
3S
1
O
HN
HO
OH
$C_{12}H_{25}$
$C_{13}H_{27}$
f2 (ppm)
f1 (ppm)

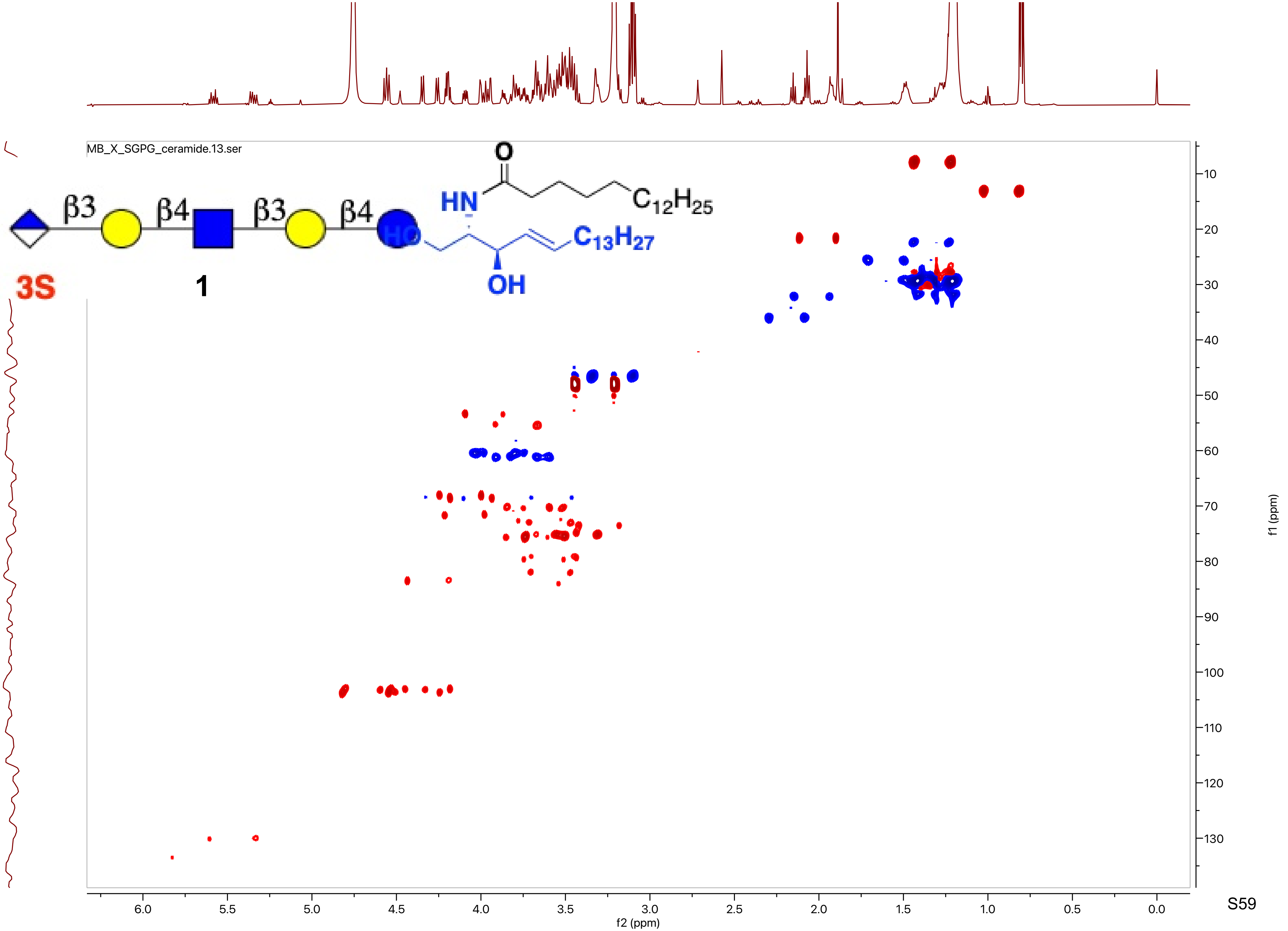
MB_X_SGPG_ceramide.13.ser
β3
β4
β3
β4
3S
1
O
HN
$C_{12}H_{25}$
HO
$C_{13}H_{27}$
OH
f2 (ppm)
f1 (ppm)

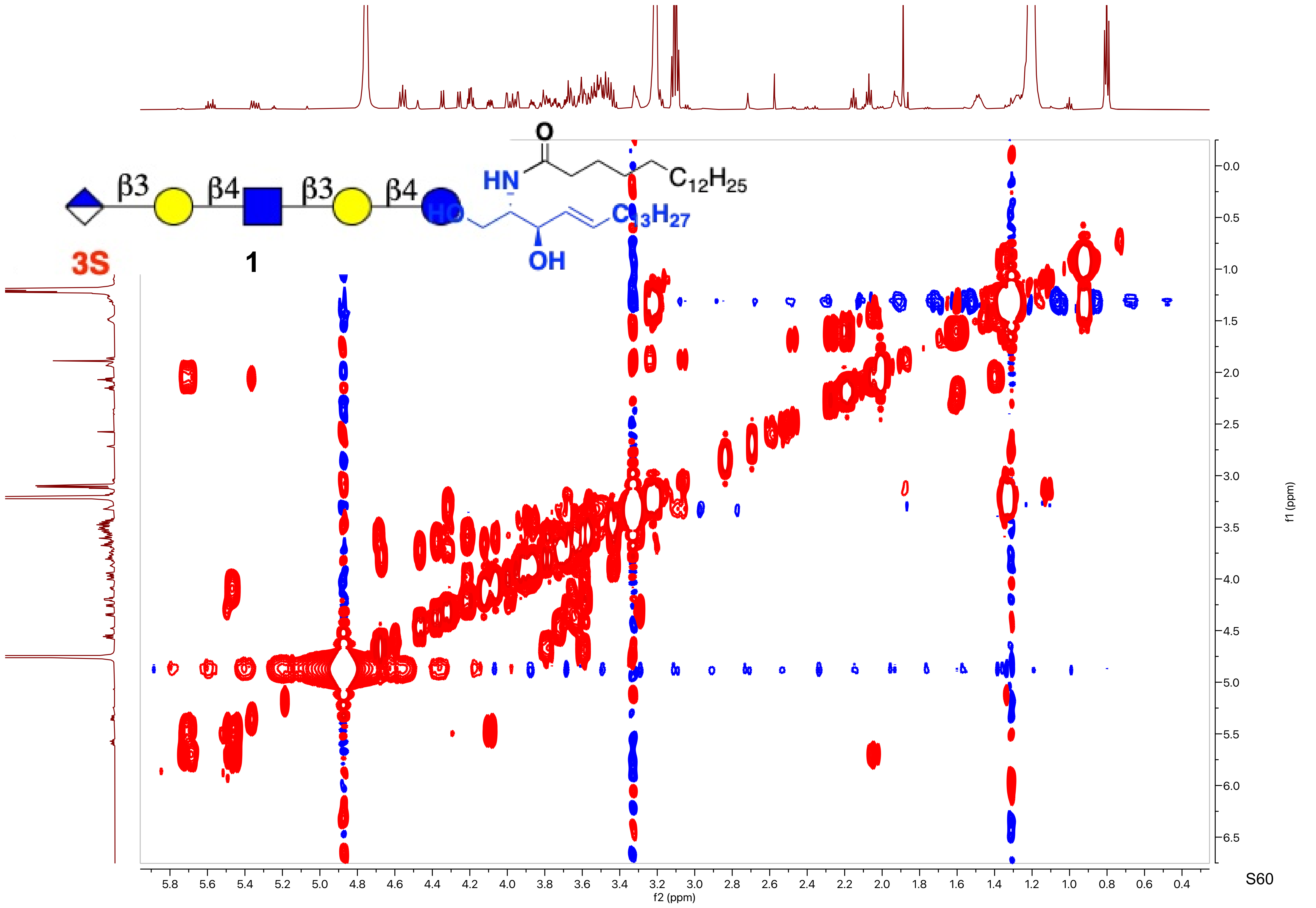

β3
β4
β3
β4
3S
1
O
HN
$C_{12}H_{25}$
HO
$C_{13}H_{27}$
OH
f1 (ppm)
f2 (ppm)

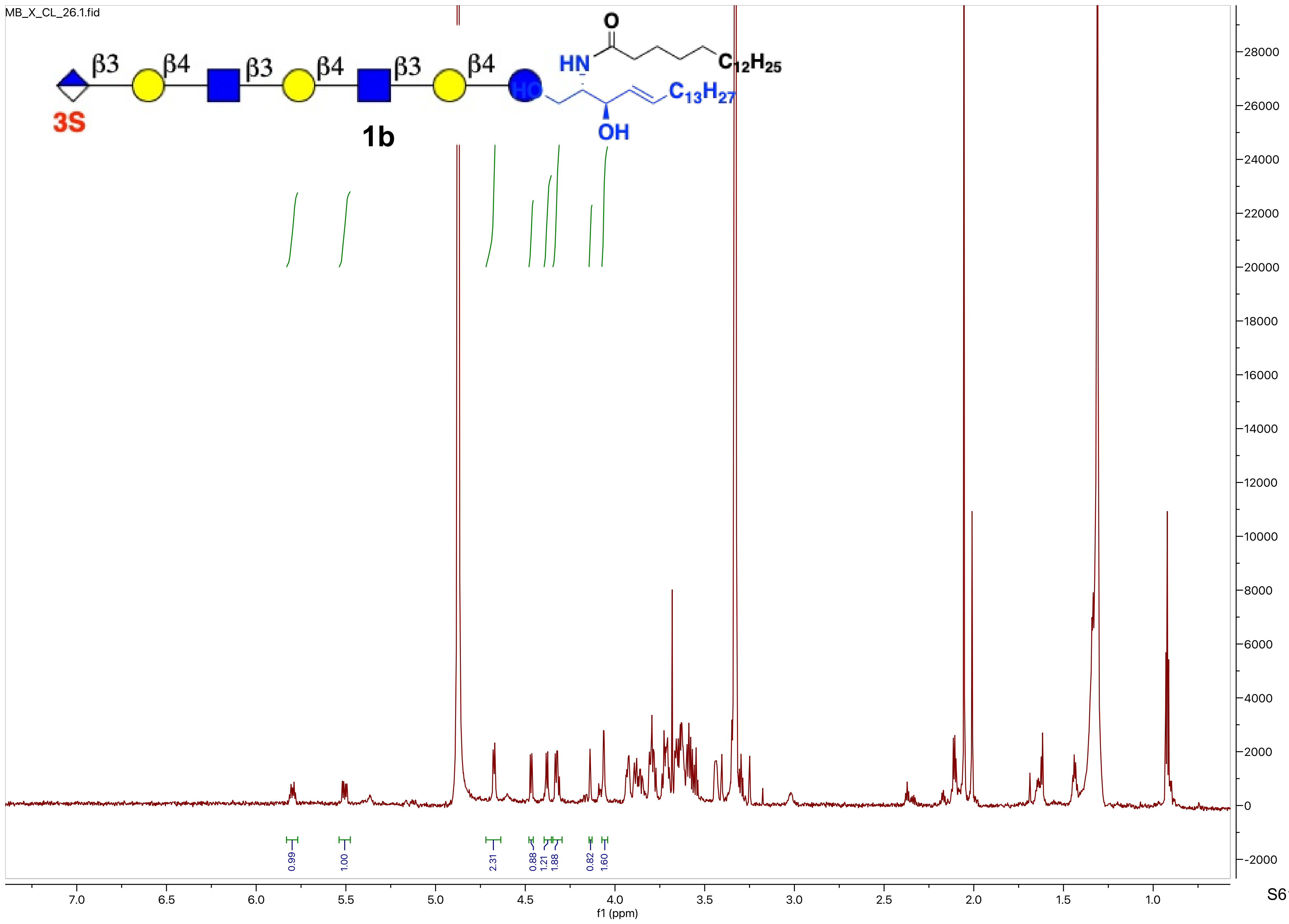
MB_X_CL_26.1.fid
β3
β4
β3
β4
β3
β4
3S
1b
O
HN
$C_{12}H_{25}$
HO
$C_{13}H_{27}$
OH
0.99
1.00
2.31
0.88
1.21
1.88
0.82
1.60
7.0
6.5
6.0
5.5
5.0
4.5
4.0
3.5
3.0
2.5
2.0
1.5
1.0
f1 (ppm)
28000
26000
24000
22000
20000
18000
16000
14000
12000
10000
8000
6000
4000
2000
0
-2000

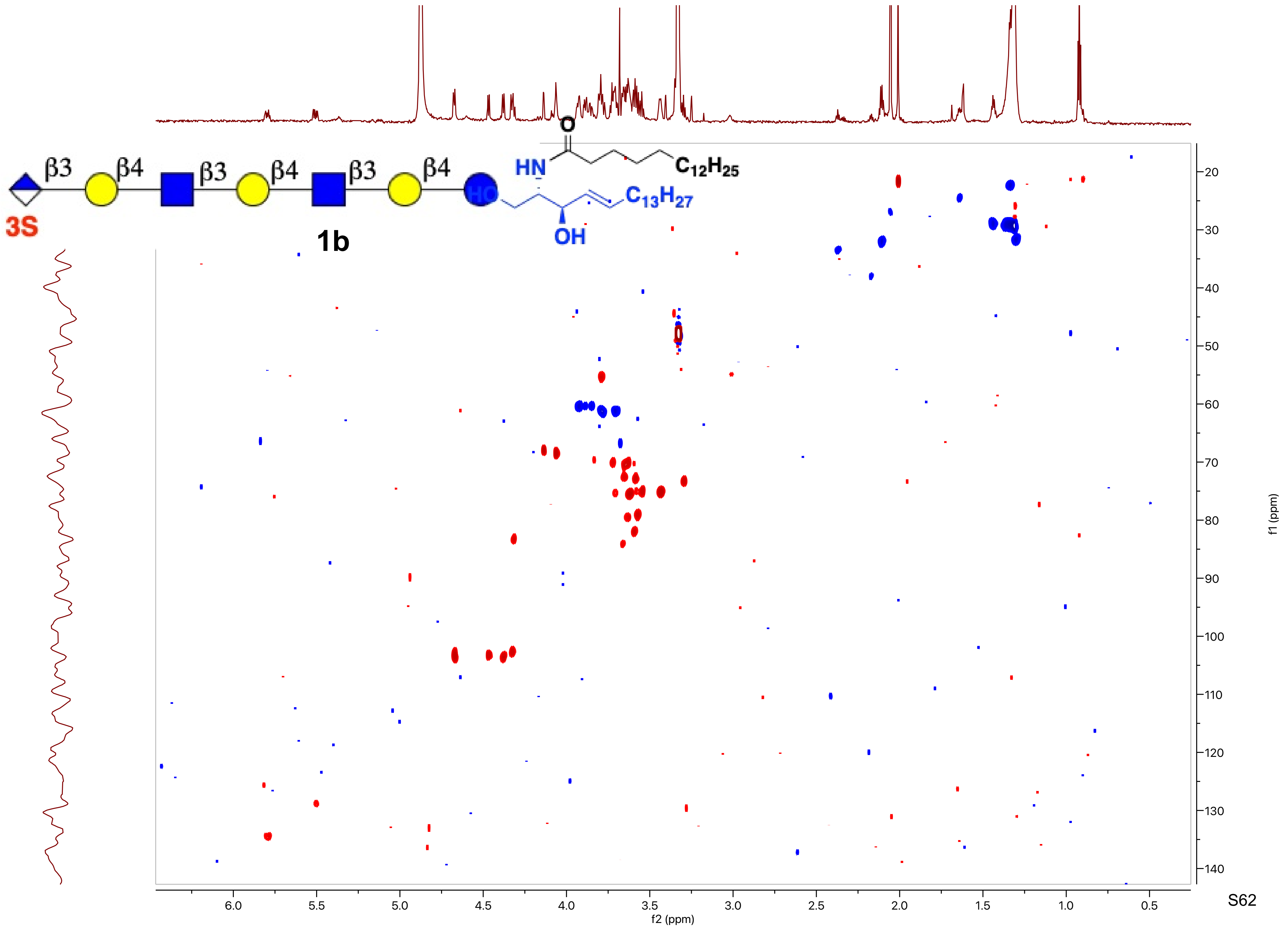

β3
β4
β3
β4
β3
β4
3S
1b
O
HN
HO
OH
$C_{12}H_{25}$
$C_{13}H_{27}$
f2 (ppm)
f1 (ppm)

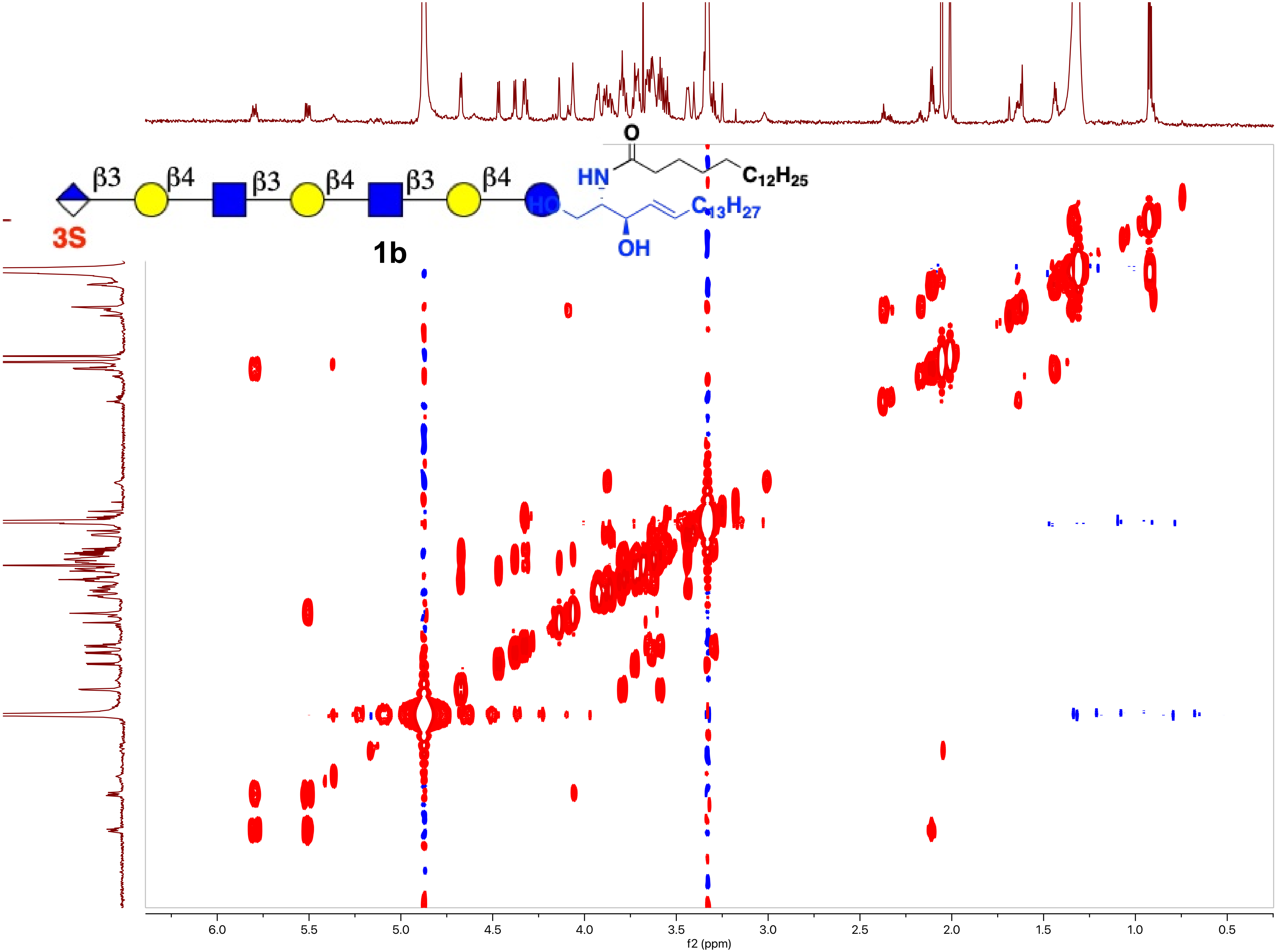
β3
β4
β3
β4
β3
β4
3S
HN
O
$C_{12}H_{25}$
HO
OH
$C_{13}H_{27}$
1b
f2 (ppm)
f1 (ppm)

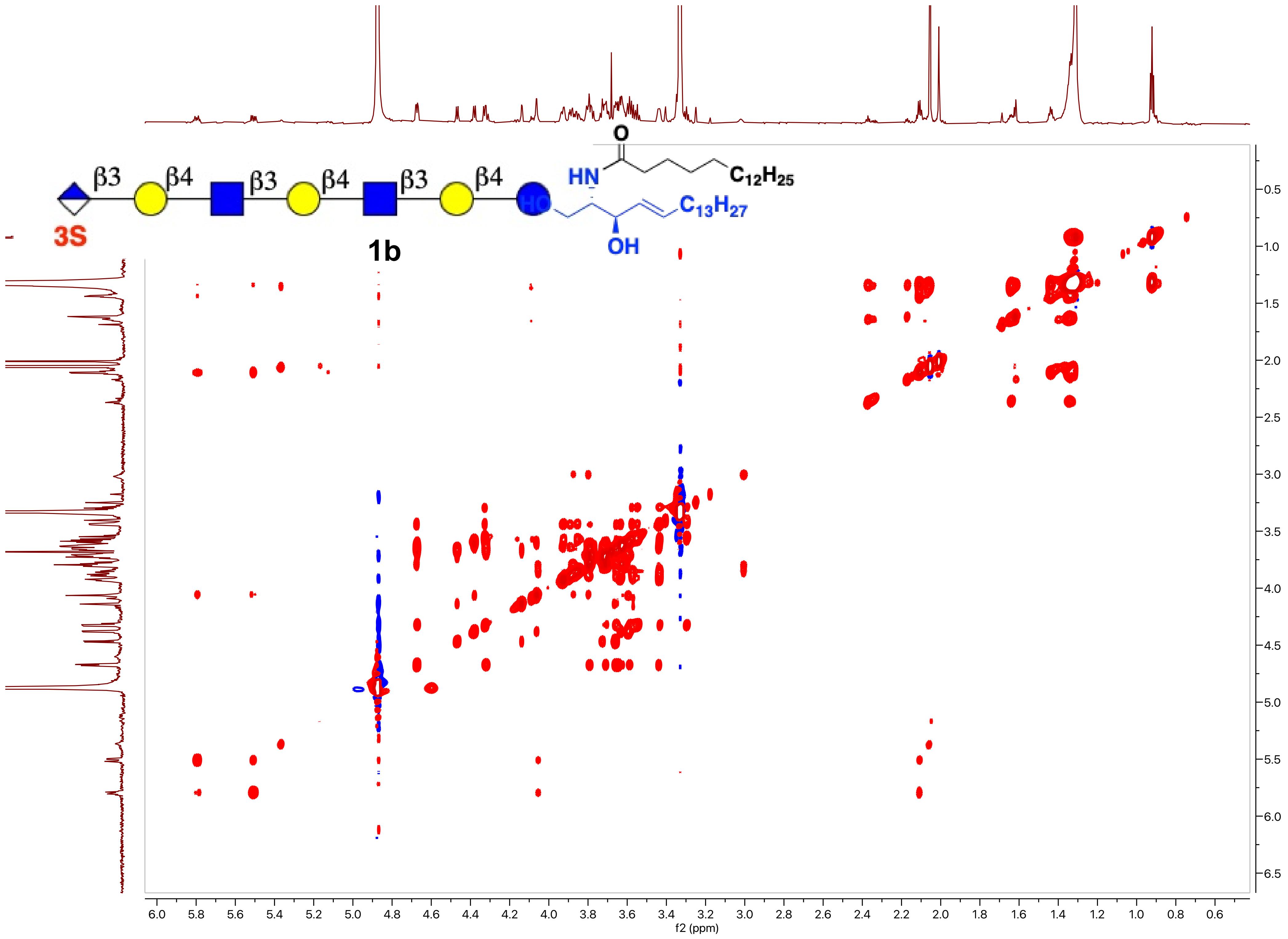
β3
β4
β3
β4
β3
β4
3S
1b
HN
O
$C_{12}H_{25}$
HO
OH
$C_{13}H_{27}$
f1 (ppm)
f2 (ppm)

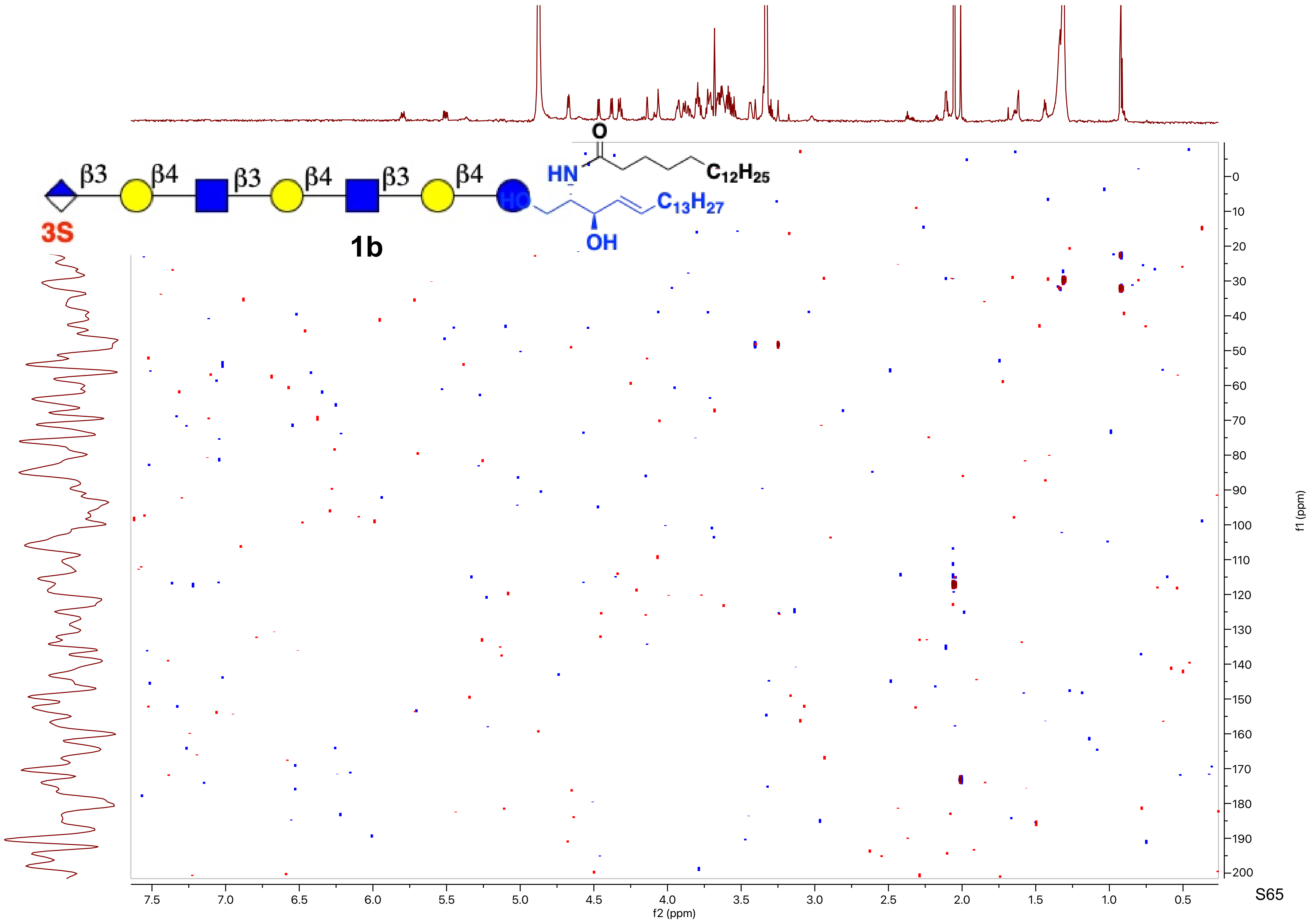
β3
β4
β3
β4
β3
β4
3S
1b
O
HN
C12H25
HO
C13H27
OH
f1 (ppm)
f2 (ppm)

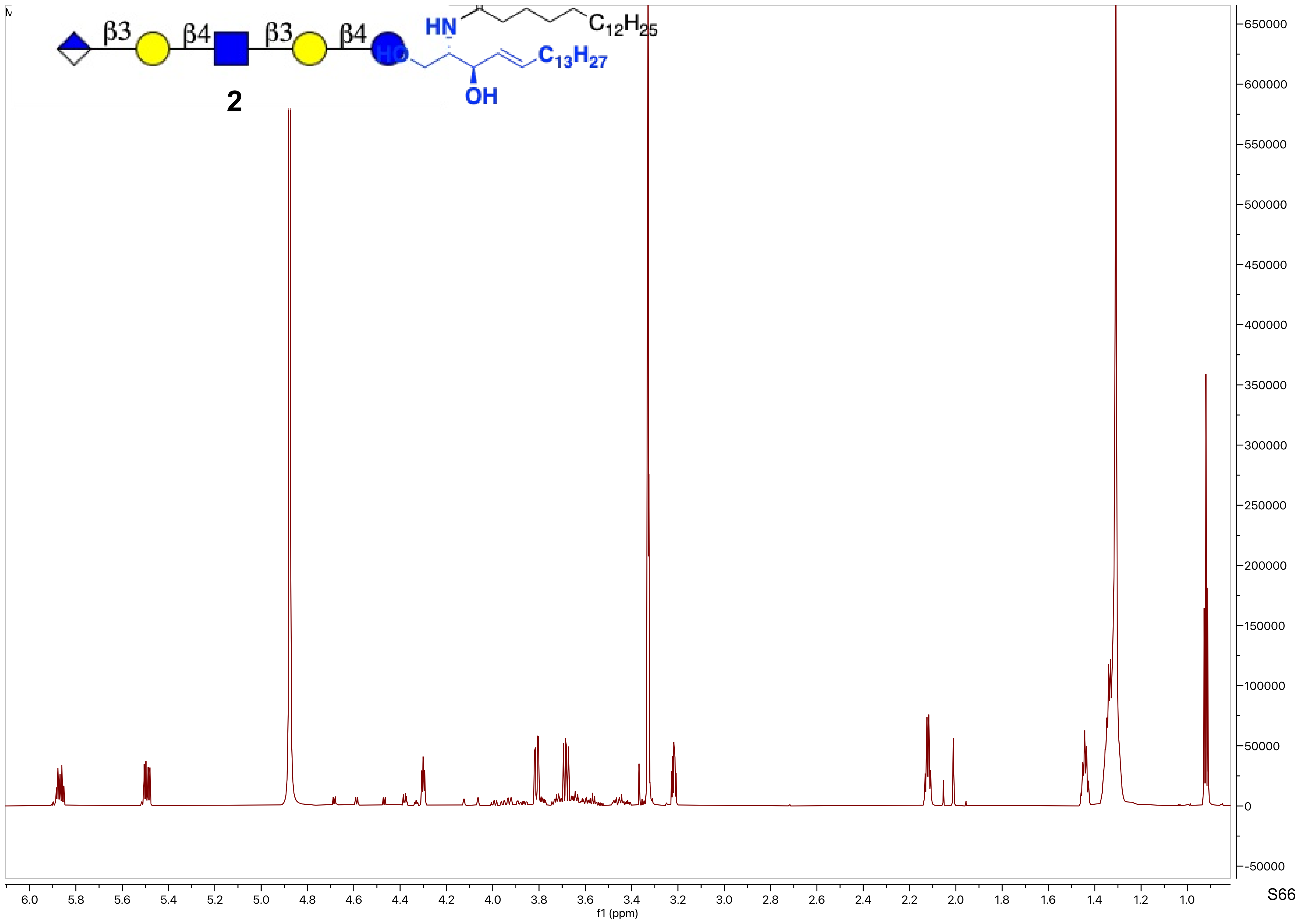

β3
β4
β3
β4
HN
HO
$C_{12}H_{25}$
$C_{13}H_{27}$
OH
2
f1 (ppm)

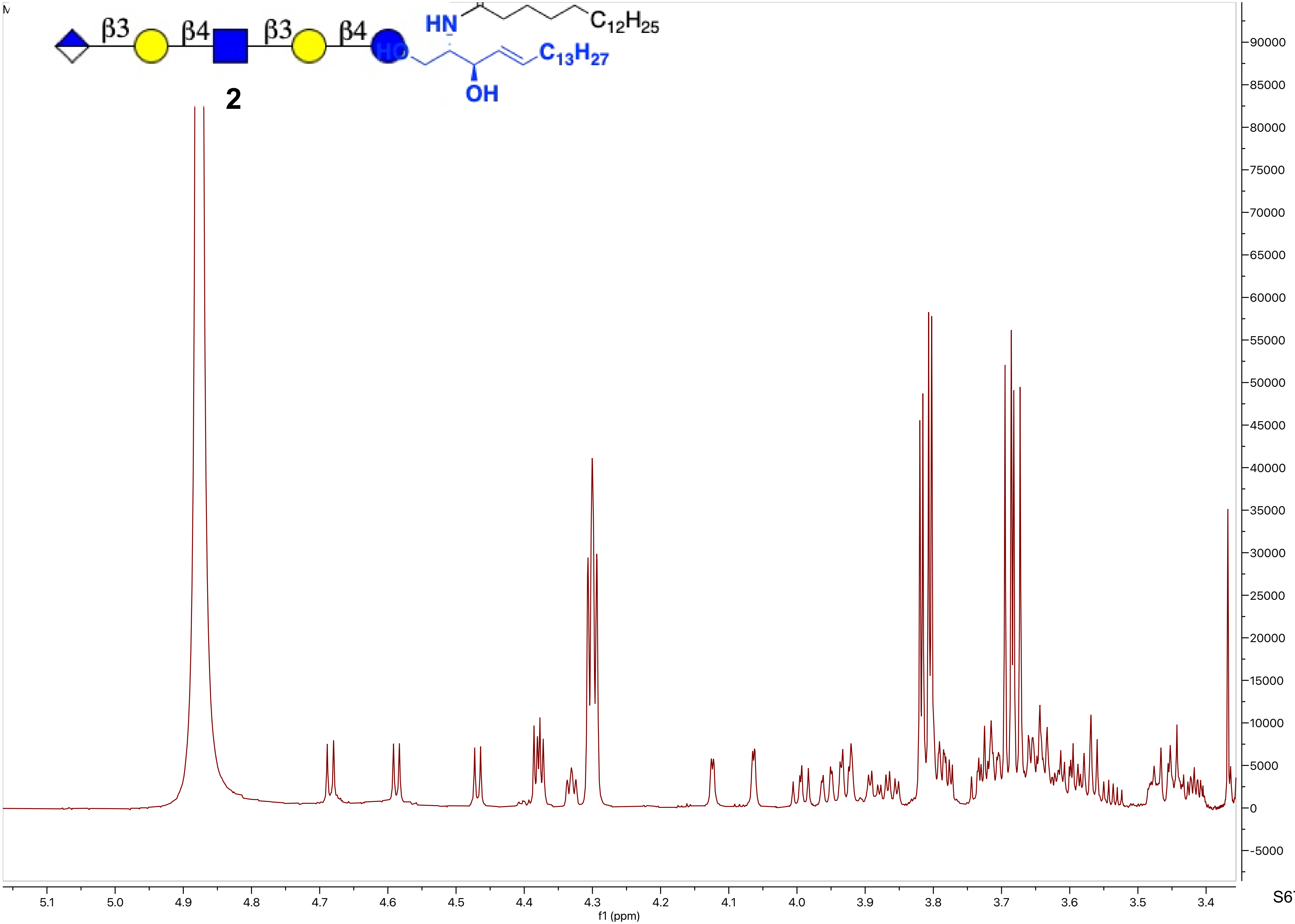
β3
β4
β3
β4
HN
$C_{12}H_{25}$
HO
$C_{13}H_{27}$
OH
2
f1 (ppm)

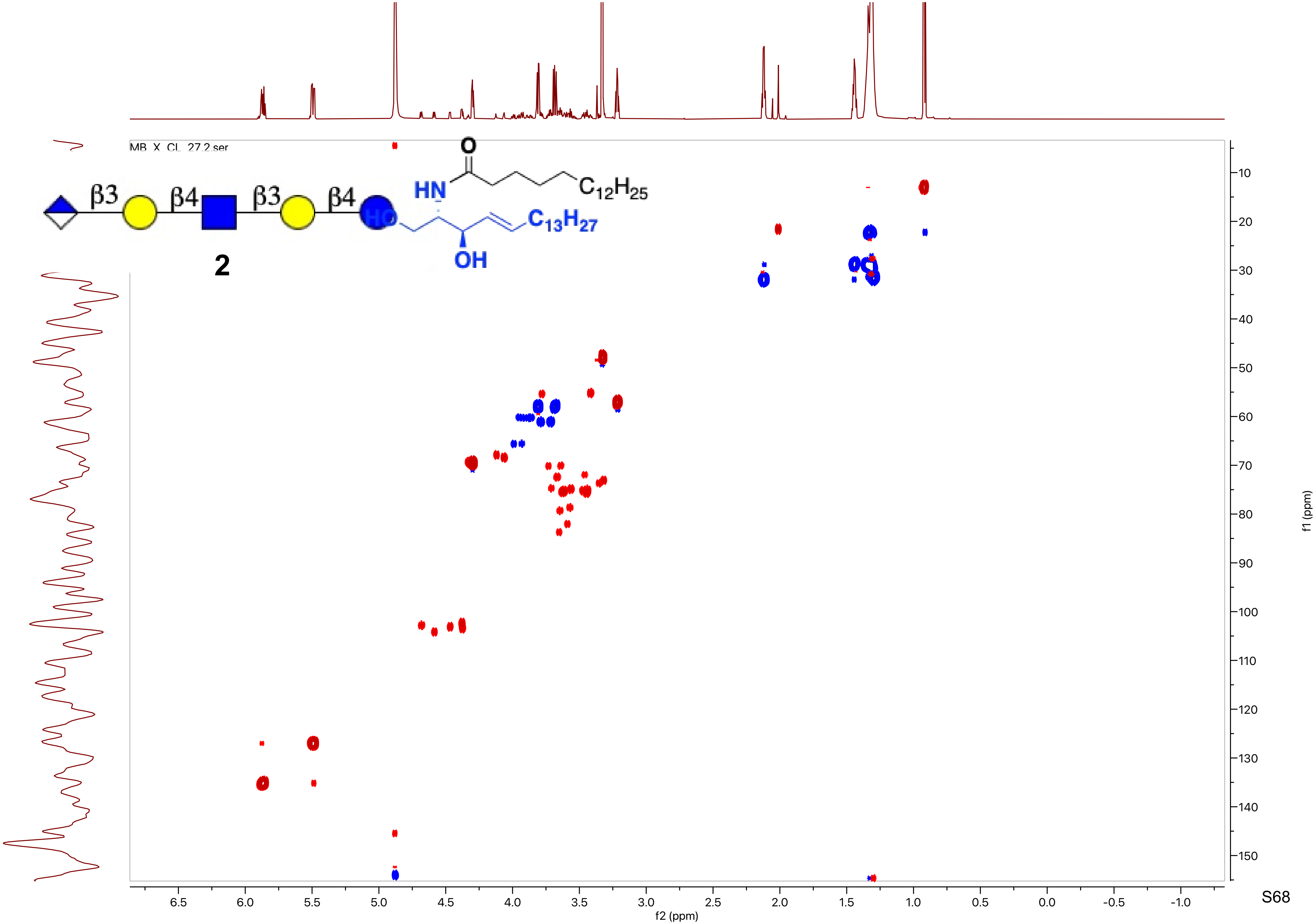
MB_X_CL_27.2.ser
β3
β4
β3
β4
O
HN
C12H25
HO
C13H27
OH
2
f2 (ppm)
f1 (ppm)

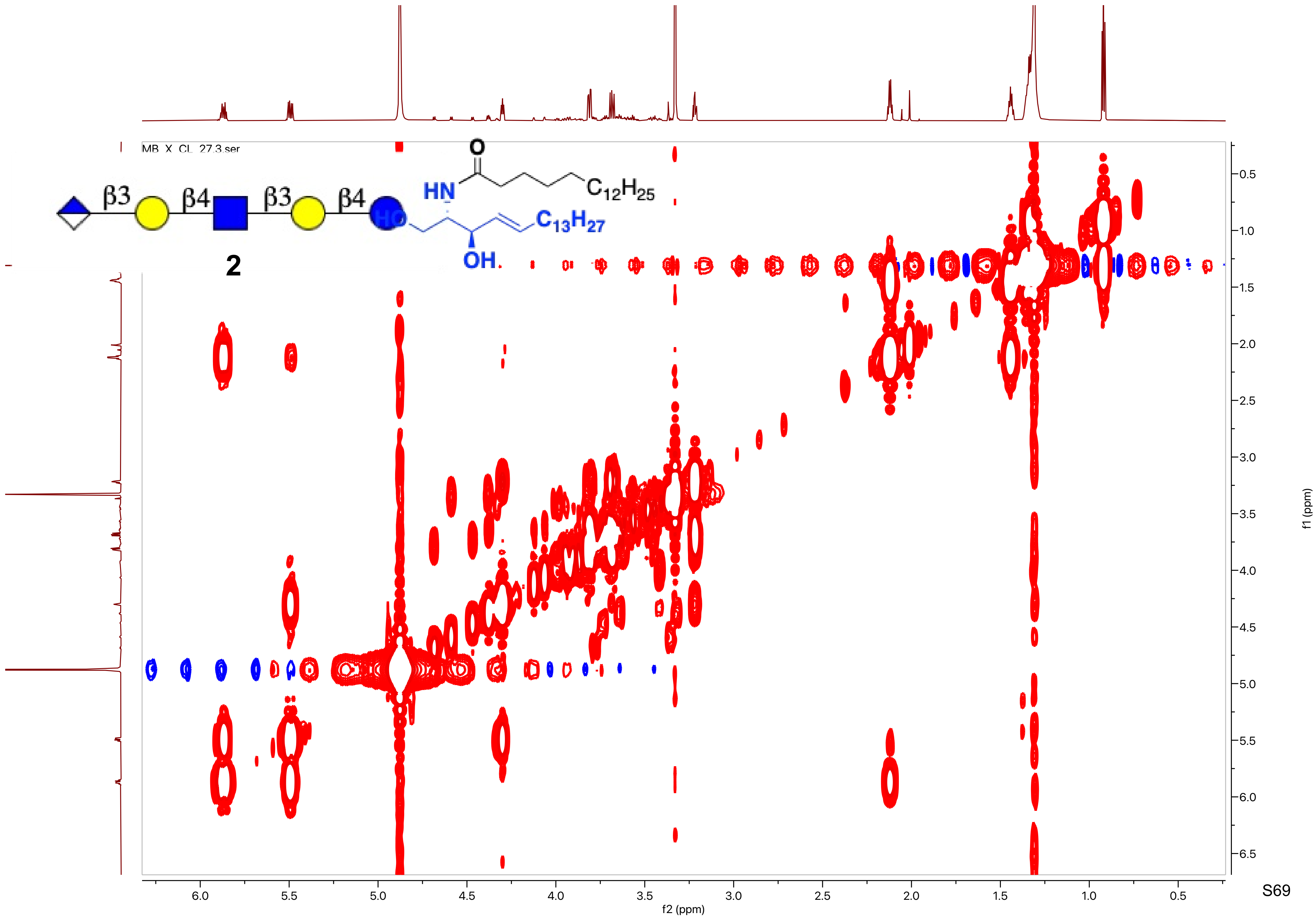

MB_X_CL_27.3.ser
β3
β4
β3
β4
HN
O
$C_{12}H_{25}$
HO
$C_{13}H_{27}$
OH
2
f2 (ppm)
f1 (ppm)

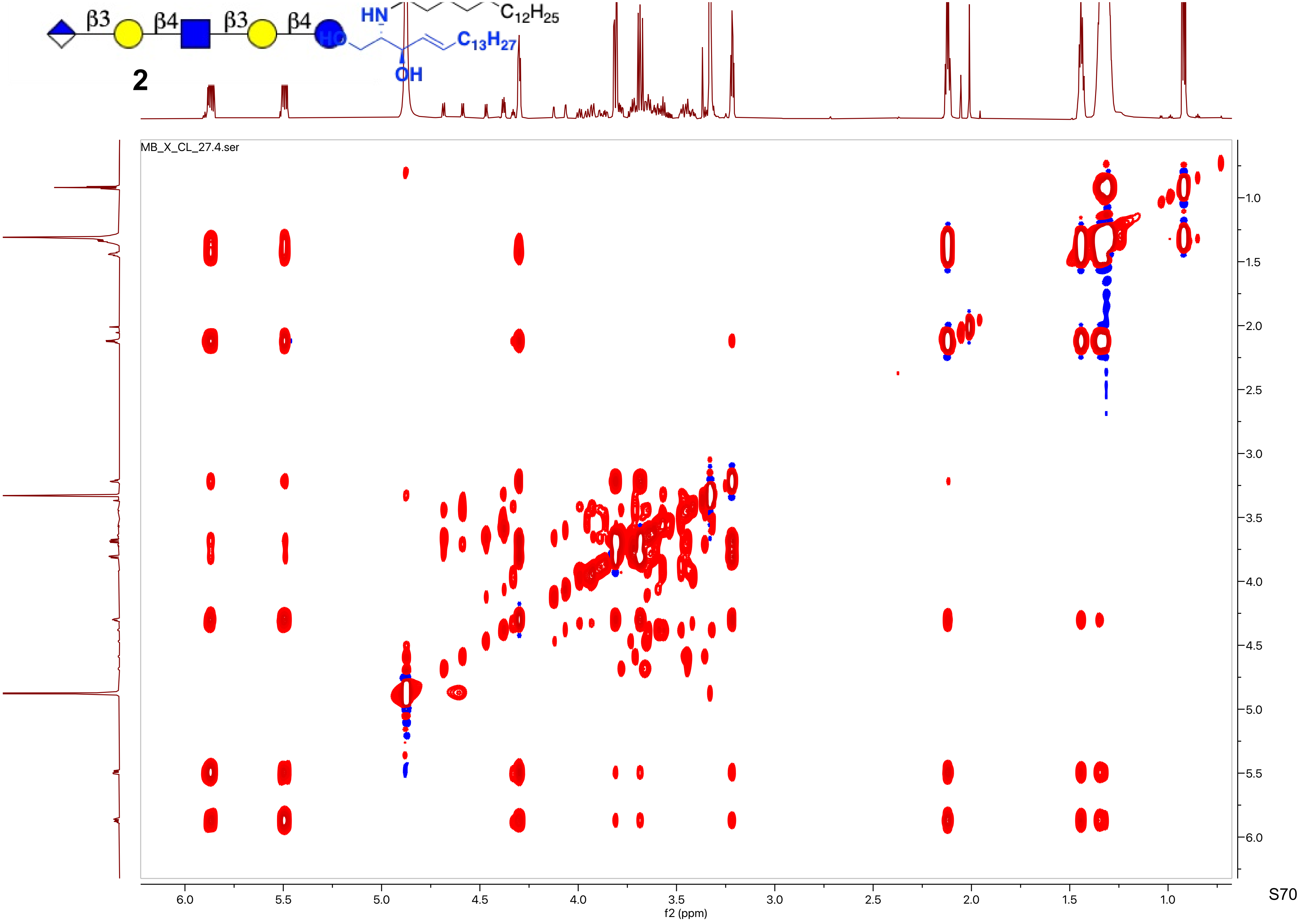

β3
β4
β3
β4
HN
$C_{12}H_{25}$
HO
$C_{13}H_{27}$
OH
2
MB_X_CL_27.4.ser
f2 (ppm)
f1 (ppm)

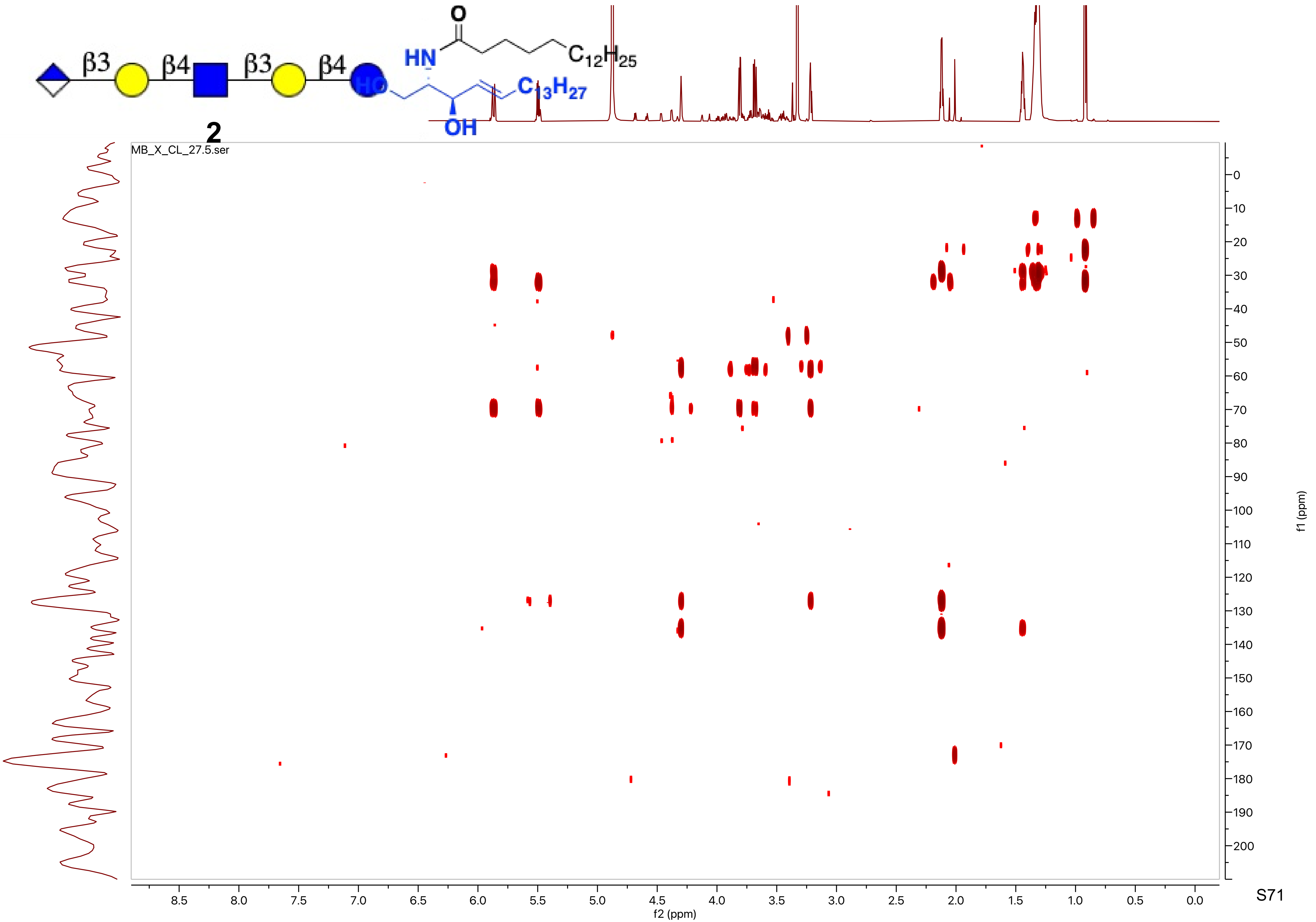
β3
β4
β3
β4
HN
O
$C_{12}H_{25}$
$C_{13}H_{27}$
OH
2
MB_X_CL_27.5.ser
f2 (ppm)
f1 (ppm)

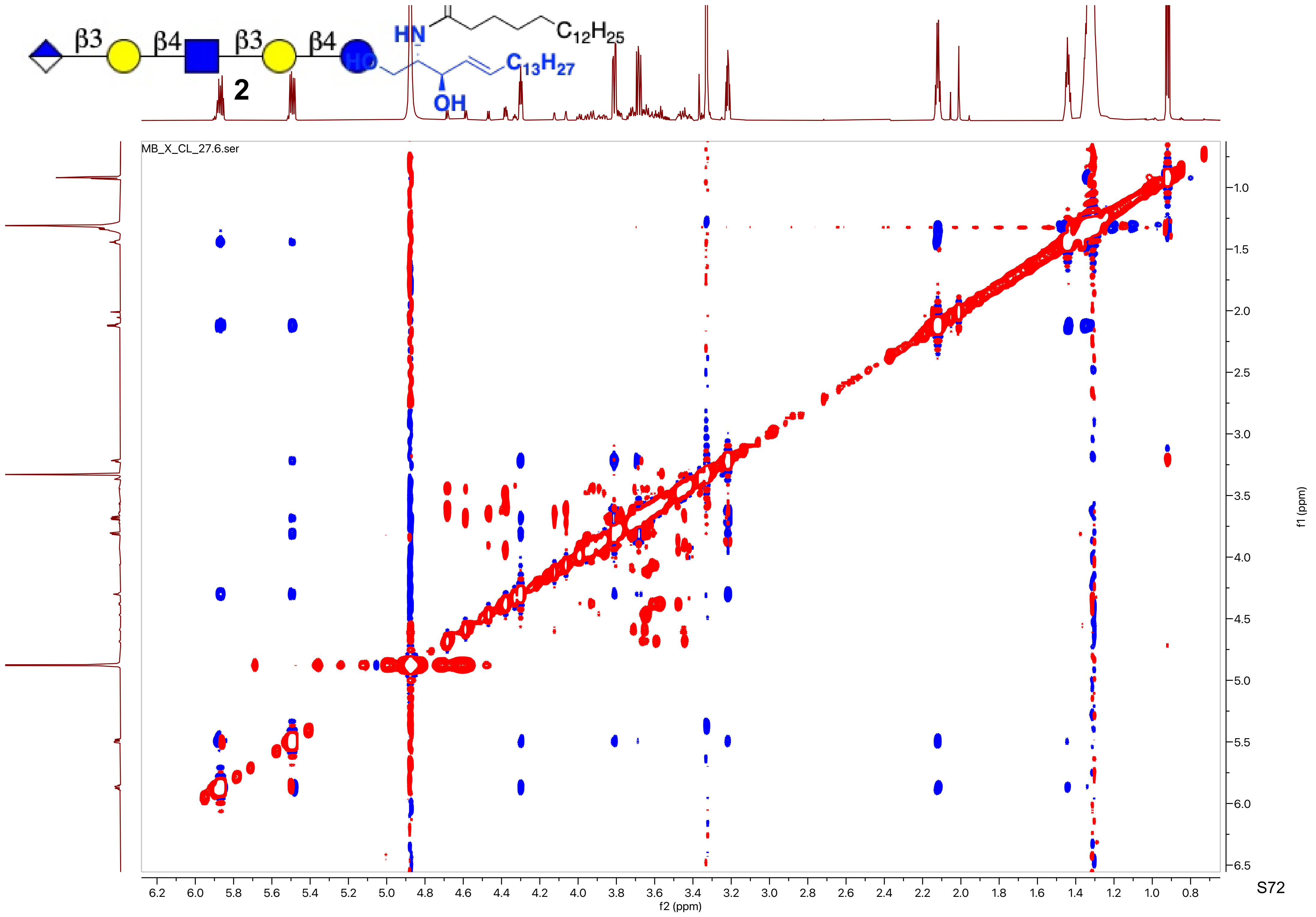
β3
β4
β3
β4
HN
C12H25
HO
C13H27
OH
2
MB_X_CL_27.6.ser
f2 (ppm)
f1 (ppm)

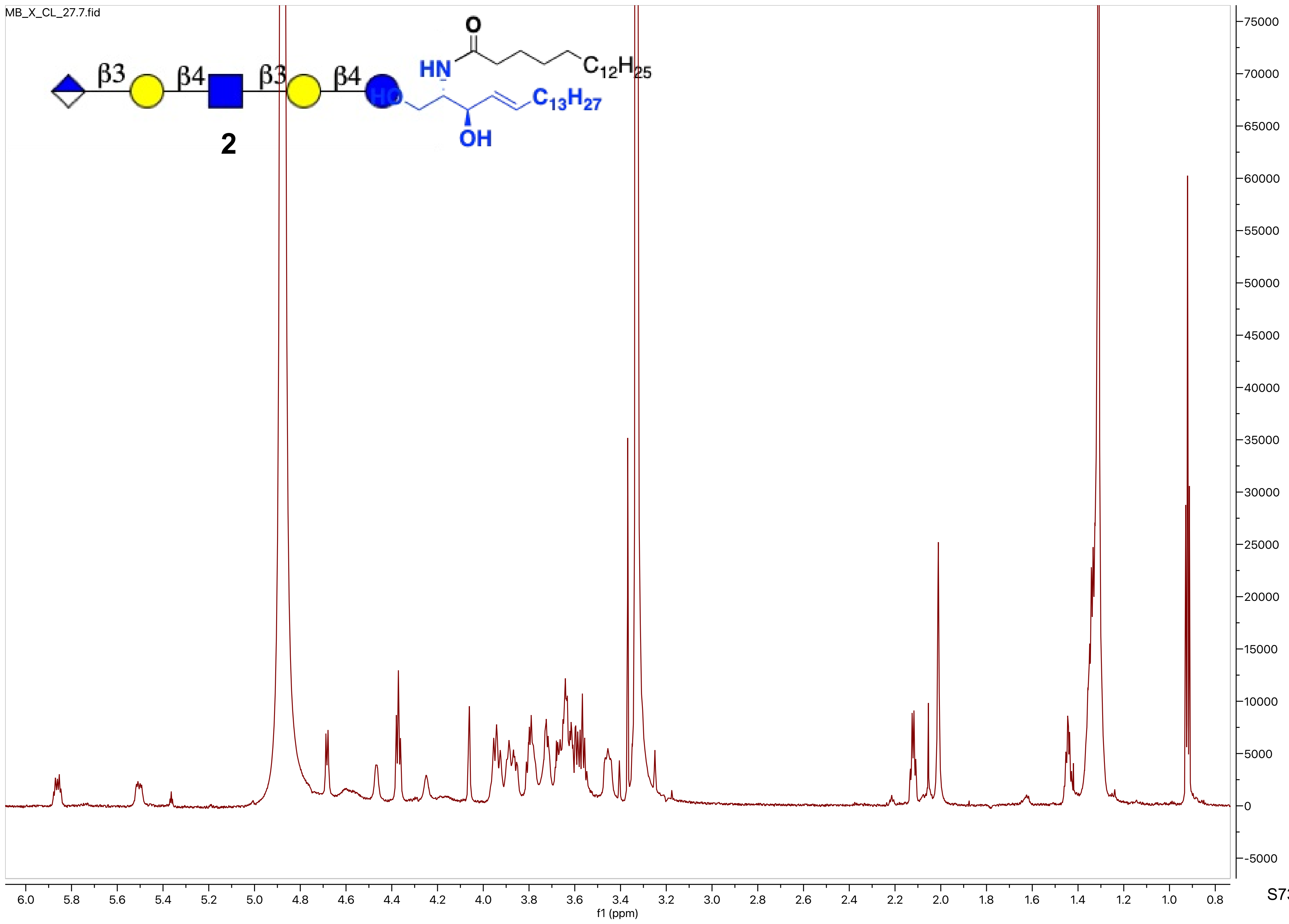
MB_X_CL_27.7.fid
β3
β4
β3
β4
O
HN
$C_{12}H_{25}$
HO
$C_{13}H_{27}$
OH
2
75000
70000
65000
60000
55000
50000
45000
40000
35000
30000
25000
20000
15000
10000
5000
0
-5000
6.0 5.8 5.6 5.4 5.2 5.0 4.8 4.6 4.4 4.2 4.0 3.8 3.6 3.4 3.2 3.0 2.8 2.6 2.4 2.2 2.0 1.8 1.6 1.4 1.2 1.0 0.8
f1 (ppm)

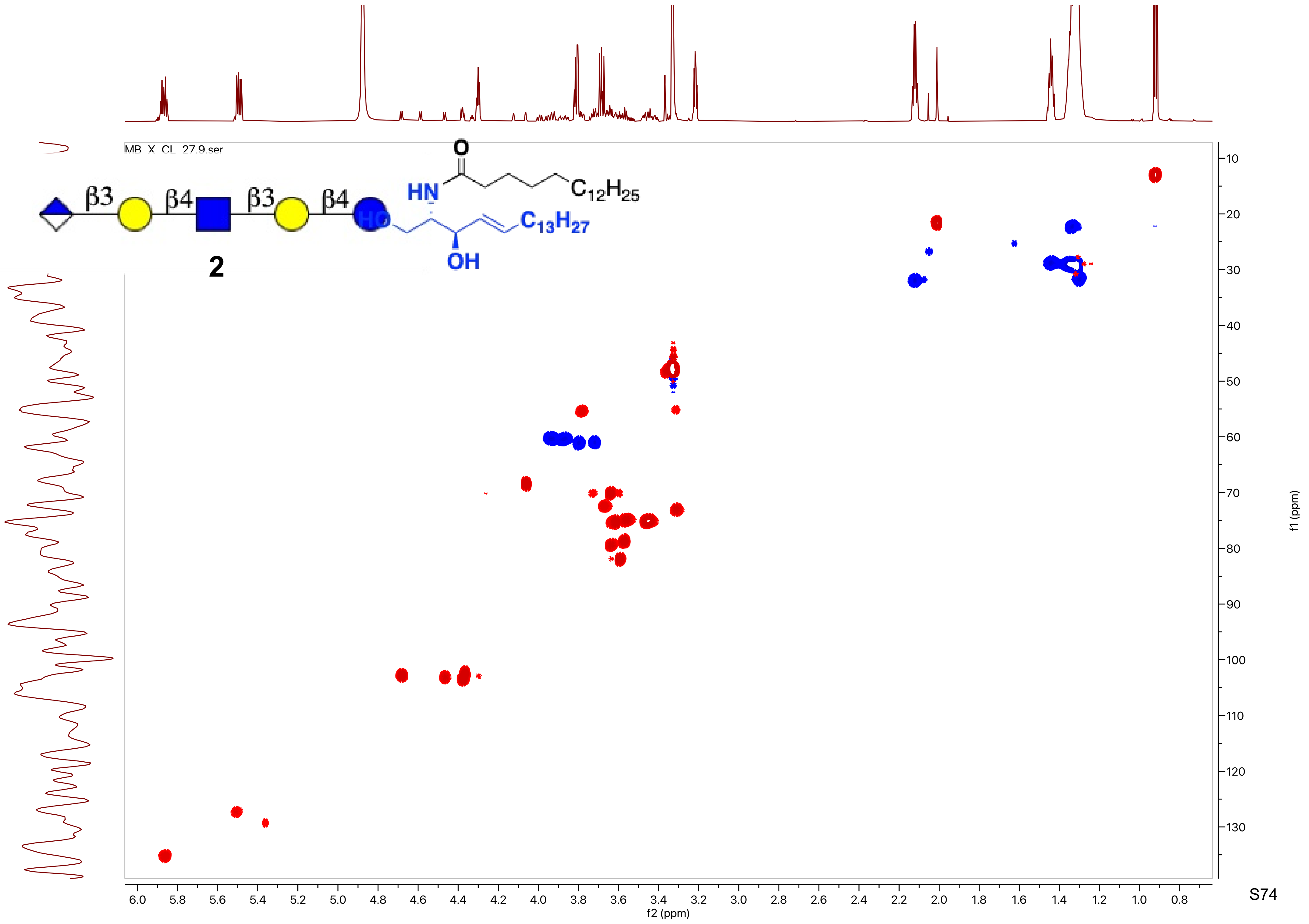
MB_X_CL_27.9.ser
β3
β4
β3
β4
HN
O
$C_{12}H_{25}$
HO
$C_{13}H_{27}$
OH
2
f1 (ppm)
f2 (ppm)

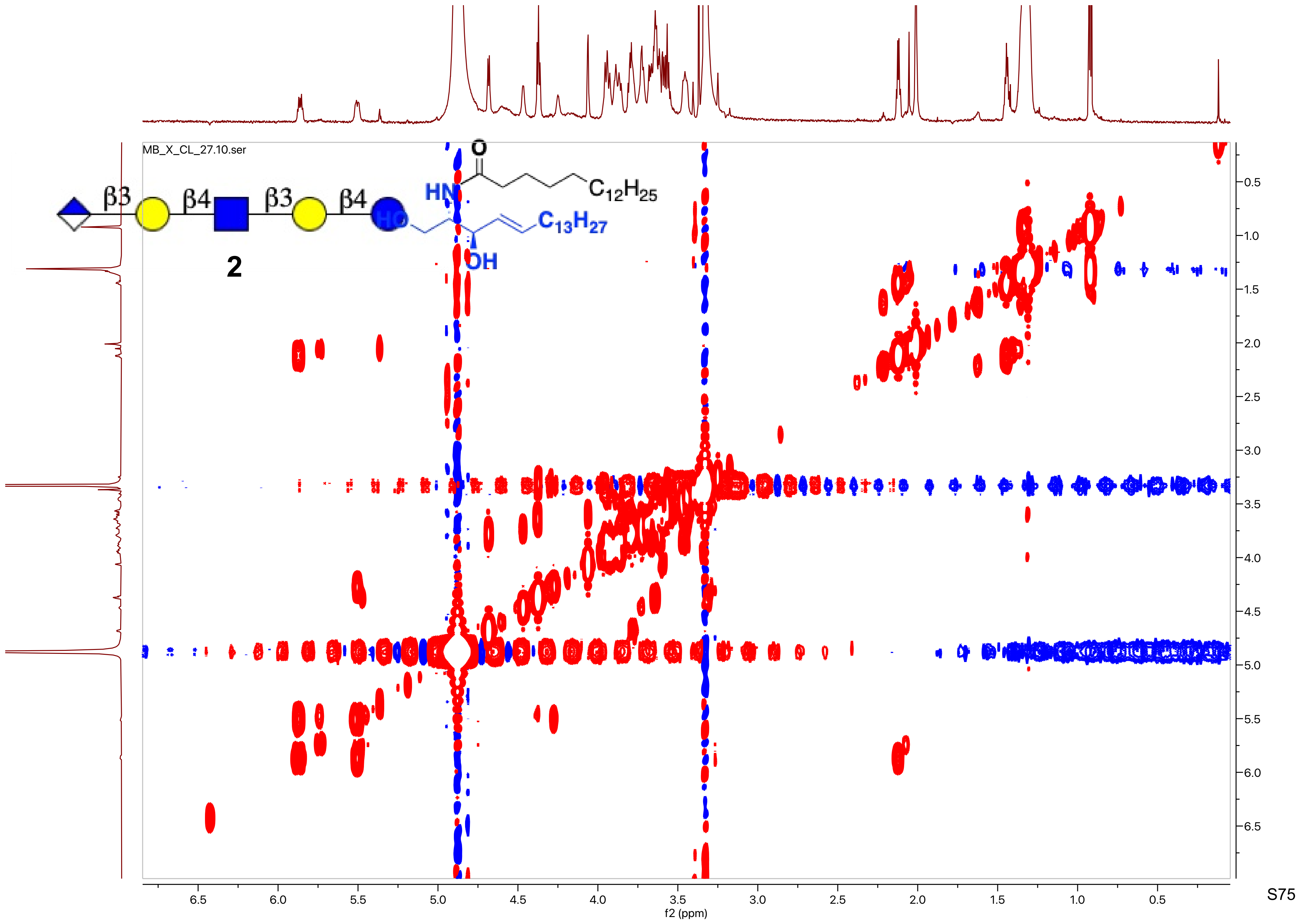
MB_X_CL_27.10.ser
β3
β4
β3
β4
HN
O
$C_{12}H_{25}$
HO
$C_{13}H_{27}$
OH
2
f2 (ppm)
f1 (ppm)

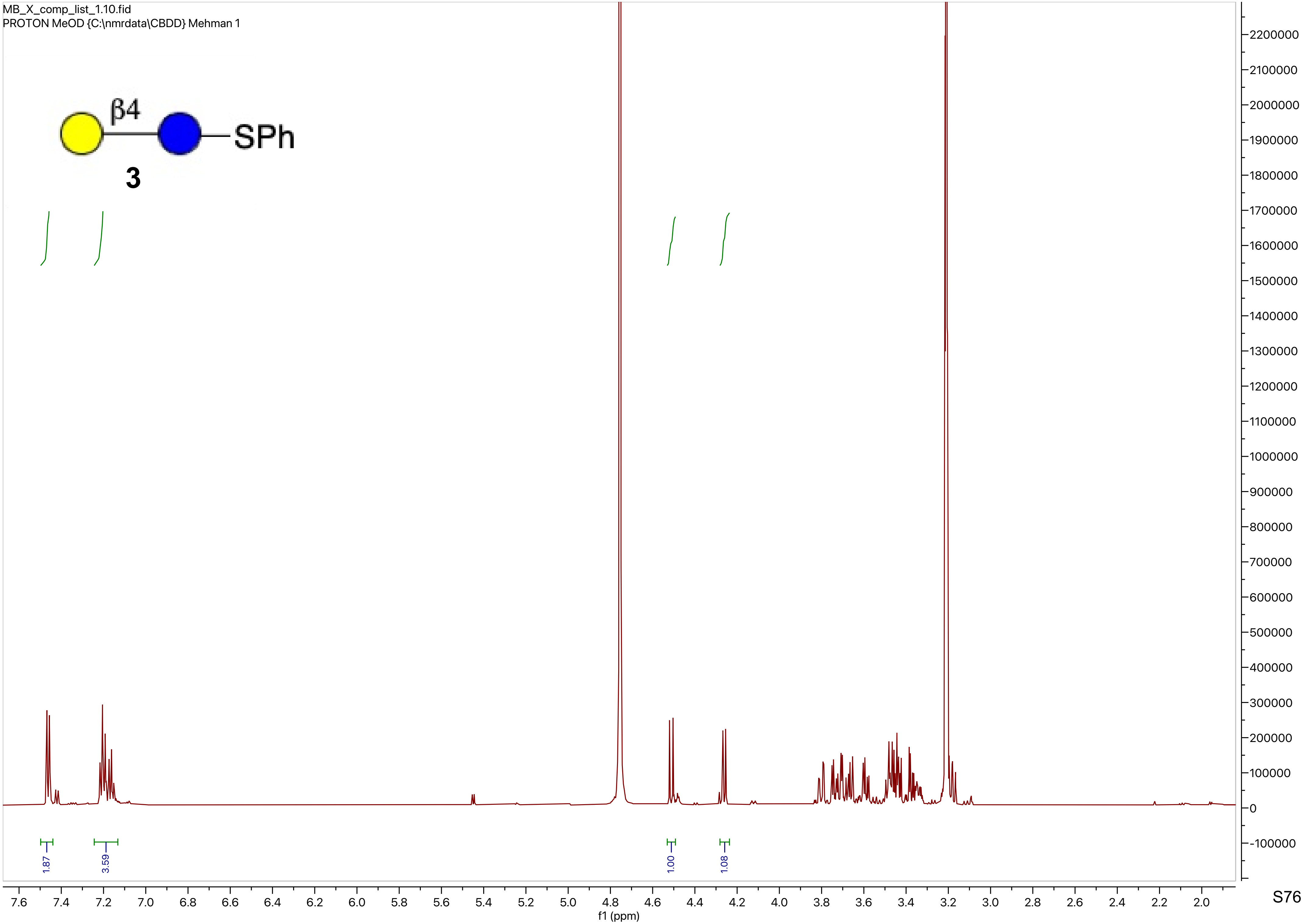
MB_X_comp_list_1.10.fid
PROTON MeOD {C:\nmrdata\CBDD} Mehman 1
β4
SPh
3
1.87
3.59
1.00
1.08
f1 (ppm)

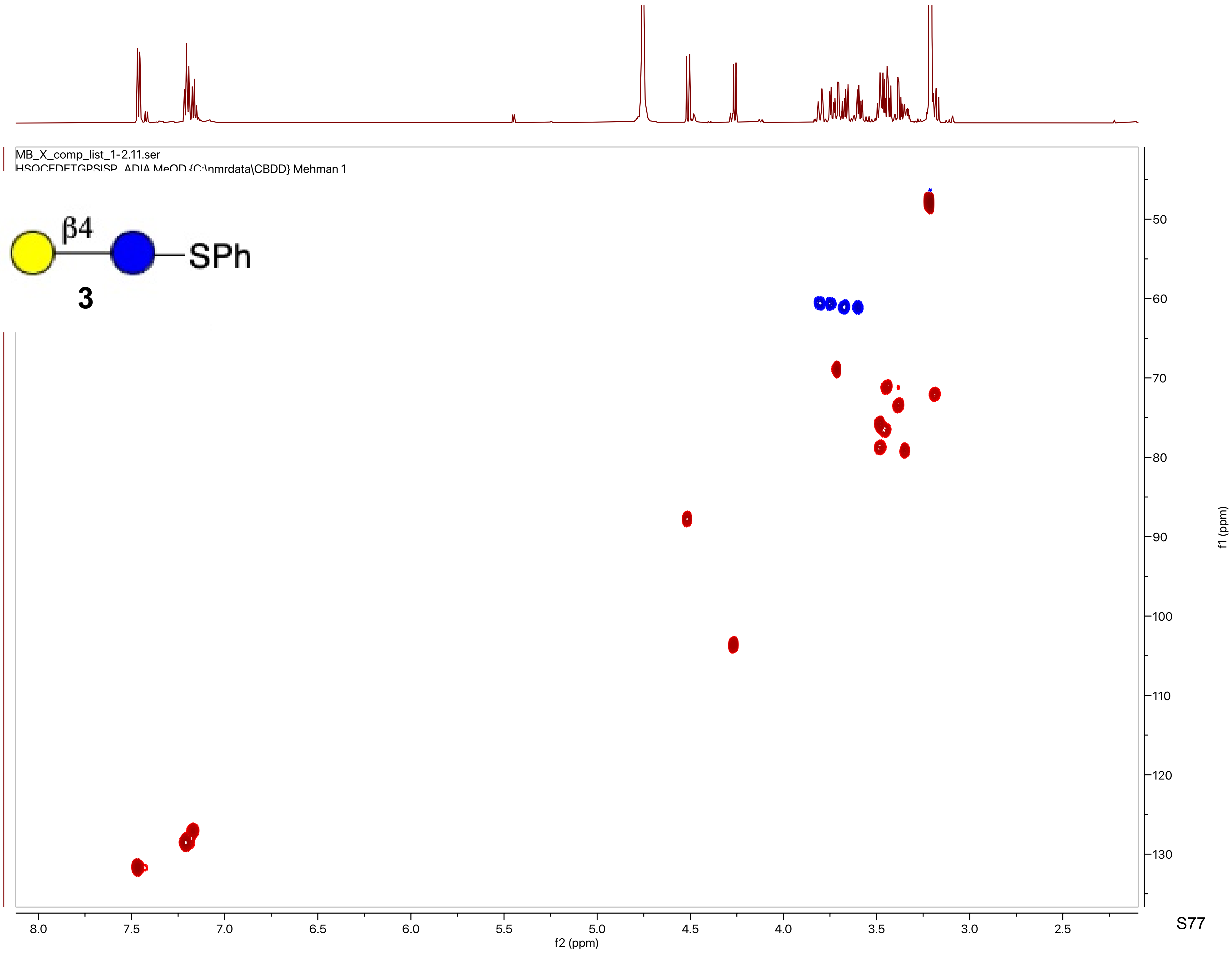
MB_X_comp_list_1-2.11.ser
HSQCEDETGPSISP_ADIA MeOD {C:\nmrdata\CBDD} Mehman 1
β4
SPh
3
f2 (ppm)
f1 (ppm)

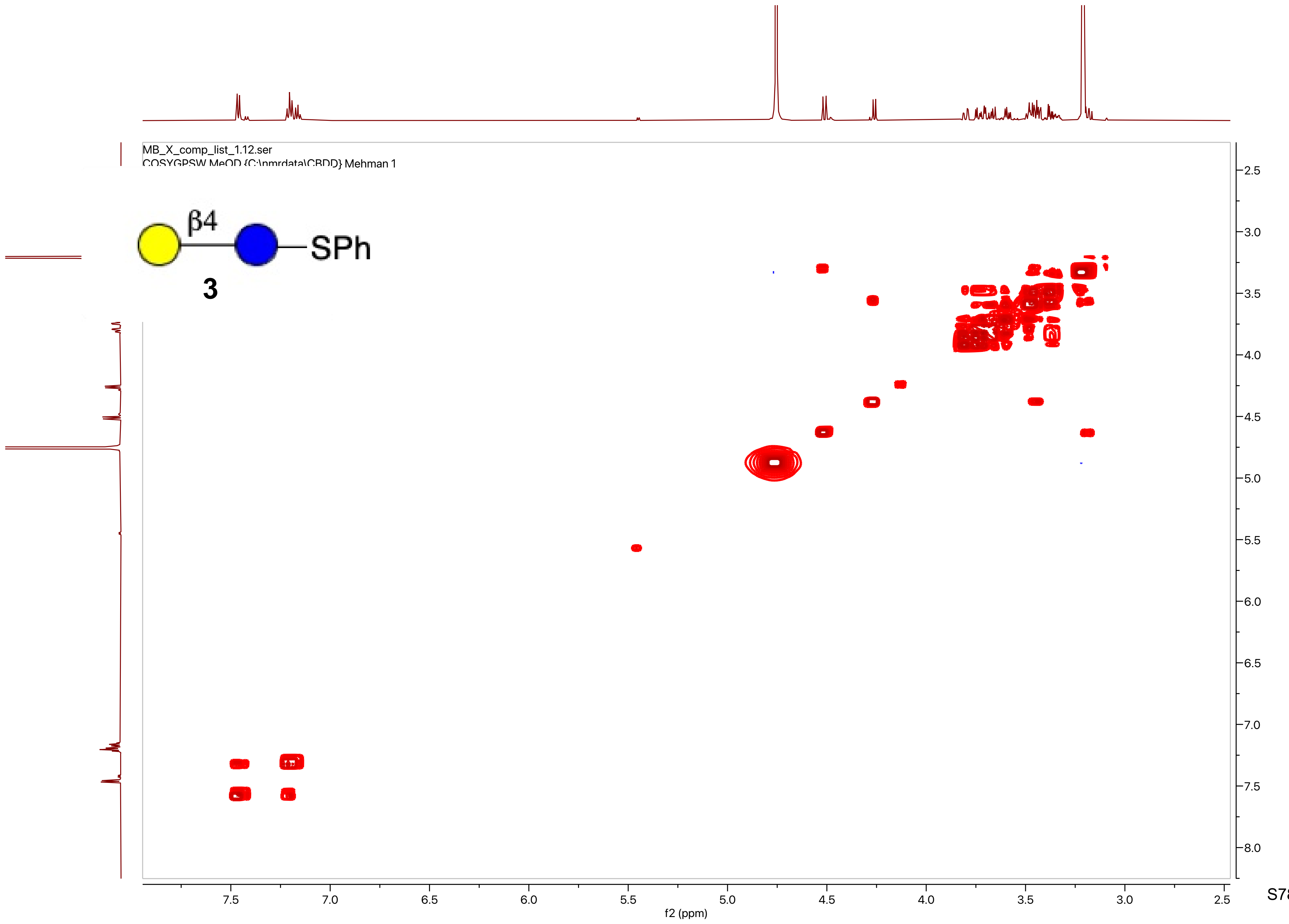
MB_X_comp_list_1.12.ser
COSYGPSW MeOD {C:\nmrdata\CBDD} Mehman 1
β4
SPh
3
f2 (ppm)
f1 (ppm)

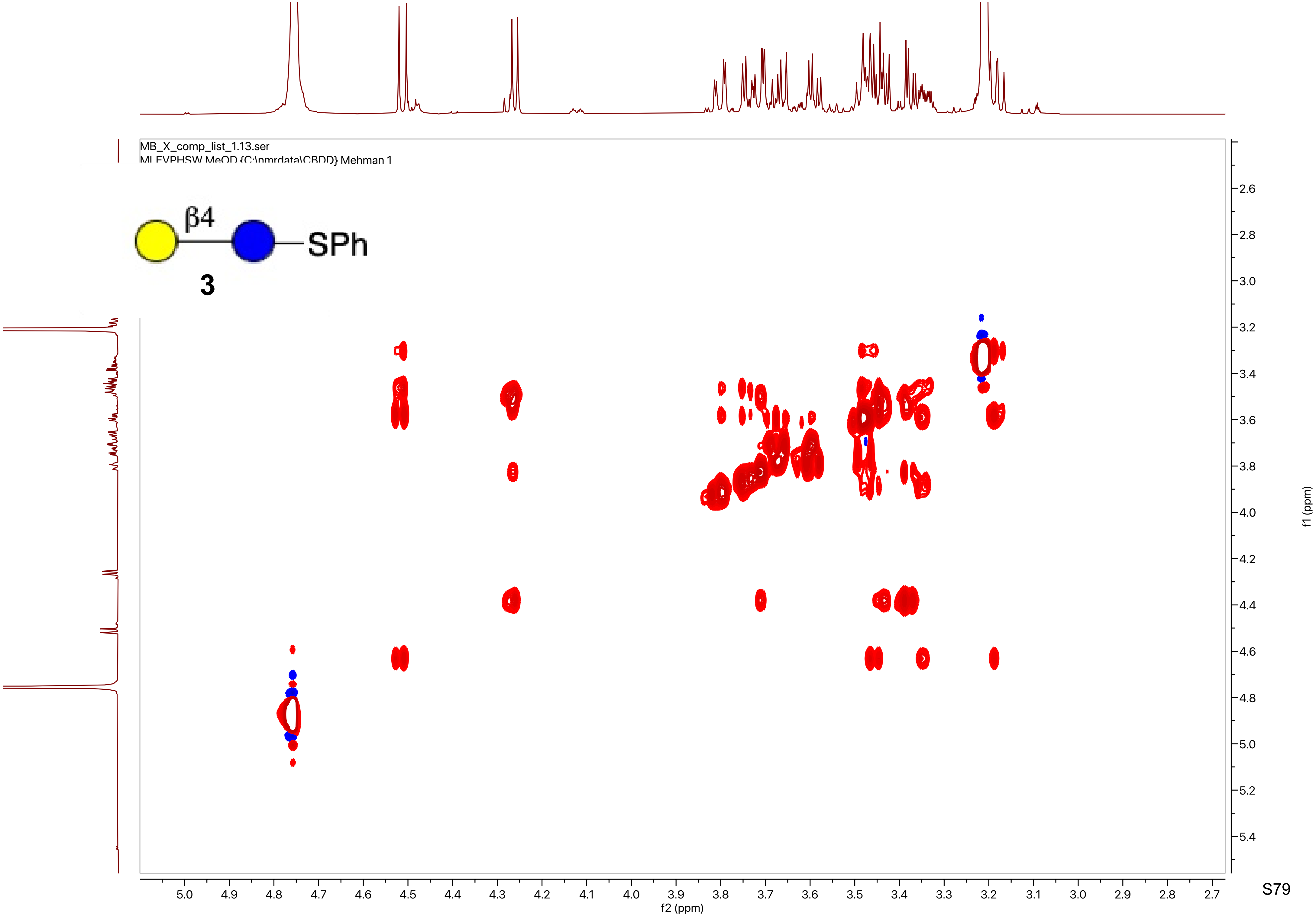

MB_X_comp_list_1.13.ser
MLEVPHSW MeOD {C:\nmrdata\CBDD} Mehman 1
β4
SPh
3
f2 (ppm)
f1 (ppm)

MB_X_24_cr.10.fid
PROTON CDCl3 {C:\nmrdata\CBDD} Mehman 57

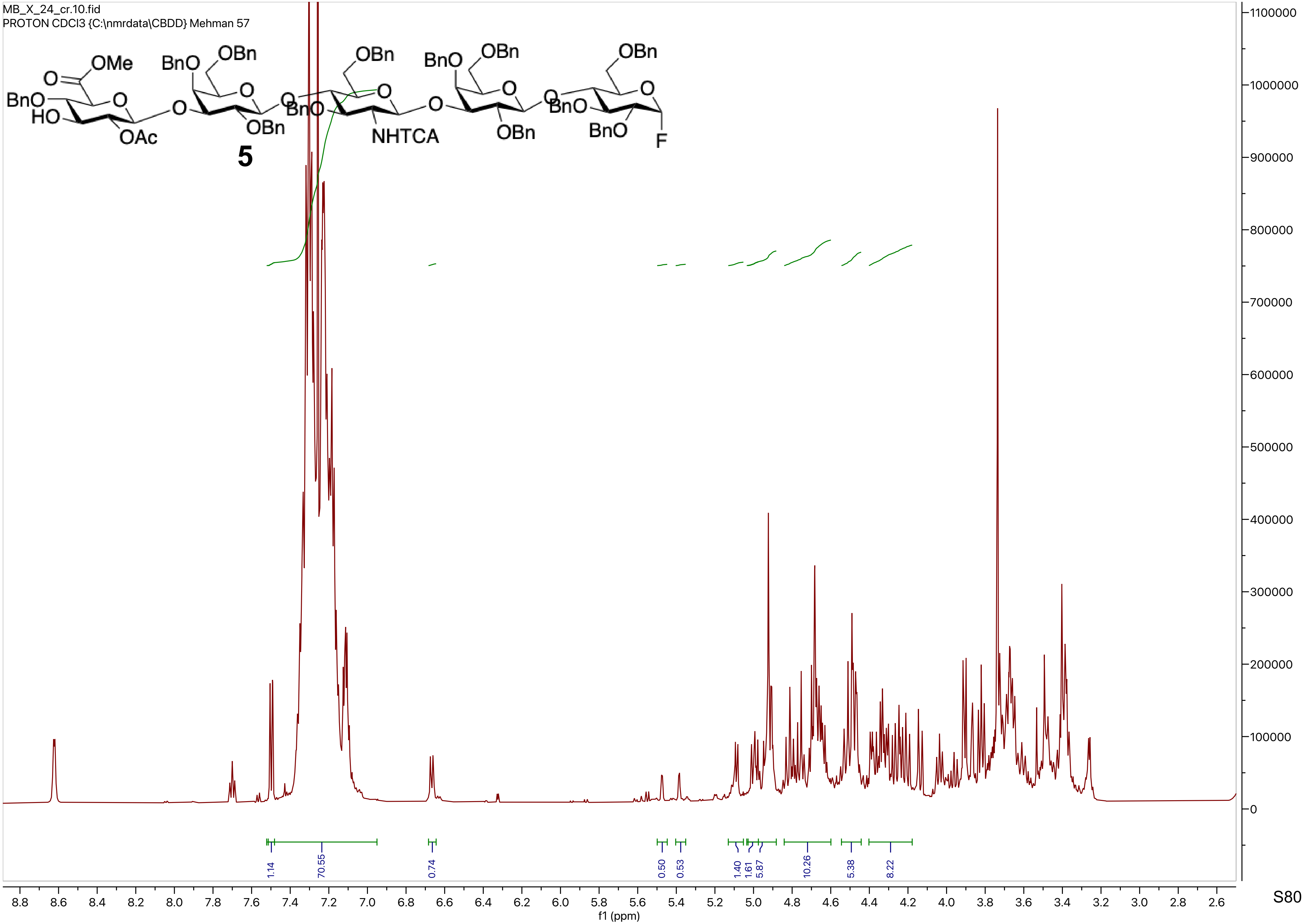

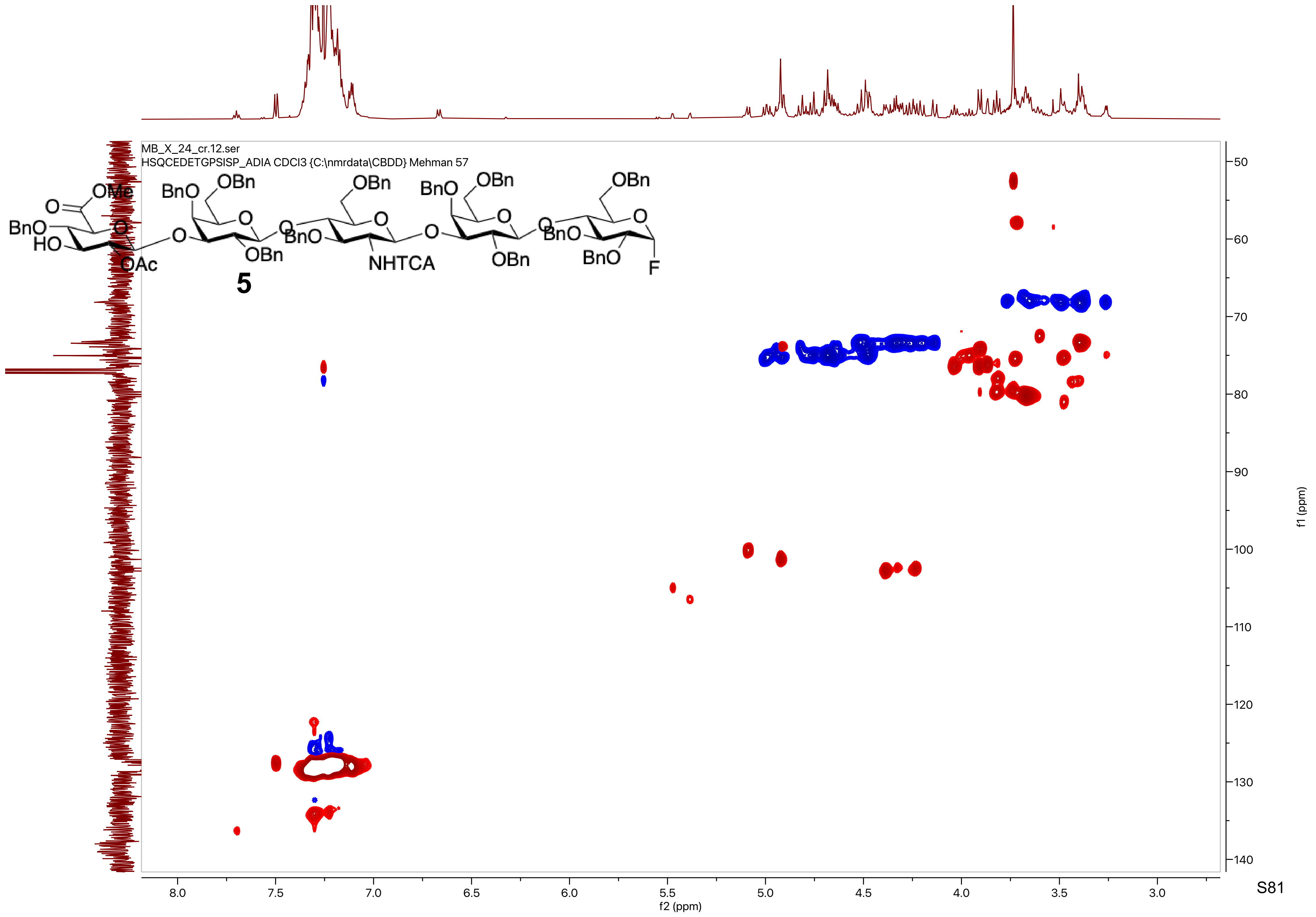
MB_X_24_cr.12.ser
HSQCEDETGPSISP_ADIA CDCl3 {C:\nmrdata\CBDD} Mehman 57
OMe
O
BnO
HO
O
OAc
BnO
OBn
O
O
OBn
O
BnO
OBn
O
NHTCA
O
BnO
OBn
O
OBn
O
BnO
OBn
O
BnO
F
5
f2 (ppm)
f1 (ppm)

MB_X_24_cr.13.fid
C13APT CDCl3 {C:\nmrdata\CBDD} Mehman 57

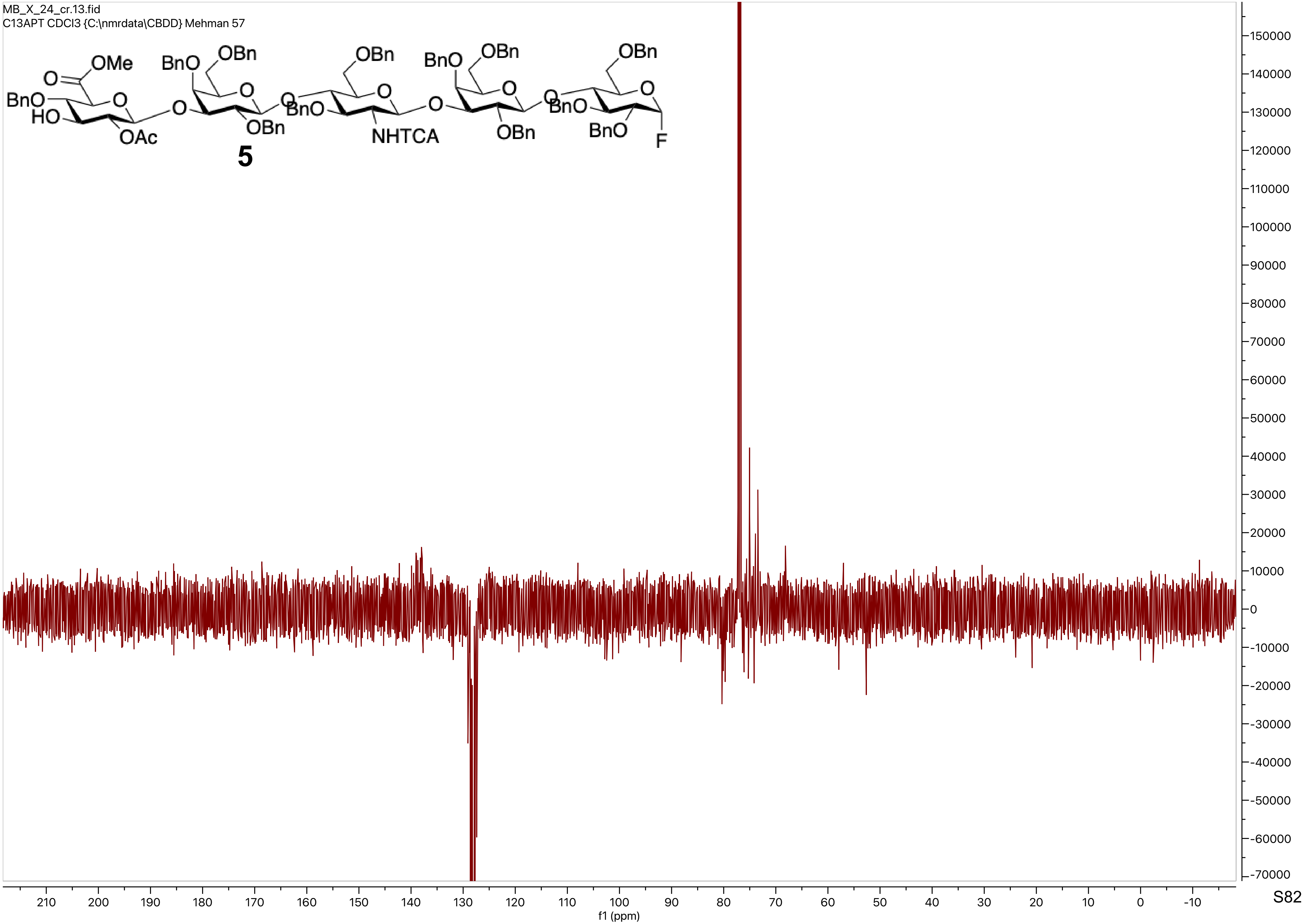

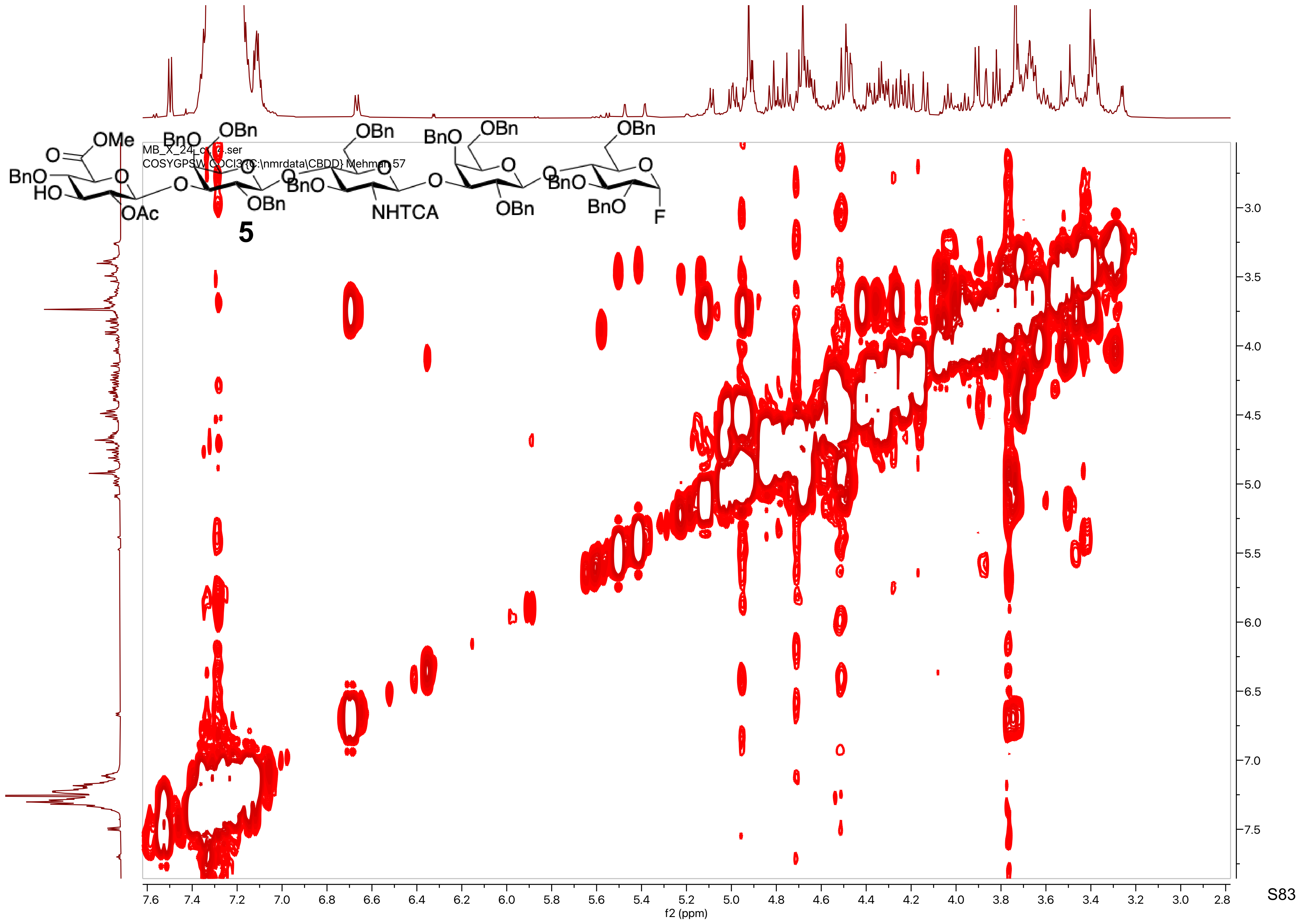
MB_X_24_c[illegible].ser
COSYGPSW CDCl3 {C:\nmrdata\CBDD} Mehman 57
5
f2 (ppm)
f1 (ppm)

MB_X_24_cr.11.fid
F19 CDCl3 {C:\nmrdata\CBDD} Mehman 57

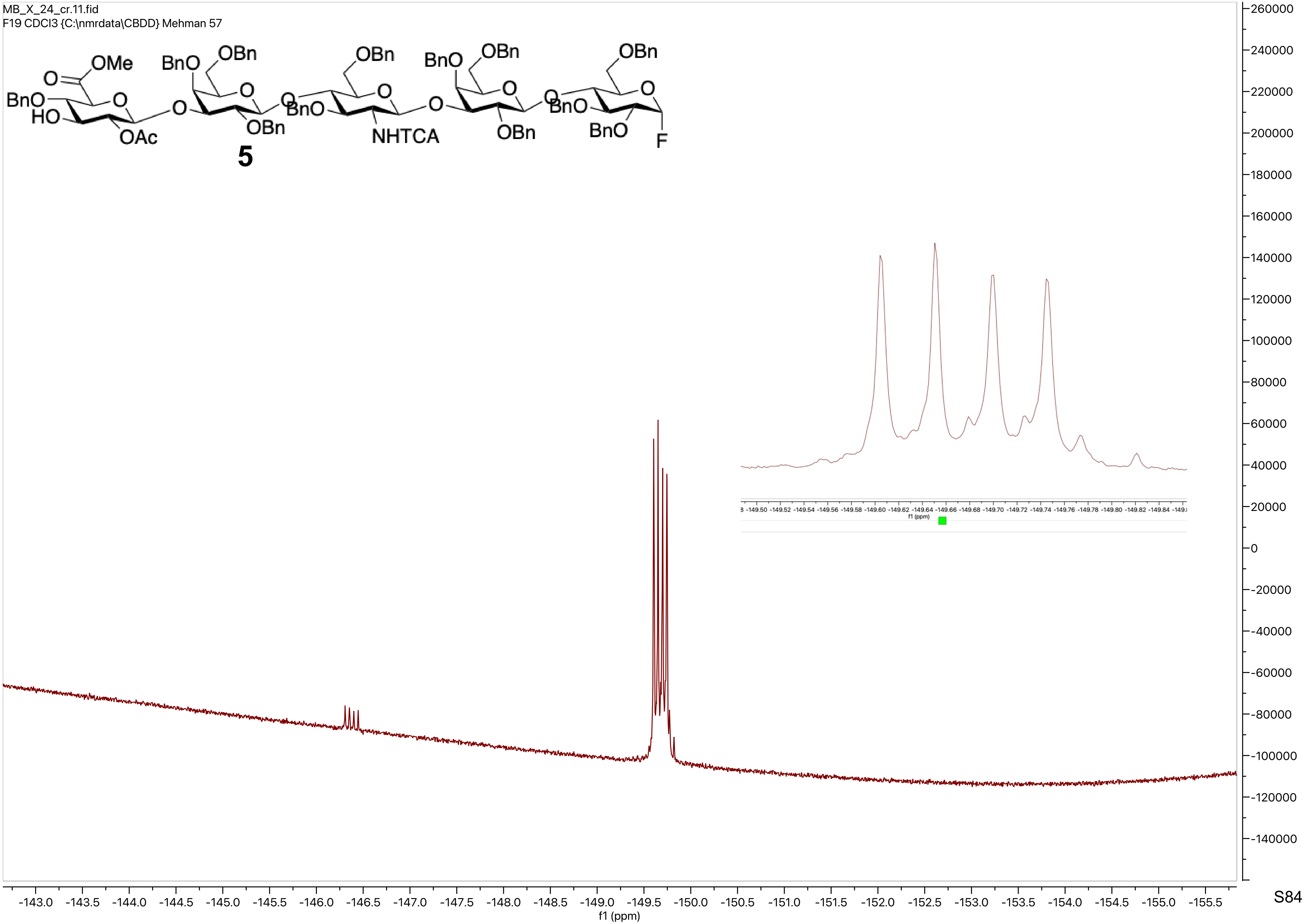

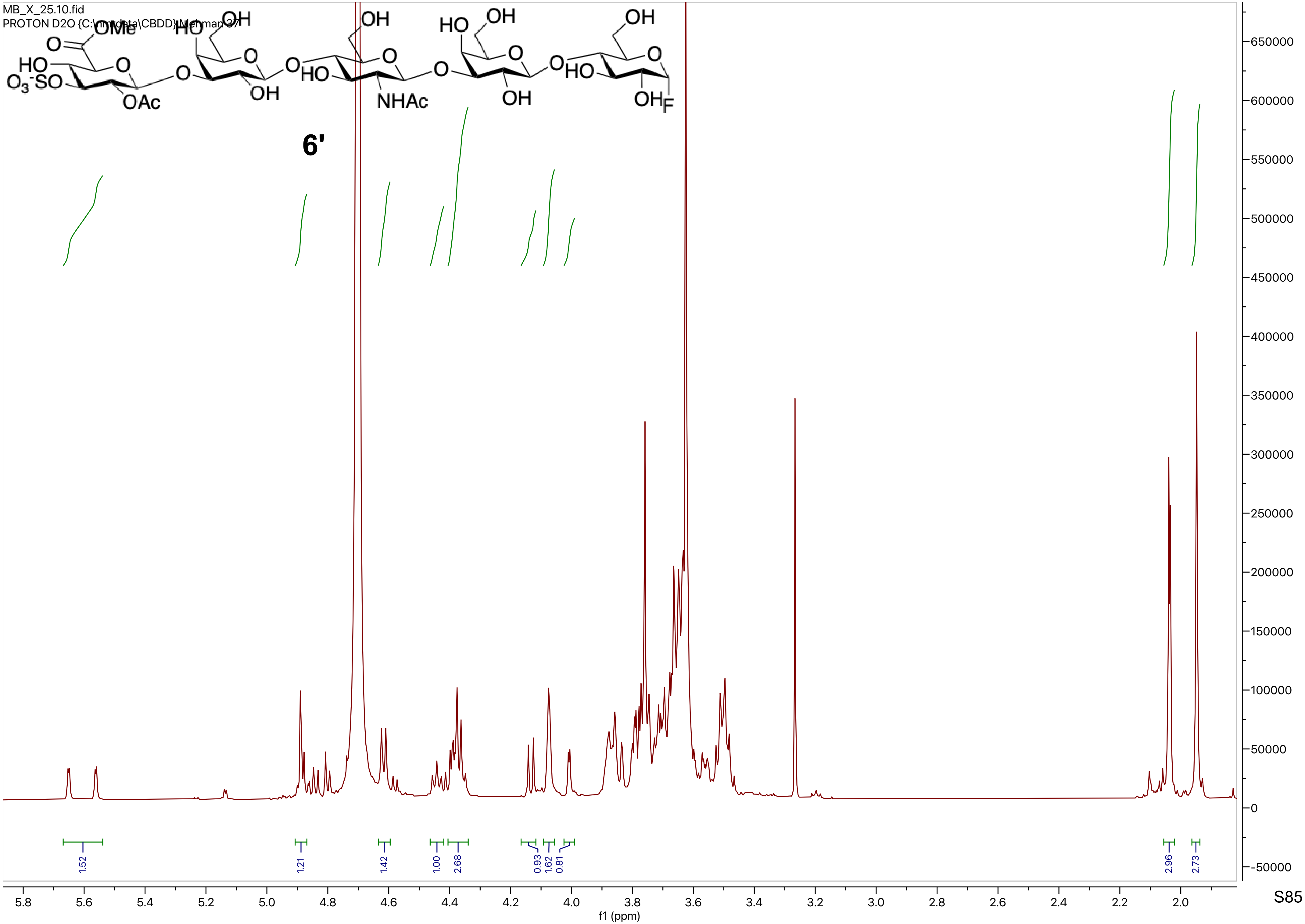

MB_X_25.10.fid
PROTON D2O {C:\nmrdata\CBDD} Mehman 37
6'
1.52
1.21
1.42
1.00
2.68
0.93
1.62
0.81
2.96
2.73
5.8
5.6
5.4
5.2
5.0
4.8
4.6
4.4
4.2
4.0
3.8
3.6
3.4
3.2
3.0
2.8
2.6
2.4
2.2
2.0
f1 (ppm)
650000
600000
550000
500000
450000
400000
350000
300000
250000
200000
150000
100000
50000
0
-50000

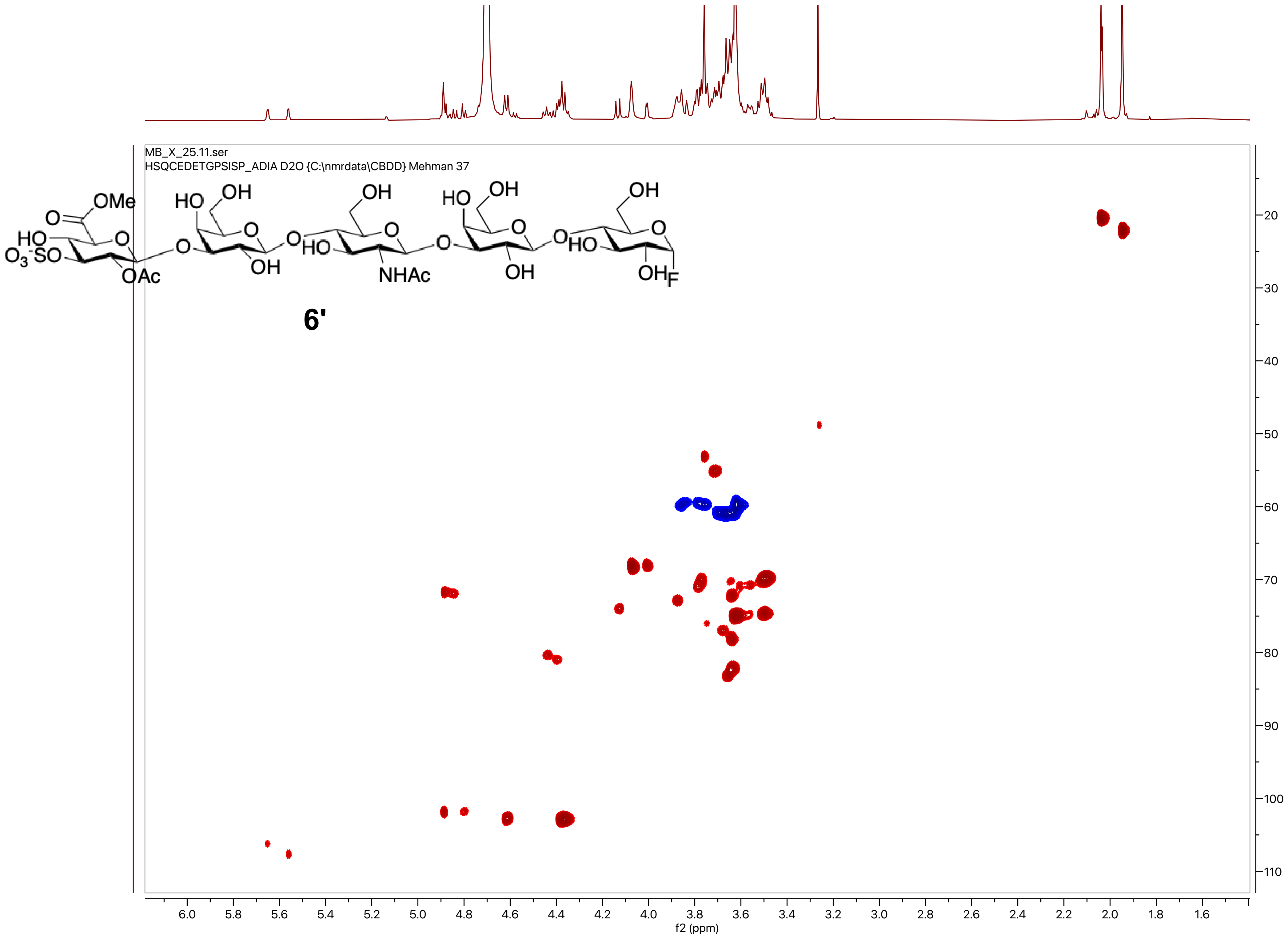
MB_X_25.11.ser
HSQCEDETGPSISP_ADIA D2O {C:\nmrdata\CBDD} Mehman 37
6'
f2 (ppm)
f1 (ppm)

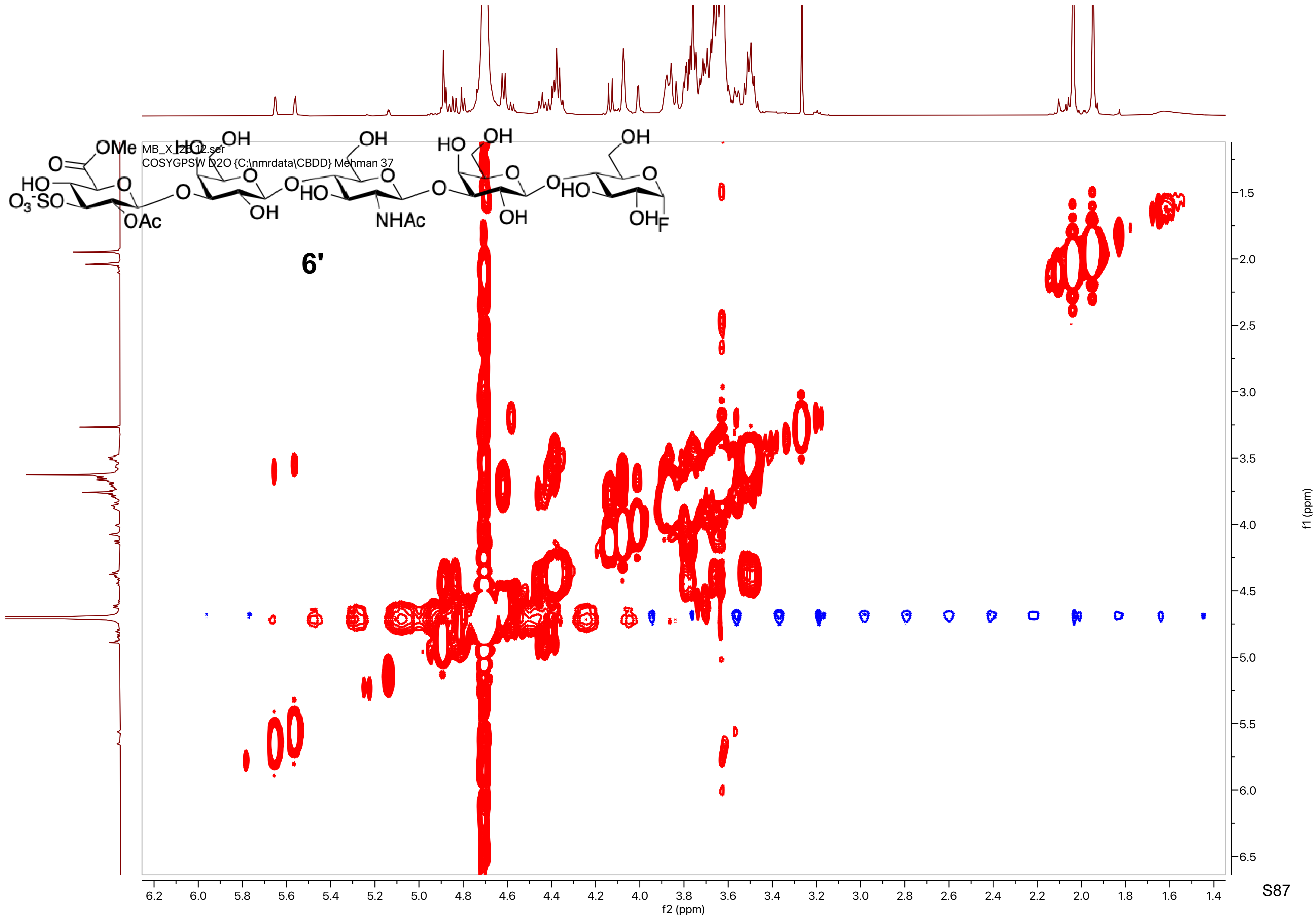
MB_X_2B 12.ser
COSYGPSW D2O {C:\nmrdata\CBDD} Mehman 37
6'
f2 (ppm)
f1 (ppm)

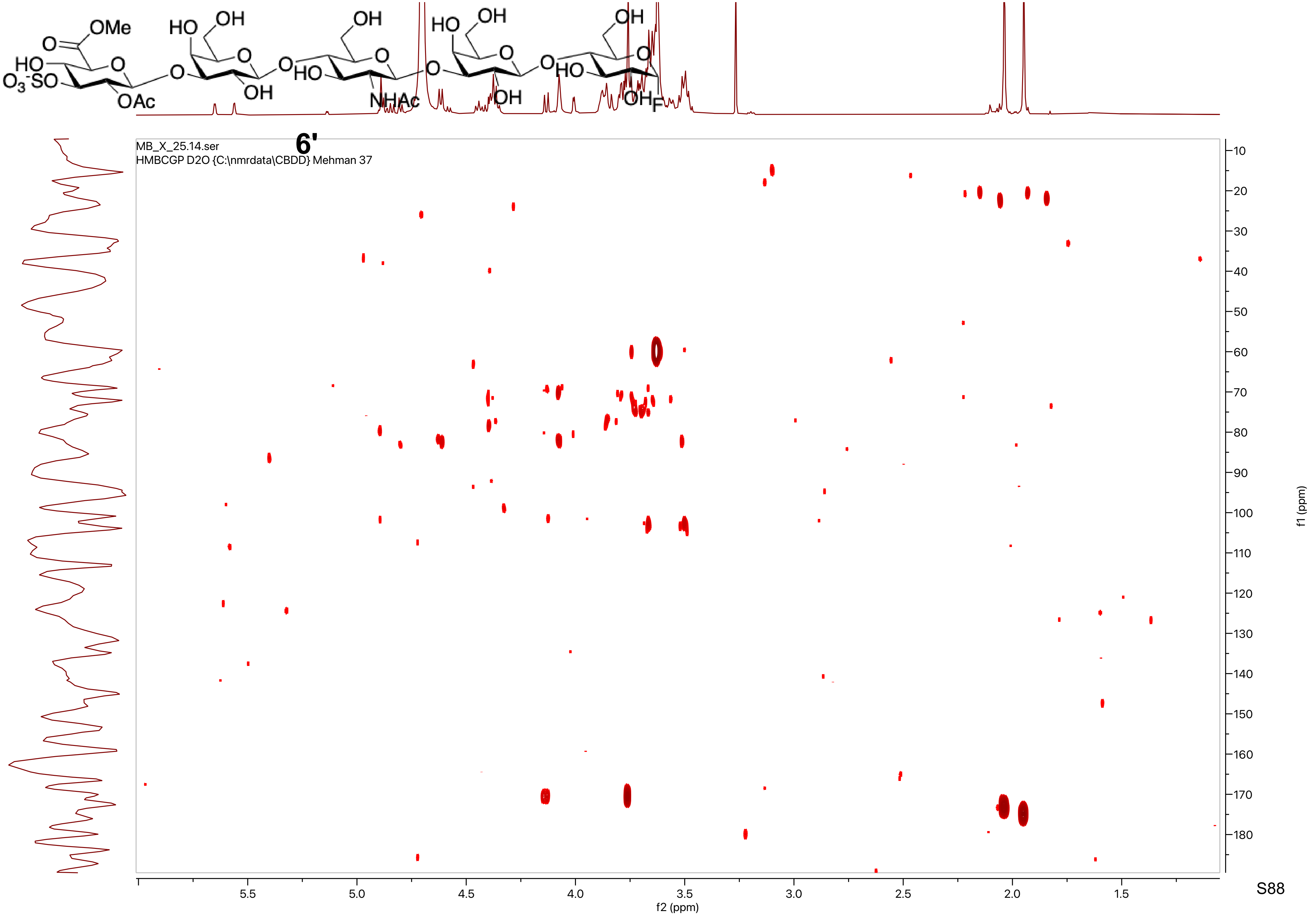

OMe
O
HO
$O_3^-SO$
O
OAc
HO
OH
O
O
OH
O
OH
O
HO
NHAc
O
HO
OH
O
OH
O
O
OH
O
HO
OH
F
6'
MB_X_25.14.ser
HMBCGP D2O {C:\nmrdata\CBDD} Mehman 37
f2 (ppm)
f1 (ppm)

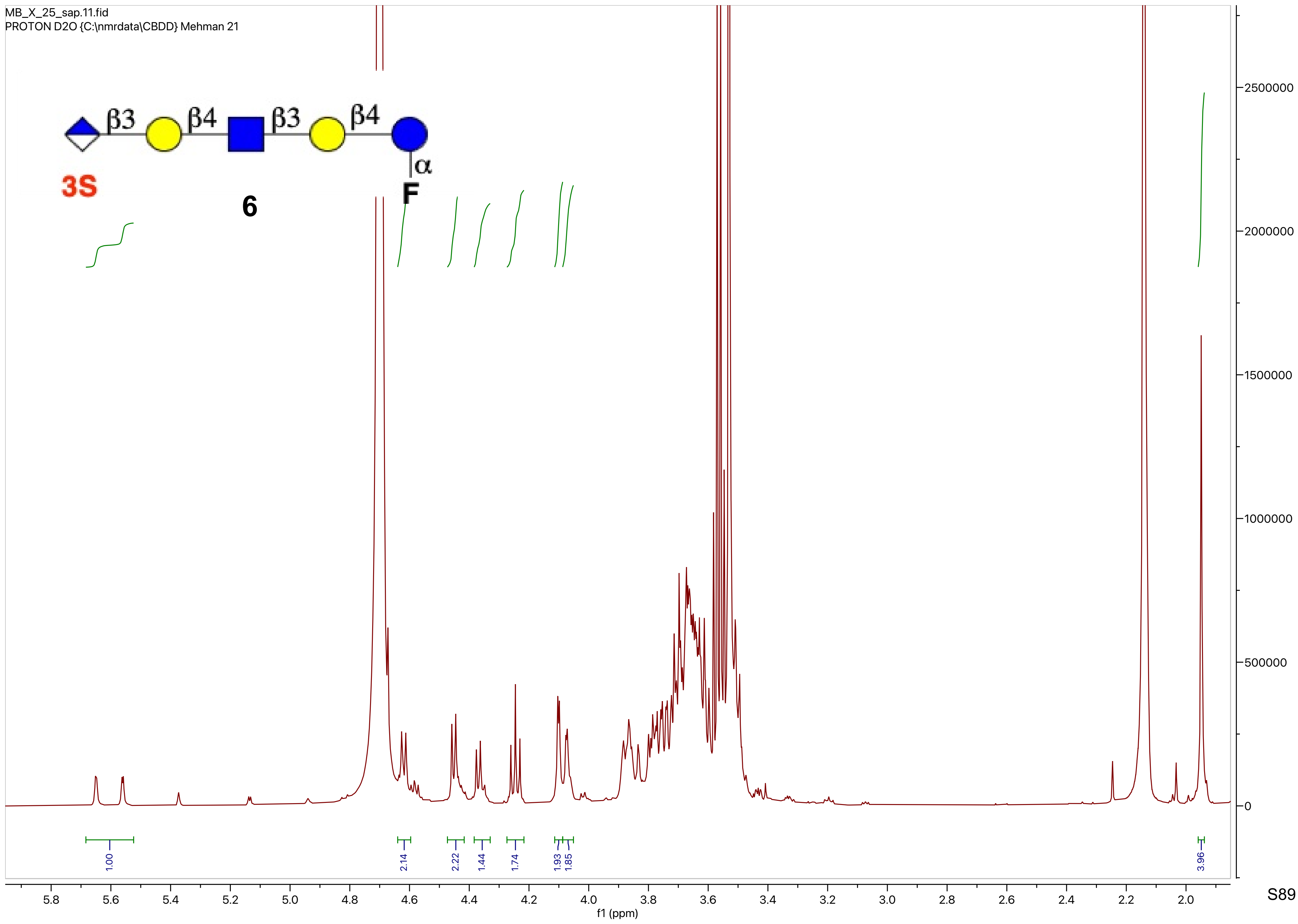

MB_X_25_sap.11.fid
PROTON D2O {C:\nmrdata\CBDD} Mehman 21
β3
β4
β3
β4
α
3S
F
6
2500000
2000000
1500000
1000000
500000
0
1.00
2.14
2.22
1.44
1.74
1.93
1.85
3.96
5.8
5.6
5.4
5.2
5.0
4.8
4.6
4.4
4.2
4.0
3.8
3.6
3.4
3.2
3.0
2.8
2.6
2.4
2.2
2.0
f1 (ppm)

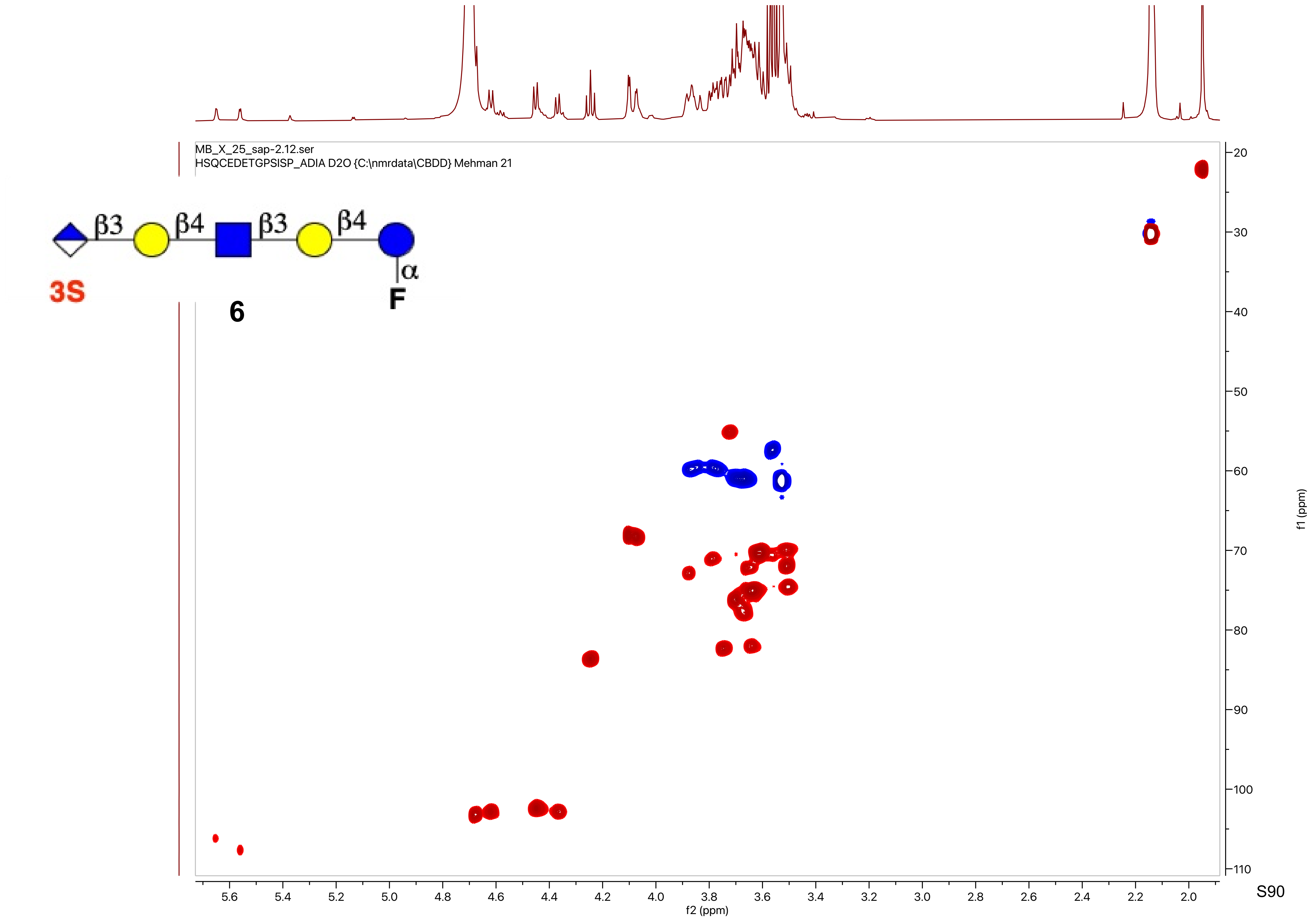
MB_X_25_sap-2.12.ser
HSQCEDETGPSISP_ADIA D2O {C:\nmrdata\CBDD} Mehman 21
β3
β4
β3
β4
α
3S
6
F
f1 (ppm)
f2 (ppm)

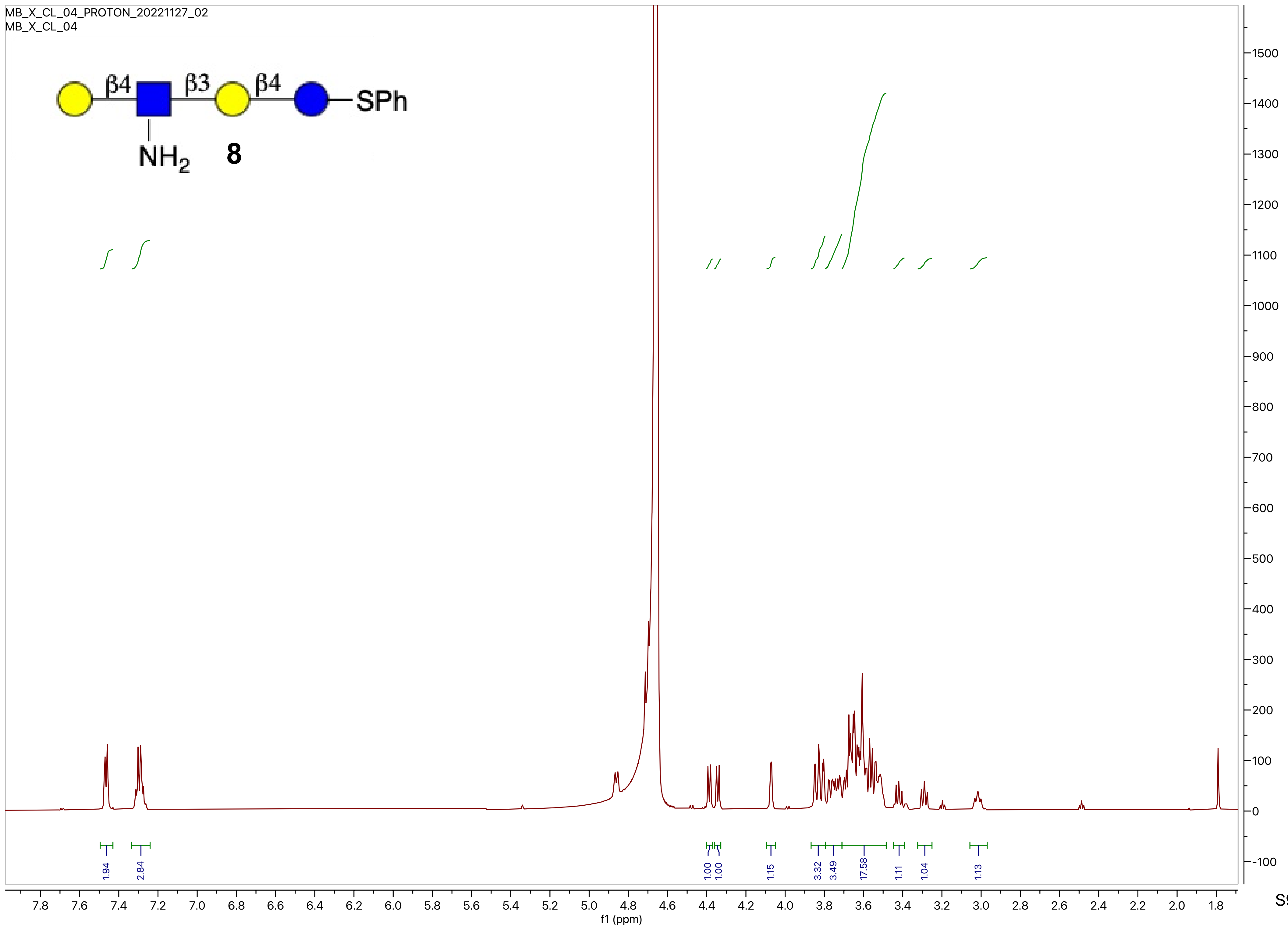
MB_X_CL_04_PROTON_20221127_02
MB_X_CL_04
β4
β3
β4
SPh
NH2
8
1.94
2.84
1.00
1.00
1.15
3.32
3.49
17.58
1.11
1.04
1.13
f1 (ppm)

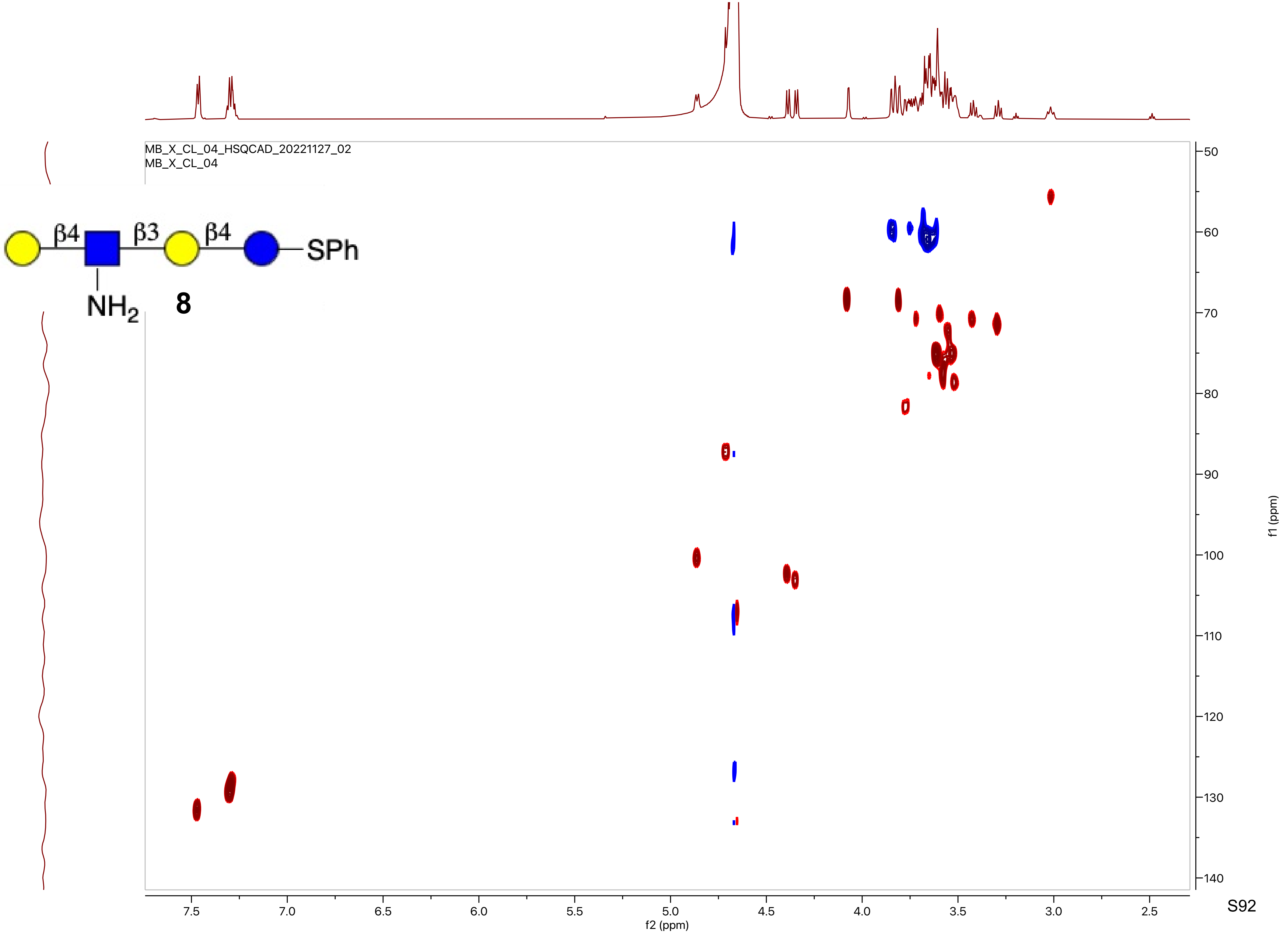
MB_X_CL_04_HSQCAD_20221127_02
MB_X_CL_04
β4
β3
β4
SPh
NH2
8
f2 (ppm)
f1 (ppm)

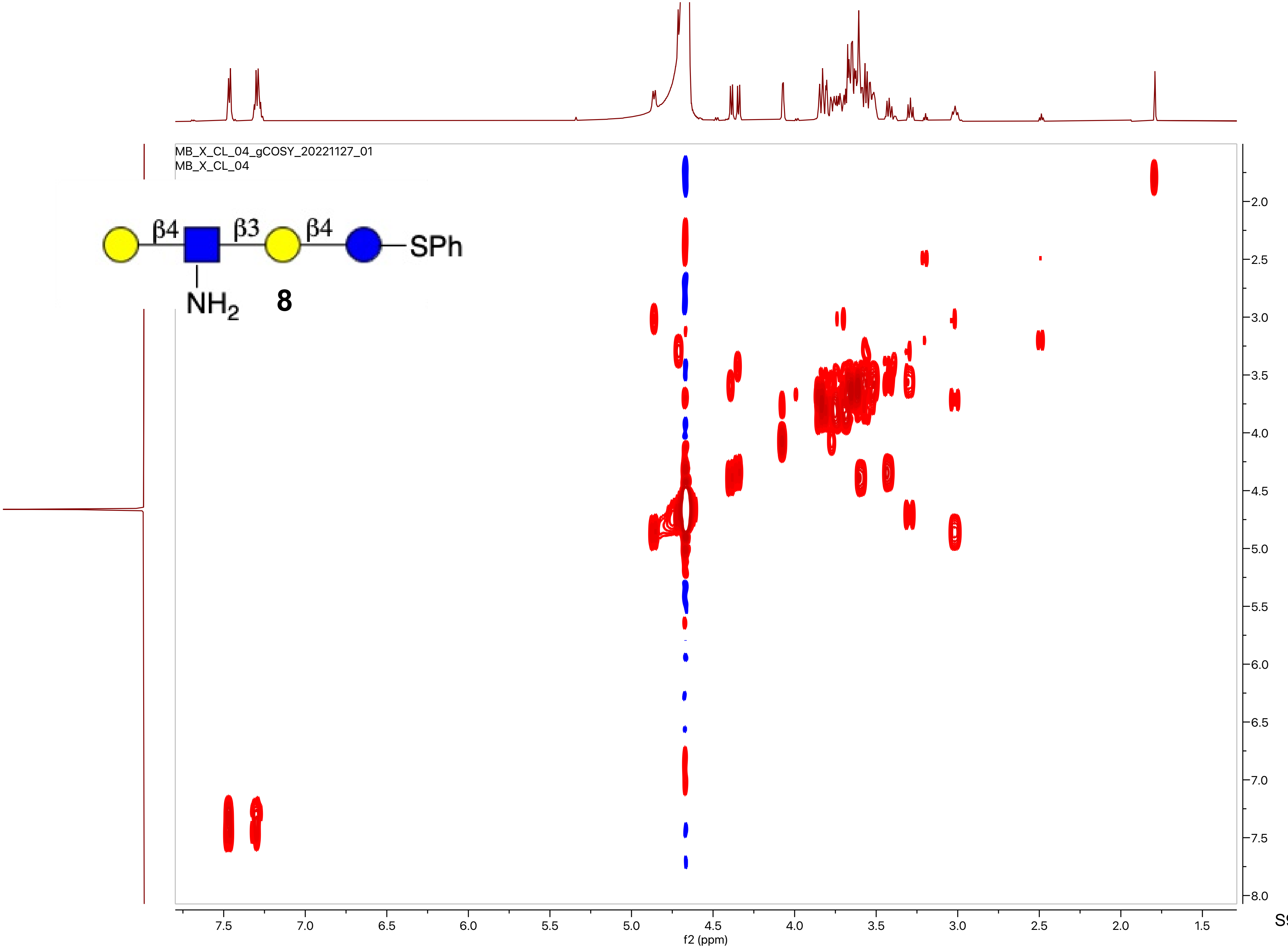
MB_X_CL_04_gCOSY_20221127_01
MB_X_CL_04
β4
β3
β4
SPh
NH2
8
f2 (ppm)
f1 (ppm)

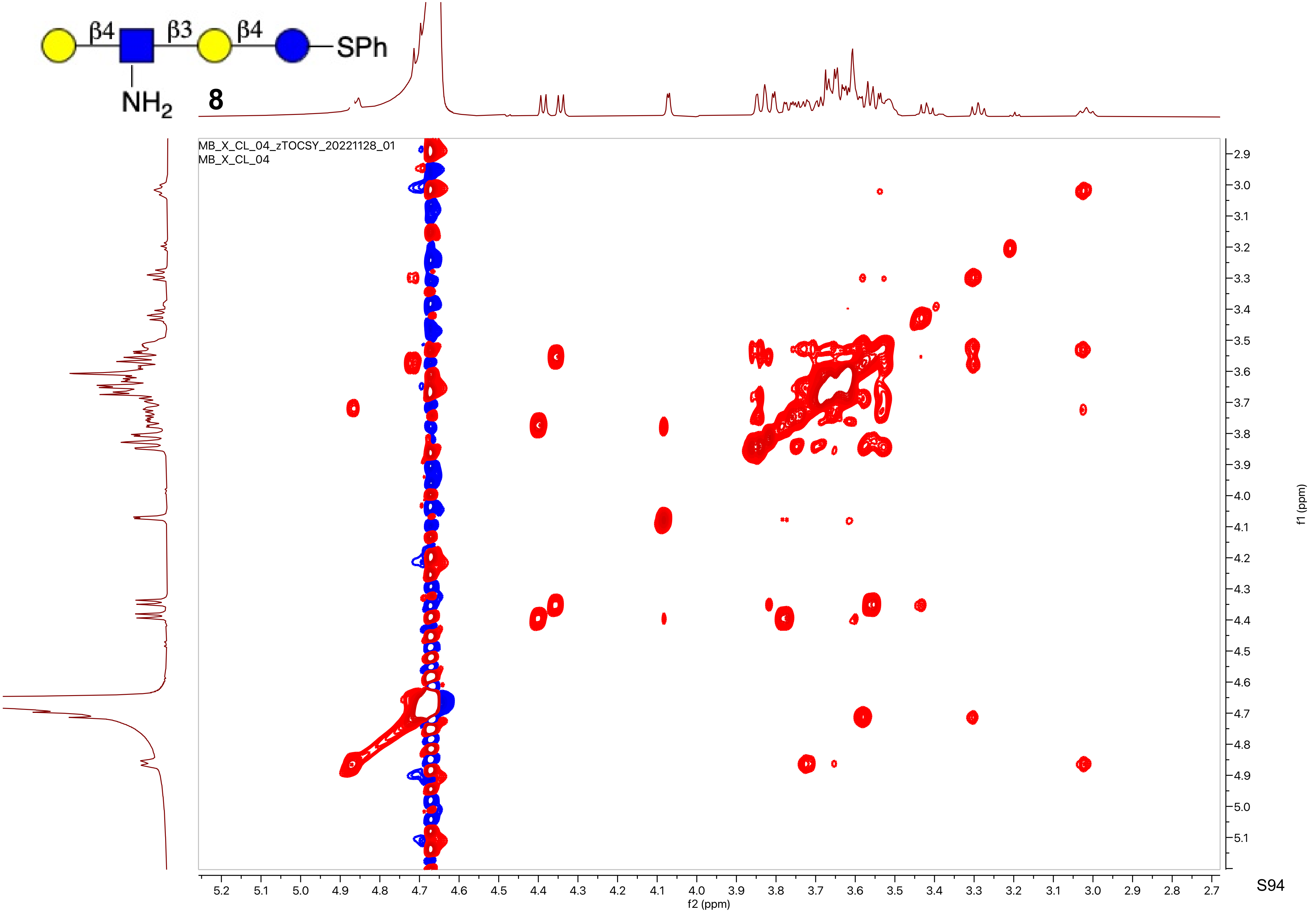
β4
β3
β4
SPh
NH2
8
MB_X_CL_04_zTOCSY_20221128_01
MB_X_CL_04
f1 (ppm)
f2 (ppm)
2.9
3.0
3.1
3.2
3.3
3.4
3.5
3.6
3.7
3.8
3.9
4.0
4.1
4.2
4.3
4.4
4.5
4.6
4.7
4.8
4.9
5.0
5.1
5.2
2.8
2.7

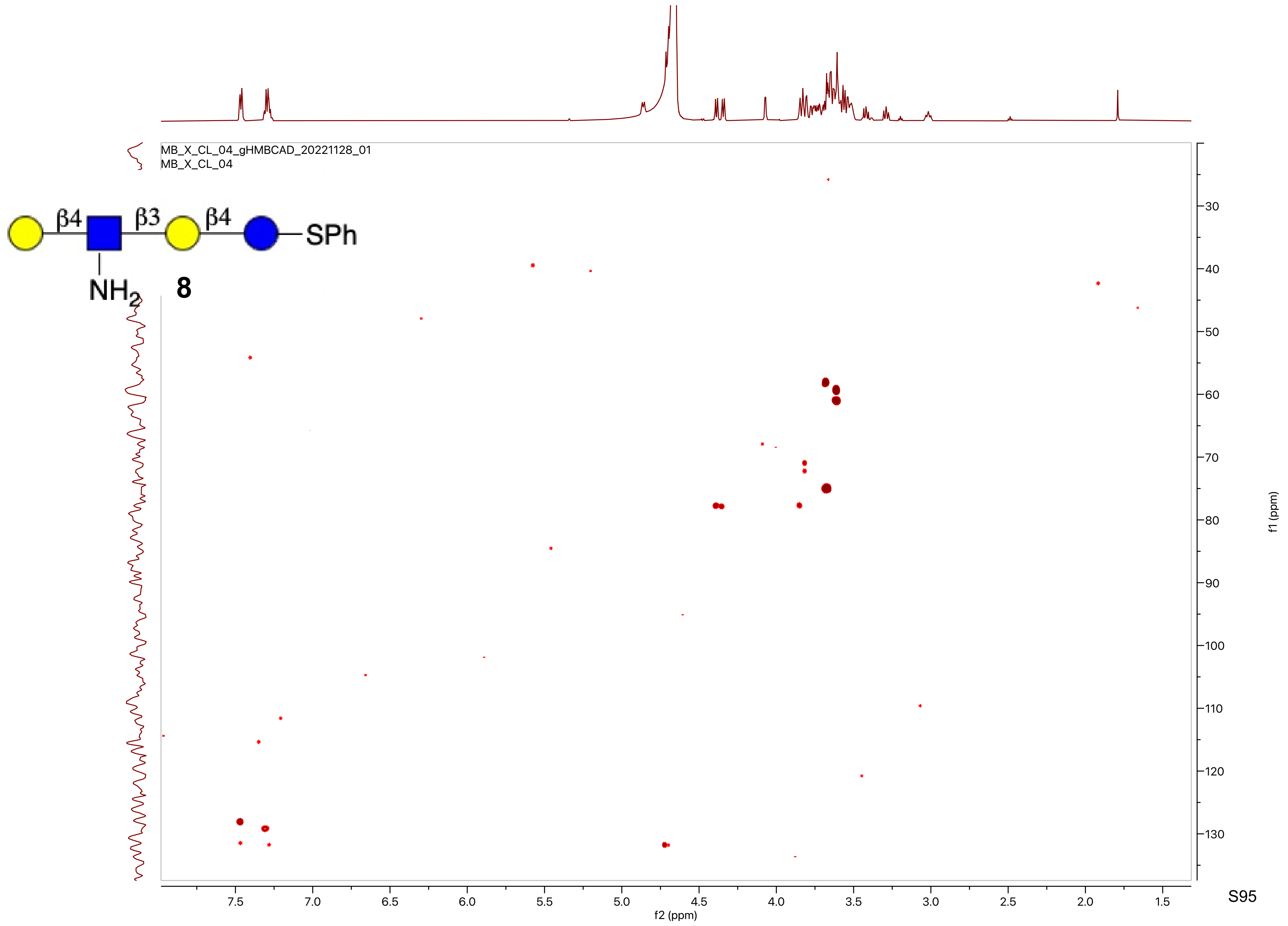
MB_X_CL_04_gHMBCAD_20221128_01
MB_X_CL_04
β4
β3
β4
SPh
NH2
8
f2 (ppm)
f1 (ppm)

MB_X_37.10.fid
PROTON CDCl3 {C:\nmrdata\CBDD} Mehman 52

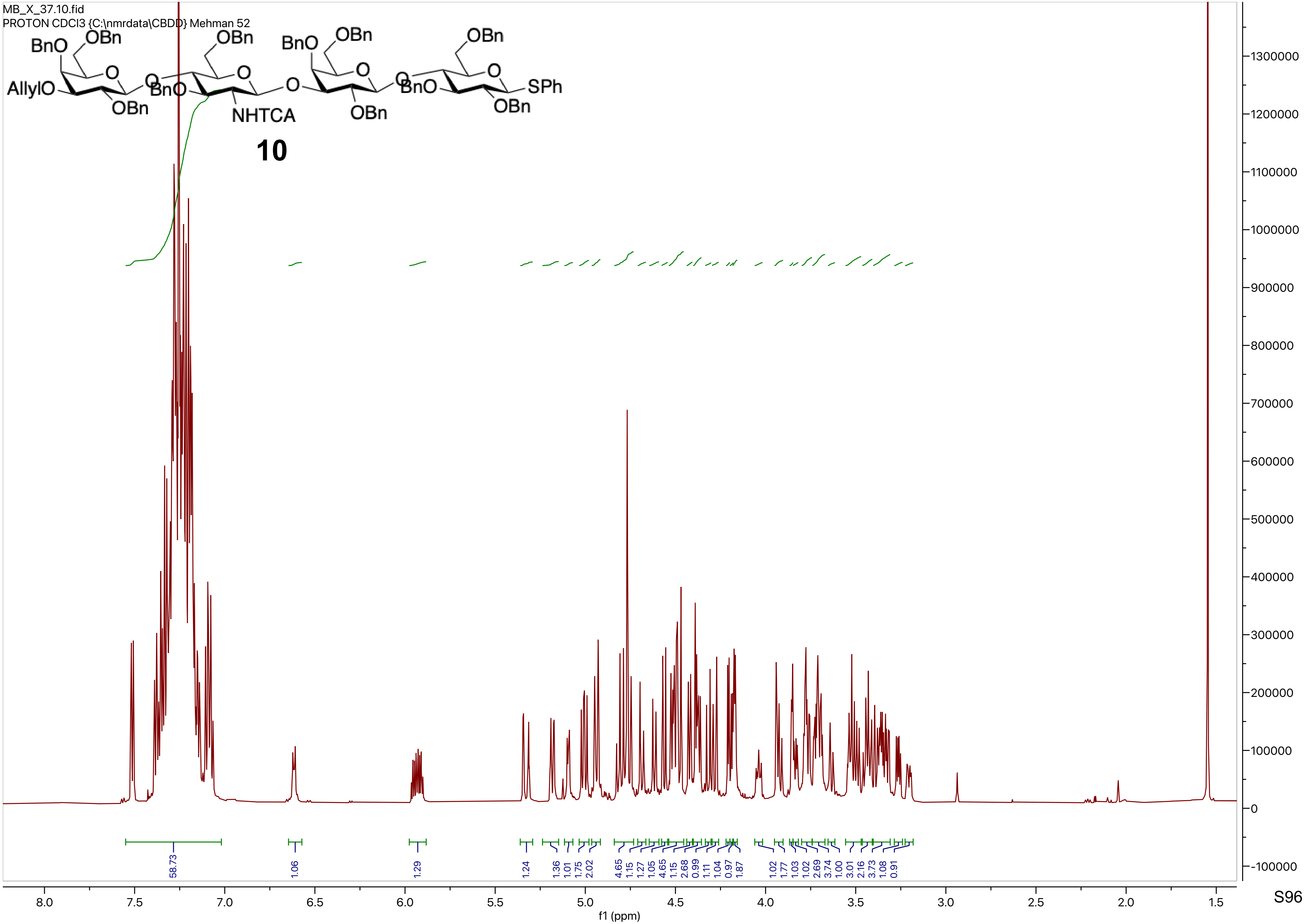

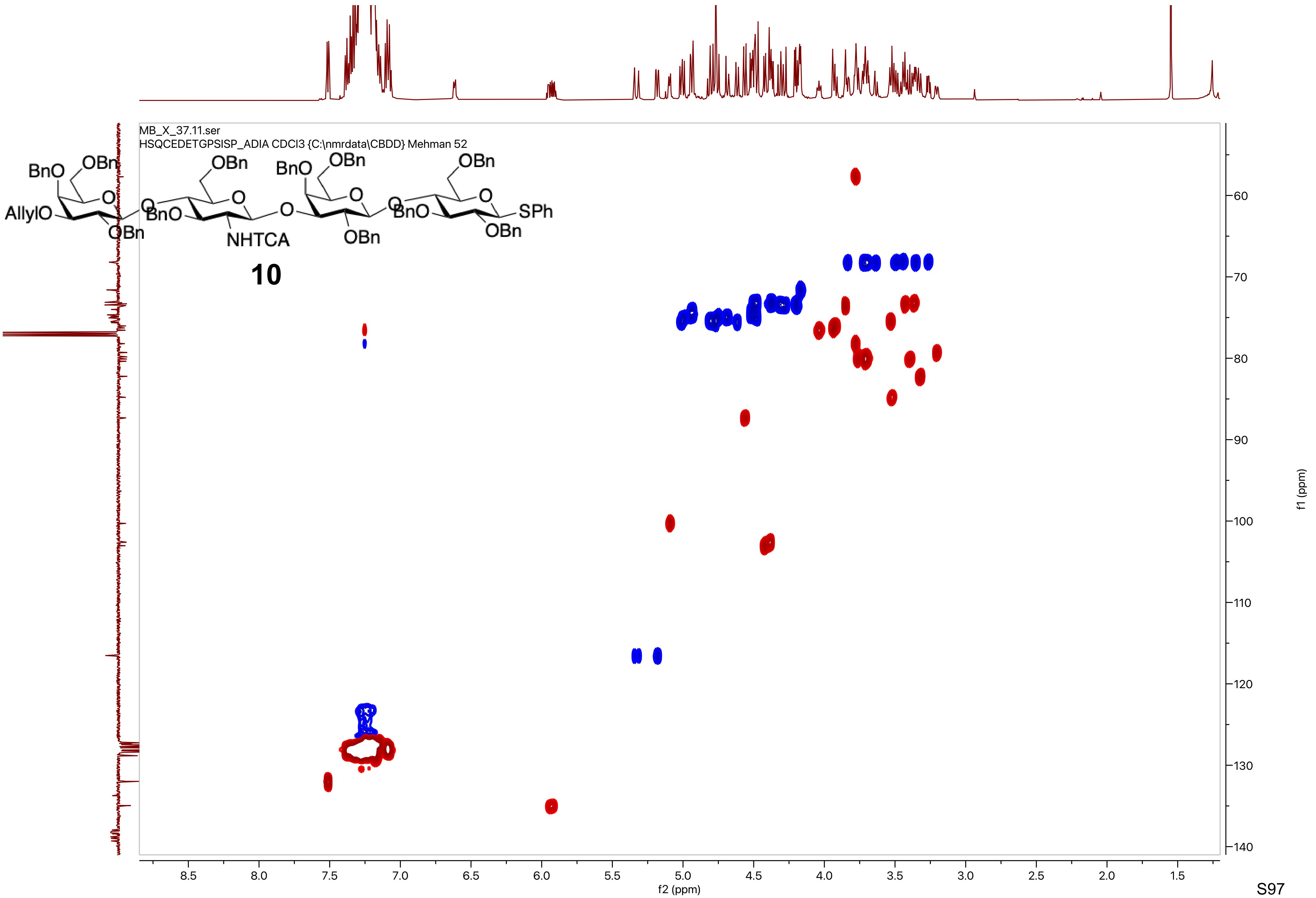
MB_X_37.11.ser
HSQCEDETGPSISP_ADIA CDCl3 {C:\nmrdata\CBDD} Mehman 52
BnO
OBn
AllylO
BnO
NHTCA
OBn
SPh
10
f2 (ppm)
f1 (ppm)

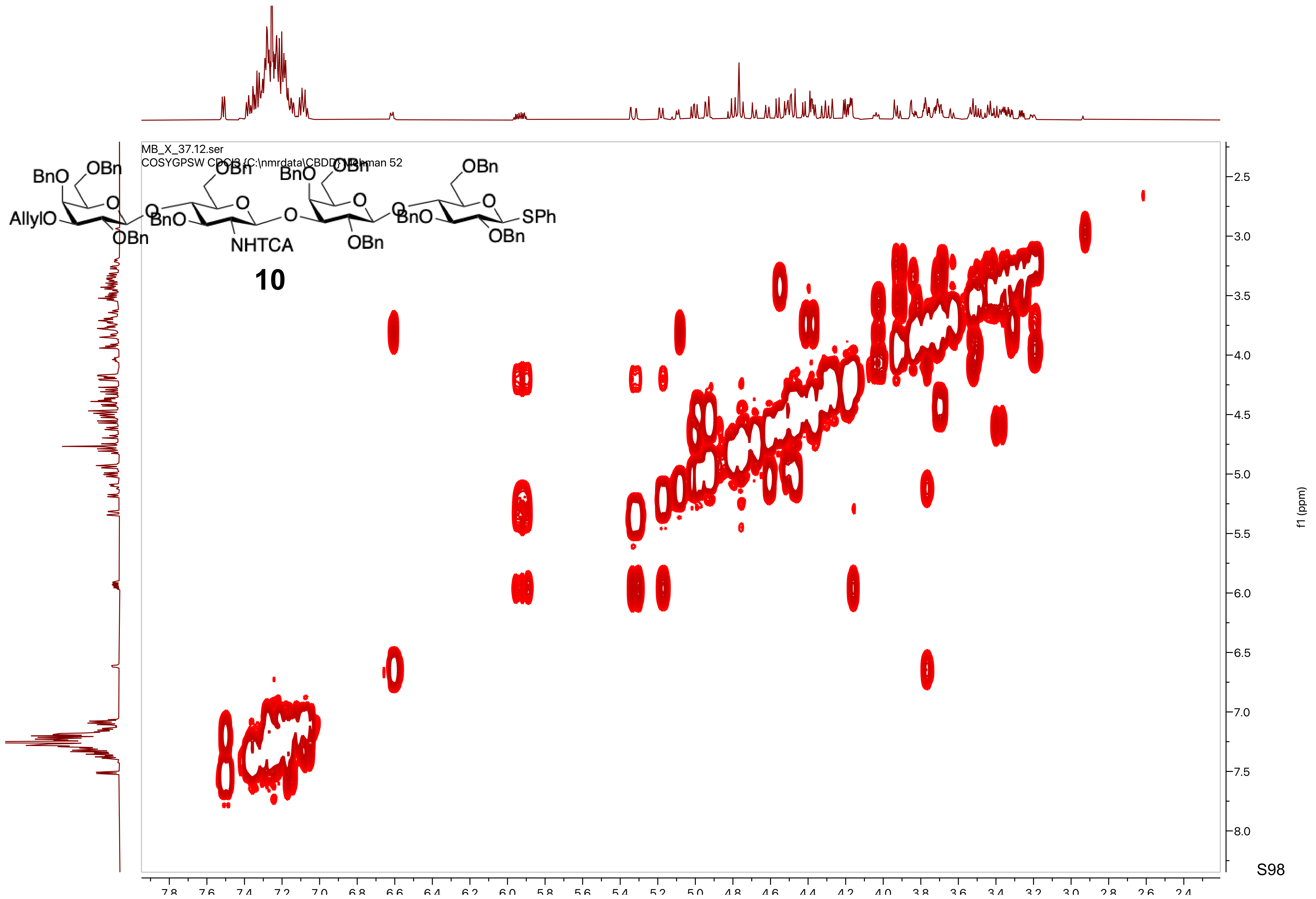
BnO
OBn
AllylO
O
OBn
BnO
NHTCA
OBn
SPh
10
MB_X_37.12.ser
COSYGPSW CDCl3 {C:\nmrdata\CBDD} [illegible] 52
f1 (ppm)
2.5
3.0
3.5
4.0
4.5
5.0
5.5
6.0
6.5
7.0
7.5
8.0

MB_X_37.13.fid
C13APT CDCl3 {C:\nmrdata\CBDD} Mehman 52

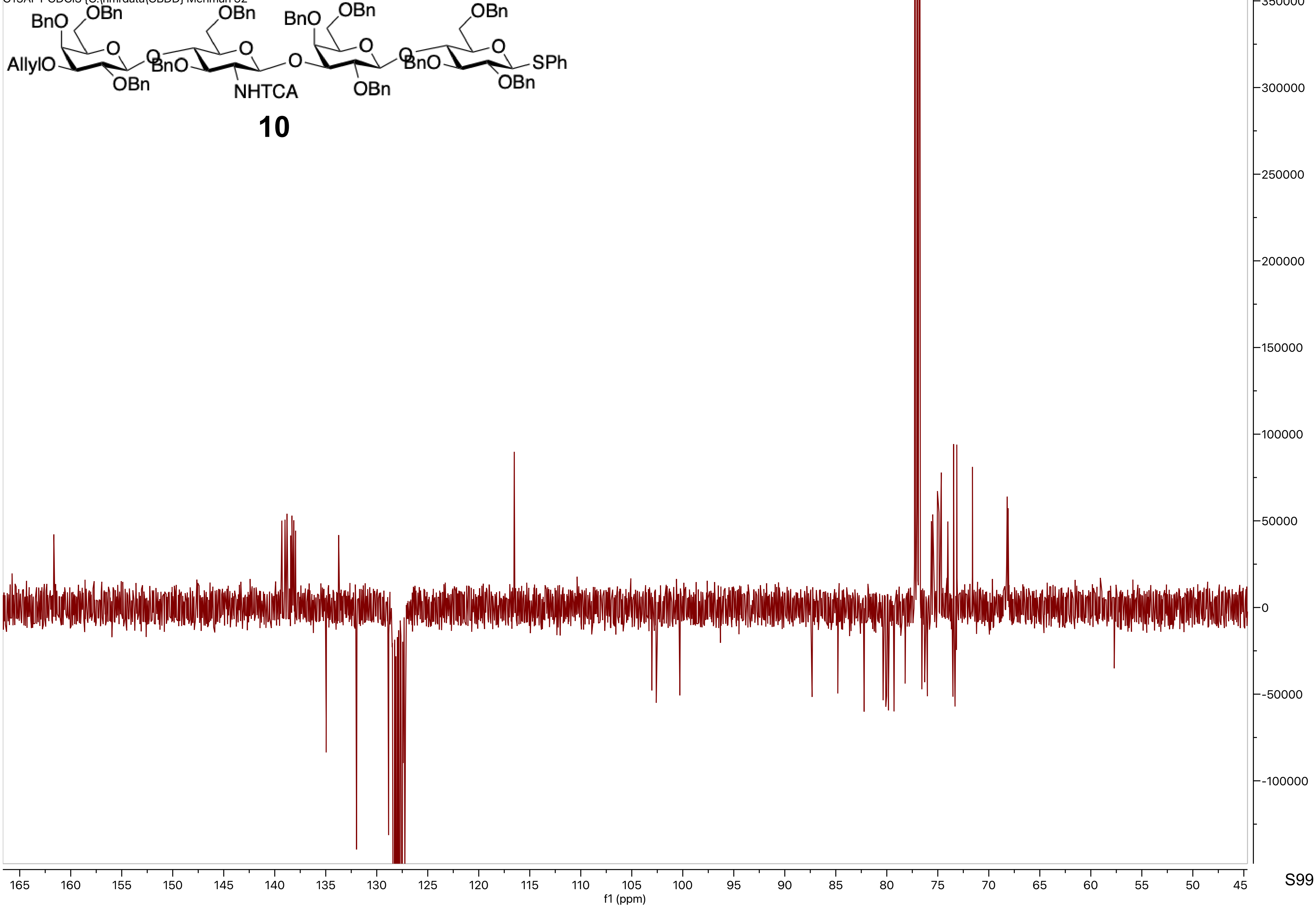

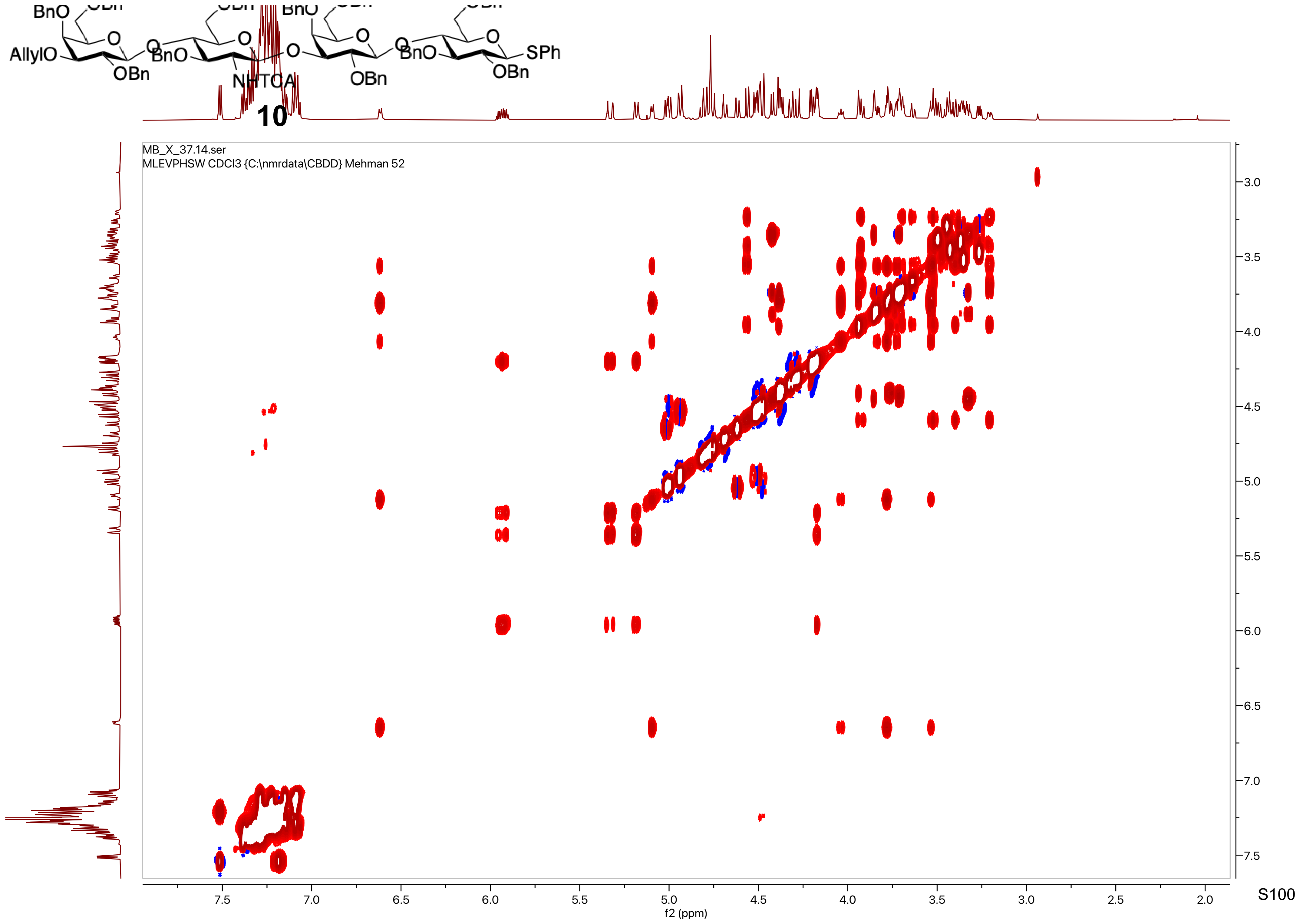

BnO
AllylO
OBn
BnO
NHTCA
O
OBn
BnO
SPh
OBn
10
MB_X_37.14.ser
MLEVPHSW CDCl3 {C:\nmrdata\CBDD} Mehman 52
7.5
7.0
6.5
6.0
5.5
5.0
4.5
4.0
3.5
3.0
2.5
2.0
f2 (ppm)
3.0
3.5
4.0
4.5
5.0
5.5
6.0
6.5
7.0
7.5
f1 (ppm)

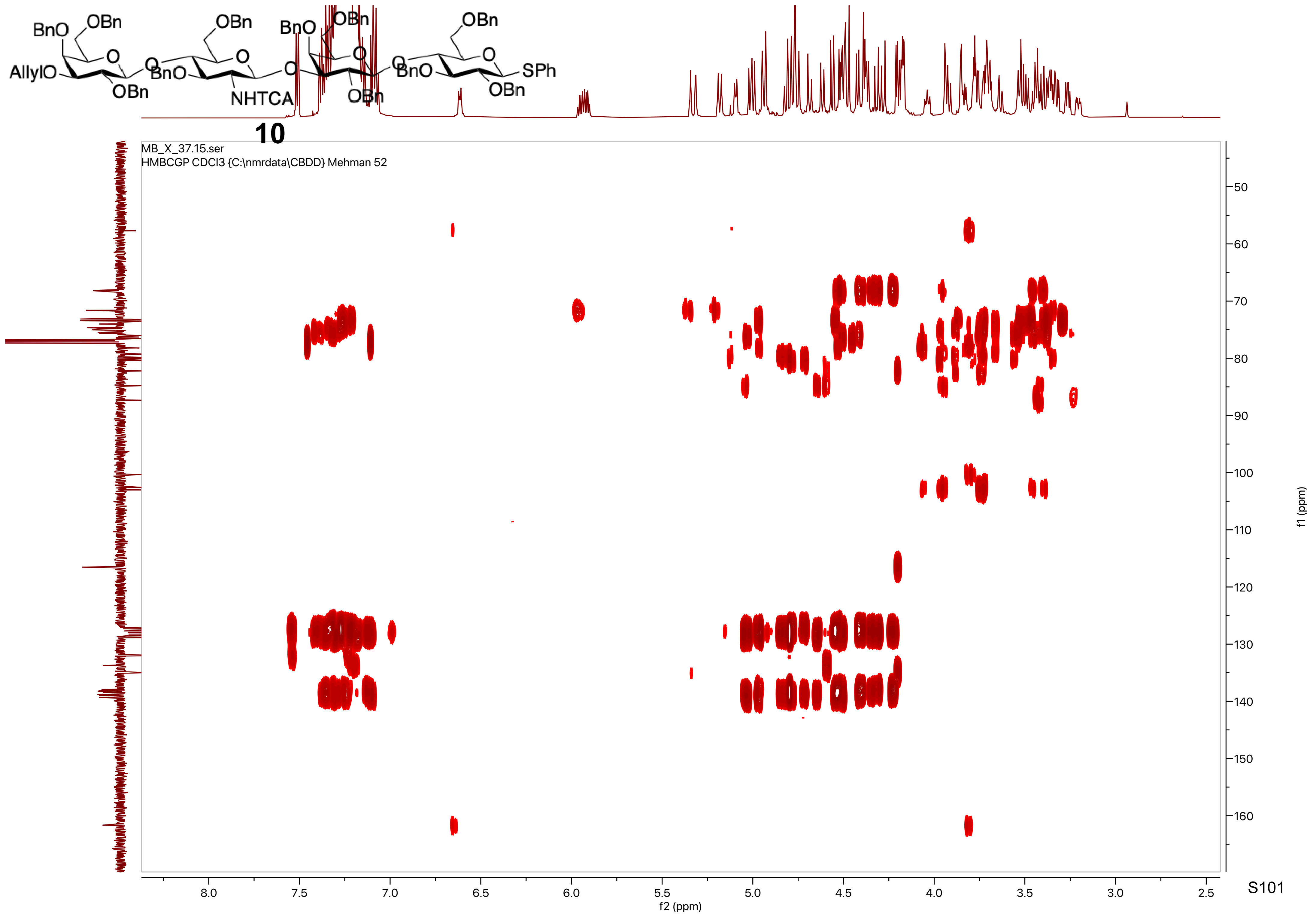

BnO
OBn
AllylO
OBn
BnO
NHTCA
OBn
BnO
OBn
BnO
SPh
OBn
MB_X_37.15.ser
HMBCGP CDCl3 {C:\nmrdata\CBDD} Mehman 52
f2 (ppm)
f1 (ppm)


10

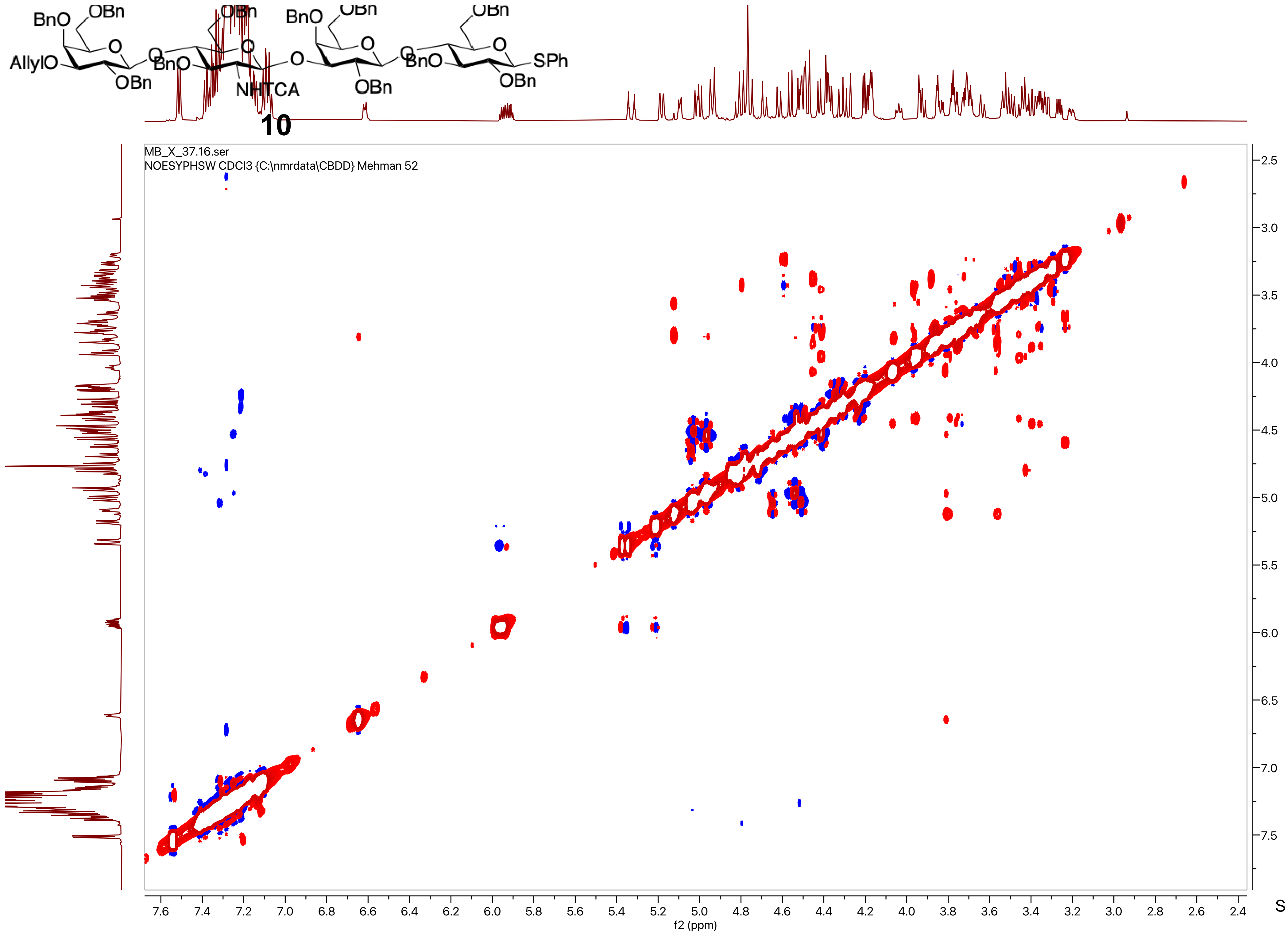
10
MB_X_37.16.ser
NOESYPHSW CDCl3 {C:\nmrdata\CBDD} Mehman 52
f2 (ppm)
f1 (ppm)

MB_VII_94_tst_if.10.fid
PROTON CDCl3 {C:\nmrdata\CBDD} Mehman 11

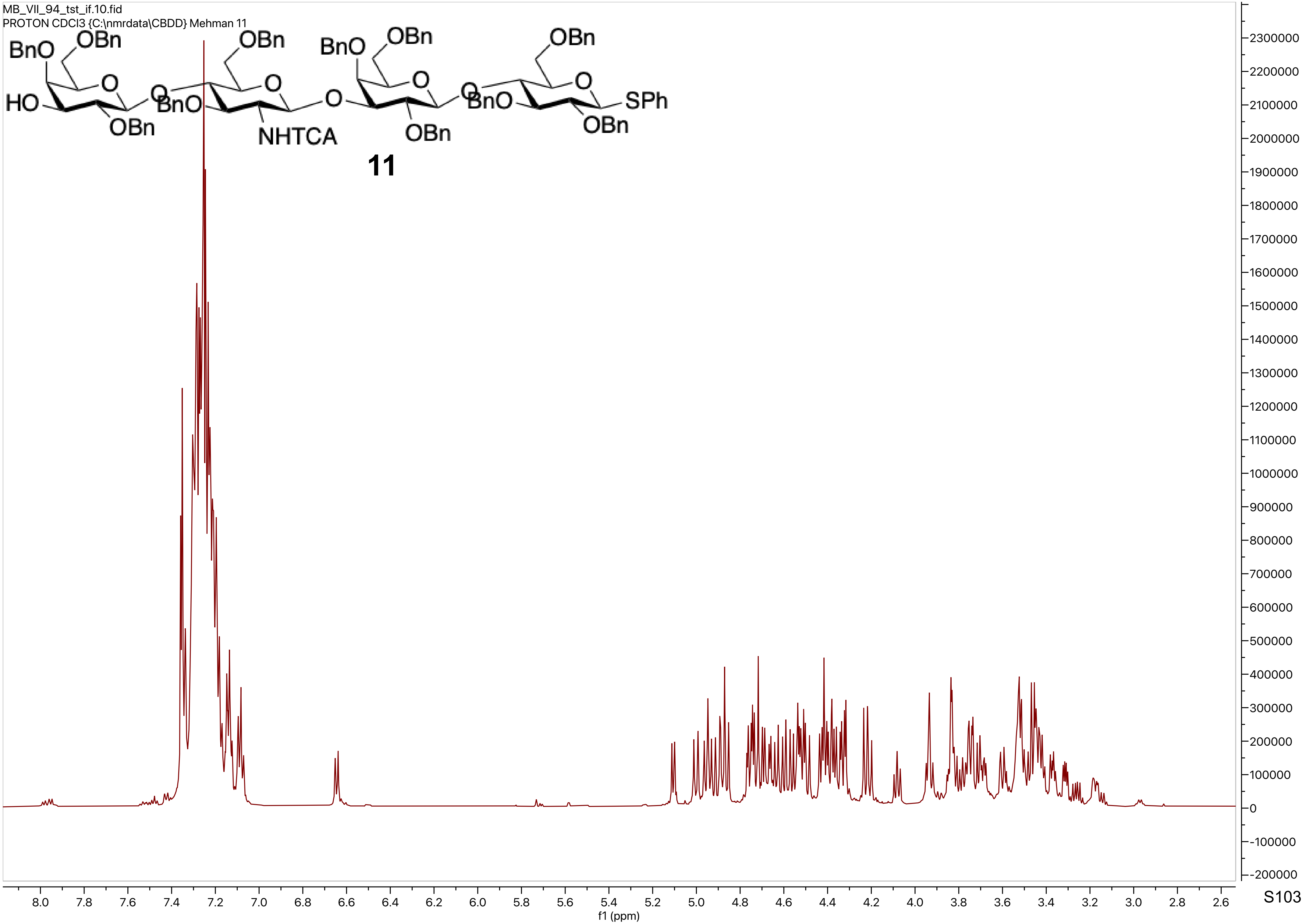

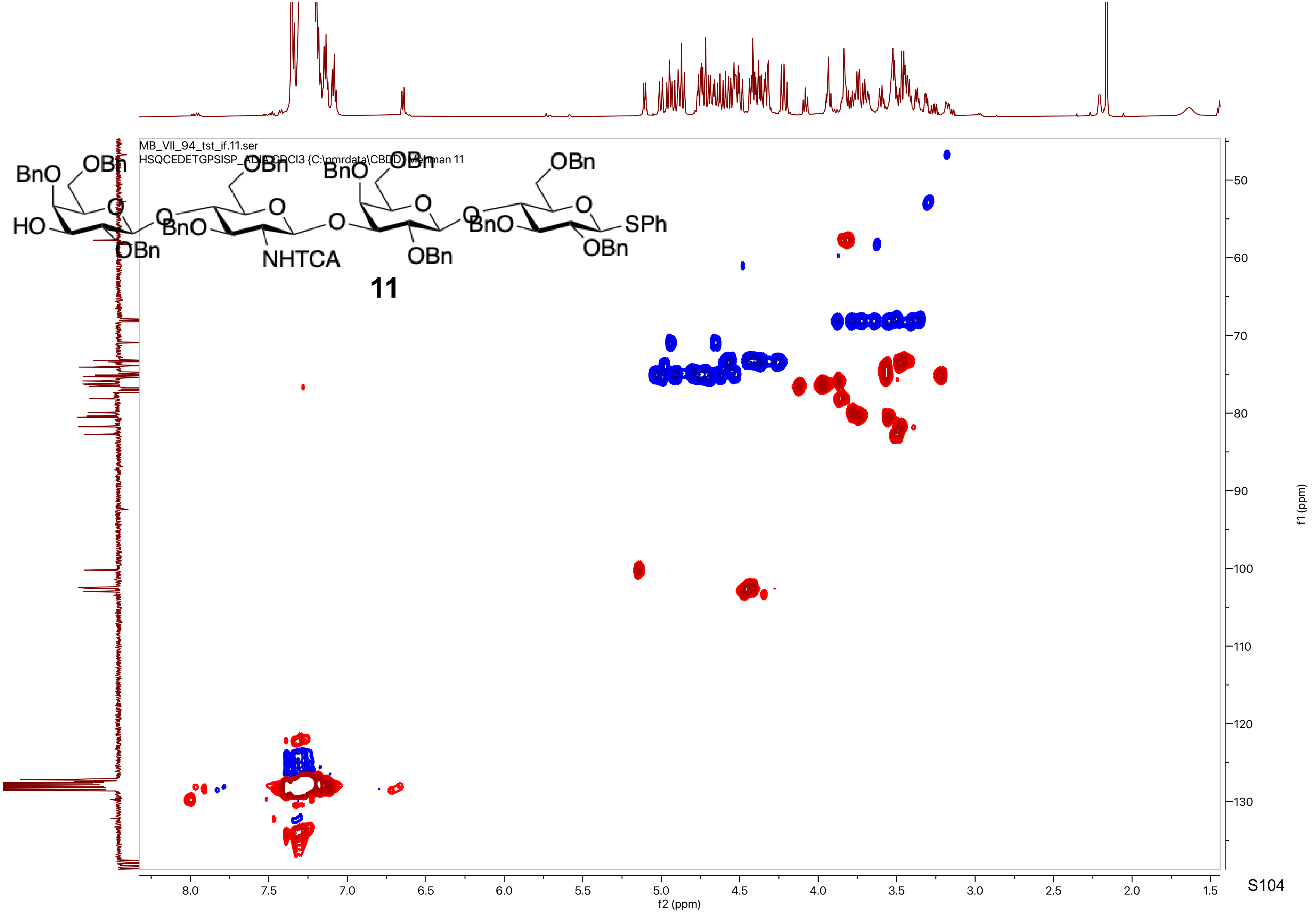

MB_VII_94_tst_if.11.ser
HSQCEDETGPSISP_ADIA CDCl3 {C:\nmrdata\CBDD} Mehman 11
BnO
OBn
HO
OBn
NHTCA
SPh
11
f2 (ppm)
f1 (ppm)

MB_VII_94_tst_if.12.ser
HSQCEDETGPSISP_ADIA_nd CDCl3 {C:\nmrdata\CBDD} Mehman 11

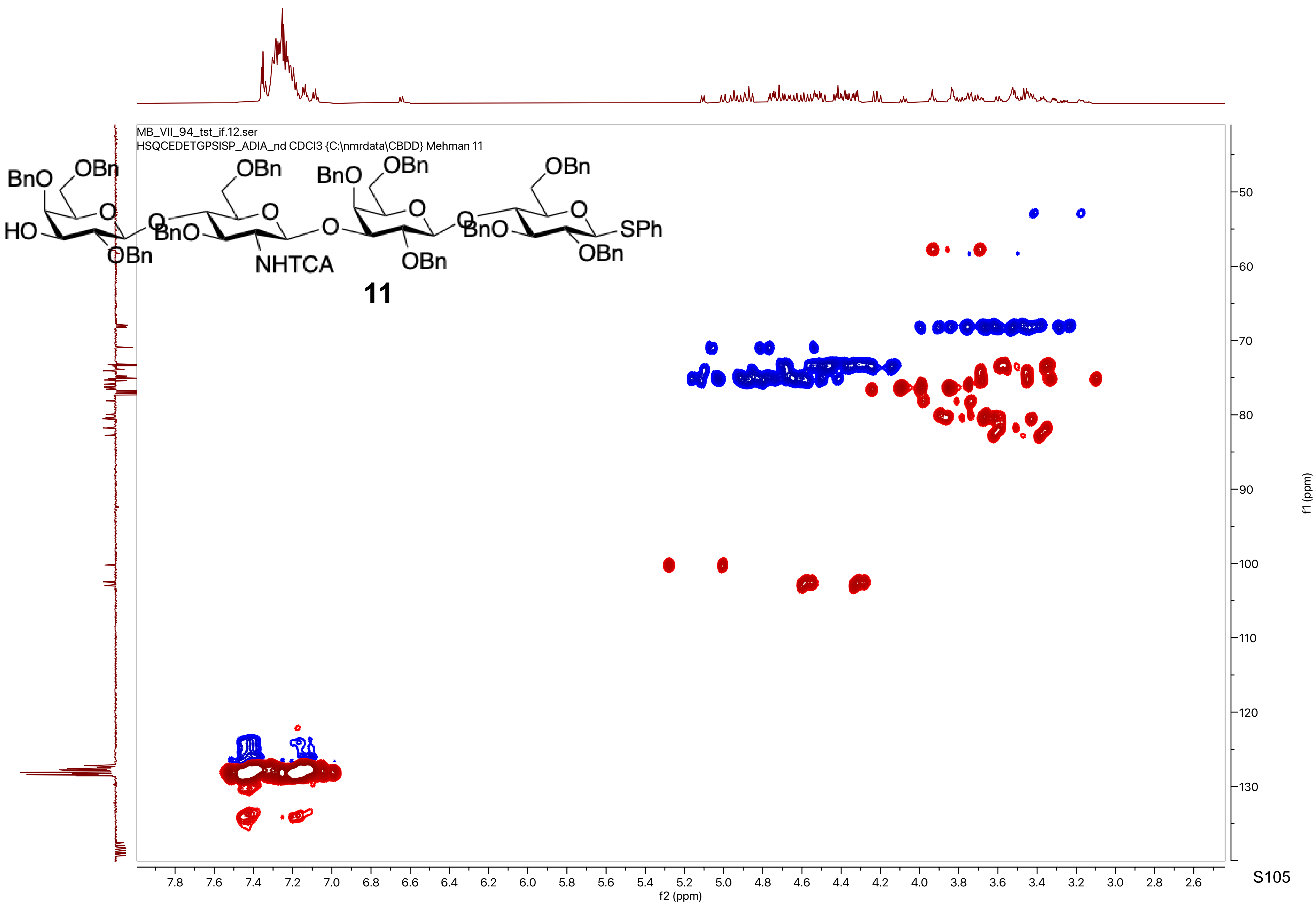


**11**

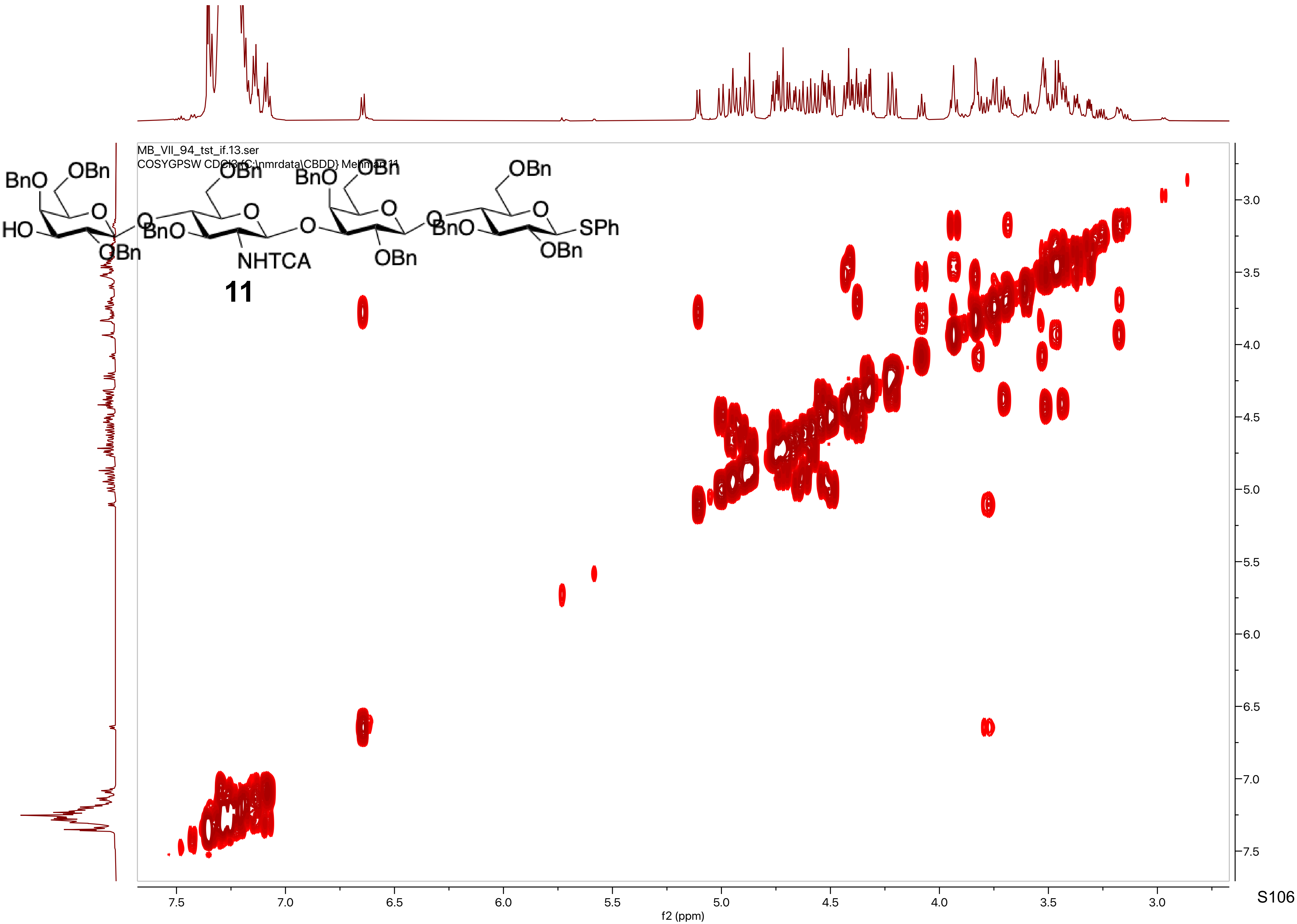

MB_VII_94_tst_if.13.ser
COSYGPSW CDCl3 {C:\nmrdata\CBDD} Mehman 11
BnO
OBn
HO
O
BnO
NHTCA
OBn
SPh
11
f2 (ppm)
f1 (ppm)

MB_VII_94_tst_if.14.fid
C13APT CDCl3 {C:\nmrdata\CBDD} Mehman 11

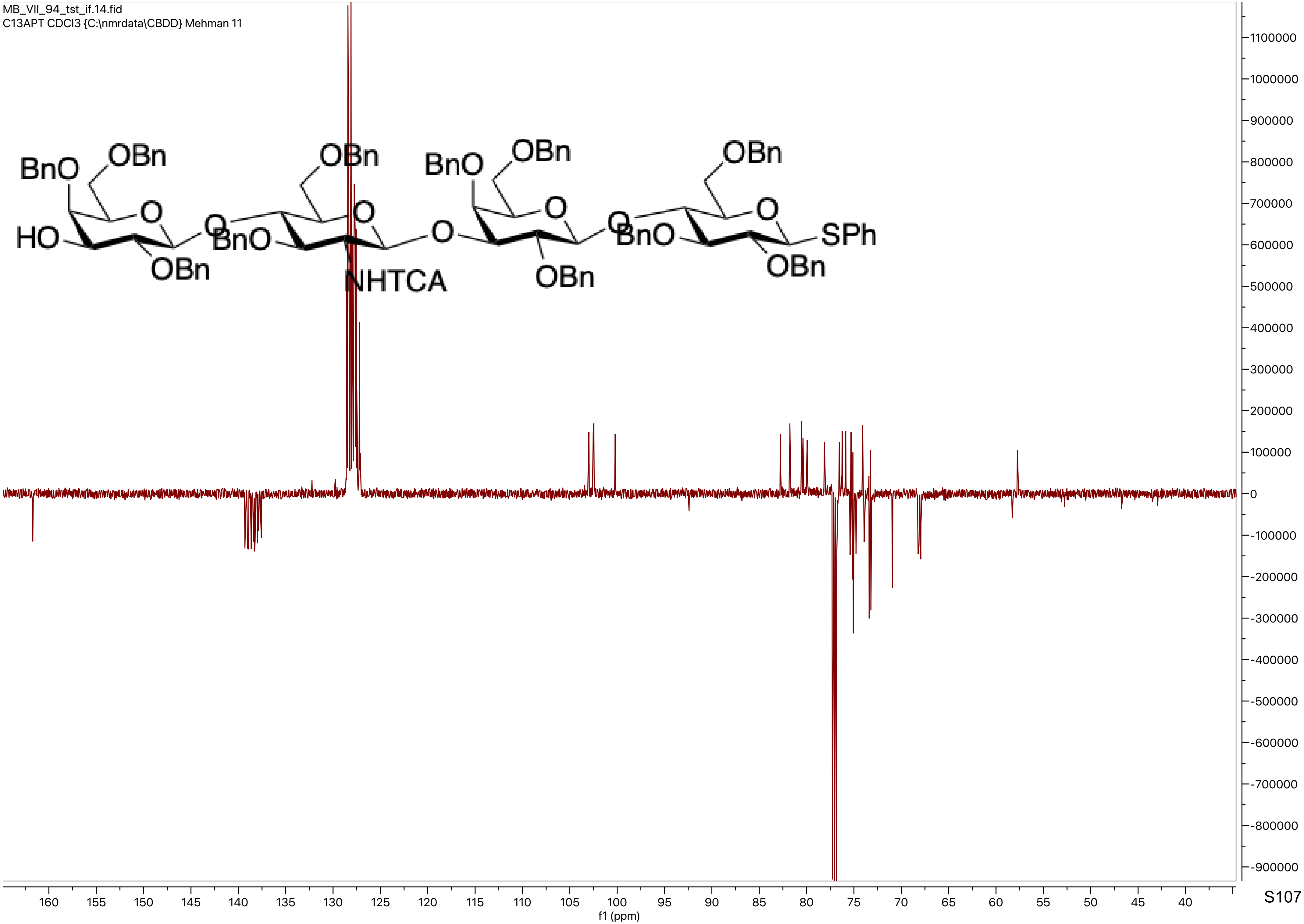

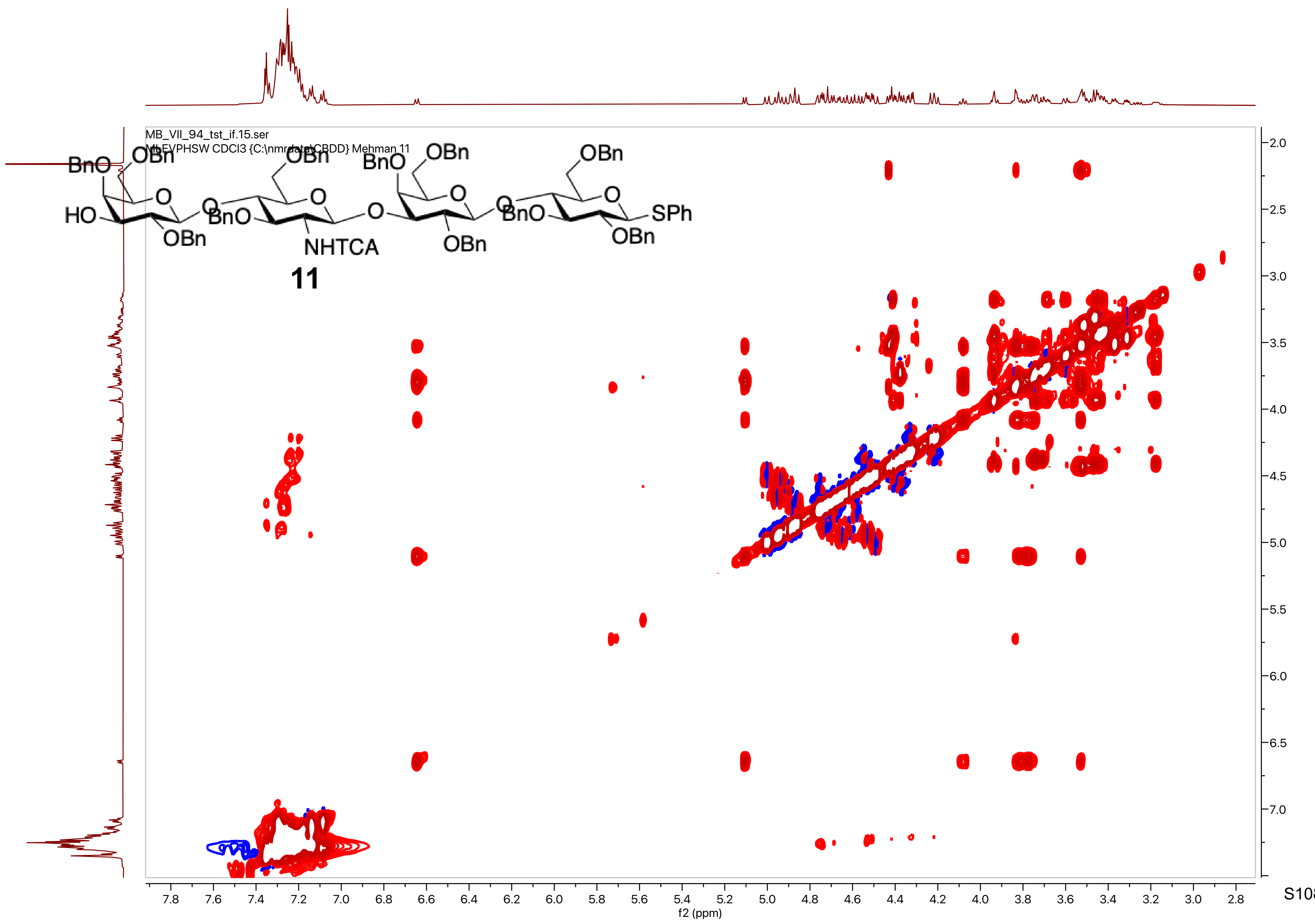

MB_VII_94_tst_if.15.ser
MLEVPHSW CDCl3 {C:\nmrdata\CBDD} Mehman 11
BnO
OBn
HO
OBn
BnO
NHTCA
OBn
SPh
11
f2 (ppm)
f1 (ppm)

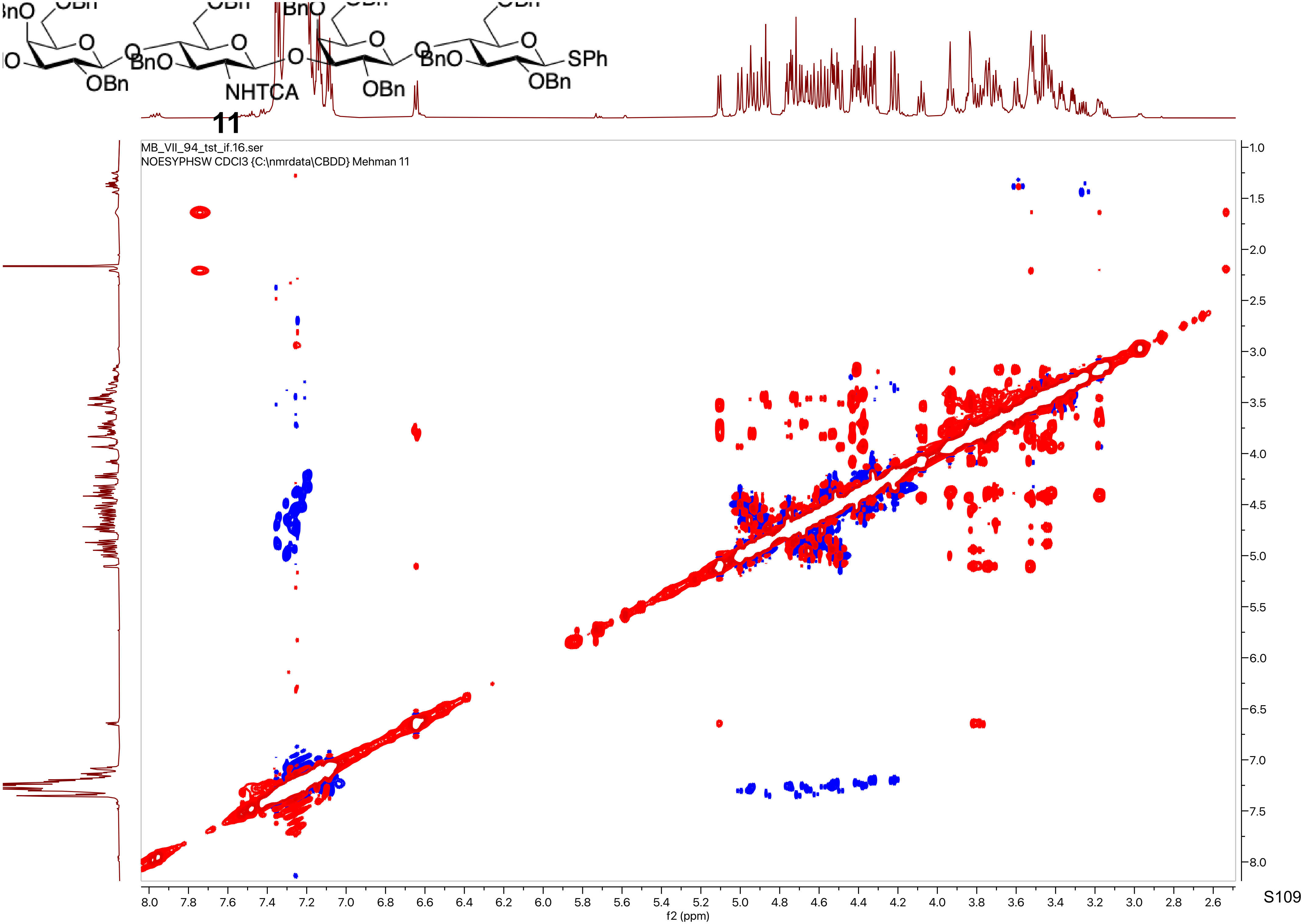
11
MB_VII_94_tst_if.16.ser
NOESYPHSW CDCl3 {C:\nmrdata\CBDD} Mehman 11
f2 (ppm)
f1 (ppm)

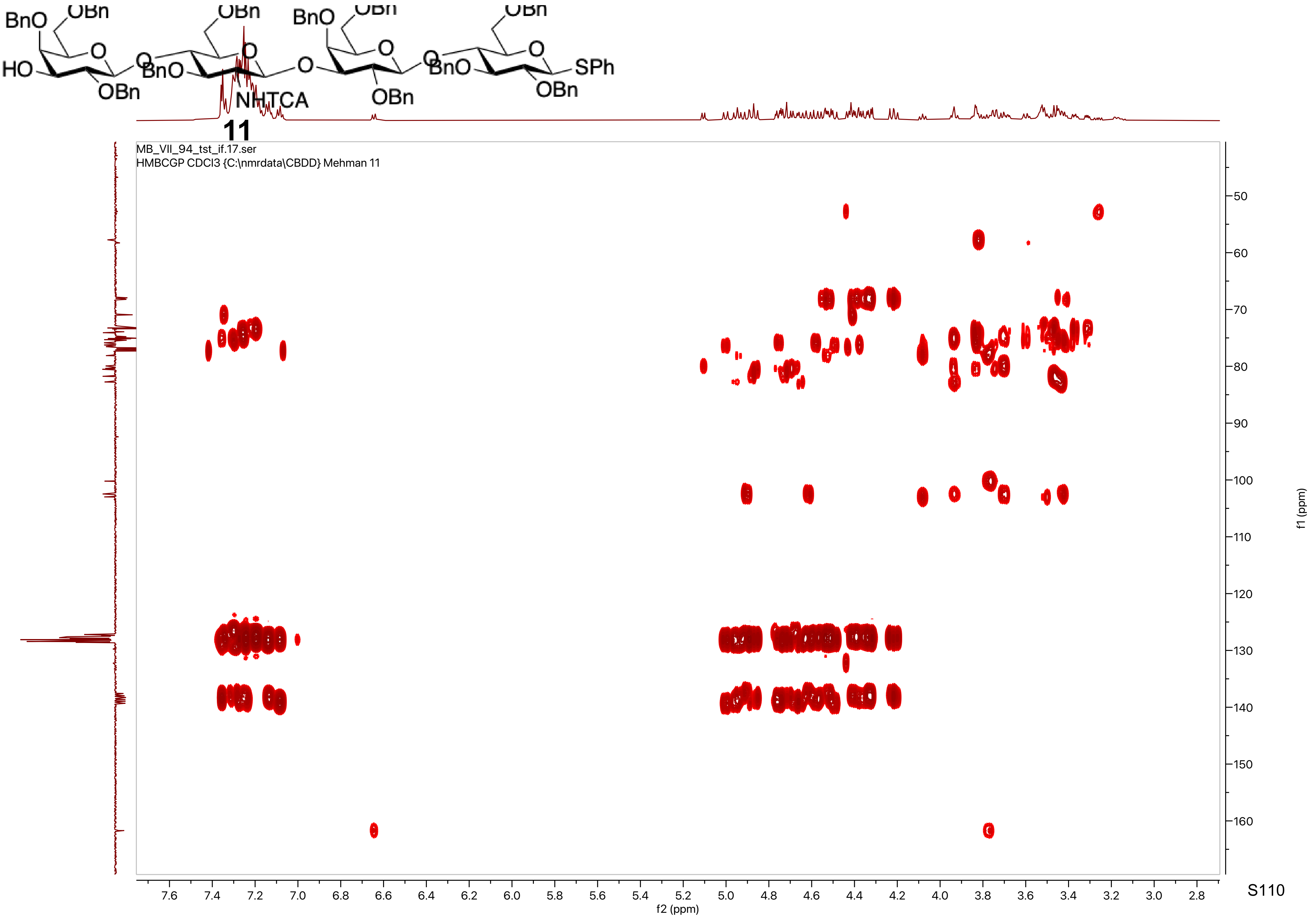

BnO
OBn
HO
O
OBn
BnO
NHTCA
SPh
11
MB_VII_94_tst_if.17.ser
HMBCGP CDCl3 {C:\nmrdata\CBDD} Mehman 11
f2 (ppm)
f1 (ppm)


S110

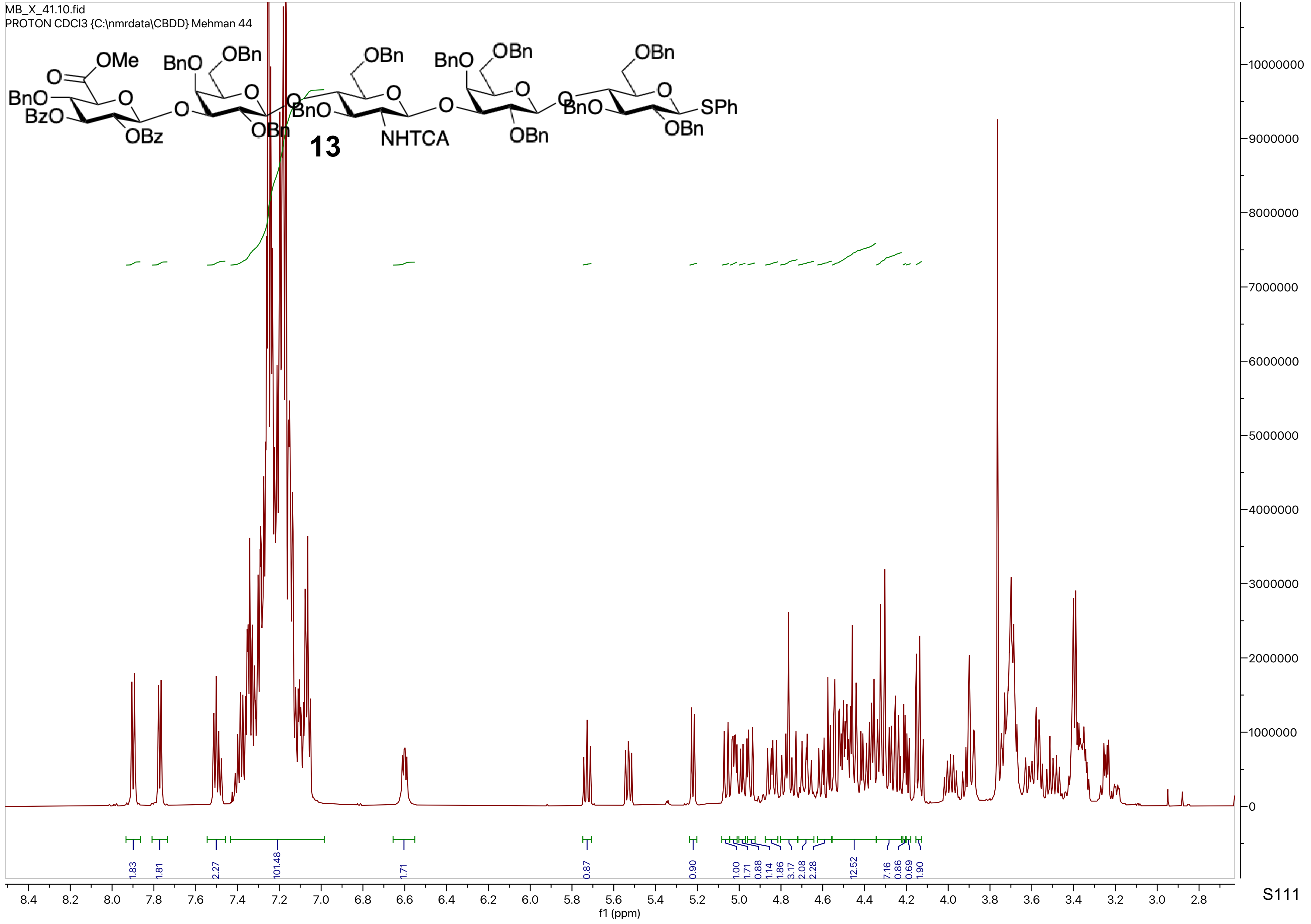
MB_X_41.10.fid
PROTON CDCl3 {C:\nmrdata\CBDD} Mehman 44
13
f1 (ppm)

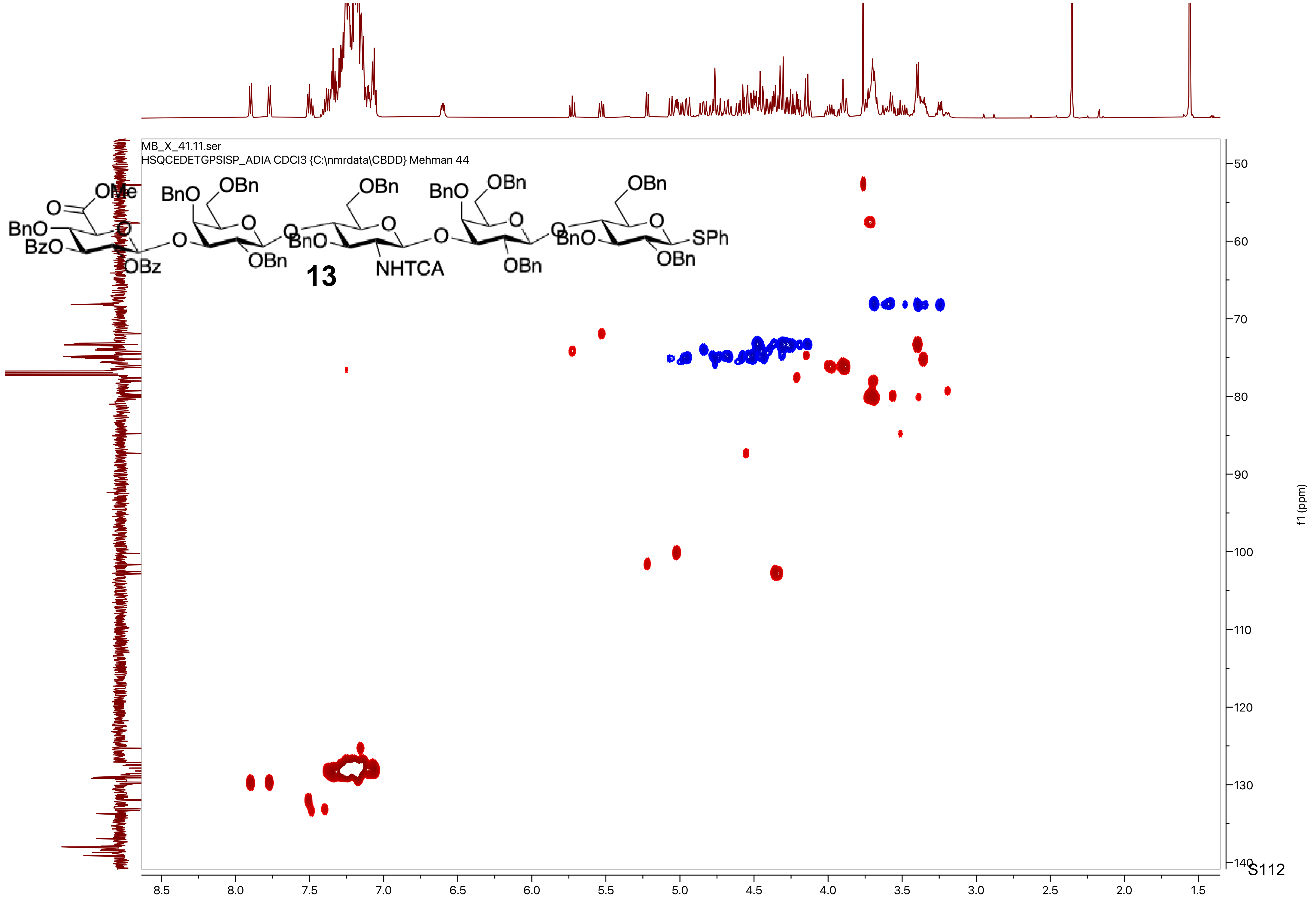
MB_X_41.11.ser
HSQCEDETGPSISP_ADIA CDCl3 {C:\nmrdata\CBDD} Mehman 44
13
f1 (ppm)
8.5
8.0
7.5
7.0
6.5
6.0
5.5
5.0
4.5
4.0
3.5
3.0
2.5
2.0
1.5
50
60
70
80
90
100
110
120
130
140

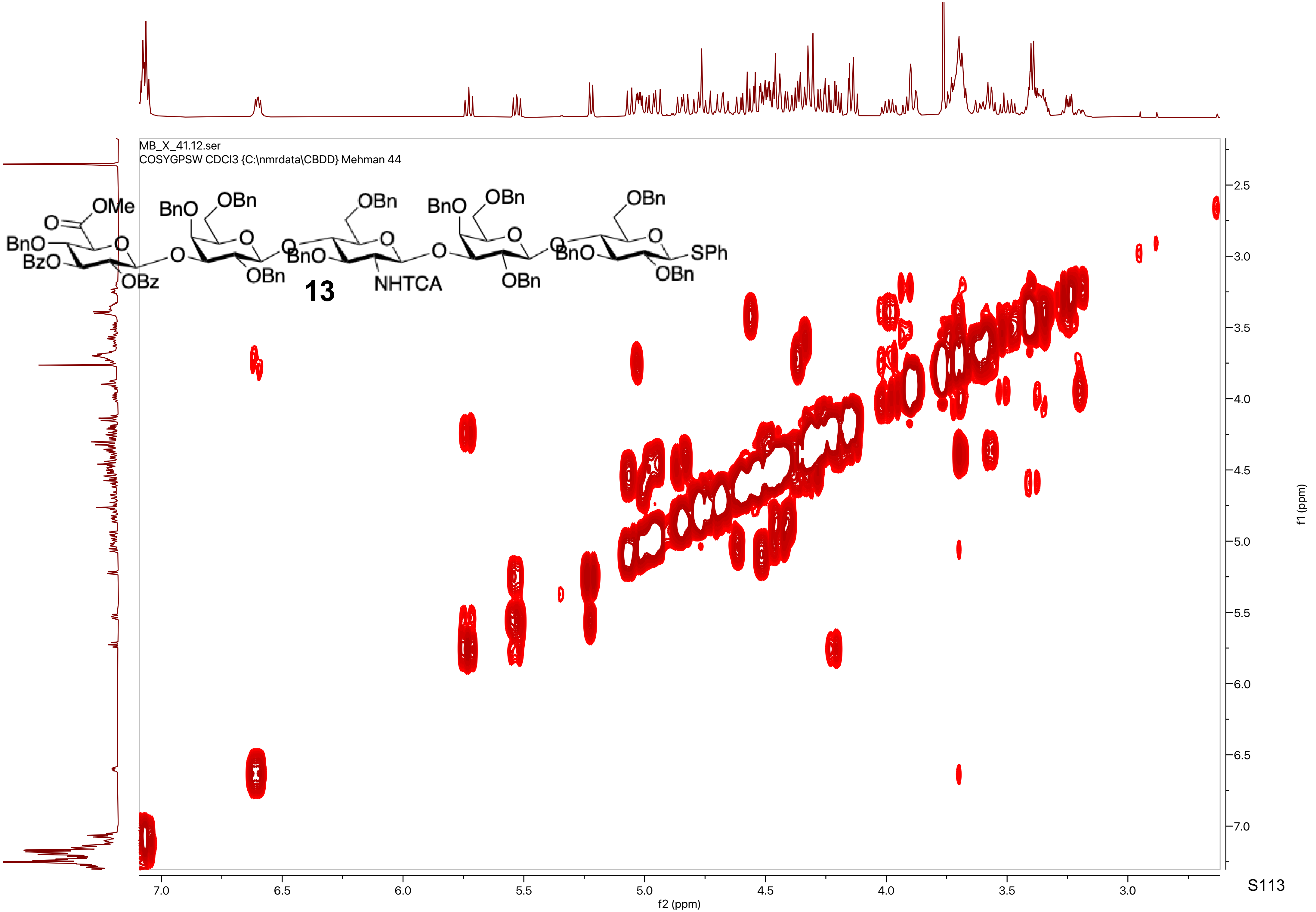
MB_X_41.12.ser
COSYGPSW CDCl3 {C:\nmrdata\CBDD} Mehman 44
OMe
O
BnO
BzO
OBz
BnO
OBn
OBn
13
OBn
BnO
NHTCA
BnO
OBn
OBn
BnO
OBn
OBn
SPh
f2 (ppm)
f1 (ppm)

MB_X_41.13.fid
C13APT CDCl3 {C:\nmrdata\CBDD} Mehman 44

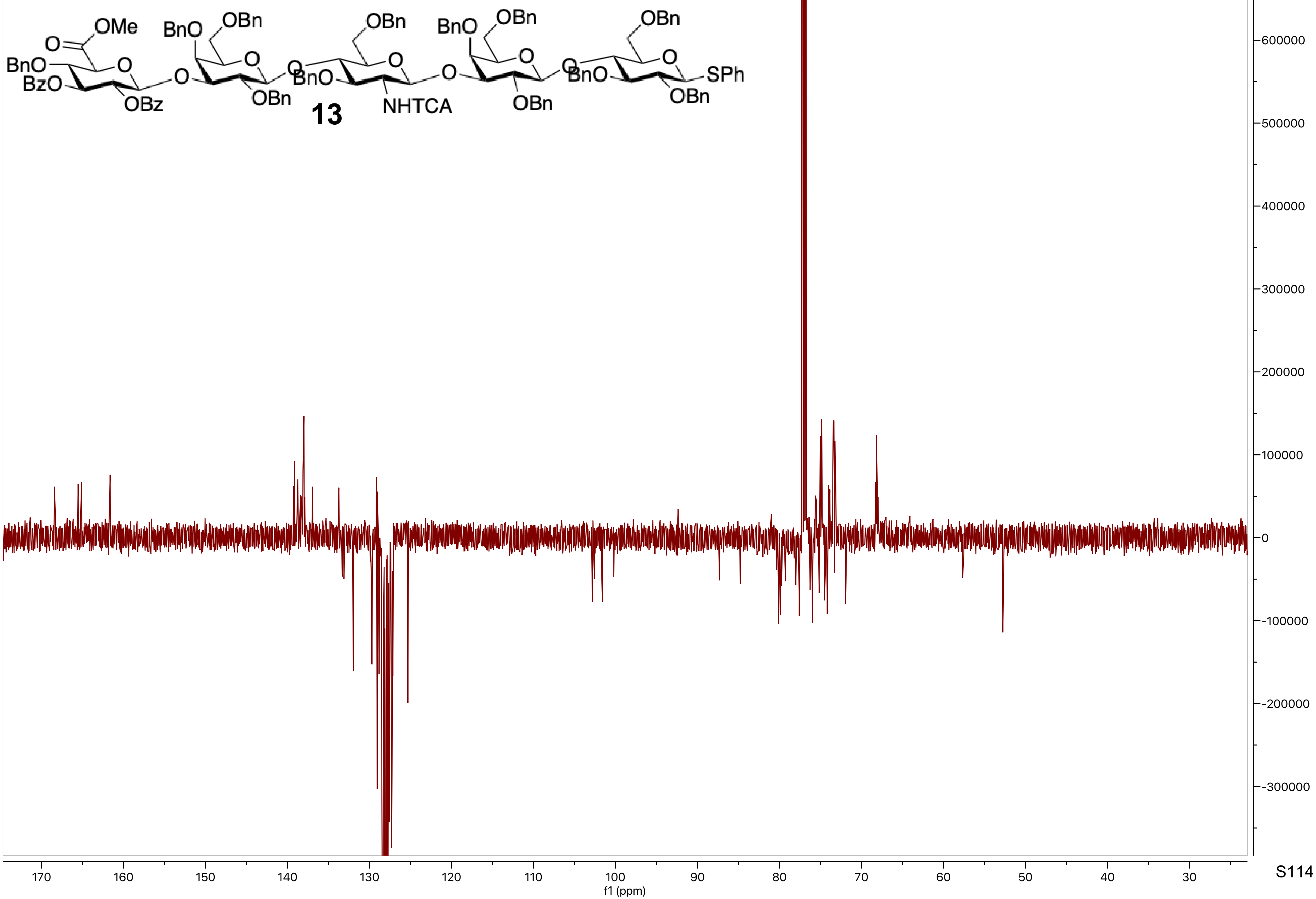

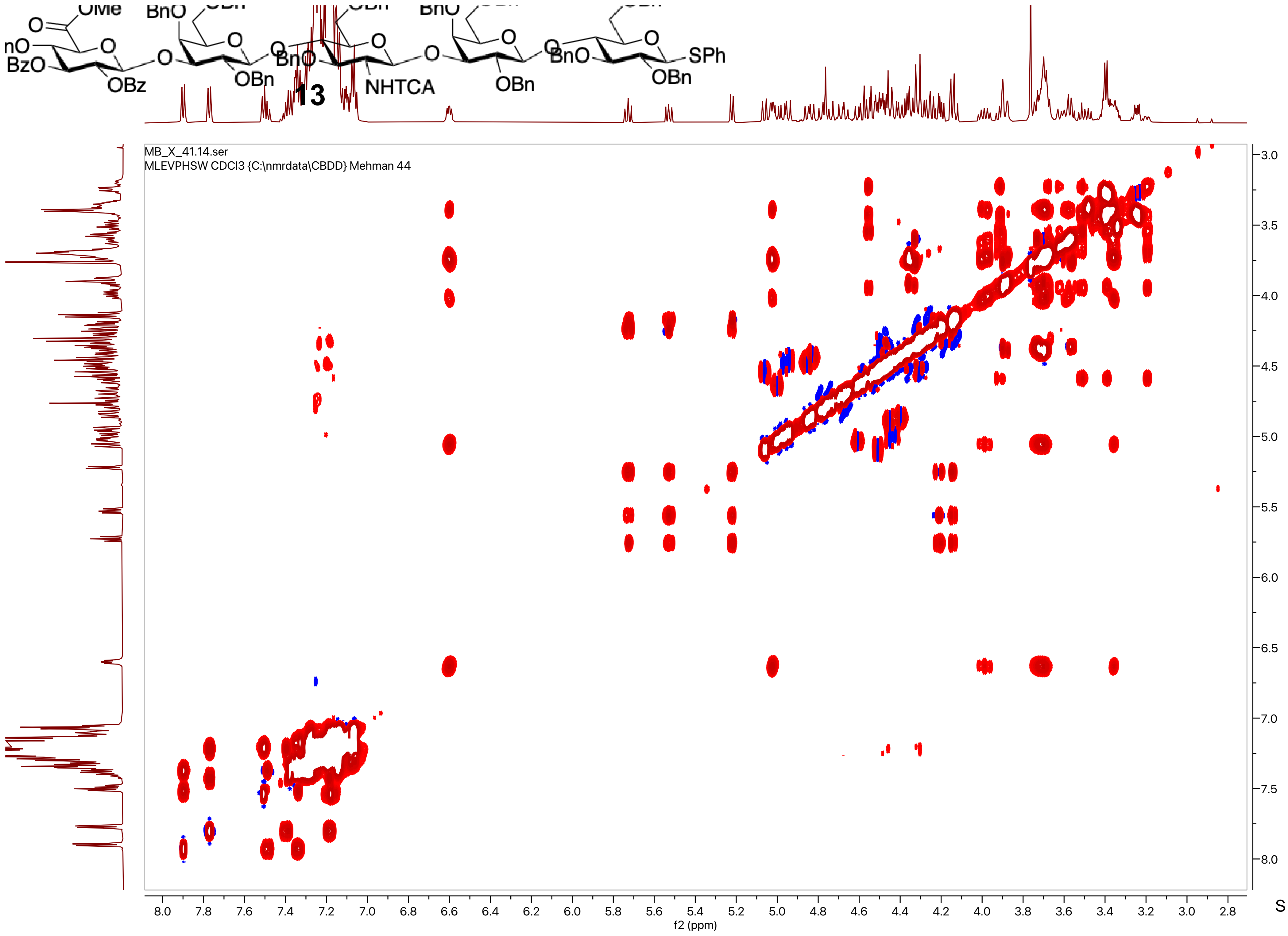

13
MB_X_41.14.ser
MLEVPHSW CDCl3 {C:\nmrdata\CBDD} Mehman 44
f2 (ppm)
f1 (ppm)

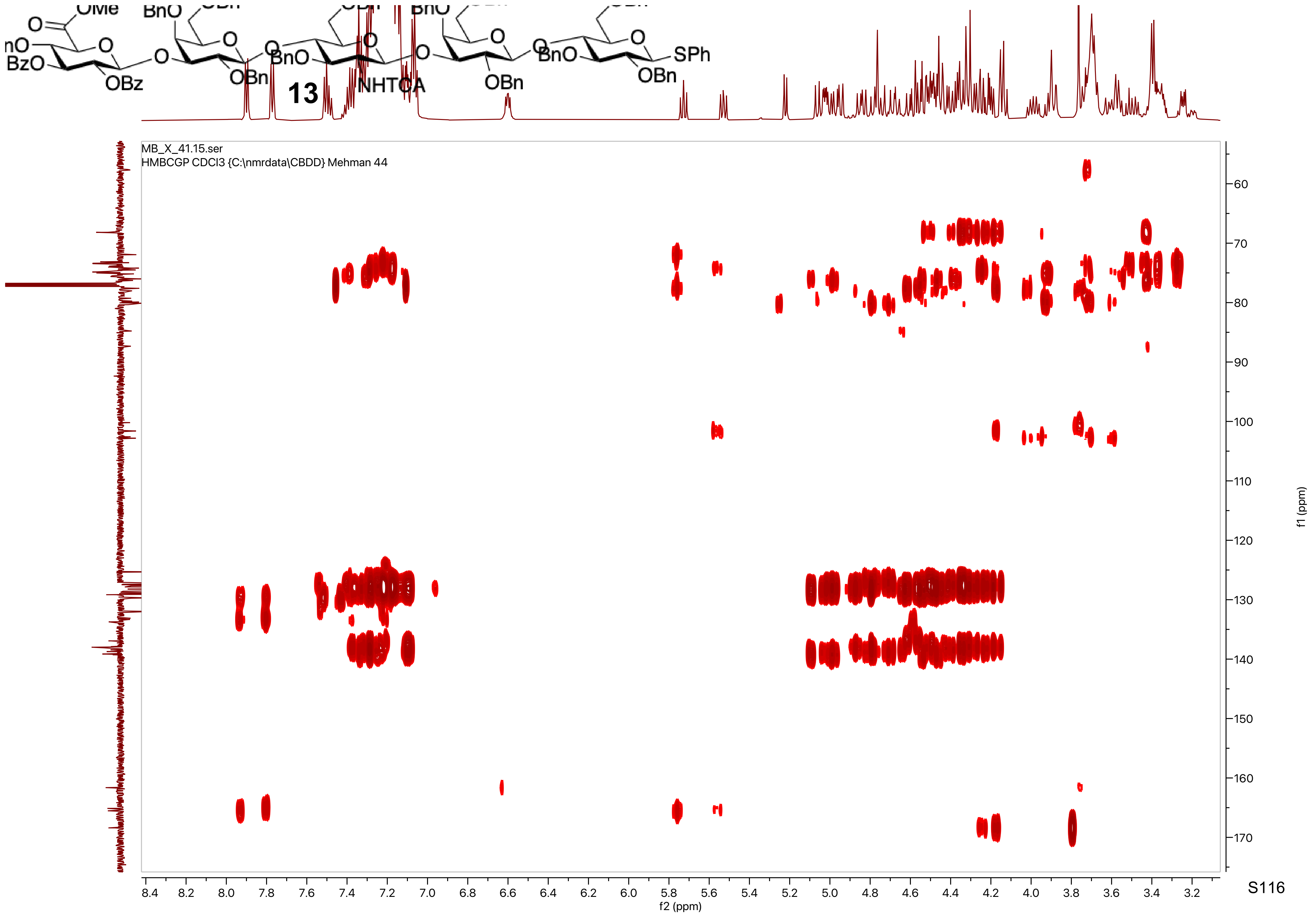

MB_X_41.15.ser
HMBCGP CDCl3 {C:\nmrdata\CBDD} Mehman 44
13
SPh
NHTCA
OBn
OBz
BzO
f2 (ppm)
f1 (ppm)

MB_X_19.10.fid
PROTON CDCl3 {C:\nmrdata\CBDD} Mehman 15

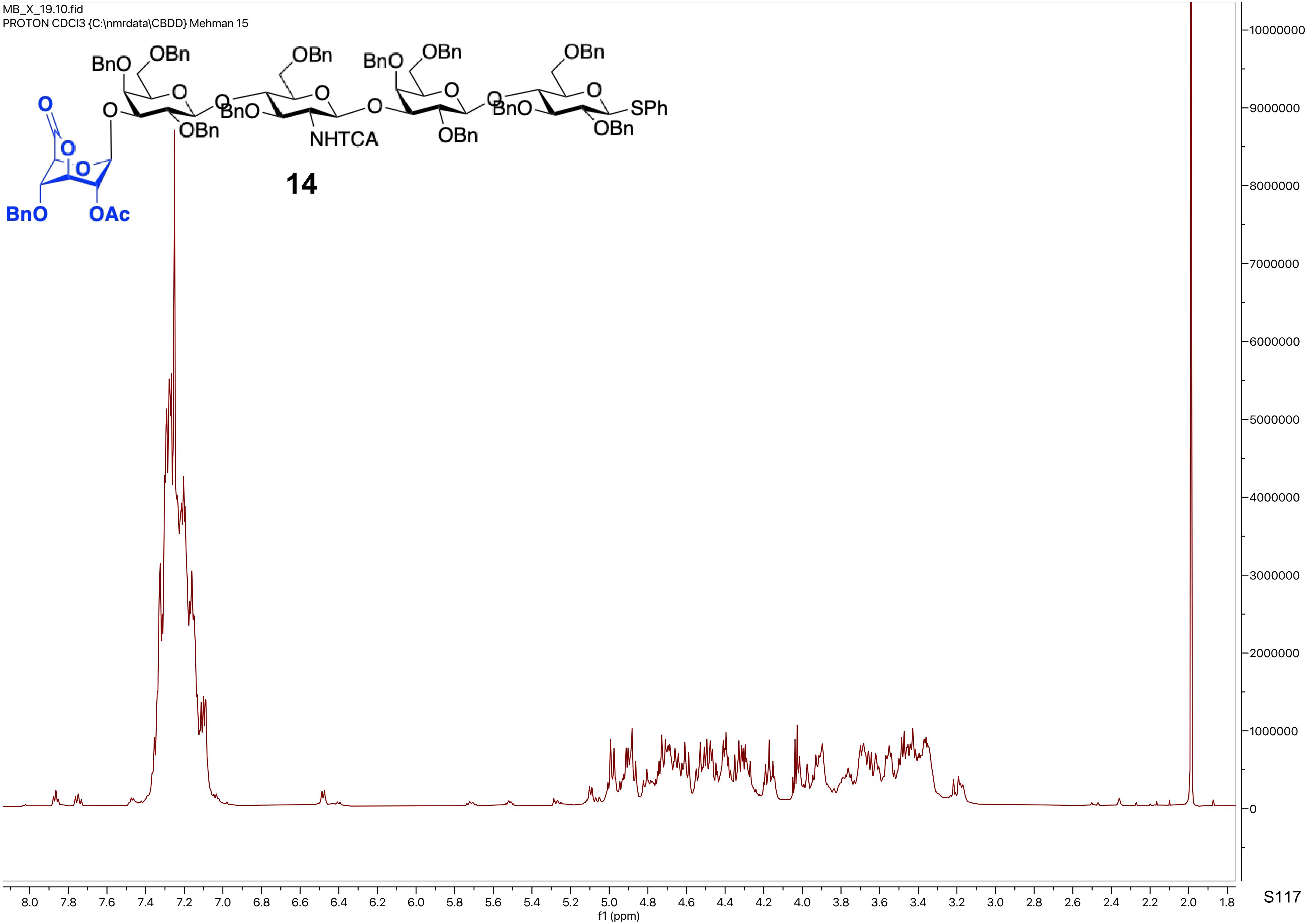

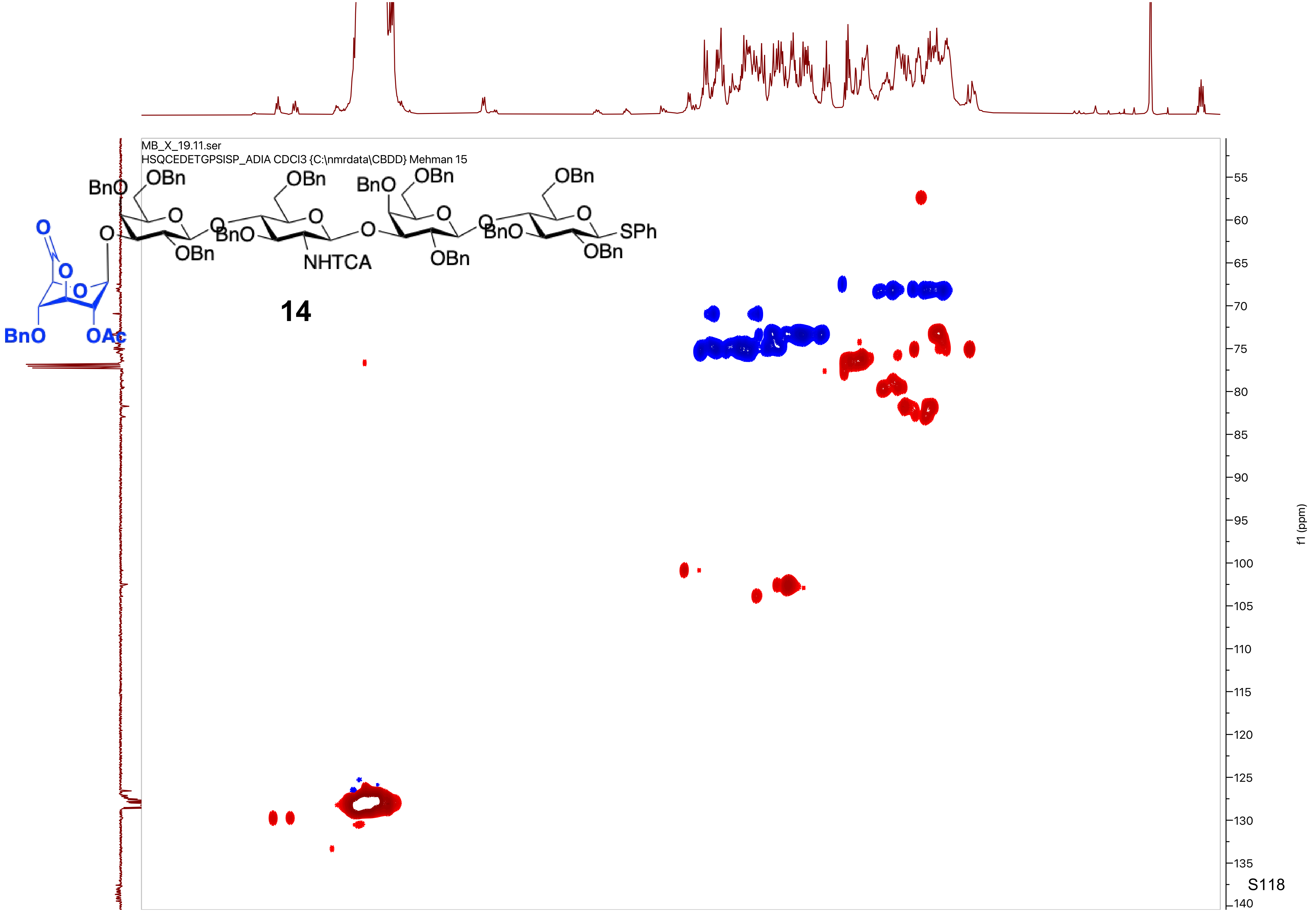
MB_X_19.11.ser
HSQCEDETGPSISP_ADIA CDCl3 {C:\nmrdata\CBDD} Mehman 15
BnO
OBn
O
OBn
NHTCA
BnO
SPh
OAc
14
f1 (ppm)
55
60
65
70
75
80
85
90
95
100
105
110
115
120
125
130
135
140

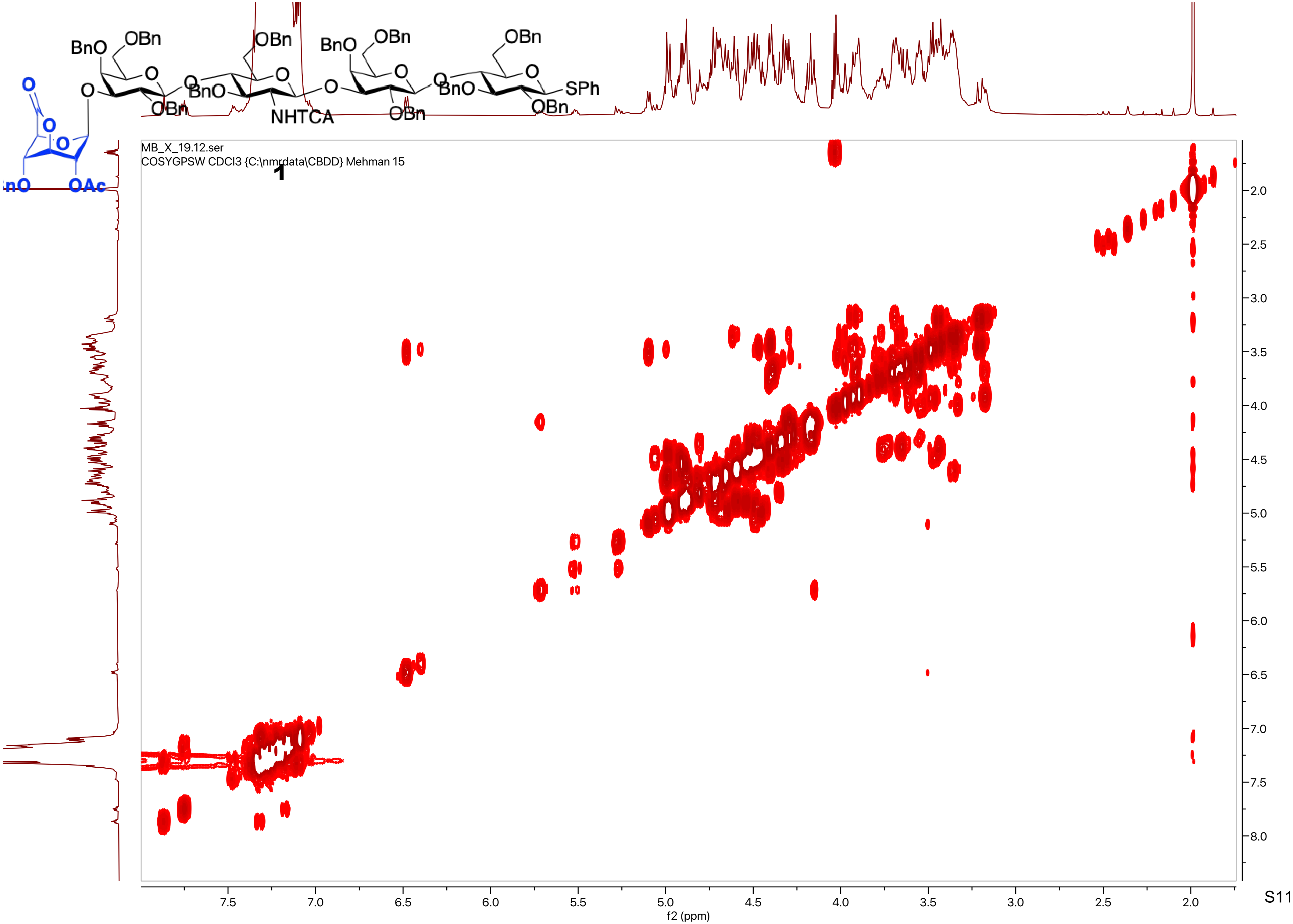
MB_X_19.12.ser
COSYGPSW CDCl3 {C:\nmrdata\CBDD} Mehman 15
1
f2 (ppm)
f1 (ppm)

MB_X_19.13.fid
C13APT CDCl3 {C:\nmrdata\CBDD} Mehman 15

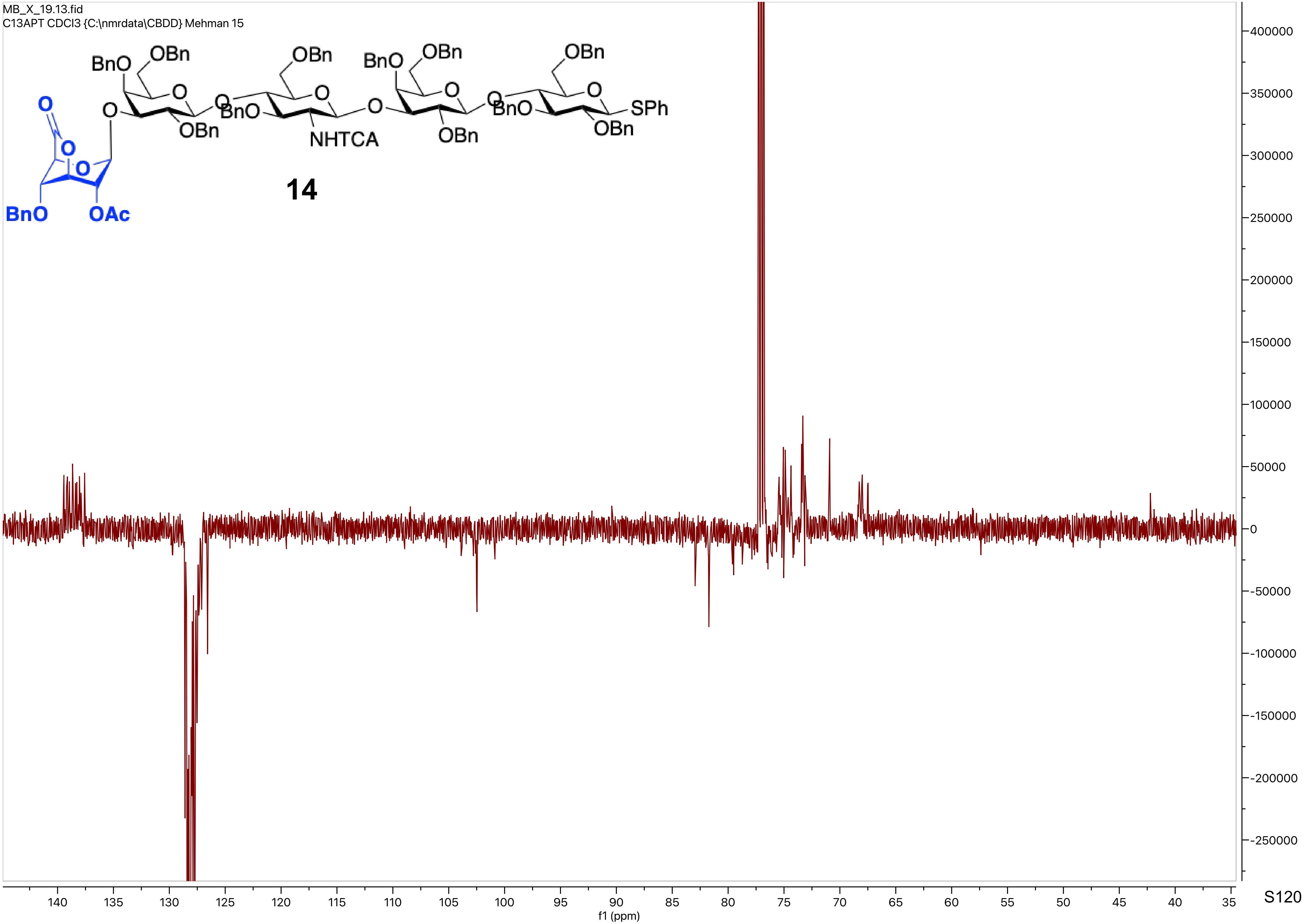

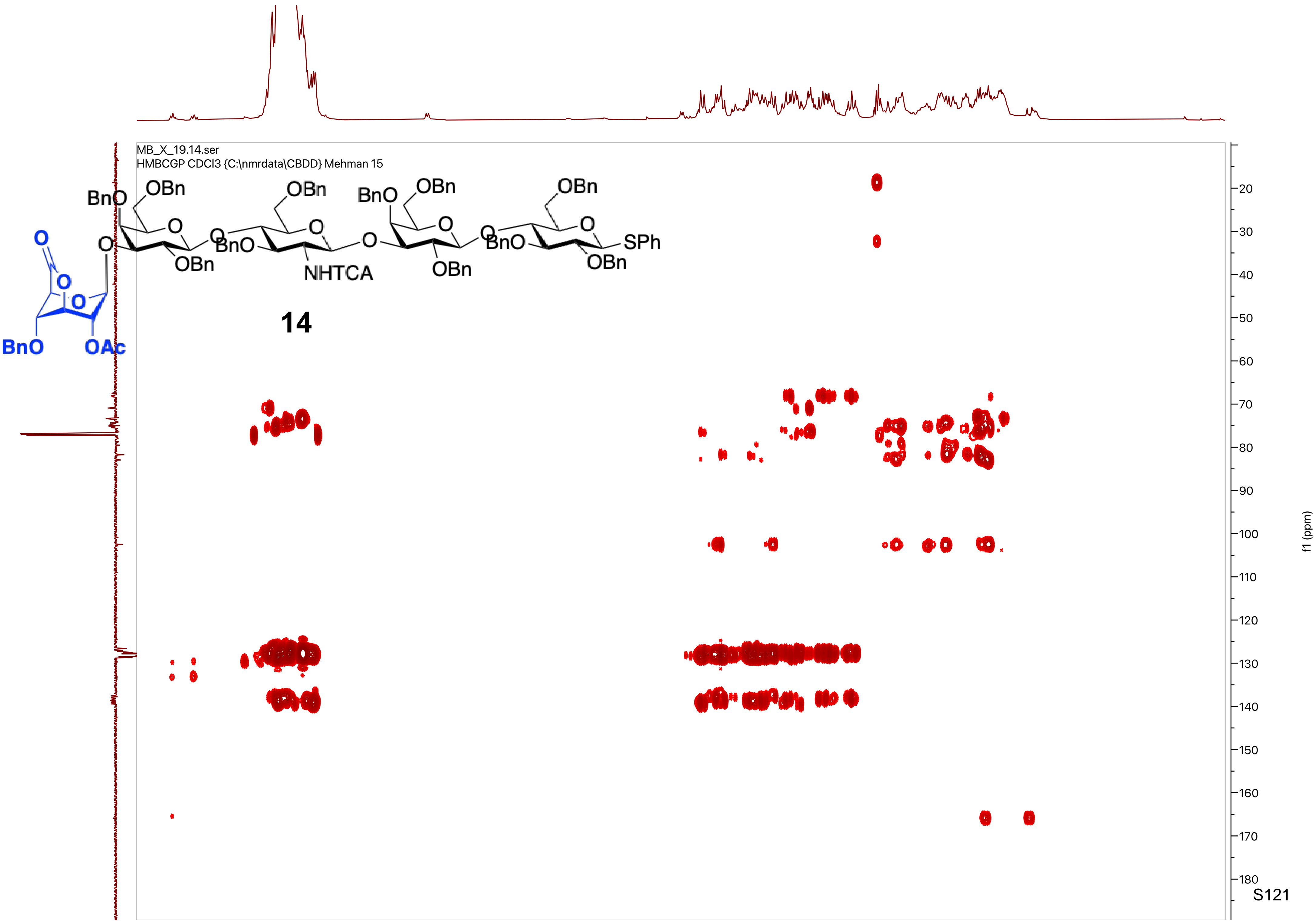
MB_X_19.14.ser
HMBCGP CDCl3 {C:\nmrdata\CBDD} Mehman 15
14
f1 (ppm)

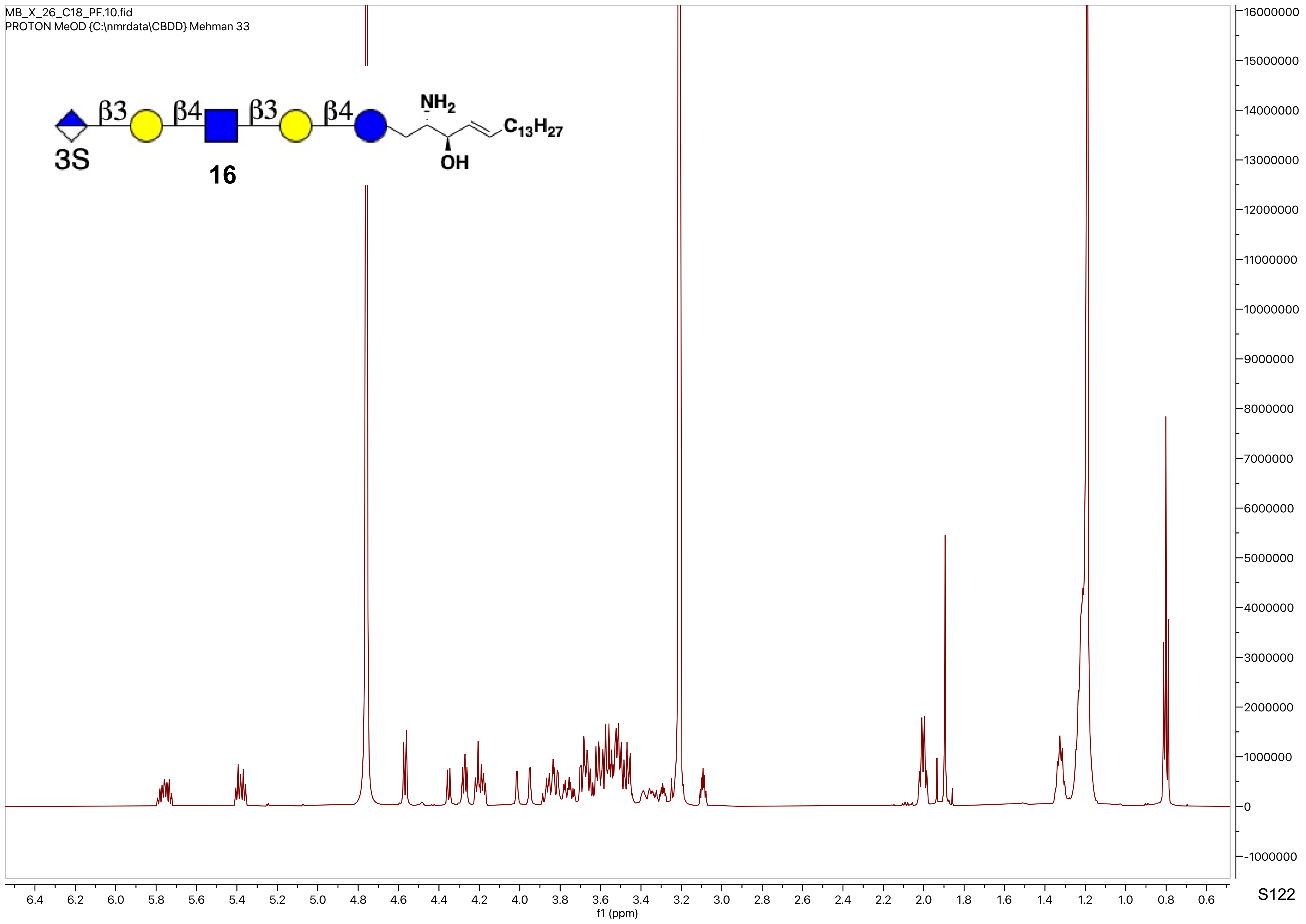
MB_X_26_C18_PF.10.fid
PROTON MeOD {C:\nmrdata\CBDD} Mehman 33
β3
β4
β3
β4
3S
16
$NH_2$
OH
$C_{13}H_{27}$
f1 (ppm)

MB_X_26_C18_PF.10.fid
PROTON MeOD {C:\nmrdata\CBDD} Mehman 33

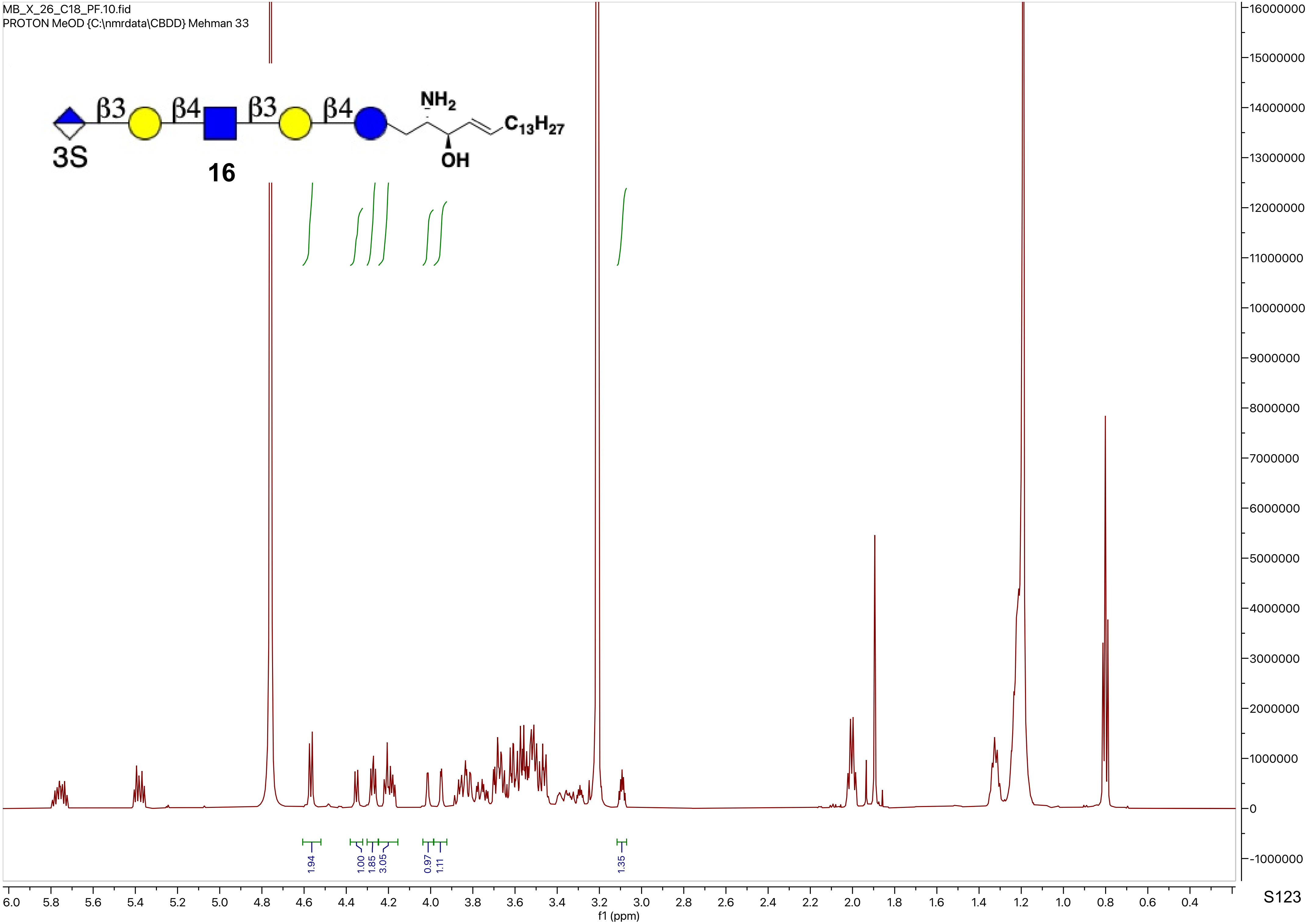

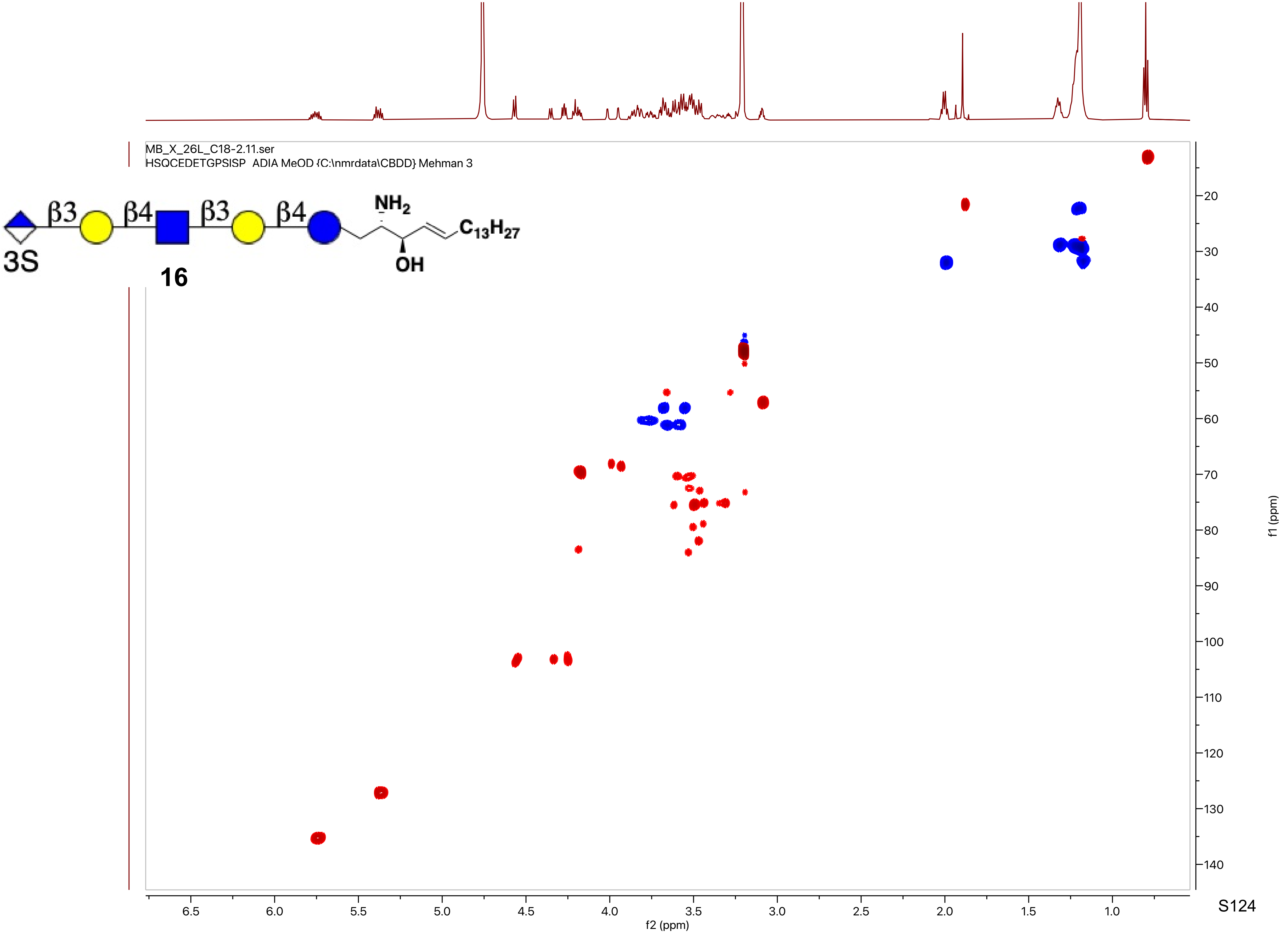
MB_X_26L_C18-2.11.ser
HSQCEDETGPSISP ADIA MeOD {C:\nmrdata\CBDD} Mehman 3
β3
β4
β3
β4
NH2
OH
C13H27
3S
16
f2 (ppm)
f1 (ppm)

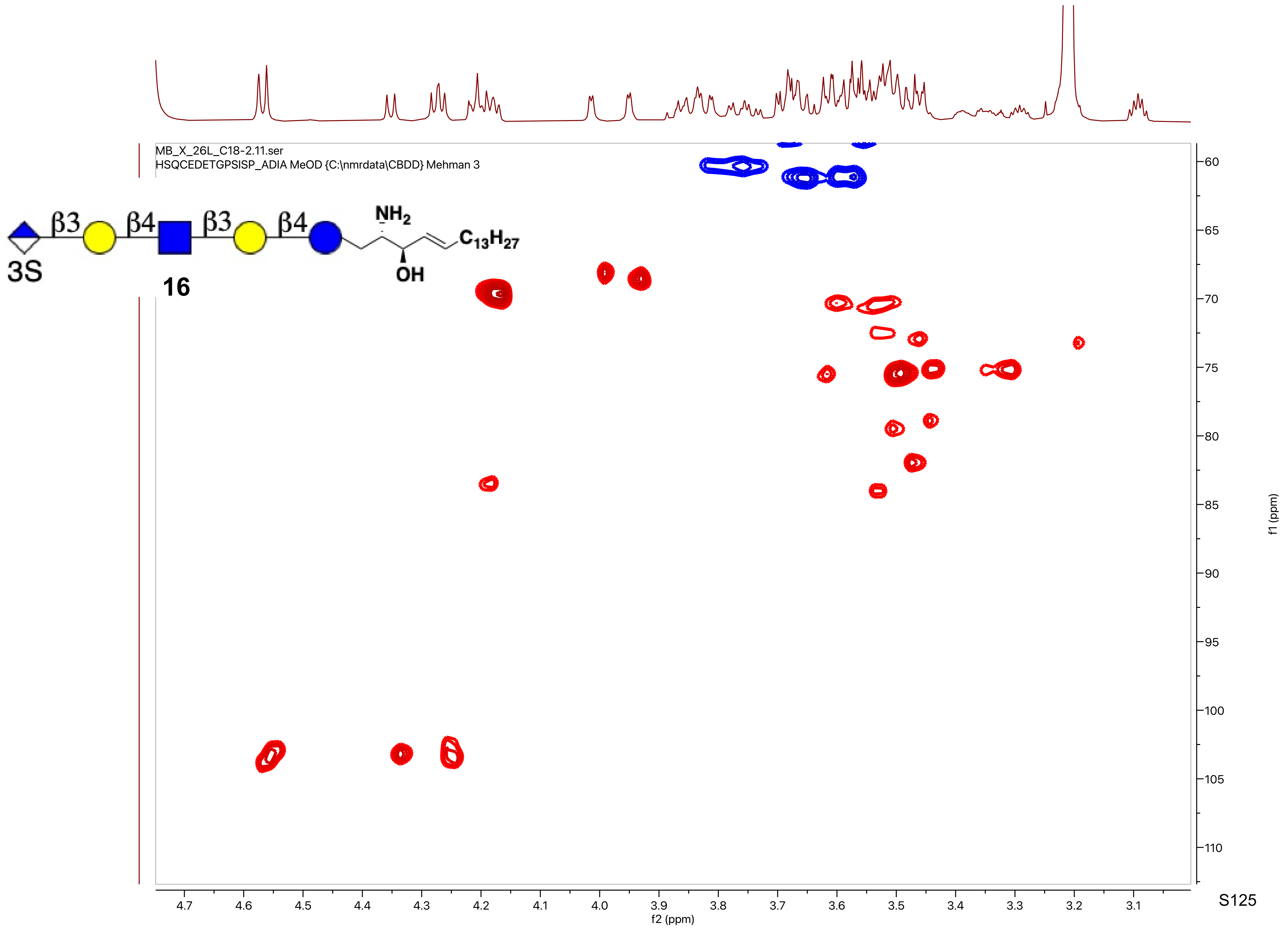
MB_X_26L_C18-2.11.ser
HSQCEDETGPSISP_ADIA MeOD {C:\nmrdata\CBDD} Mehman 3
β3
β4
β3
β4
NH2
C13H27
OH
3S
16
f2 (ppm)
f1 (ppm)

MB_X_38.10.fid
PROTON CDCl3 {C:\nmrdata\CBDD} Mehman 5

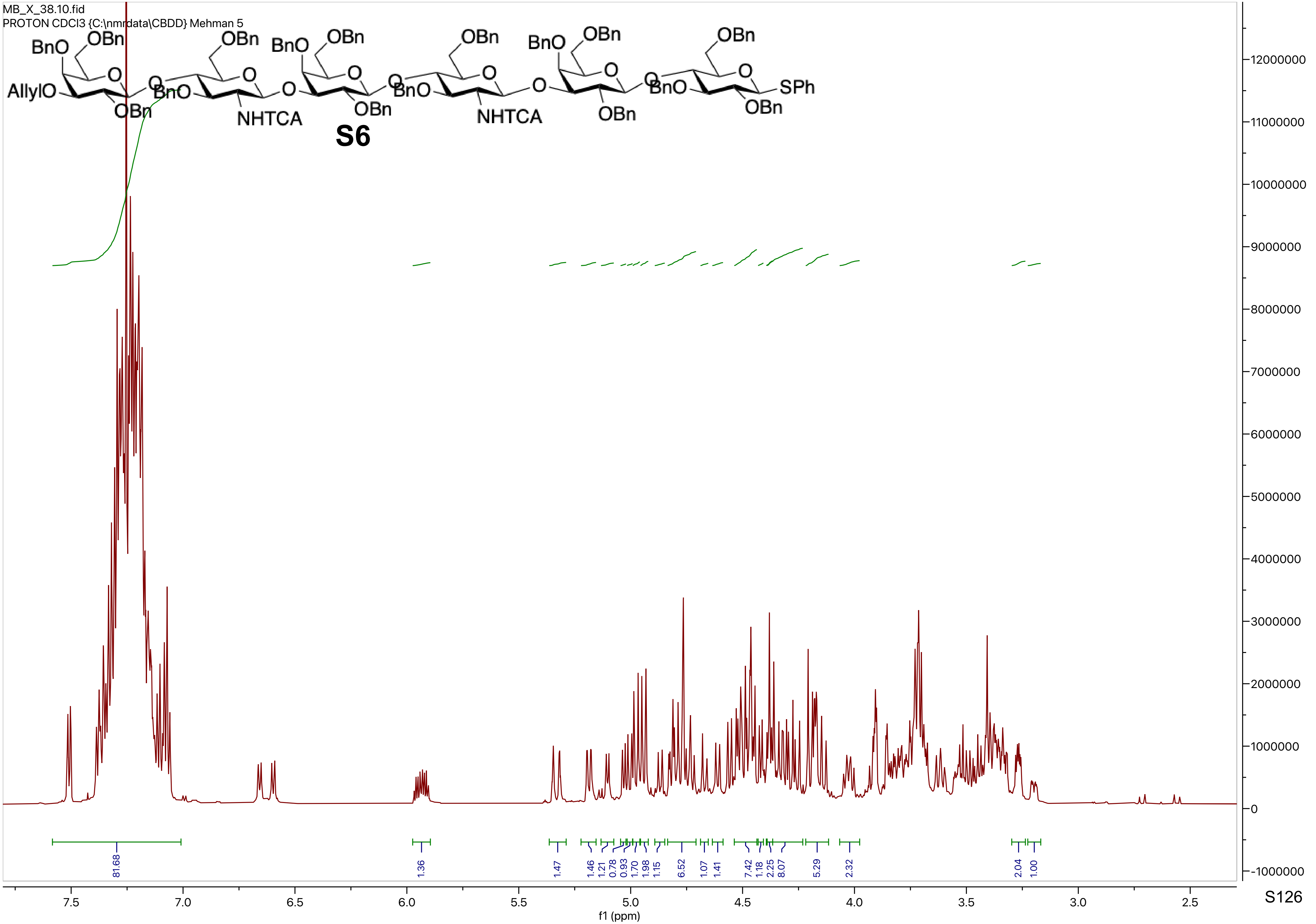

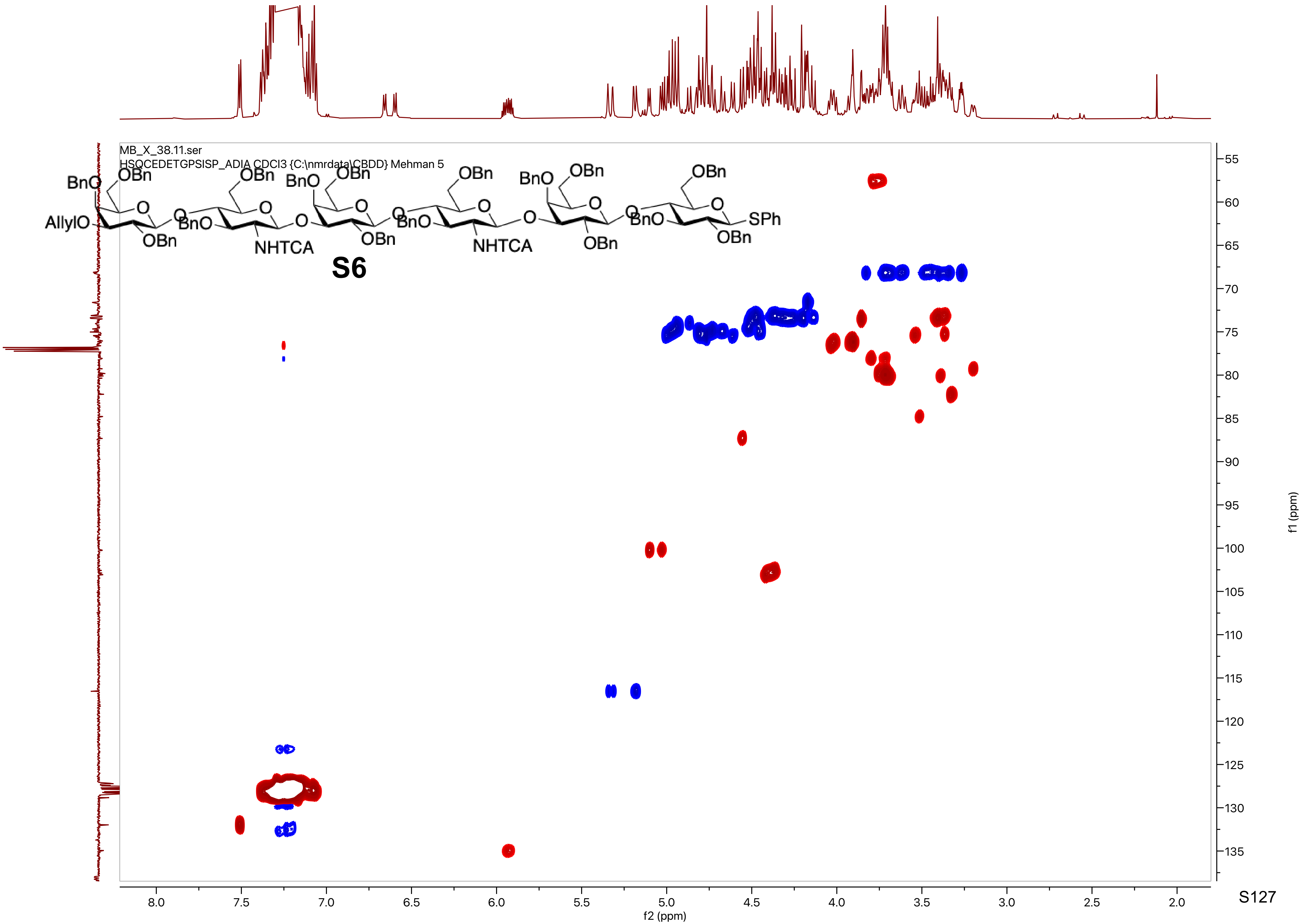
MB_X_38.11.ser
HSQCEDETGPSISP_ADIA CDCl3 {C:\nmrdata\CBDD} Mehman 5
BnO
OBn
AllylO
NHTCA
SPh
S6
f2 (ppm)
f1 (ppm)

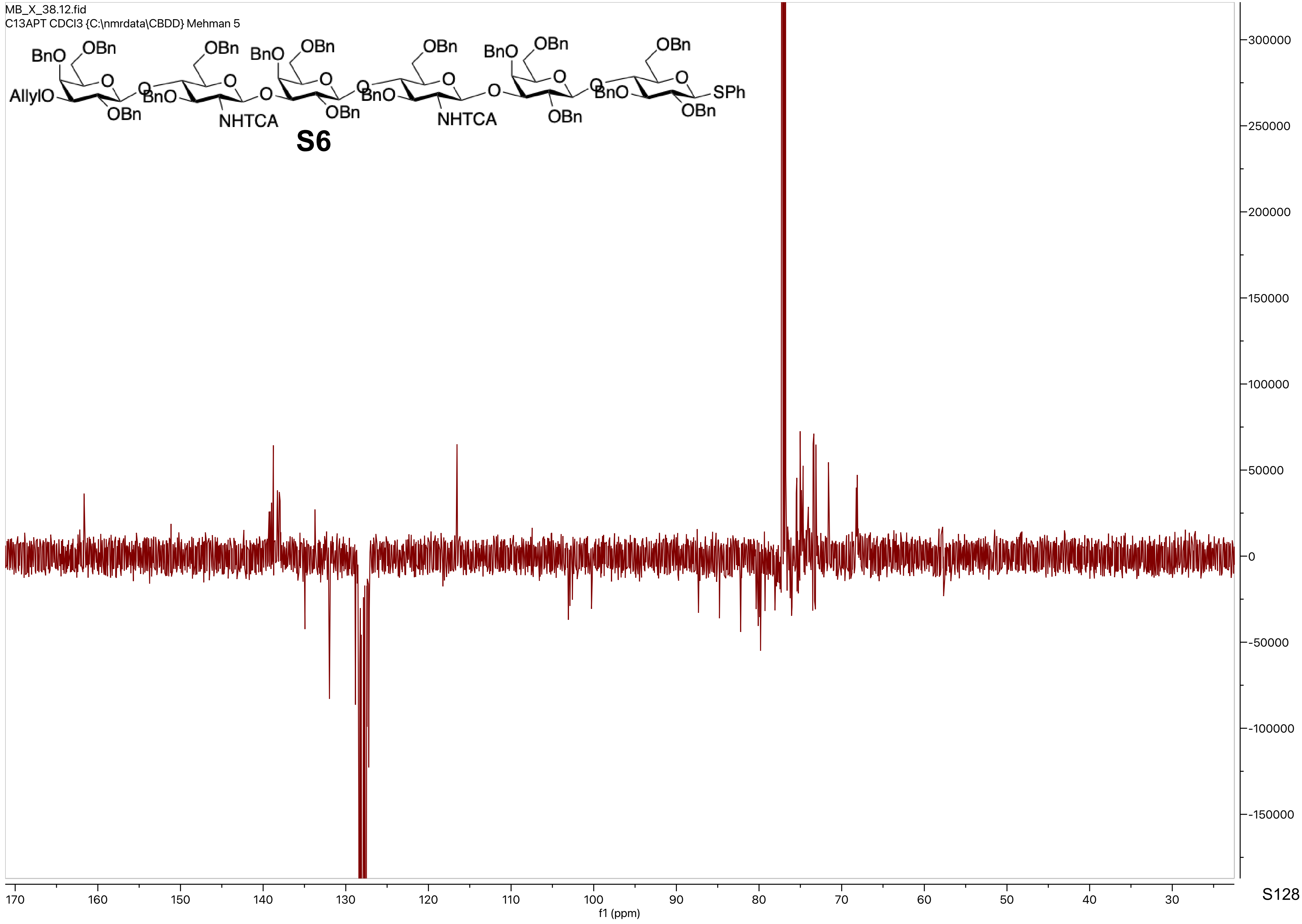
MB_X_38.12.fid
C13APT CDCl3 {C:\nmrdata\CBDD} Mehman 5
BnO
OBn
AllylO
BnO
NHTCA
SPh
S6
f1 (ppm)

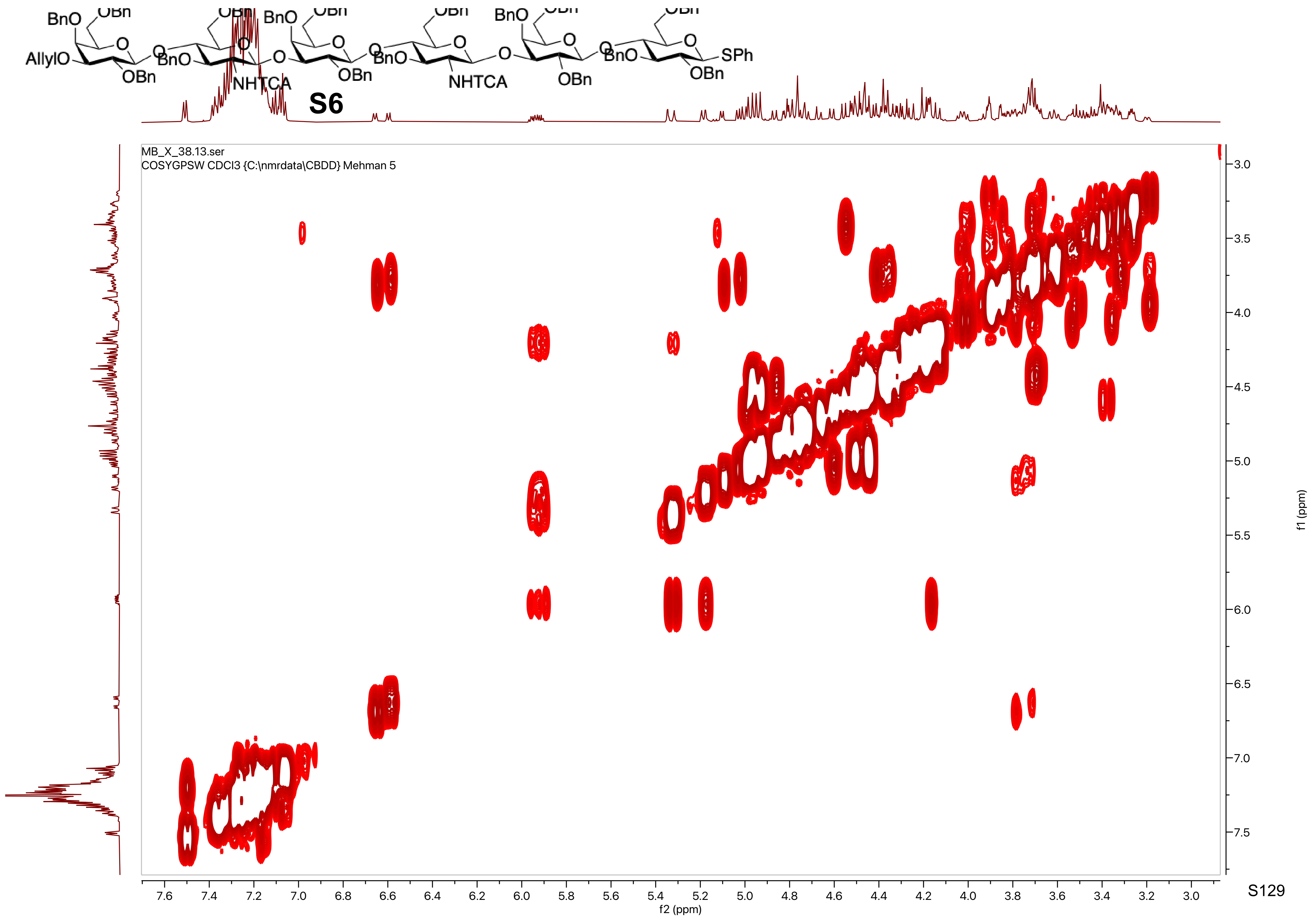

BnO
OBn
AllylO
OBn
BnO
NHTCA
NHTCA
SPh
S6
MB_X_38.13.ser
COSYGPSW CDCl3 {C:\nmrdata\CBDD} Mehman 5
f2 (ppm)
f1 (ppm)

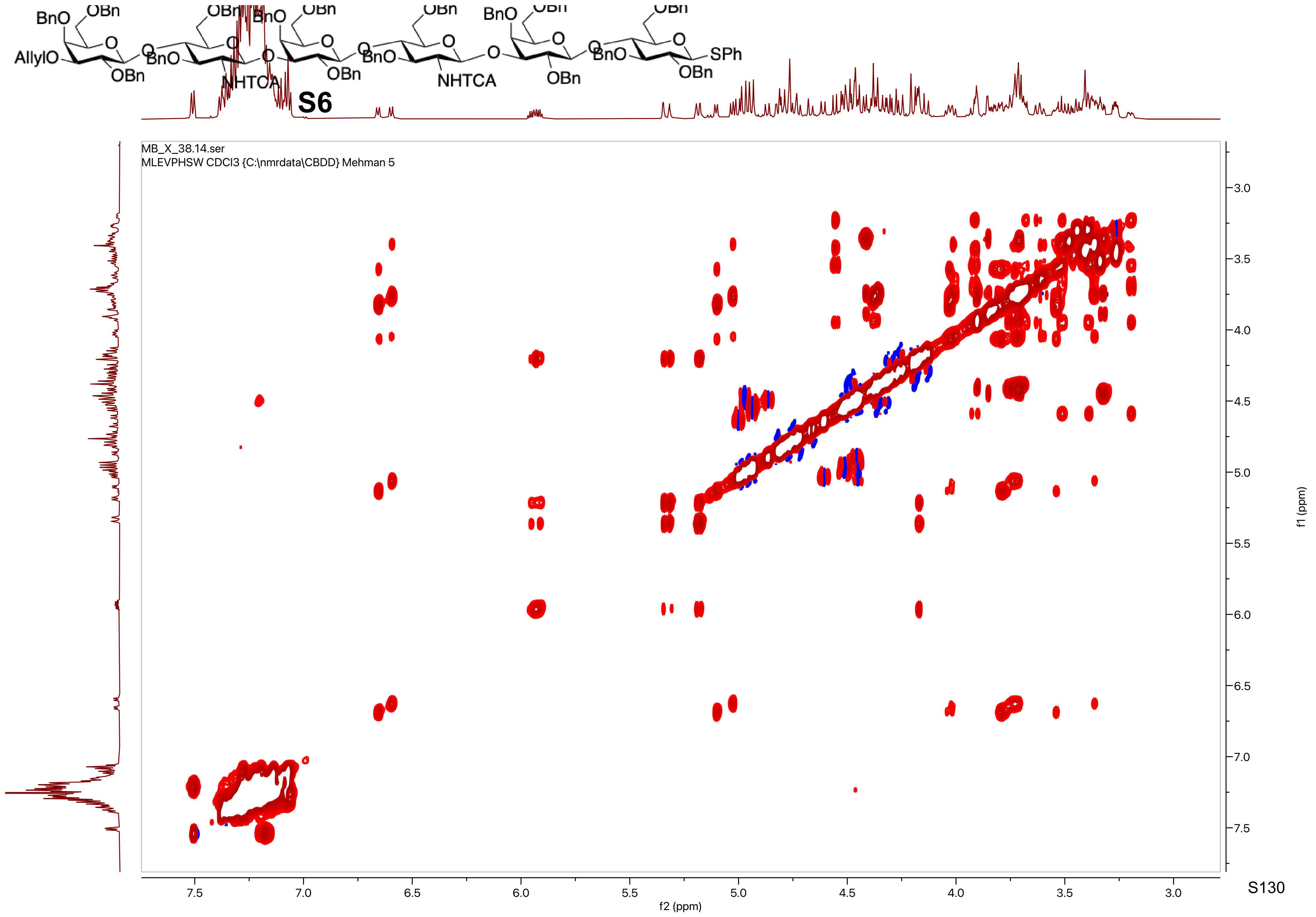
S6
MB_X_38.14.ser
MLEVPHSW CDCl3 {C:\nmrdata\CBDD} Mehman 5
f2 (ppm)
f1 (ppm)

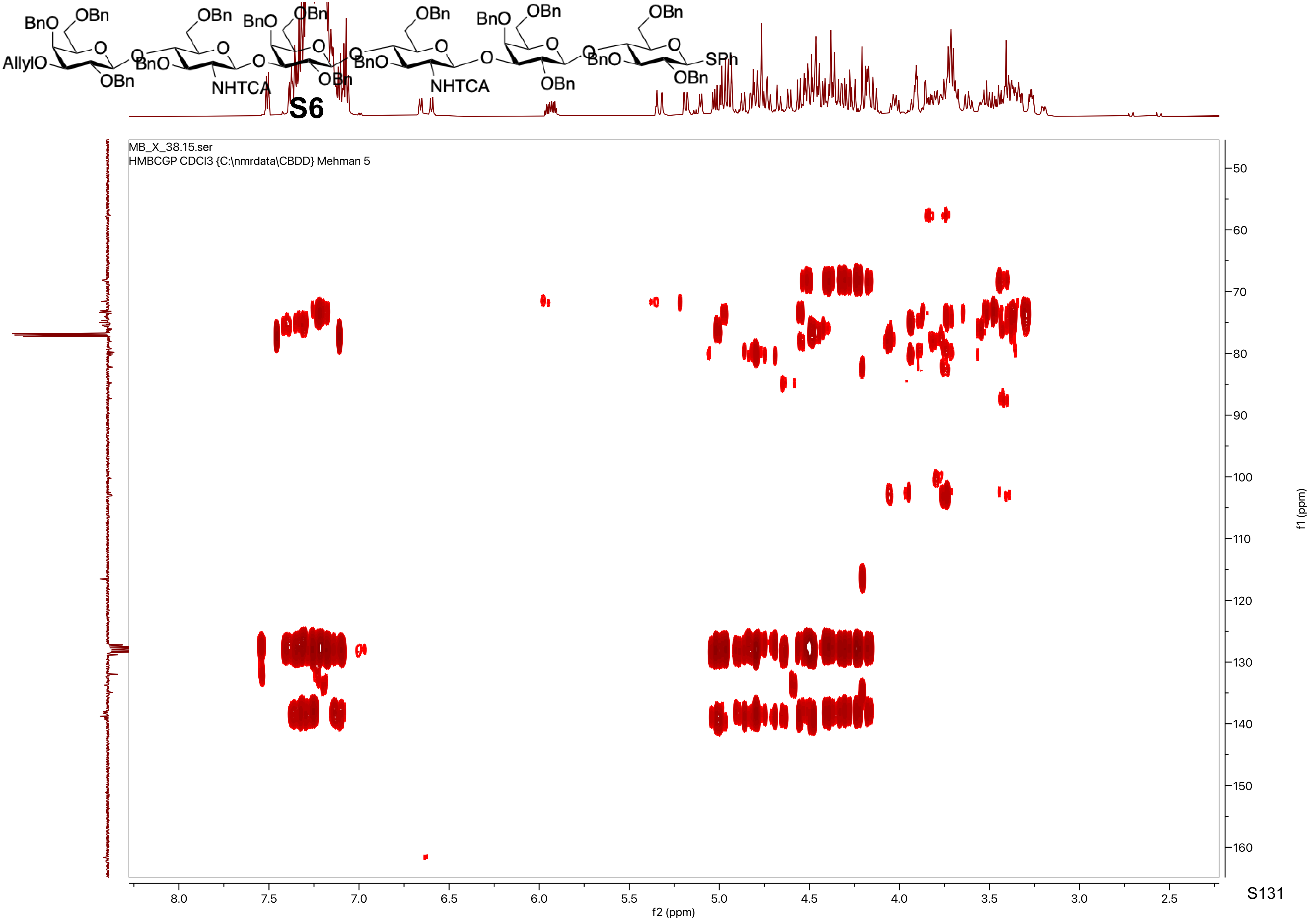

BnO
OBn
AllylO
OBn
BnO
NHTCA
BnO
OBn
OBn
BnO
NHTCA
BnO
OBn
OBn
BnO
OBn
SPh
S6
MB_X_38.15.ser
HMBCGP CDCl3 {C:\nmrdata\CBDD} Mehman 5
f2 (ppm)
f1 (ppm)

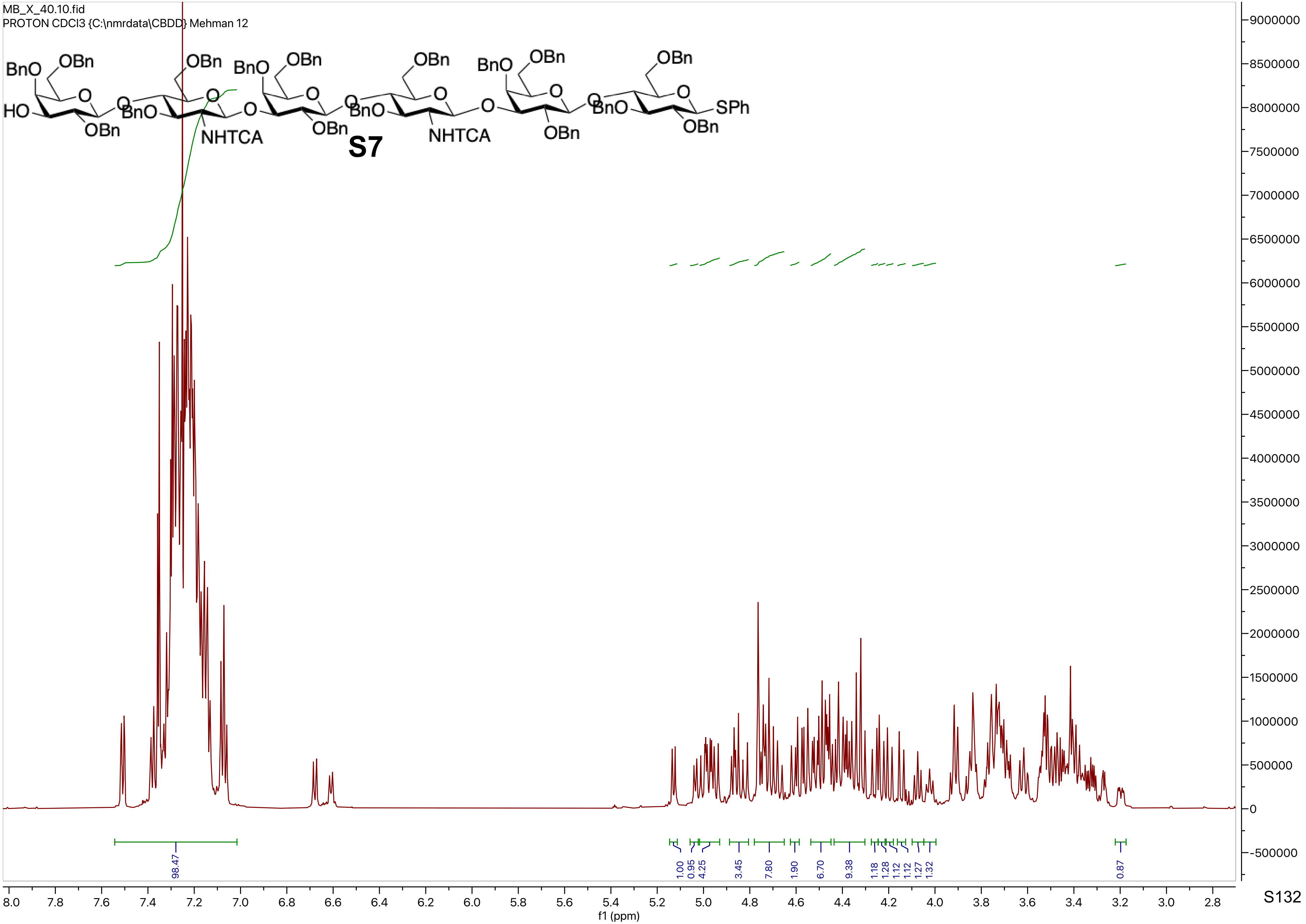

MB_X_40.10.fid
PROTON CDCl3 {C:\nmrdata\CBDD} Mehman 12
S7
BnO
OBn
HO
NHTCA
SPh
98.47
1.00
0.95
4.25
3.45
7.80
1.90
6.70
9.38
1.18
1.28
1.12
1.12
1.27
1.32
0.87
f1 (ppm)

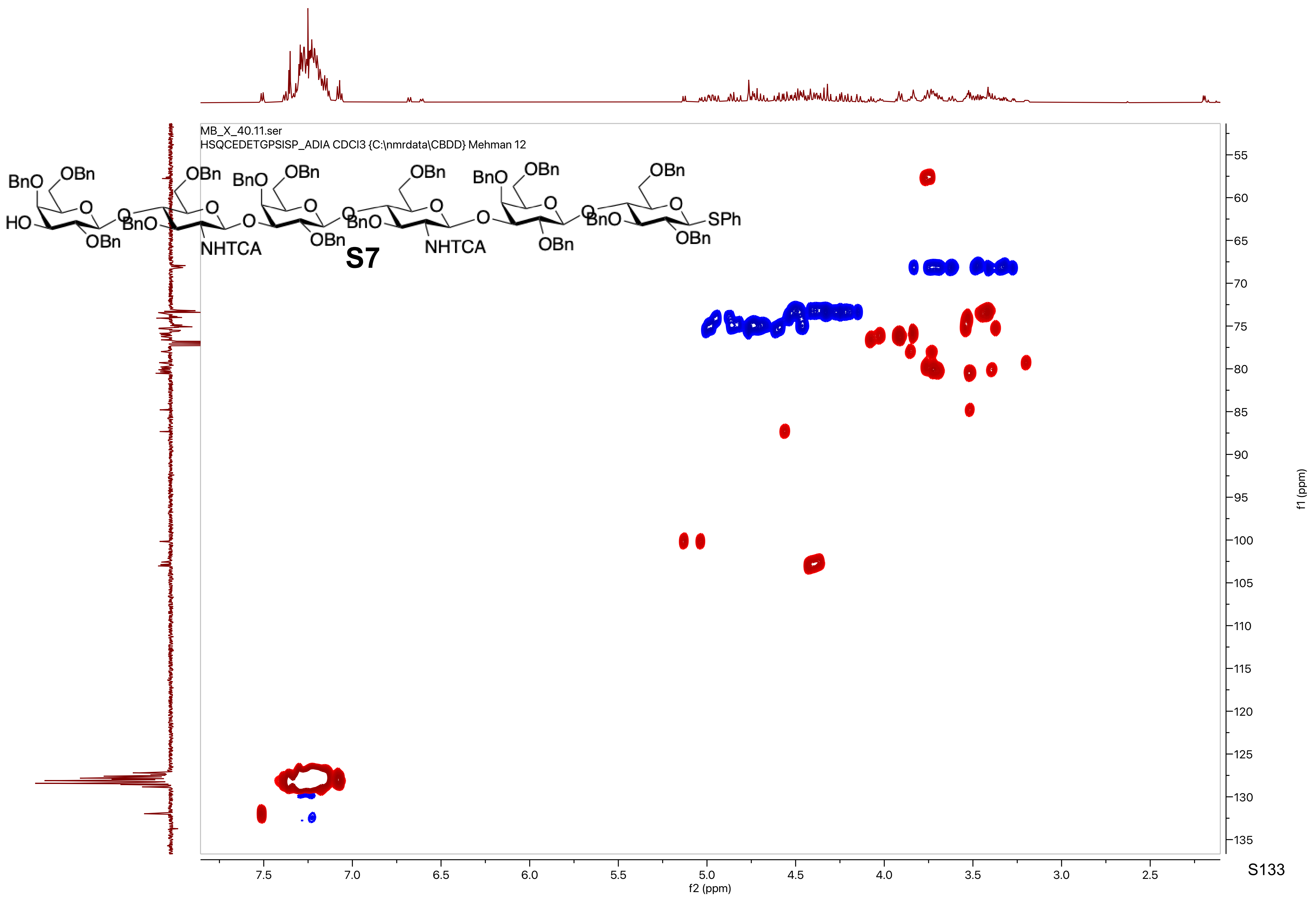
MB_X_40.11.ser
HSQCEDETGPSISP_ADIA CDCl3 {C:\nmrdata\CBDD} Mehman 12
BnO
OBn
HO
OBn
BnO
NHTCA
OBn
S7
SPh
f2 (ppm)
f1 (ppm)

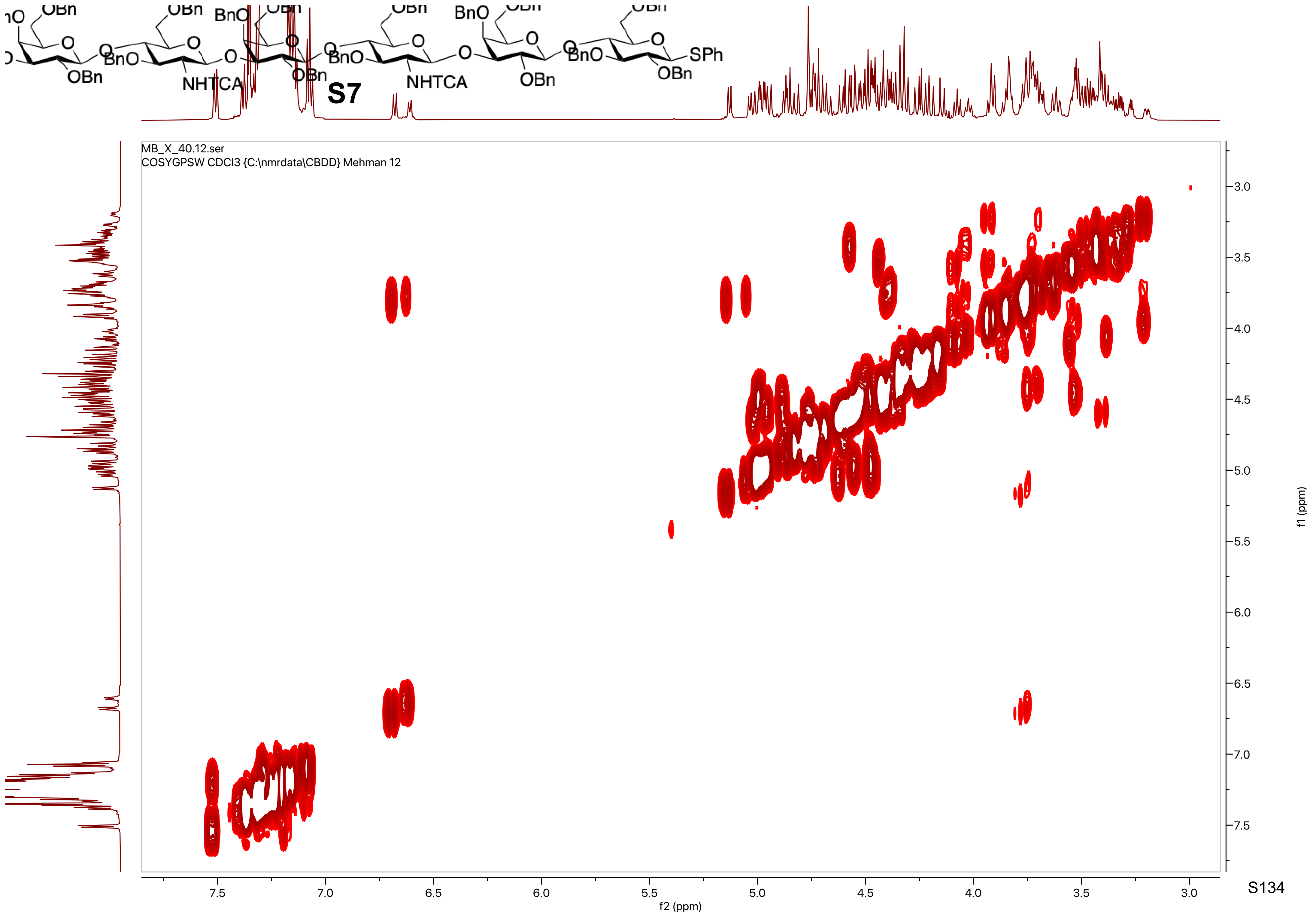
OBn
BnO
NHTCA
OBn
S7
SPh
MB_X_40.12.ser
COSYGPSW CDCl3 {C:\nmrdata\CBDD} Mehman 12
f2 (ppm)
f1 (ppm)
7.5
7.0
6.5
6.0
5.5
5.0
4.5
4.0
3.5
3.0

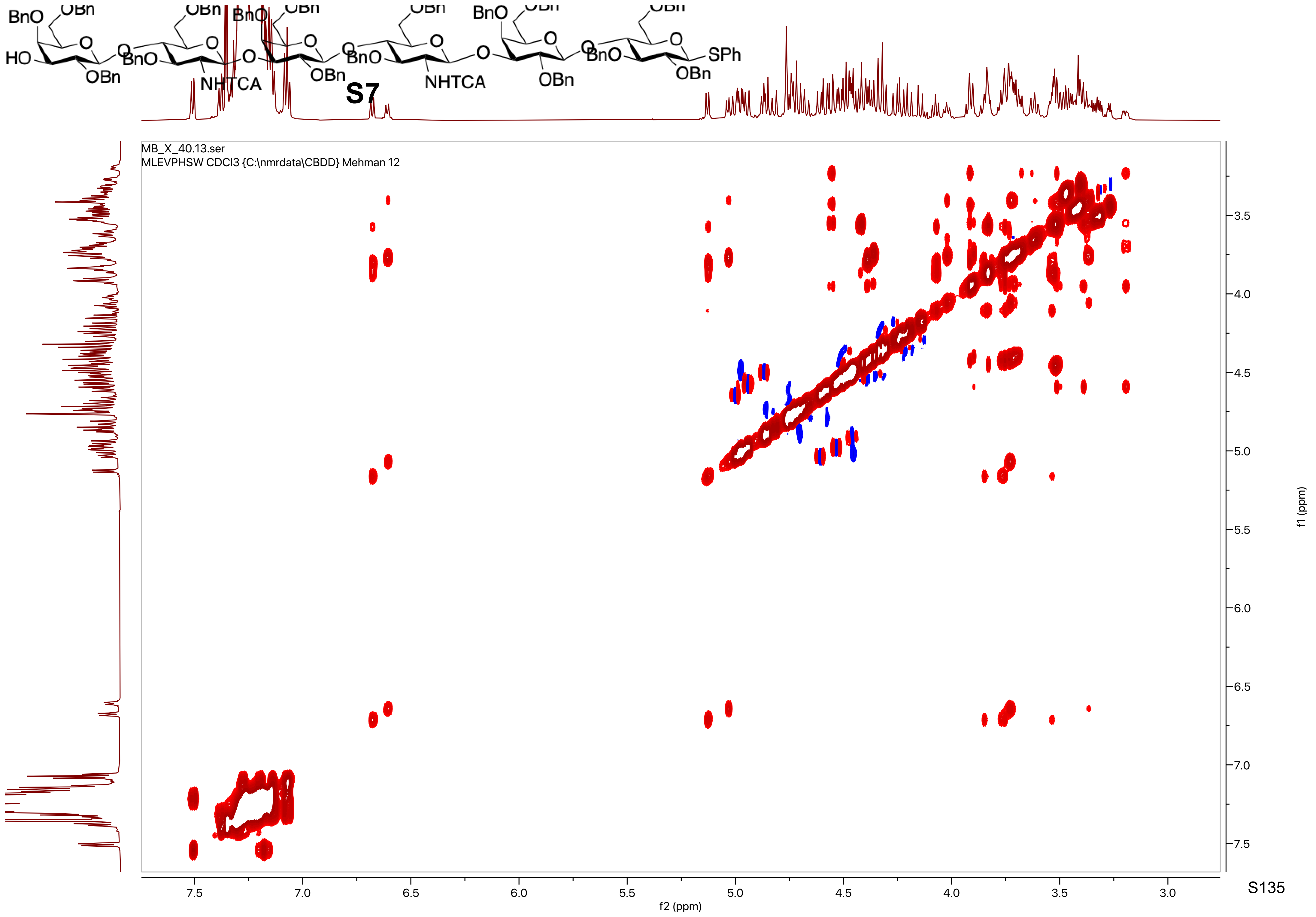

HO
BnO
OBn
NHTCA
SPh
S7
MB_X_40.13.ser
MLEVPHSW CDCl3 {C:\nmrdata\CBDD} Mehman 12
f2 (ppm)
f1 (ppm)

MB_X_40.14.fid
C13APT CDCl3 {C:\nmrdata\CBDD} Mehman 12

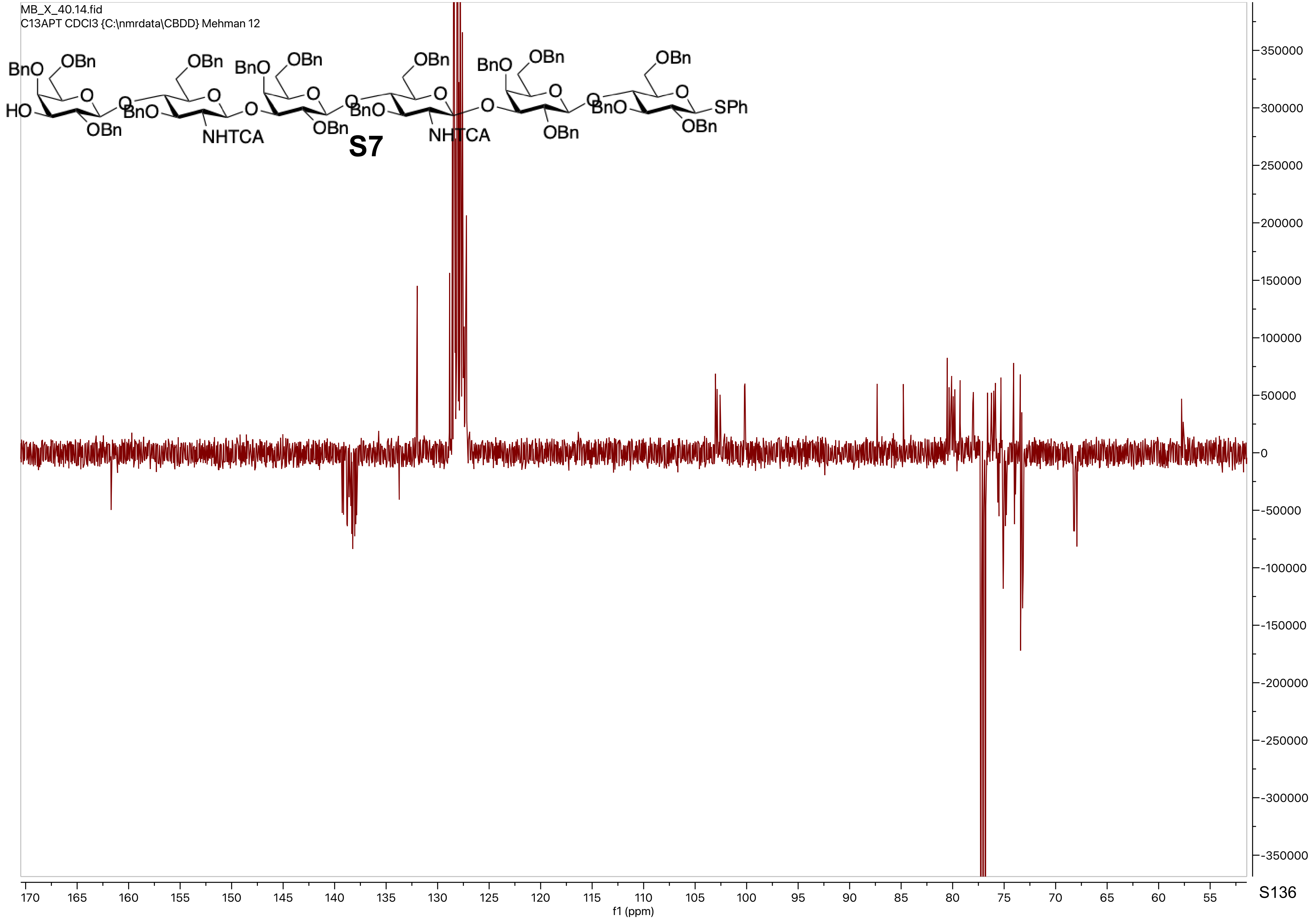

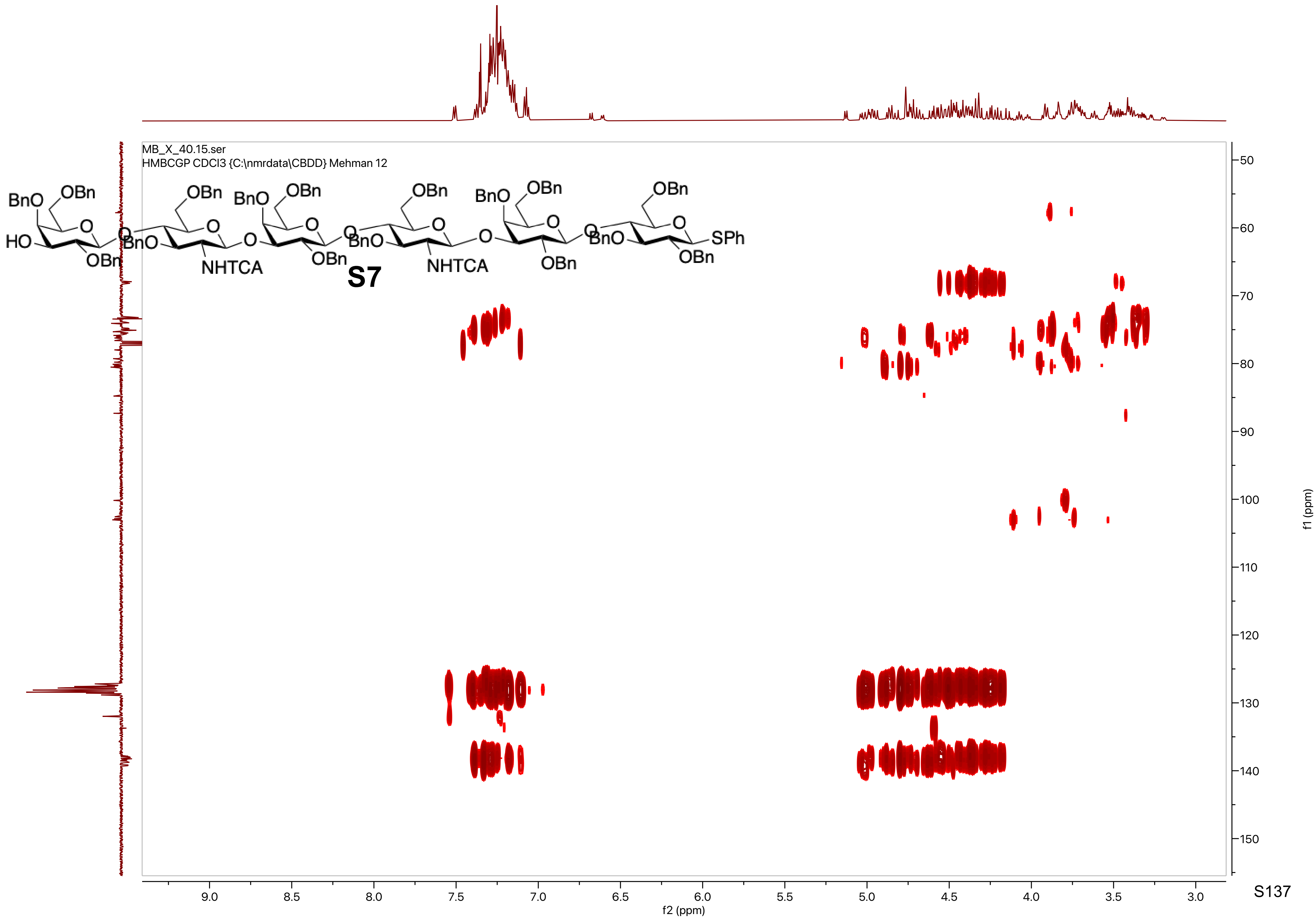
MB_X_40.15.ser
HMBCGP CDCl3 {C:\nmrdata\CBDD} Mehman 12
BnO
OBn
HO
O
OBn
NHTCA
SPh
S7
f2 (ppm)
f1 (ppm)
9.0
8.5
8.0
7.5
7.0
6.5
6.0
5.5
5.0
4.5
4.0
3.5
3.0
50
60
70
80
90
100
110
120
130
140
150

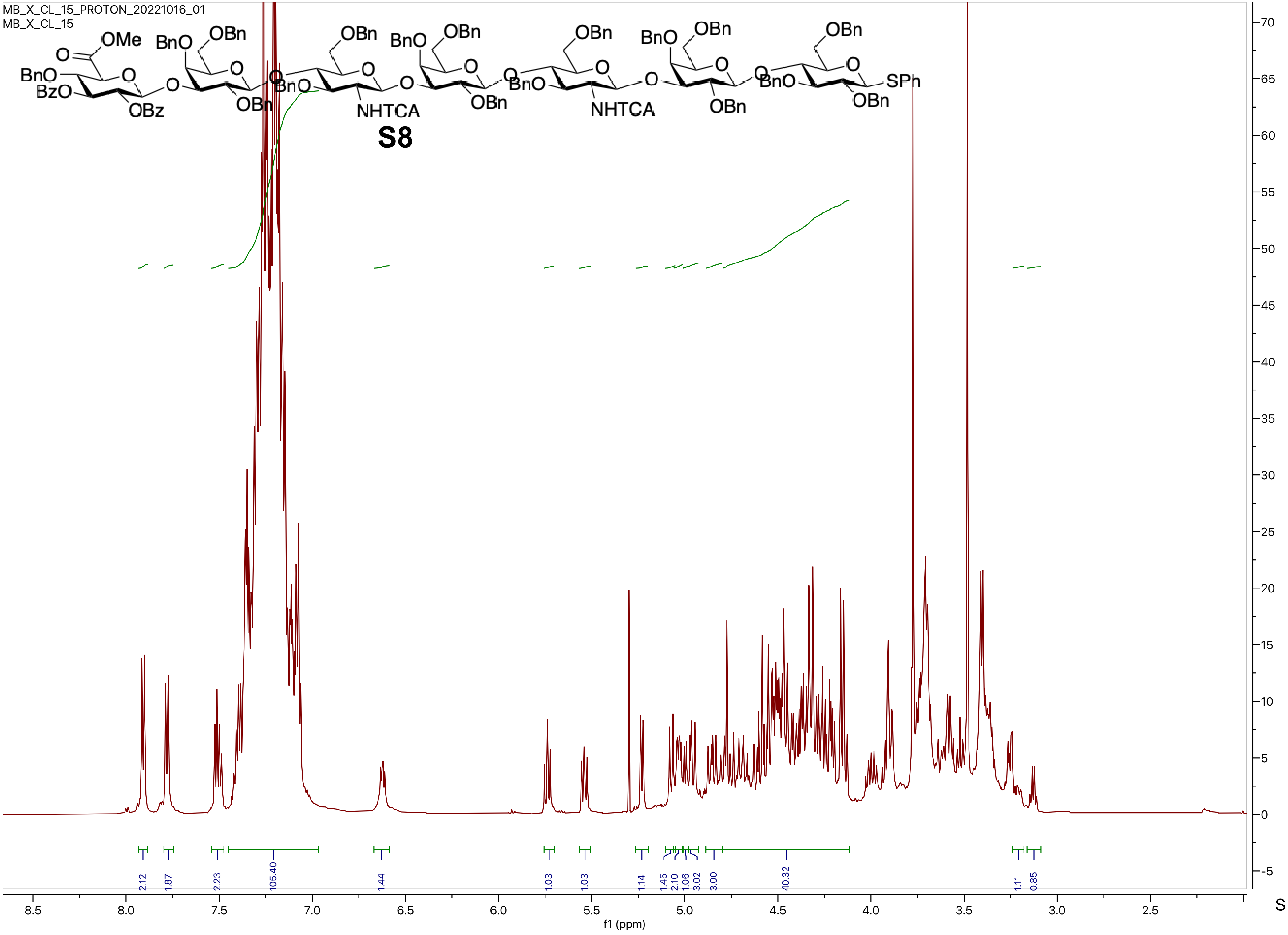
MB_X_CL_15_PROTON_20221016_01
MB_X_CL_15
S8
2.12
1.87
2.23
105.40
1.44
1.03
1.03
1.14
1.45
2.10
1.06
3.02
3.00
40.32
1.11
0.85
f1 (ppm)

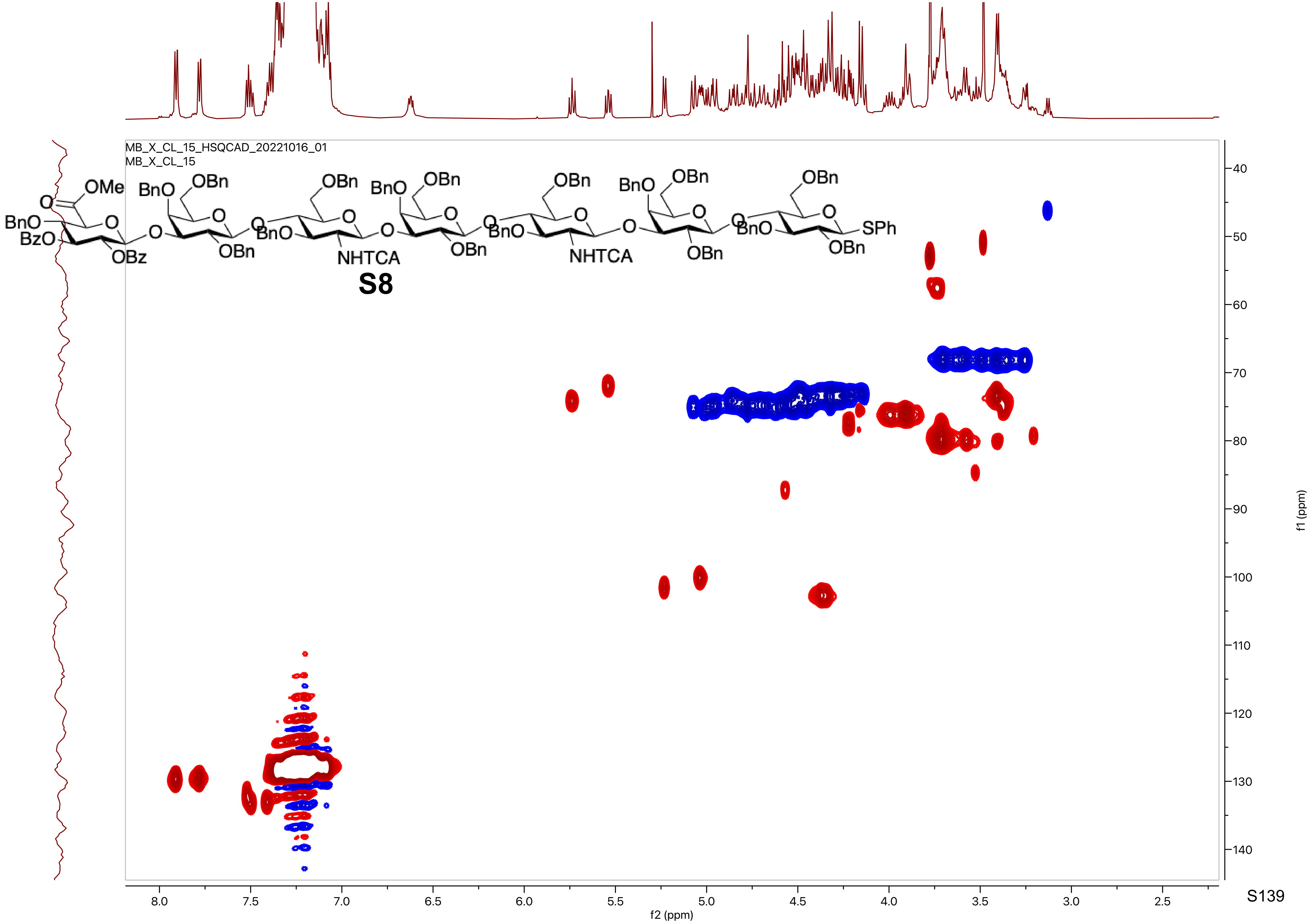
MB_X_CL_15_HSQCAD_20221016_01
MB_X_CL_15
OMe
BnO
BzO
OBz
OBn
NHTCA
SPh
S8
f2 (ppm)
f1 (ppm)

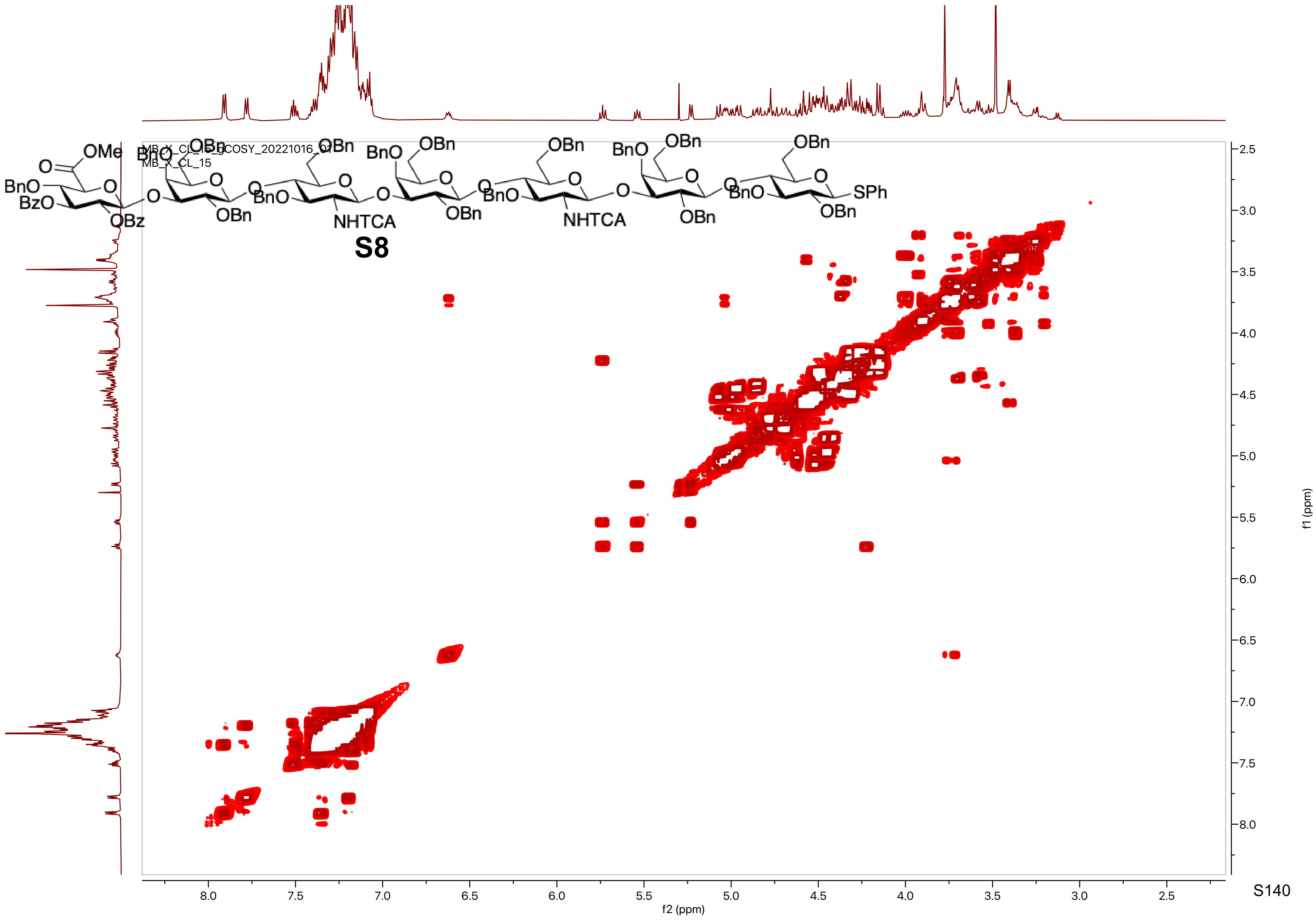

MB_X_CL_15_gCOSY_20221016_01
MB_X_CL_15
S8
f2 (ppm)
f1 (ppm)

MB_delactone_GLPG_hepta.10.fid
PROTON CDCl3 {C:\nmrdata\CBDD} Mehman 17

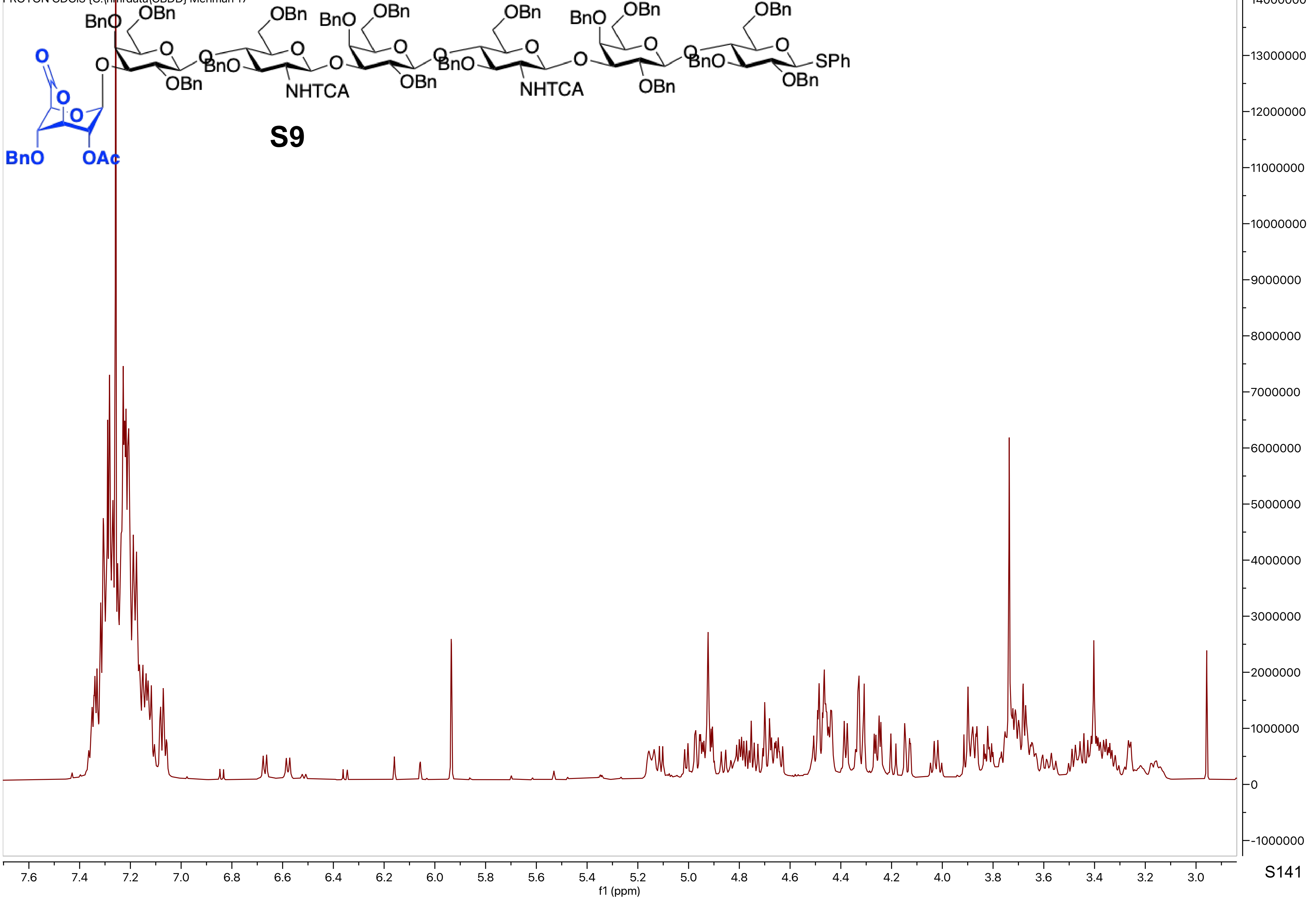

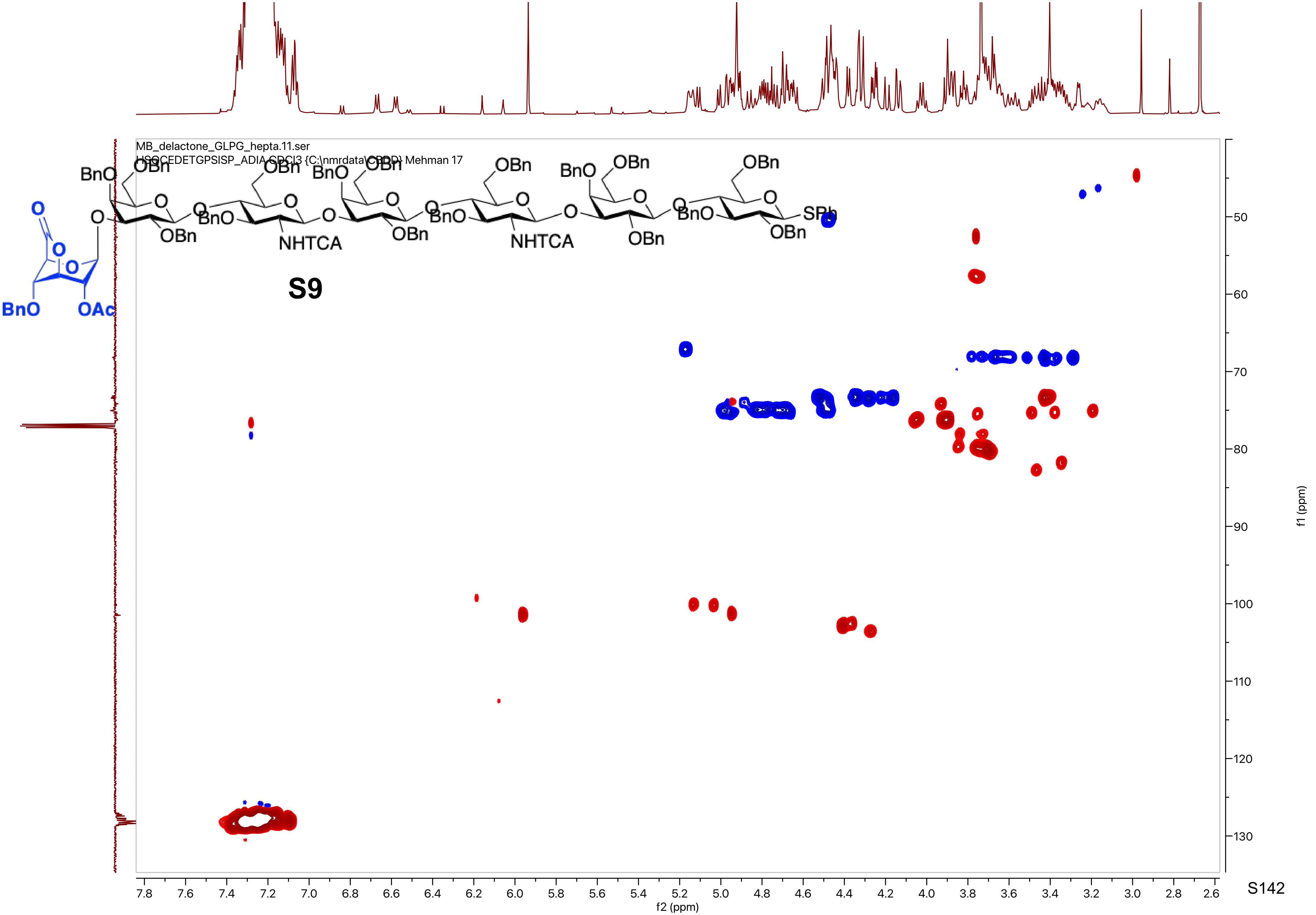

MB_delactone_GLPG_hepta.11.ser
HSQCEDETGPSISP_ADIA CDCl3 {C:\nmrdata\CBDD} Mehman 17
S9
f2 (ppm)
f1 (ppm)

MB_delactone_GLPG_hepta.12.fid
C13APT CDCl3 {C:\nmrdata\CBDD} Mehman 17

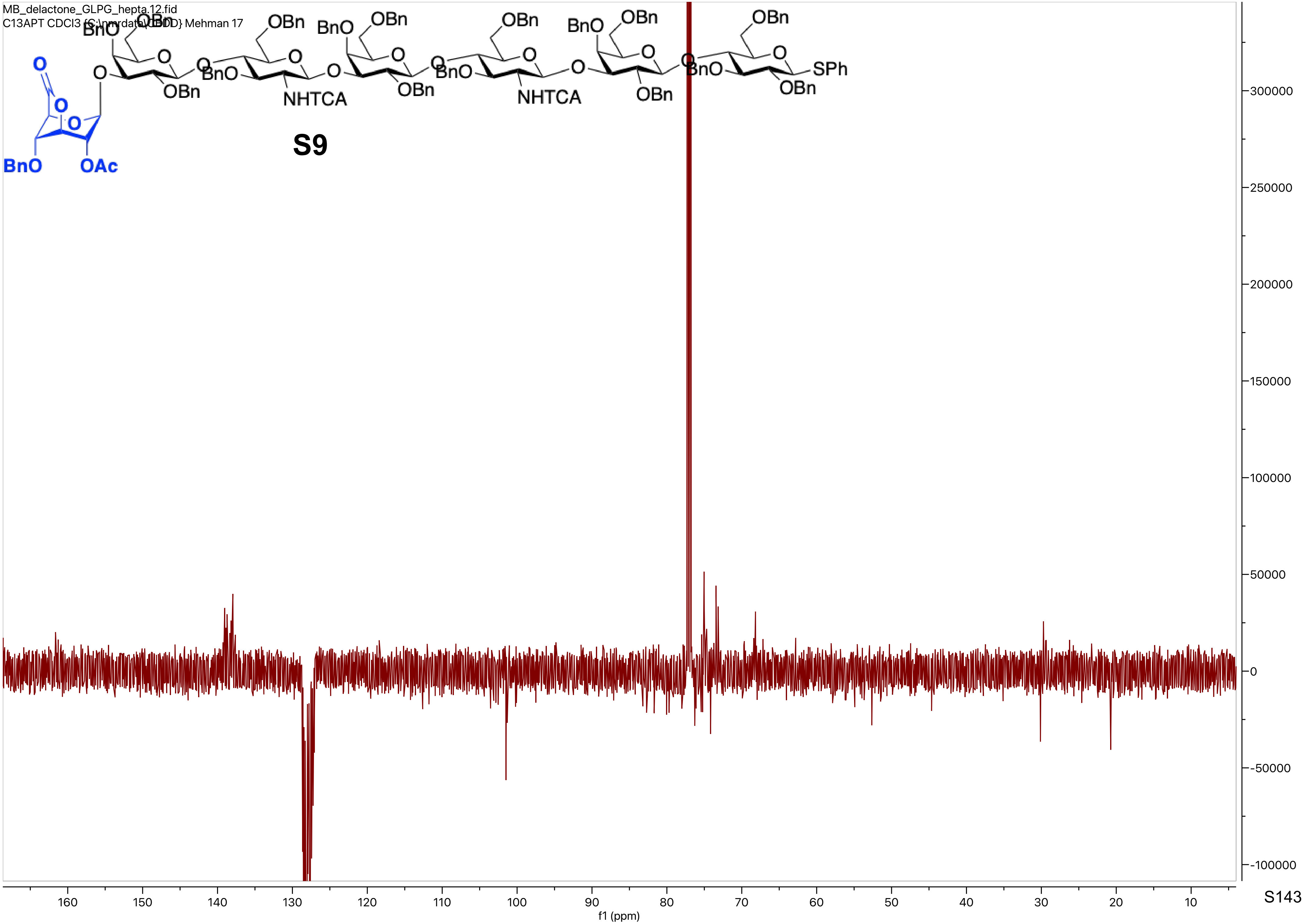

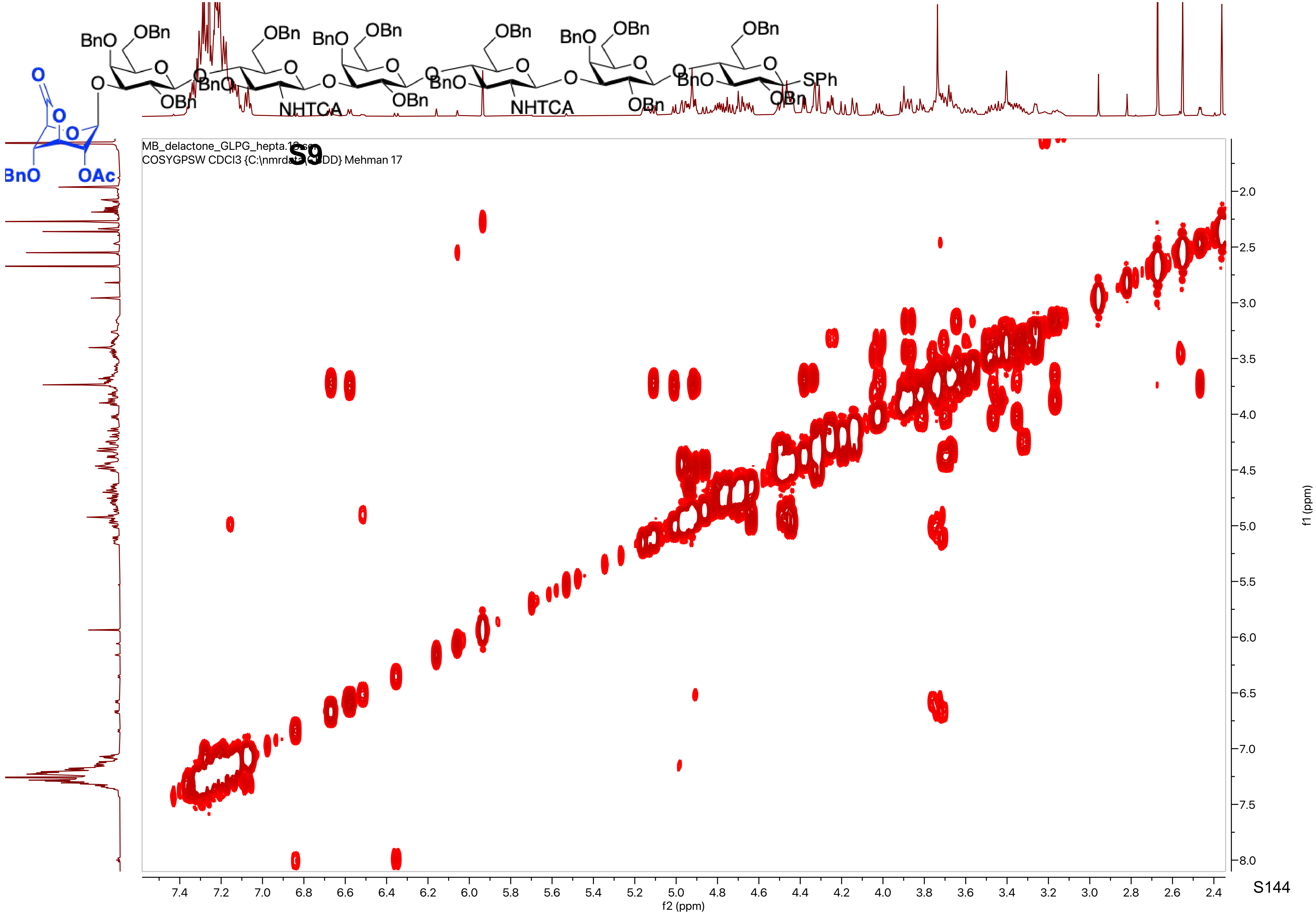
S9
COSYGPSW CDCl3
Mehman 17
f2 (ppm)
f1 (ppm)
BnO
OBn
OAc
NHTCA
SPh

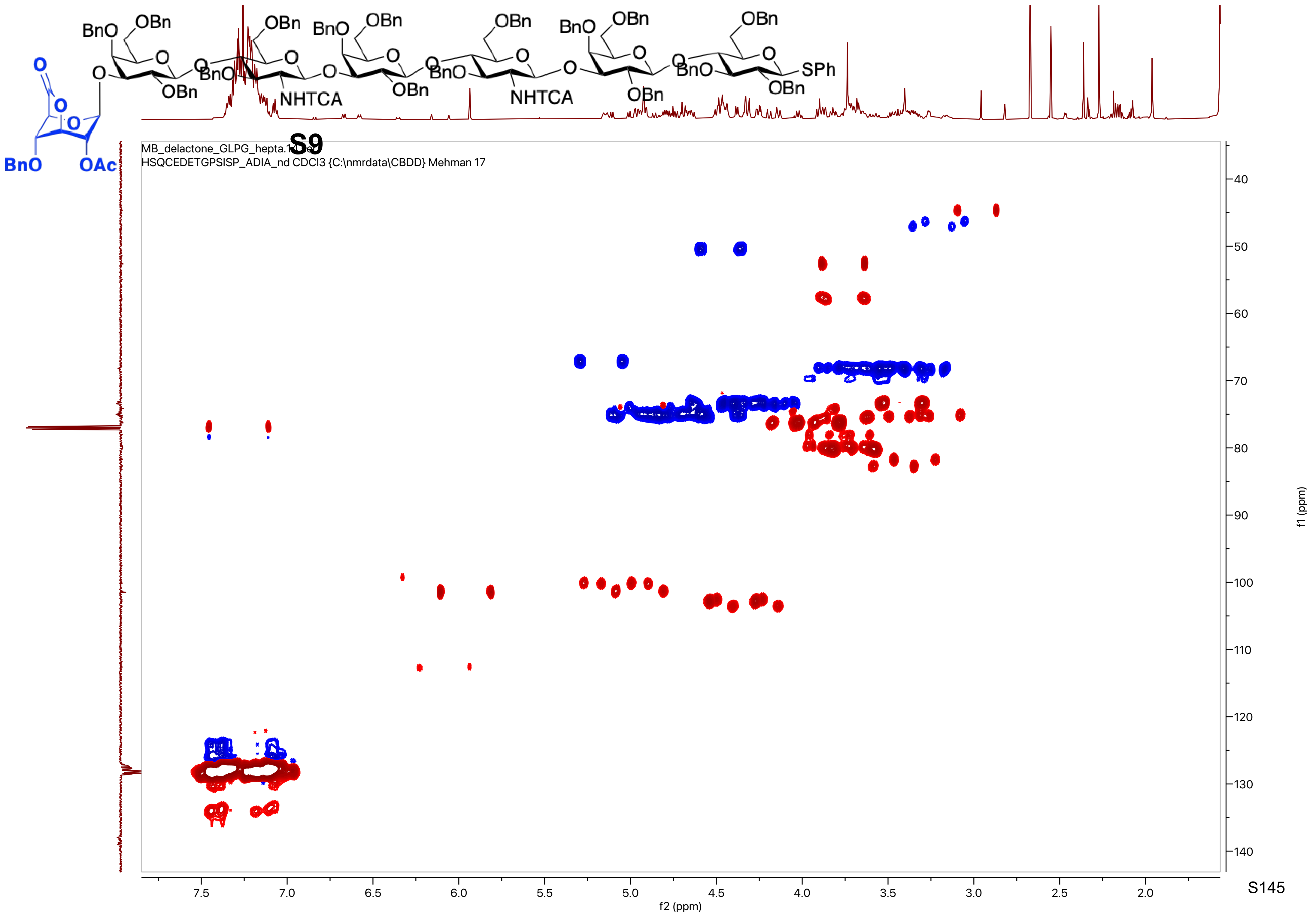
S9
MB_delactone_GLPG_hepta.1.fid
HSQCEDETGPSISP_ADIA_nd CDCl3 {C:\nmrdata\CBDD} Mehman 17
f2 (ppm)
f1 (ppm)
SPh
NHTCA
OAc

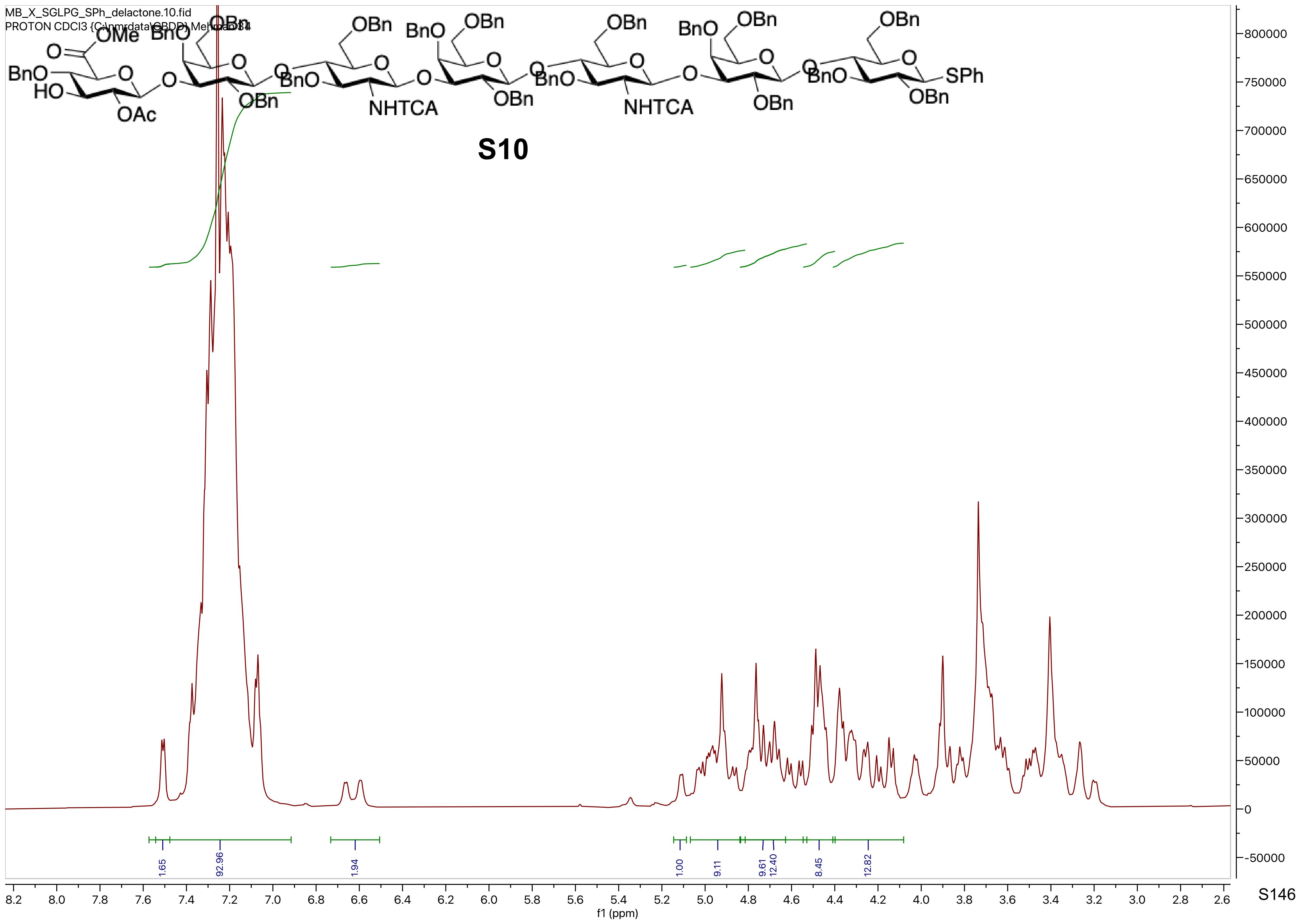

MB_X_SGLPG_SPh_delactone.10.fid
PROTON CDCl3 {C:\nmrdata\GBDD} Mehran 84
S10
f1 (ppm)

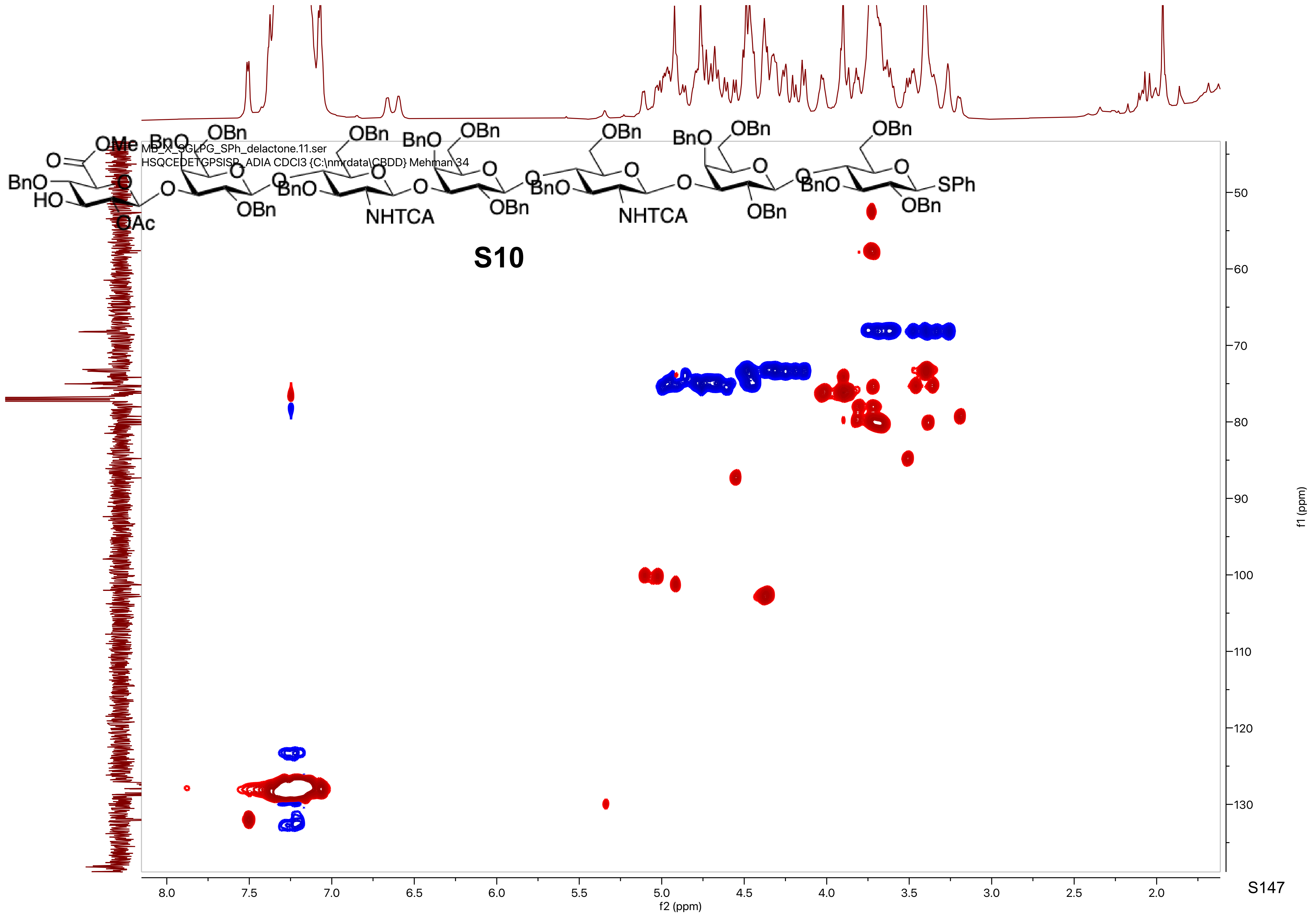

MB_X_SGLPG_SPh_delactone.11.ser
HSQCEDETGPSISP_ADIA CDCl3 {C:\nmrdata\CBDD} Mehman 34
S10
f2 (ppm)
f1 (ppm)

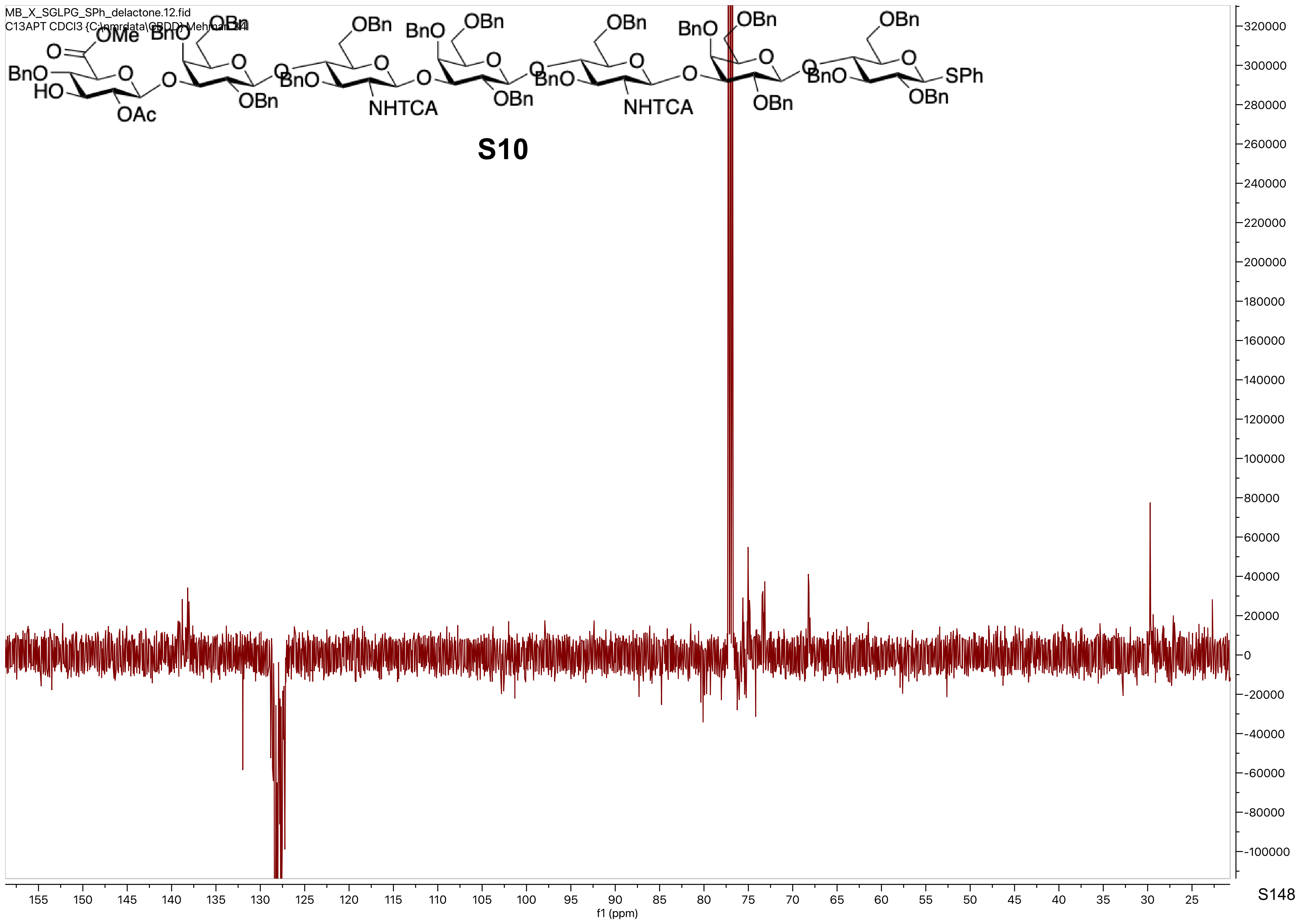
MB_X_SGLPG_SPh_delactone.12.fid
C13APT CDCl3 {C:\nmrdata\GBDD} Mehman 34
OMe
BnO
OBn
O
BnO
HO
OAc
NHTCA
SPh
S10
f1 (ppm)

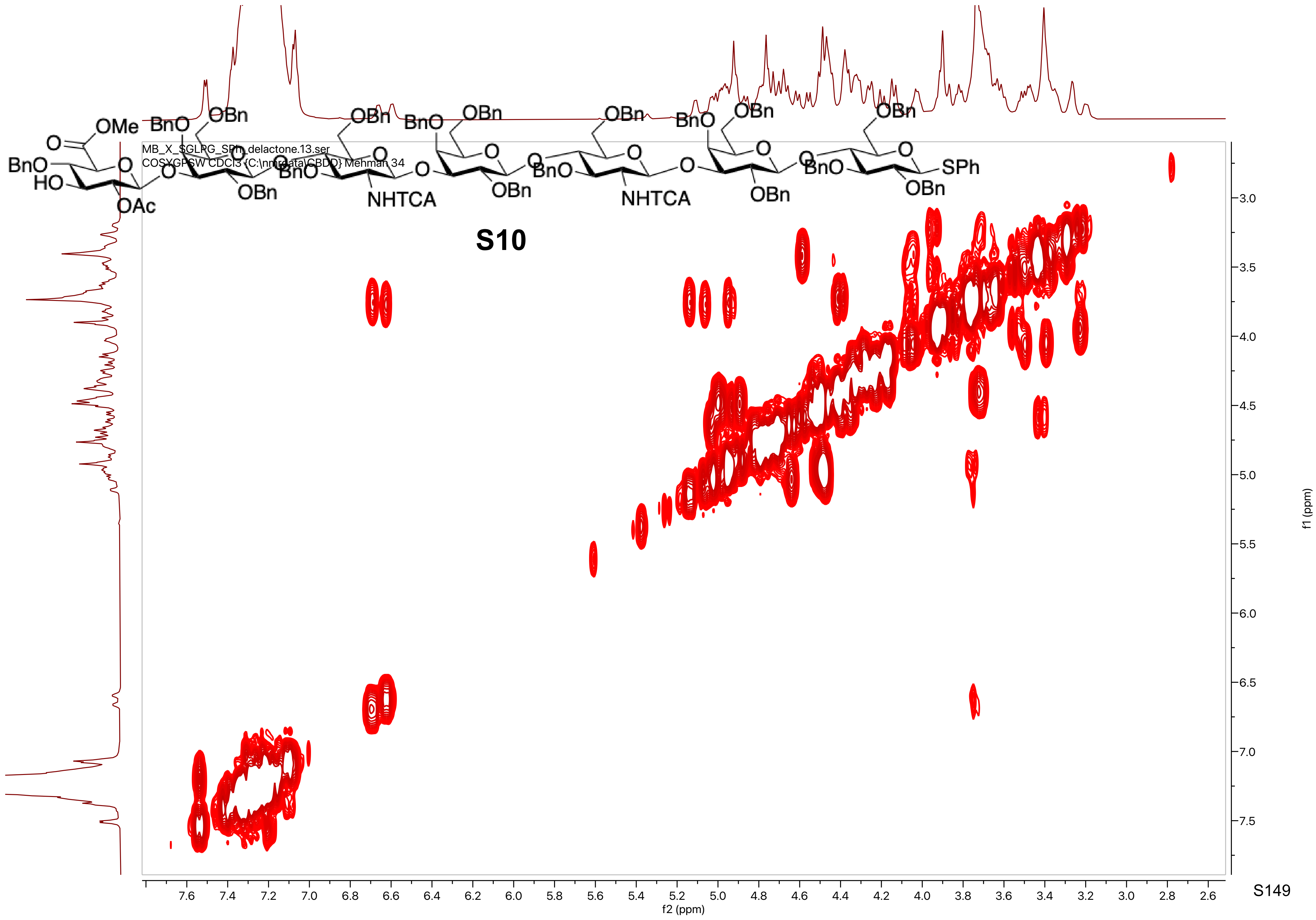

MB_X_SGLPG_SPh_delactone.13.ser
COSYGPSW CDCl3 {C:\nmrdata\CBDD} Mehman 34
OMe
O
BnO
HO
OAc
OBn
NHTCA
SPh
S10
f2 (ppm)
f1 (ppm)

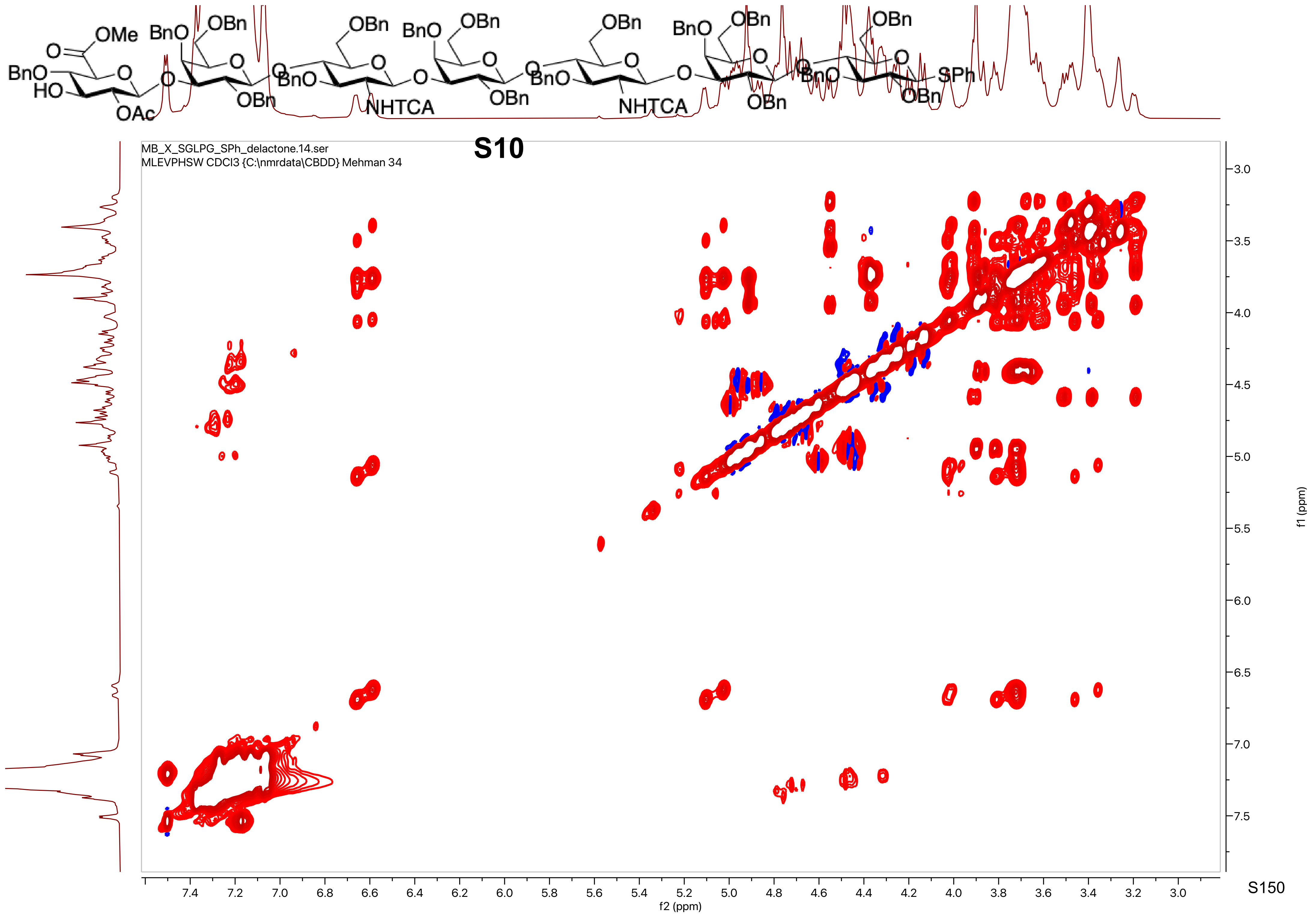
S10
MB_X_SGLPG_SPh_delactone.14.ser
MLEVPHSW CDCl3 {C:\nmrdata\CBDD} Mehman 34
f2 (ppm)
f1 (ppm)

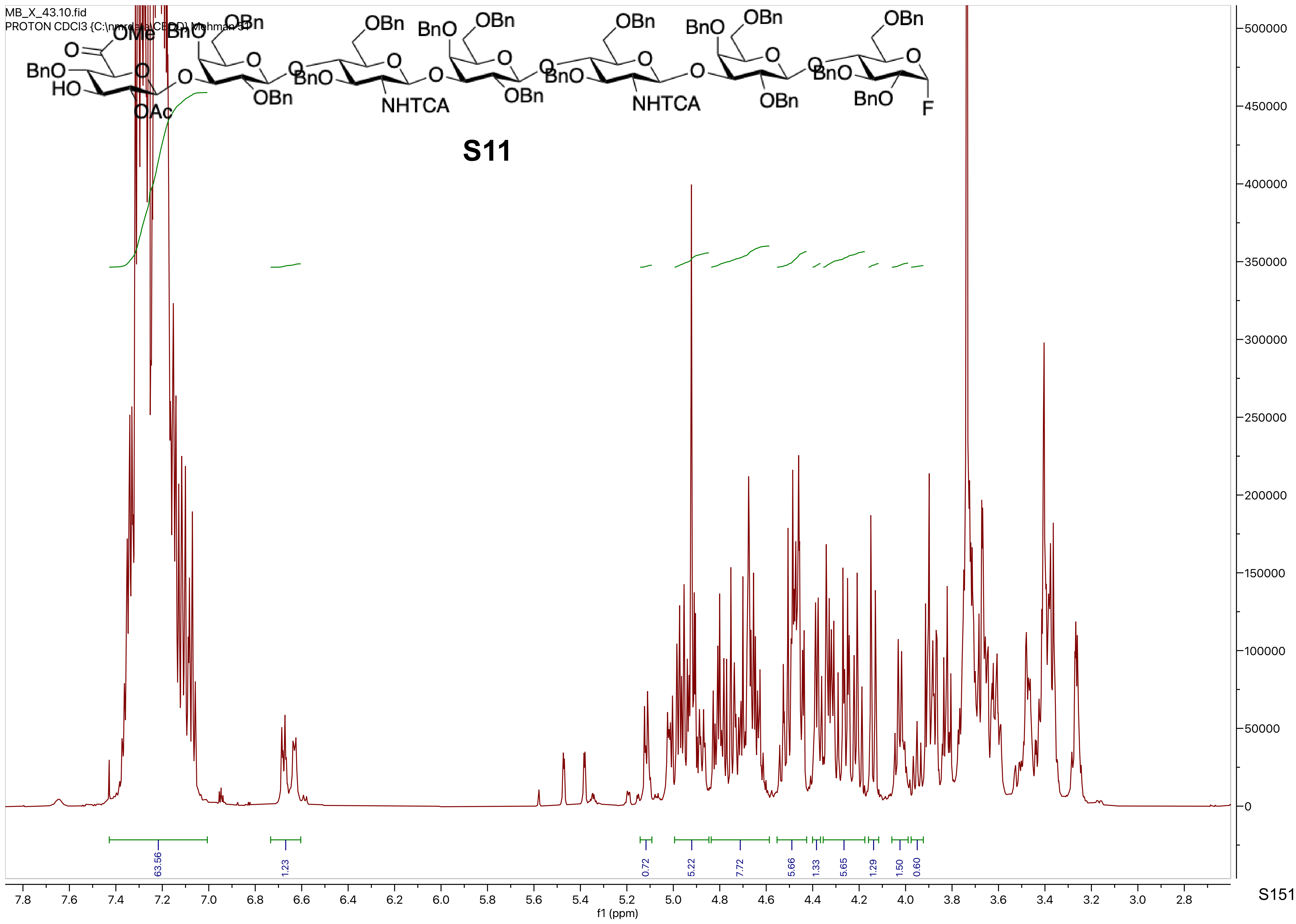
MB_X_43.10.fid
PROTON CDCl3 {C:\nmrdata\CEDD} Mehman 51
OMe
BnO
OBn
O
OAc
HO
NHTCA
F
S11
63.56
1.23
0.72
5.22
7.72
5.66
1.33
5.65
1.29
1.50
0.60
f1 (ppm)

MB_X_43.10.fid
PROTON CDCl3 {C:\nmrdata\CBDD} Mehman 51

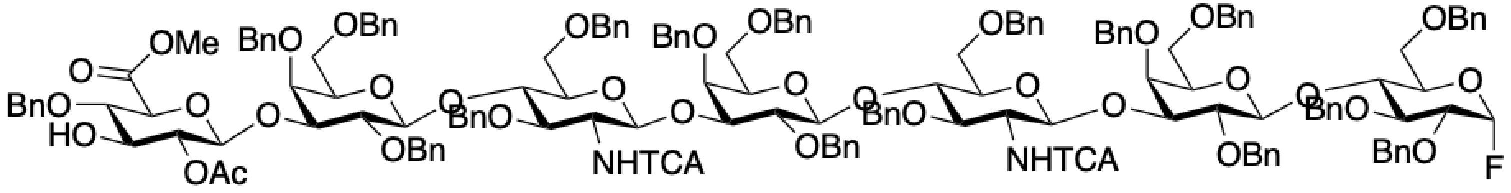


**S11**

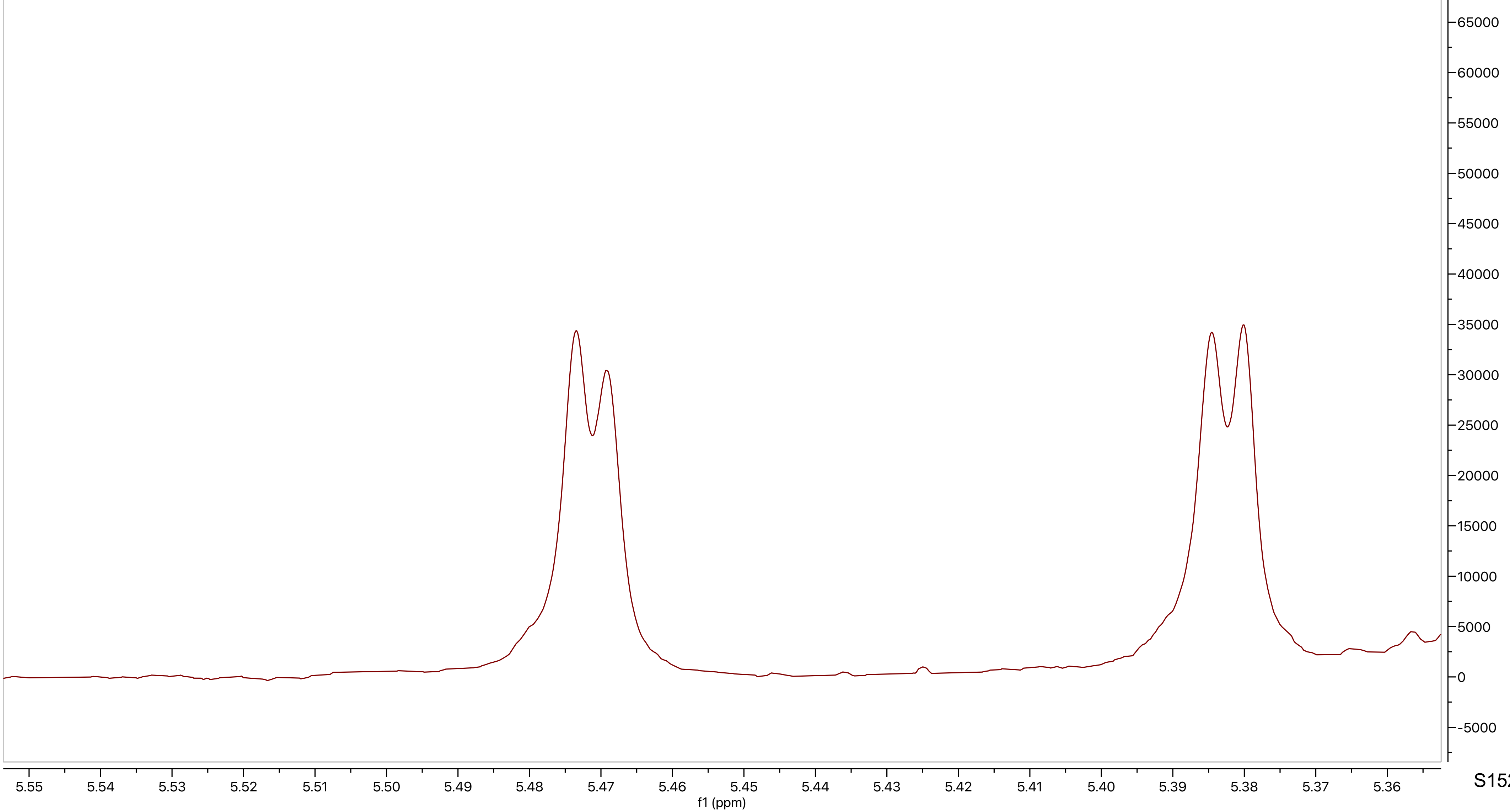

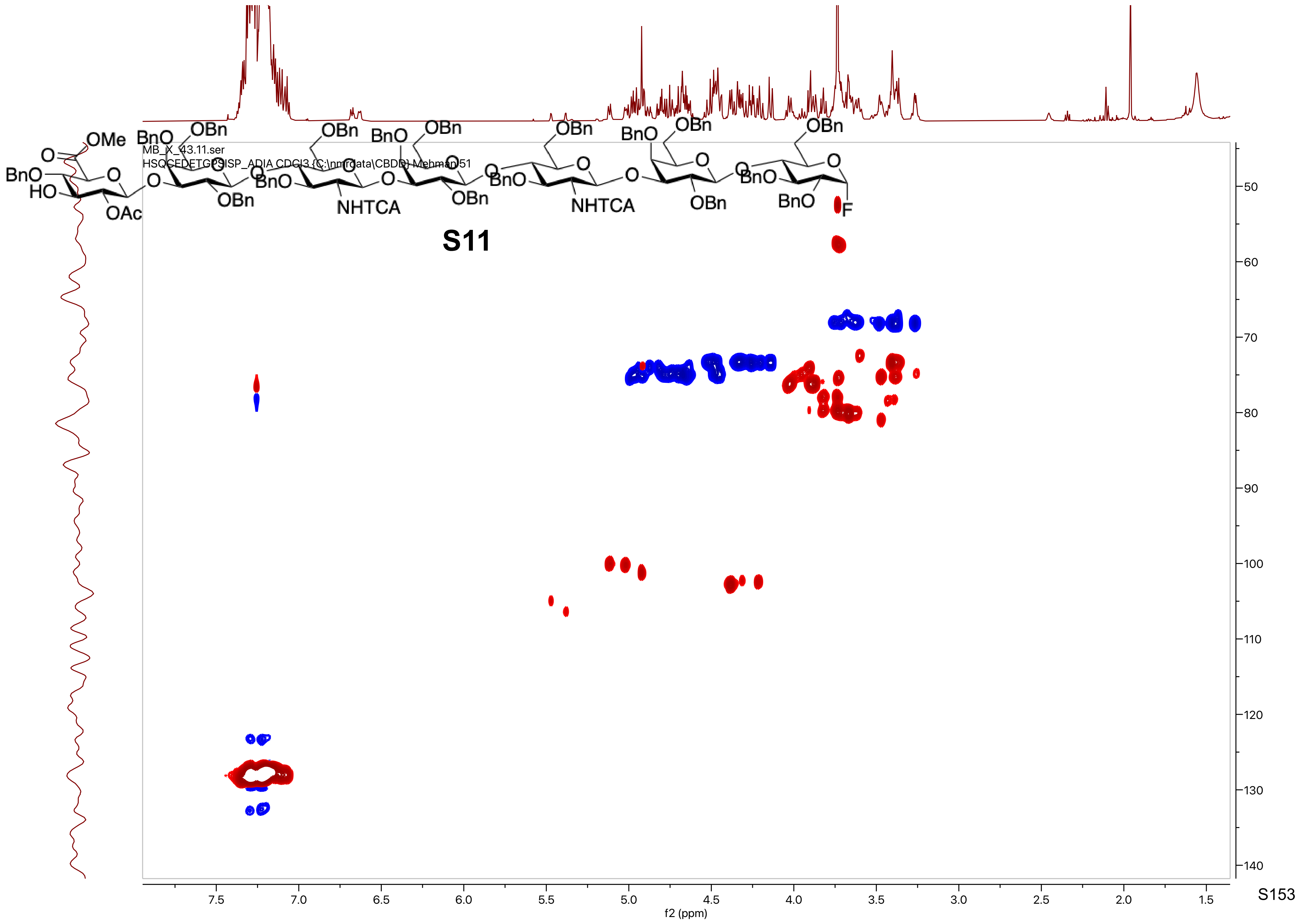
MB_X_43.11.ser
HSQCEDETGPSISP_ADIA CDCl3 {C:\nmrdata\CBDD} Mehman 51
S11
f2 (ppm)
f1 (ppm)

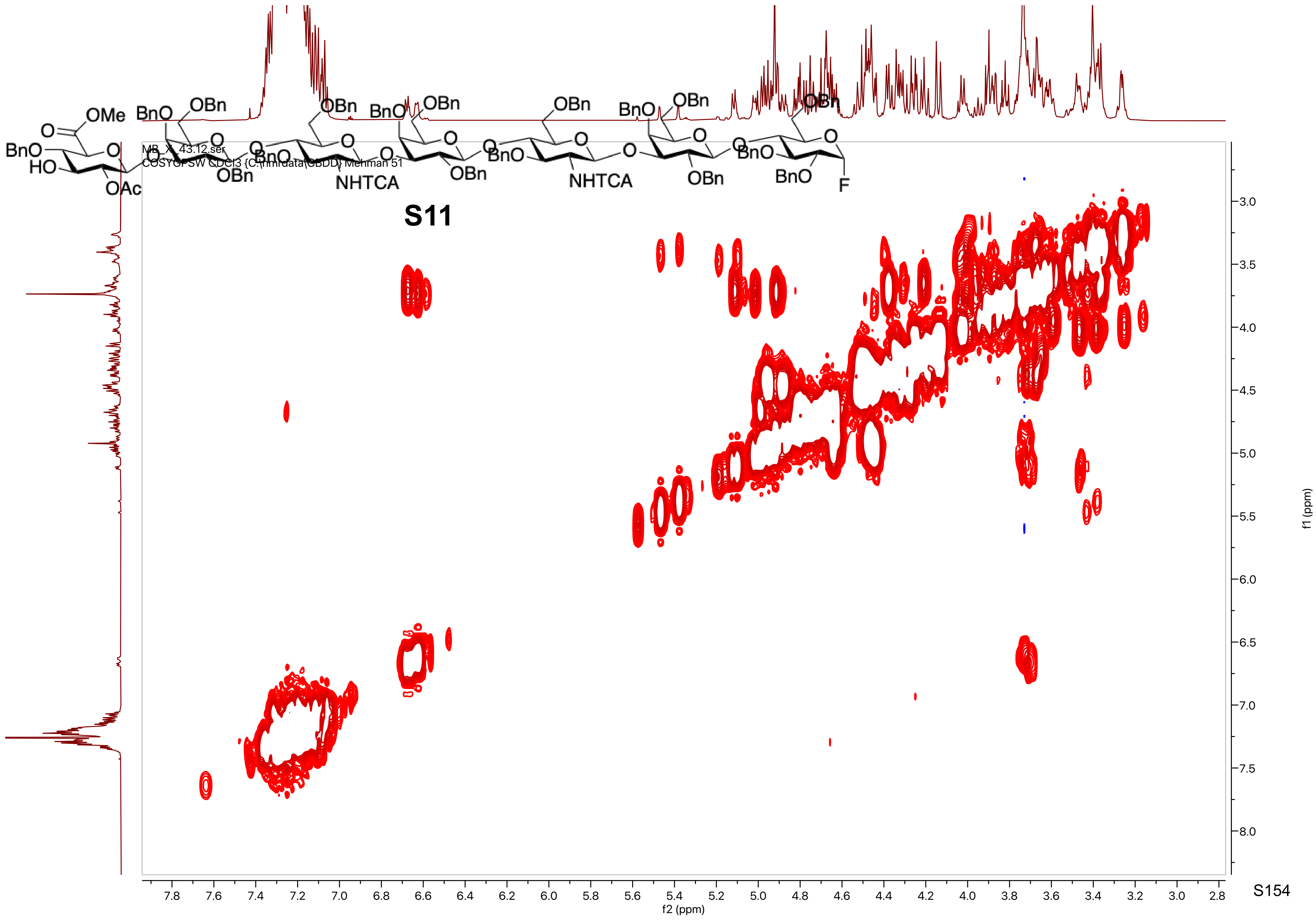

MB_XI_43.12.ser
COSYGPSW CDCl3 {C:\nmrdata\CBDD} Mehman 51
OMe
O
BnO
HO
OAc
OBn
BnO
NHTCA
F
S11
f2 (ppm)
f1 (ppm)

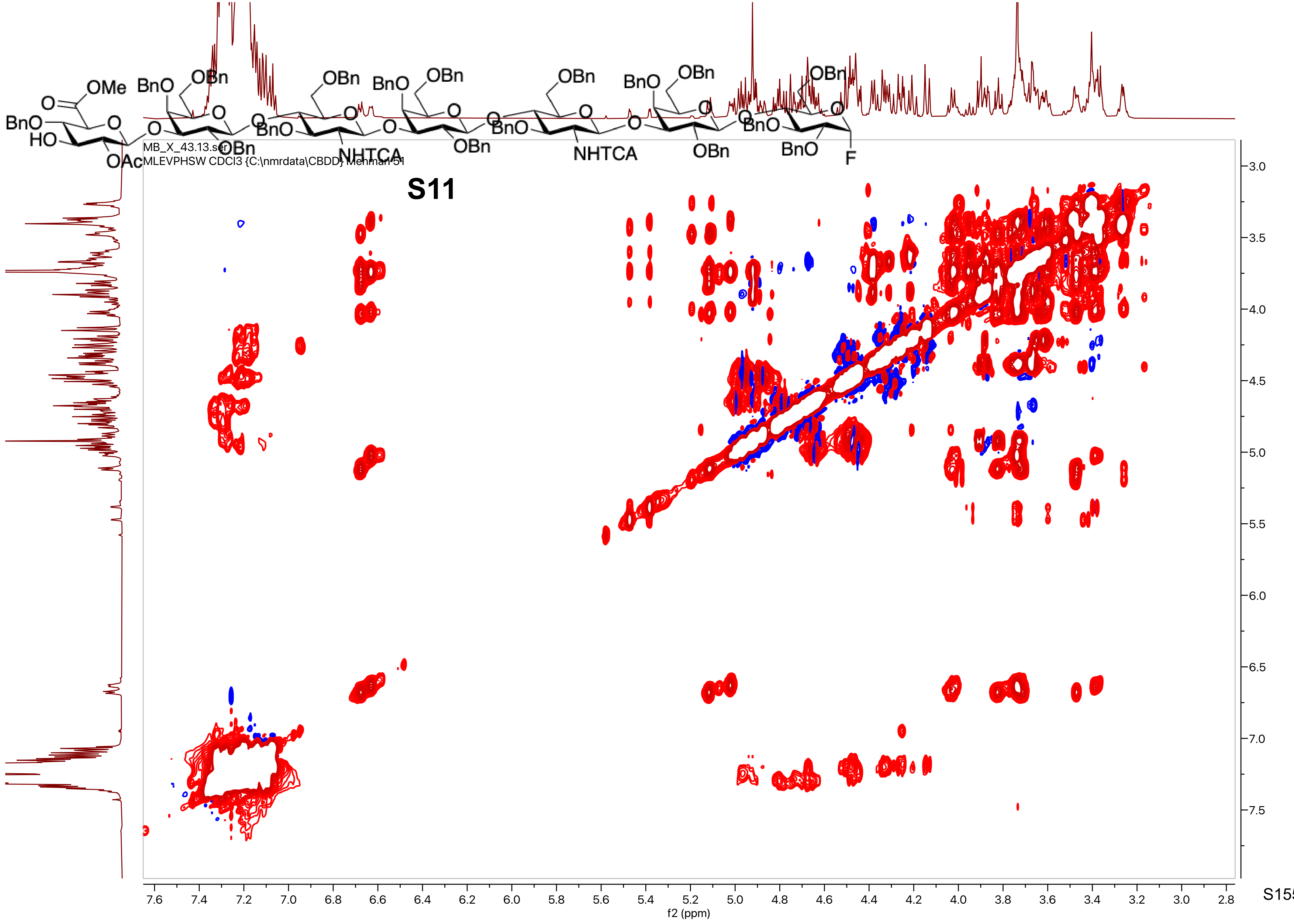

S11
MB_X_43.13.ser
MLEVPHSW CDCl3 {C:\nmrdata\CBDD} Mehman 51
f2 (ppm)
f1 (ppm)

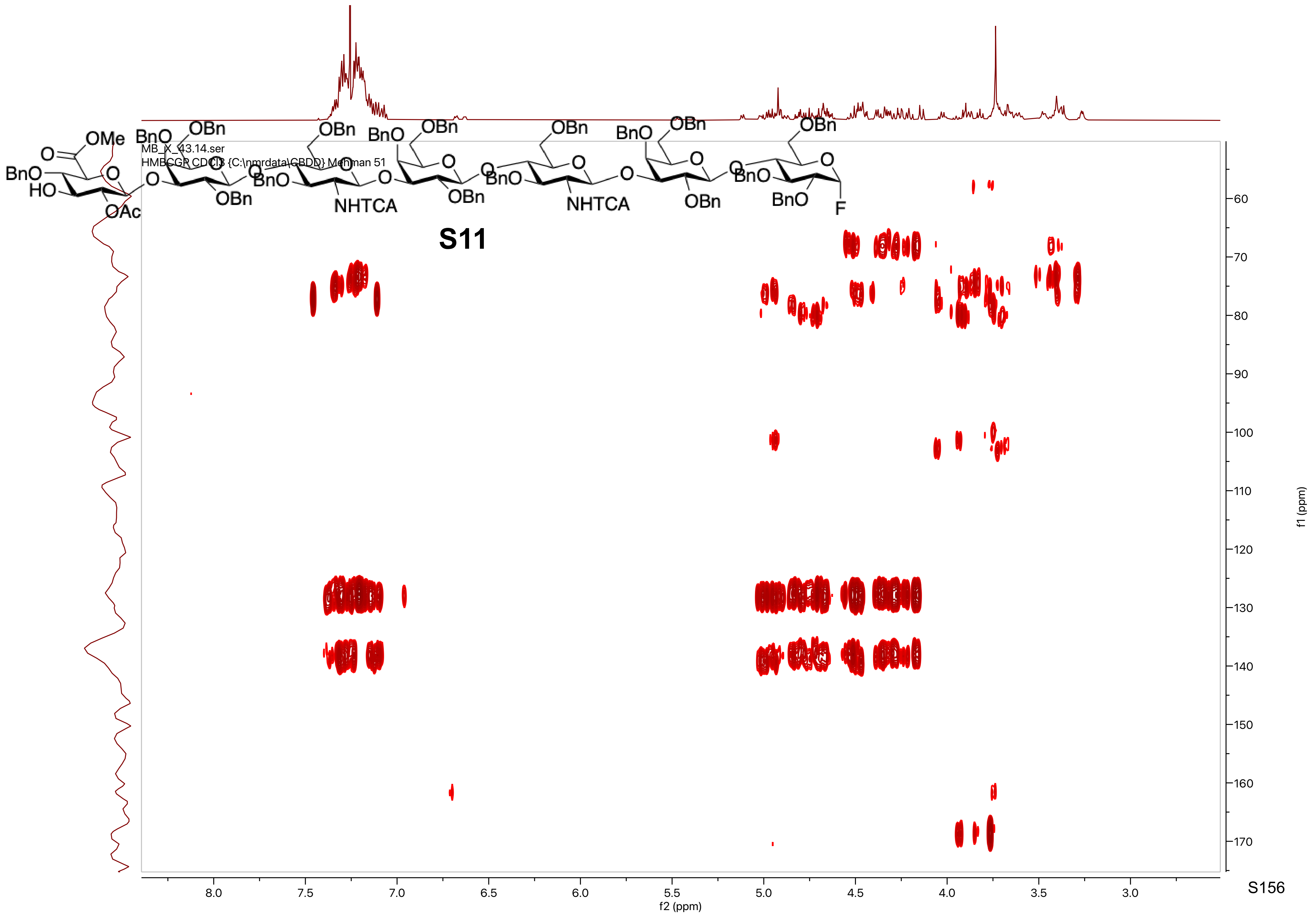
MB_X_43.14.ser
HMBCGP CDCl3 {C:\nmrdata\CBDD} Mehman 51
S11
f2 (ppm)
f1 (ppm)

MB_X_43.15.fid
F19 CDCl3 {C:\nmrdata\CBDD} Mehman 51

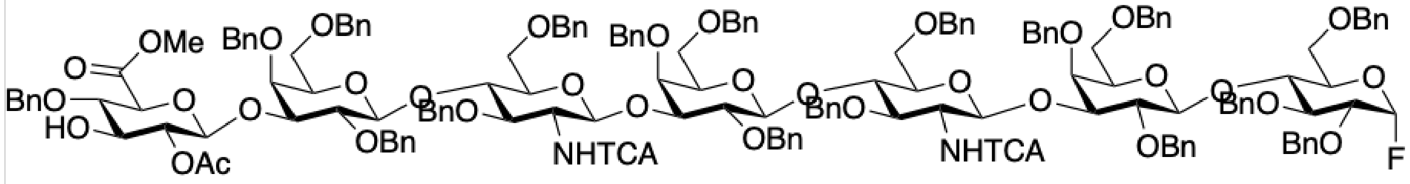

**S11**

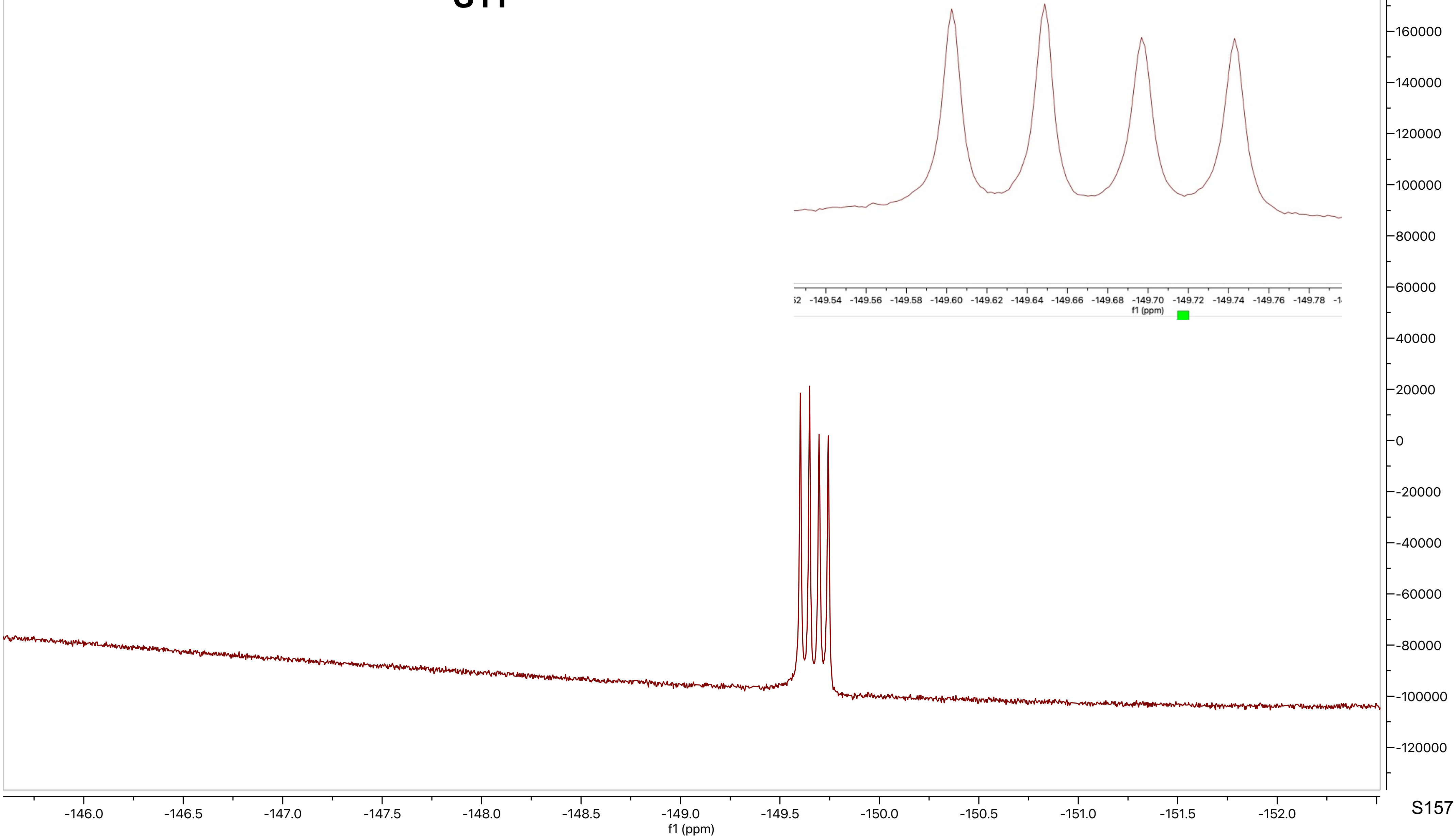

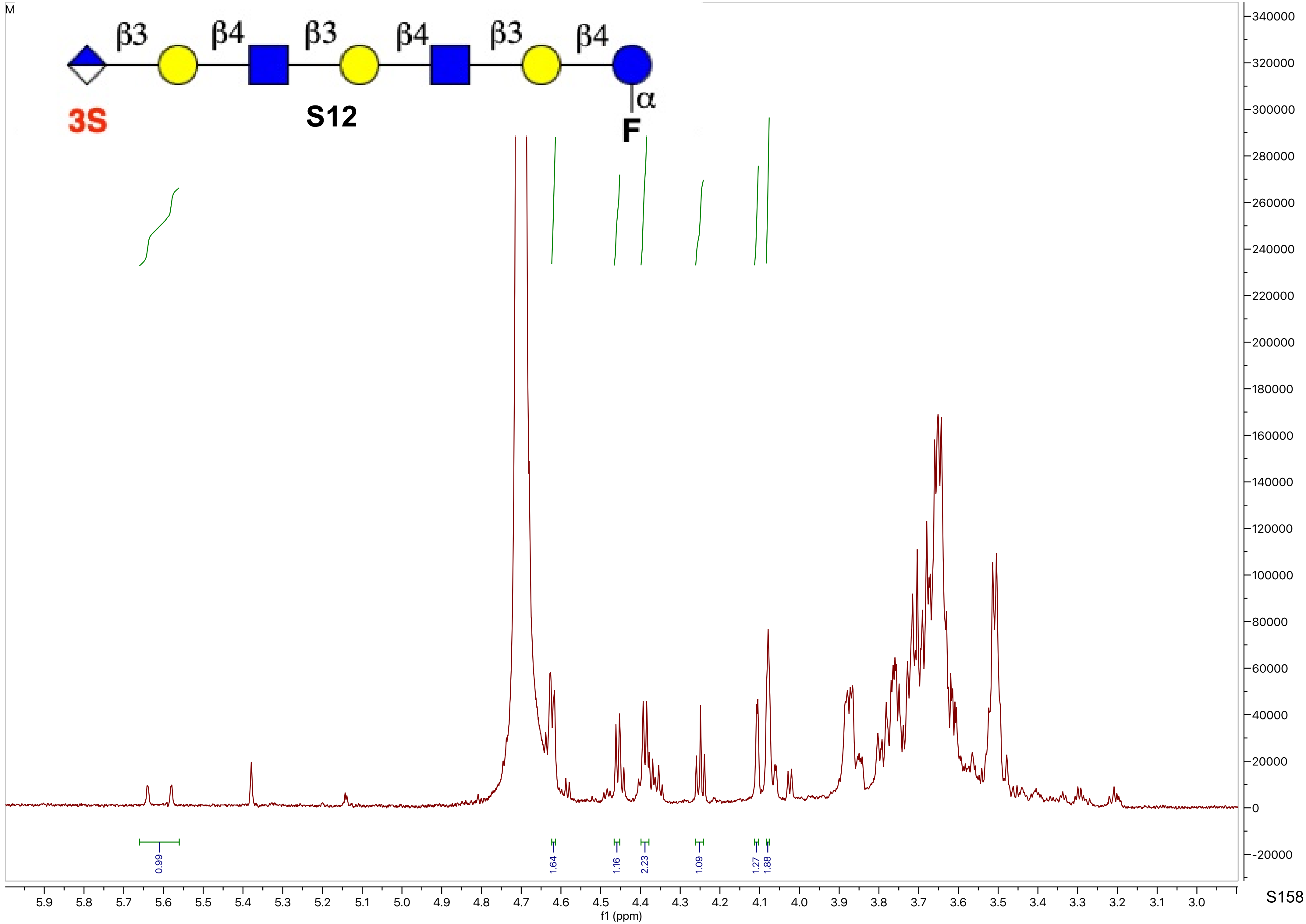

M
β3
β4
β3
β4
β3
β4
α
3S
S12
F
0.99
1.64
1.16
2.23
1.09
1.27
1.88
f1 (ppm)

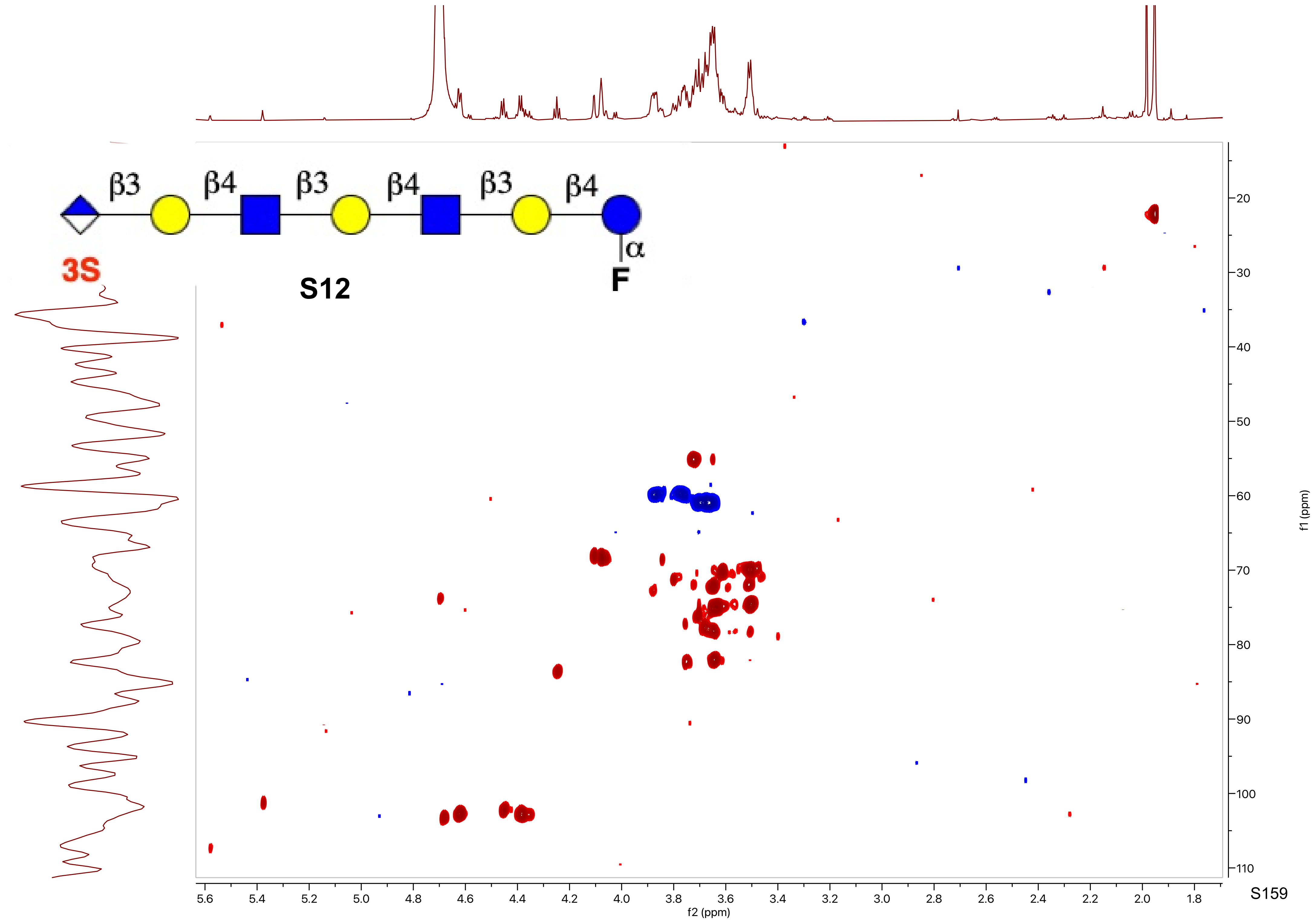
β3
β4
β3
β4
β3
β4
α
3S
F
S12
f2 (ppm)
f1 (ppm)

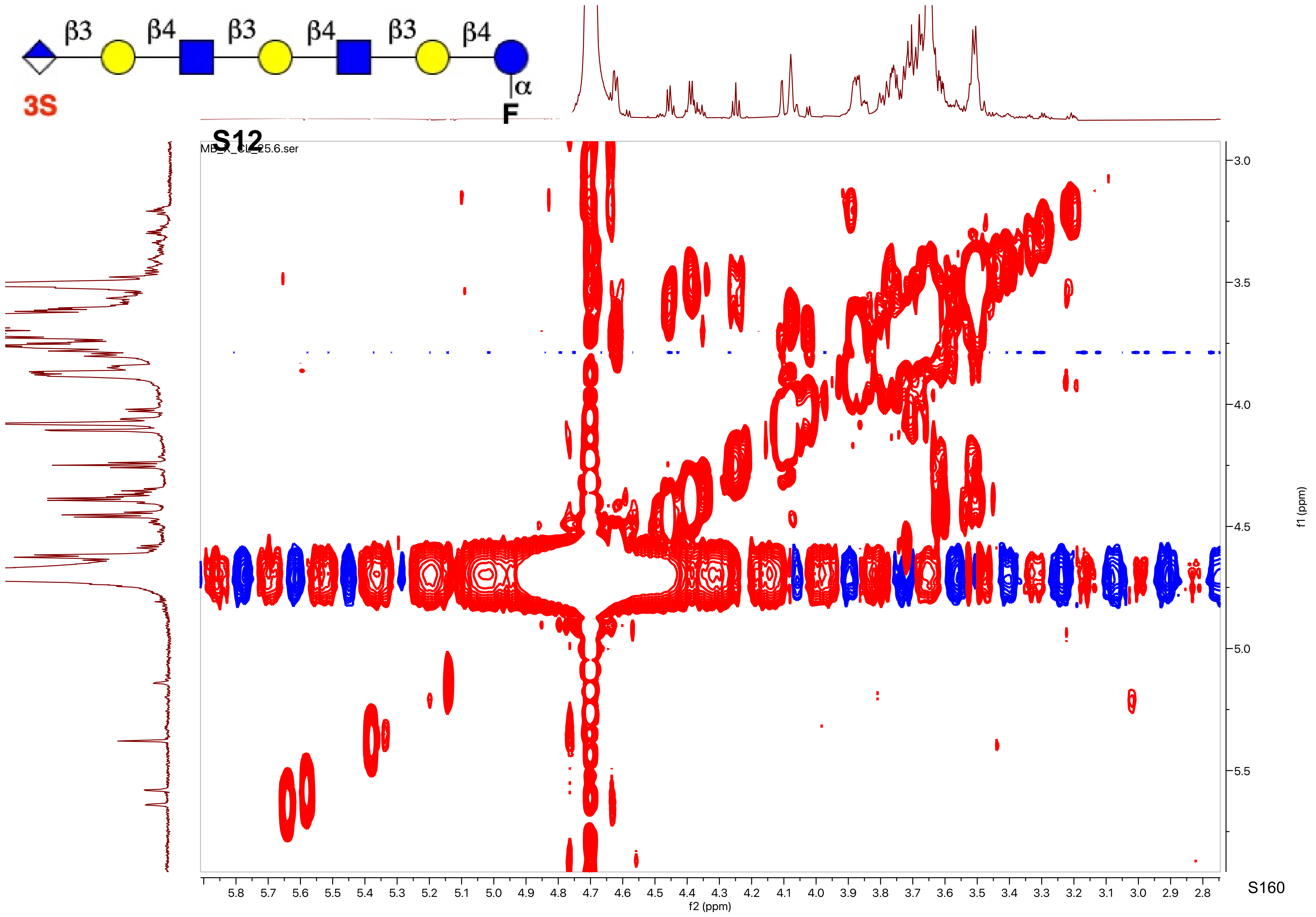
β3
β4
β3
β4
β3
β4
α
3S
F
S12
MB_X_CL_25.6.ser
f2 (ppm)
f1 (ppm)

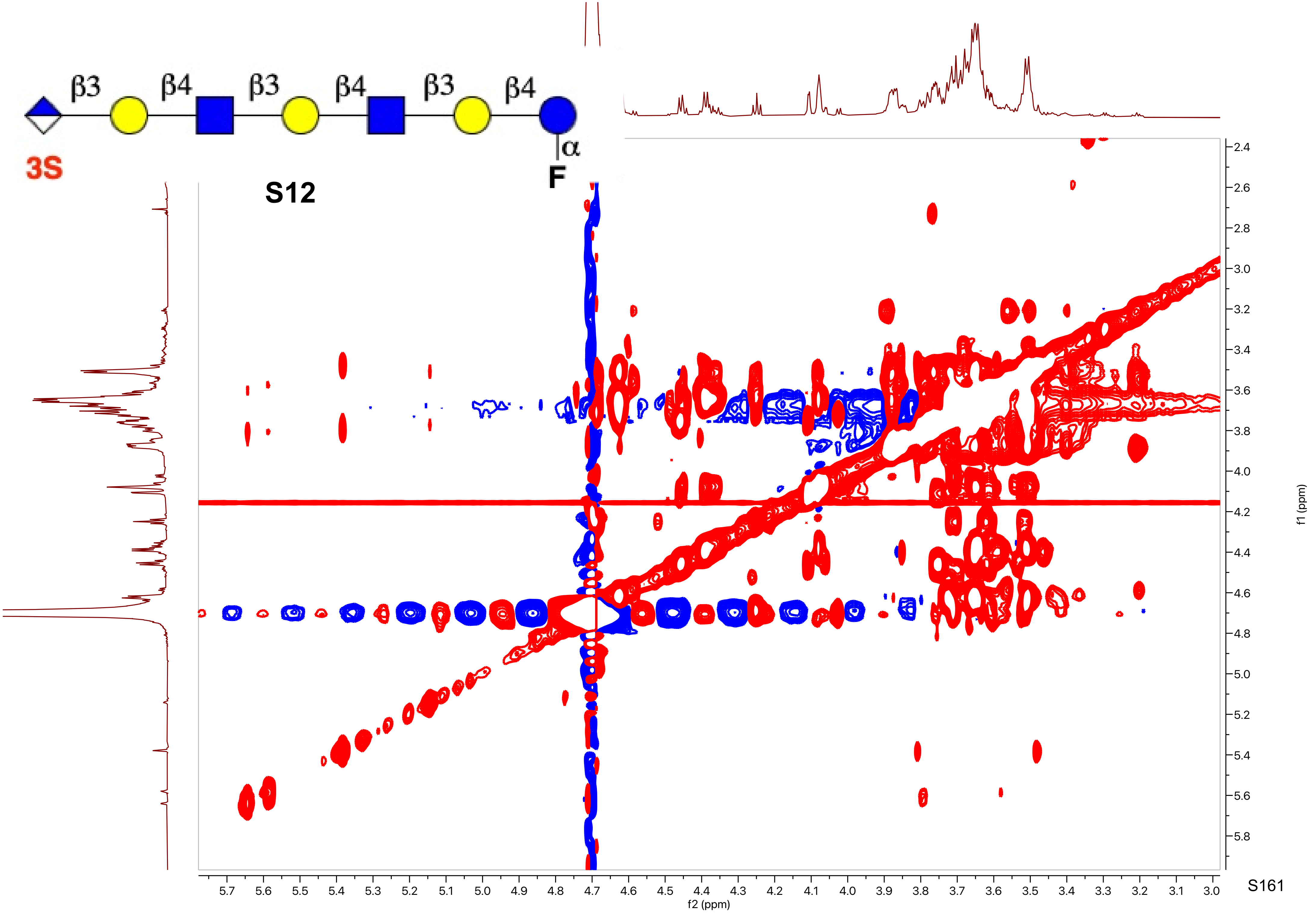
β3
β4
β3
β4
β3
β4
α
3S
F
S12
f1 (ppm)
f2 (ppm)

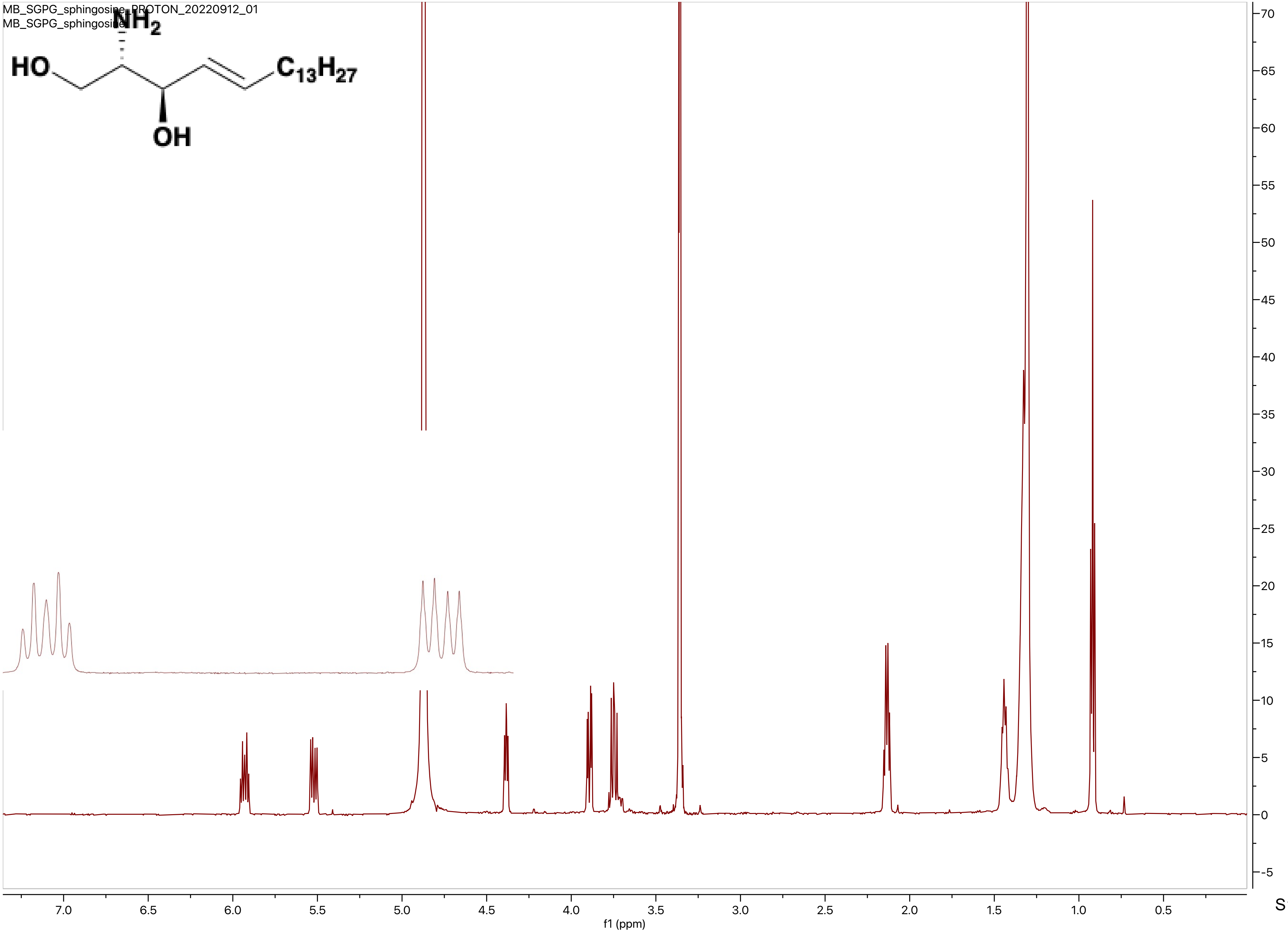
MB_SGPG_sphingosine_PROTON_20220912_01
MB_SGPG_sphingosine
NH2
HO
C13H27
OH
f1 (ppm)

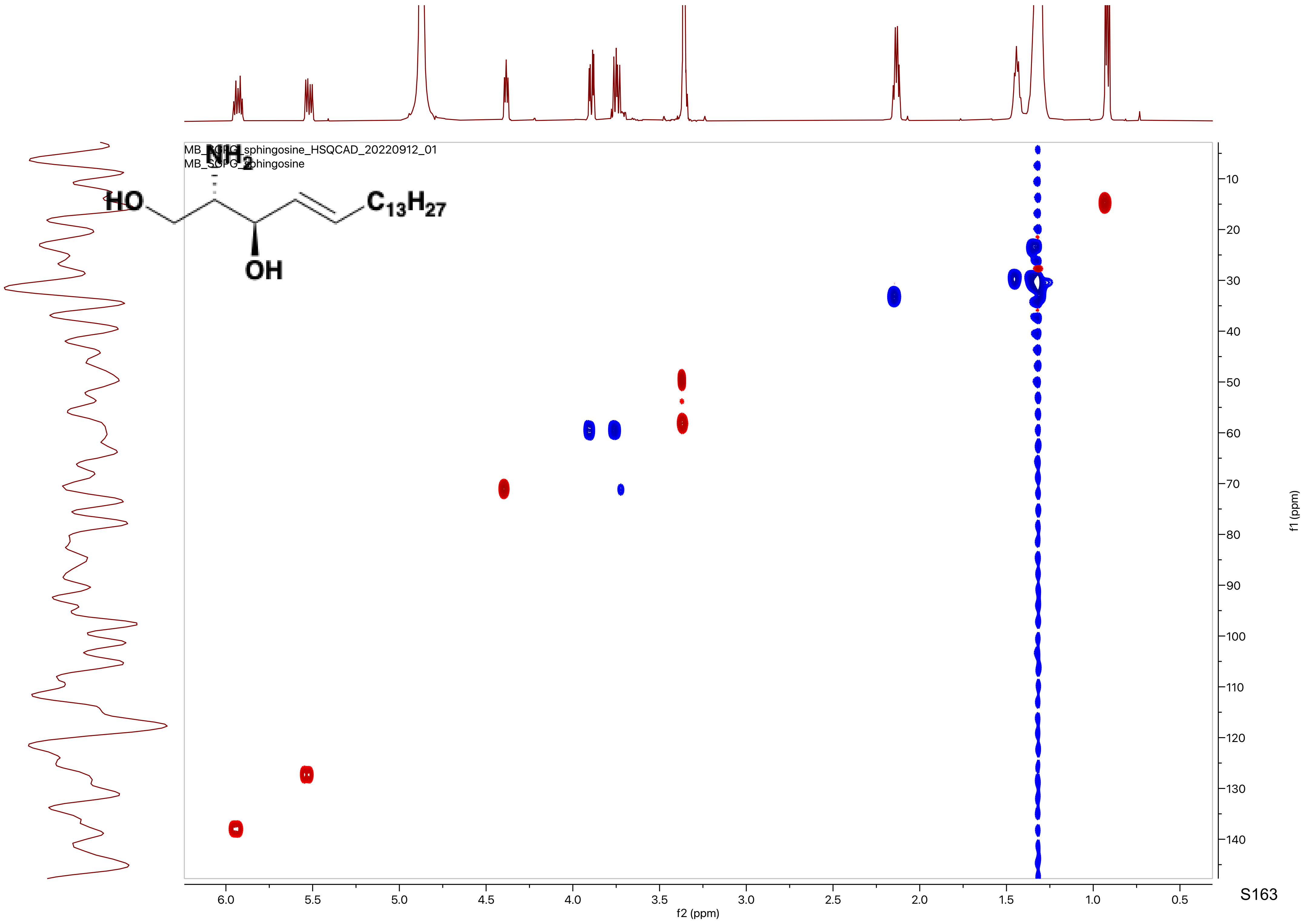
MB_SGPG_sphingosine_HSQCAD_20220912_01
MB_SGPG_sphingosine
NH2
HO
OH
C13H27
6.0
5.5
5.0
4.5
4.0
3.5
3.0
2.5
2.0
1.5
1.0
0.5
f2 (ppm)
10
20
30
40
50
60
70
80
90
100
110
120
130
140
f1 (ppm)
